# Lung derived extracellular vesicles have distinct pro-repair effects, depending on the age of their source tissue

**DOI:** 10.64898/2025.12.23.694250

**Authors:** Qi Chen, Paulson Tseng, Celia Diaz Nicieza, Fiona J. Culley, Charlotte H. Dean, Sally Yunsun Kim

## Abstract

Many age-related lung diseases encompass aberrant repair mechanisms. To develop novel treatments for these diseases, it is critical to understand how biology and repair signals are altered in aged lungs. Extracellular vesicles (EVs) carry bioactive signals reflecting their source tissue; therefore, we used EVs derived from precision-cut lung slices (PCLS) as tools to investigate differences in lung biology and repair signals that occur upon ageing and injury.

We compared the physicochemical properties and biological content of four groups of EVs, obtained from uninjured and injured, young and aged PCLS. Treatment with EVs obtained from young, uninjured (YU) PCLS decreased apoptotic cells in both young and aged spatially injured (AIR)-PCLS. YU EVs also increased the percentage of alveolar progenitor cells in aged AIR-PCLS, indicating that aged lungs retain the capacity to mount a repair response, if provided with the right biological cues.

Small RNA sequencing revealed the miRNA content of each EV group is altered, depending on the physiological conditions of their source tissue. Bioinformatic analysis of the target genes regulated by differentially expressed miRNAs identified enrichment of genes required for lung development and repair.

These findings pave the way for future therapeutic strategies to repair or regenerate aged lungs.

## Introduction

Extracellular vesicles (EVs), are nanoparticles that transfer biological information between cells, by encapsulating proteins, nucleic acids and lipids^1^. In the past decade, EVs have been shown to play key roles in tissue repair ^2^, and various studies and trials have explored their potential as cell-free therapeutics for numerous conditions^3–6^. In the lungs, EVs have shown potential for the treatment of lung diseases by promoting repair and regeneration^7^.

The majority of these investigations utilised mesenchymal stromal/stem cell-derived EVs, due to their immunomodulatory effects. However, given that the composition and function of EVs vary depending on their parent cells, numerous alternative sources of EVs have also been investigated for potential beneficial effects on pulmonary diseases including amnion epithelial cells^8^, lung spheroids^9^ and primary human lung fibroblasts^10–12^.

On the other hand, in mouse models of lung injury, EVs released from lung cells upon stress or infection may contribute to pathophysiology, through the capacity to transmit pathological cargo, propagating pulmonary inflammation and injury^13^. For example, pathogenic neutrophil and macrophage derived EVs obtained from cigarette smokers were sufficient to induce emphysema in naïve mice^14^. More recently, EVs secreted by primary adult lung fibroblasts obtained from control mice were shown to worsen bleomycin-induced fibrosis in vivo^15^, but EVs isolated from healthy human lung fibroblasts attenuate elastase-induced lung injury in ex vivo and in vivo mouse models^10^. These differing findings highlight that the biological cargo of EVs vary, depending on the physiological conditions of the cells from which they are sourced. Since EVs are released from almost all cell types as well as from the extracellular matrix (ECM), EVs obtained from precision-cut lung slices, a three-dimensional (3D) model in which the native tissue architecture and ECM are retained, provide a useful tool to understand both the biology of EVs and their role in lung repair.

The lungs are relatively quiescent during homeostasis, but they have significant intrinsic capacity for self-repair upon injury, following which progenitor/stem cells become activated to help repair the damaged tissue^16^. Ageing is a known risk factor for various lung diseases such as chronic obstructive pulmonary disease (COPD) and idiopathic pulmonary fibrosis (IPF)^17,18^. In aged lungs, repair and regeneration is dysregulated and attenuated^19,20^. The lung resident stem/progenitor cell reservoir is depleted, demonstrated by a decrease in quantities of airway basal and club cells, along with reduced self-renewal and differentiation capacity of the alveolar epithelial progenitors, alveolar type II (ATII) cells^21–23^. Other ageing-related physiological changes such as increased cellular senescence and disrupted cytoskeletal integrity also contribute to impaired regenerative capacity^18^. Ageing particularly affects the composition and physical properties of the ECM and in turn, altered ECM leads to dysregulated cell behaviour, for example increased fibroblast proliferation and impaired differentiation of alveolar epithelial cells^24^. Aged lungs are unable to fully regenerate functional tissue and instead compensate for tissue damage with fibrotic repair, especially in chronically injured tissue when regenerative mechanisms are exhausted^25^. The altered mechanics in aged and injured lungs impact pharmacological responsiveness^26^.

To date, the biology of EVs obtained from young and aged lungs has not been investigated. Given the extensive biological differences between young and aged lungs, we hypothesised that EVs obtained from young and aged lungs would reflect the differences in their source tissue and would therefore vary in their composition and functional capabilities. Since EV-mediated cellular crosstalk plays an active role in lung homeostasis and regeneration^7^, we reasoned that determining the composition and effects of young and aged lung derived EVs would be important for future therapeutic strategies to treat age-related lung diseases. To address this knowledge gap, we isolated EVs secreted from precision-cut lung slices (PCLS) obtained from young or aged mouse lungs and investigated their content and their effects on lung repair and regeneration.

## Results

### Characterisation of EVs derived from uninjured or acid-injured PCLS obtained from young and aged mice

To compare the properties of lung tissue derived EVs investigated in this study, we obtained four distinct groups of EVs: EVs isolated from uninjured PCLS obtained from young or aged mouse lungs, and EVs isolated from young or aged PCLS briefly injured with 0.1 M HCl acid. All EVs were isolated from the same number of PCLS of similar size. These four groups of EVs are named according to their source PCLS and are hereby referred to as (1) young, uninjured EVs (“YU EVs”), (2) aged, uninjured EVs (“AU EVs”), (3) young, injured EVs (“YI EVs”) and (4) aged, injured EVs (“AI EVs”) (Fig. 1a).

**Figure 1.**
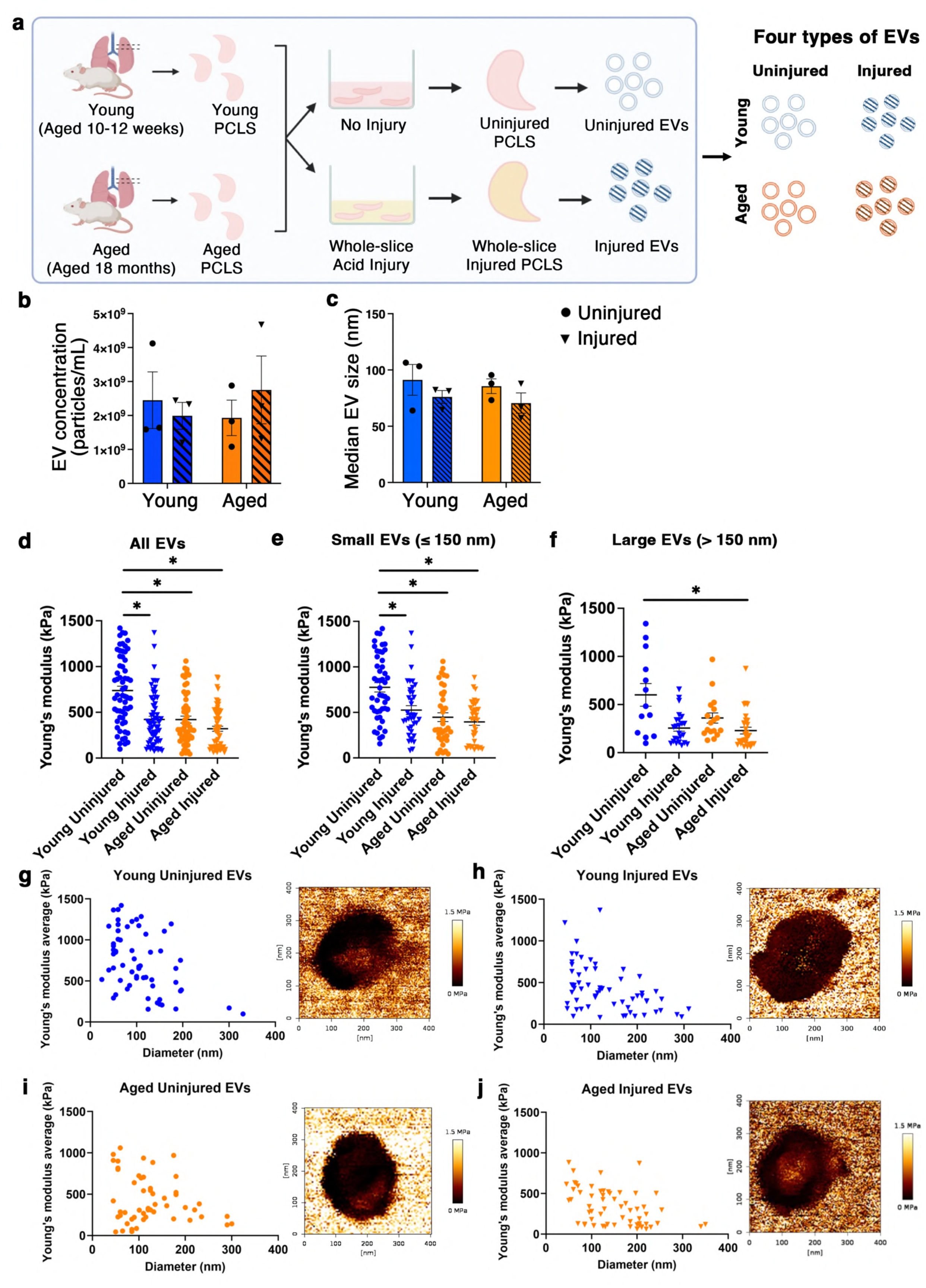
Characterisation of four types of EVs derived from uninjured or acid-injured PCLS obtained from young and aged mouse lungs. (**a**) Extracellular vesicles (EVs) were isolated from precision cut lung slices (PCLS) obtained from young or aged mouse lungs and then left uninjured or injured by briefly treating with 0.1 M HCl acid for 1 min, resulting in four groups of EVs: young uninjured (YU), young injured (YI), aged uninjured (AU) and aged injured (AI) EVs. (**b, c**) The four groups of EVs were similar in yield (**b**) and particle size (**c**). (**d-f**) Young’s modulus for the four groups of EVs measured using atomic force microscopy for: all EVs regardless of size (**d**) and smaller EVs (≤ 150nm, **e**) and larger EVs (>150nm, **f**). (**g-j**) Average Young’s modulus for each EV according to the diameter of EVs are presented for each of the four groups of EVs. Data are presented as mean ± S.E.M. *N* = 3. Two-way ANOVA test with Tukey’s multiple comparisons test with *p* < 0.05 as significant.

Nanoscale flow cytometry revealed similar quantities and particle sizes among the four groups of EVs (Fig. 1b, c). Average EV concentrations ranged between 1.93 × 10^9^ and 2.75 × 10^9^ particles/mL (Fig. 1b). The median EV size ranged from 70.58 nm to 91.22 nm, with the majority of EVs measuring between 50 nm and 120 nm in diameter (Fig. 1c).

The mechanical properties of EVs were determined using atomic force microscopy, to quantify stiffness of EVs measured in liquid. Depending on the scan size, which varied from 400 nm to 1.5 mm, typically one to ten EVs sized between 40-350 nm were captured in each field of view. The Young’s modulus of EVs was calculated by averaging the values obtained in the middle region of each EV using the JPK Data Processing software, processed with the Hertz model. The Young’s modulus of EVs averaged from four biological replicates are presented in Fig. 1d. Upon injury, the mean stiffness values reduced by 296.8 kPa for young EVs (from 687.3 kPa for YU EVs to 390.5 kPa for YI EVs, Fig. 1d) and by 72.3 kPa for aged EVs (from 388.0 kPa for AU EVs to 315.7 kPa for AI EVs, Fig. 1d). Then we investigated the Young’s modulus mean values relative to the diameter of EVs (Fig. 1e-j). Despite high levels of heterogeneity, there was a trend that the smaller EVs (≤150nm) had higher stiffness compared to larger EVs (>150nm). This difference was the highest in YI EVs, in which the average stiffness of larger EVs was 256.1 kPa and the mean stiffness of smaller EVs was 525.6 kPa. For YU, there was a smaller difference in stiffness between larger EVs (mean = 600.2 kPa) and smaller EVs (mean = 775.6 kPa). For aged EVs, the difference in the mean stiffness of smaller EVs and larger EVs was also greater in the injured group (396.1 kPa for smaller AI EVs vs. 229.3 kPa for larger AI EVs) compared to the uninjured group (447.4 kPa for smaller AU EVs vs. 360.2 kPa for larger AU EVs).

### Comparison of young and aged adult PCLS reveals key hallmarks of ageing and injury

As a prelude to investigating the effects of EVs on PCLS, we first sought to determine whether key hallmarks of ageing and injury could be detected in PCLS. To do this we chose to examine critical components of the ECM and the cytoskeleton because dysregulation of these key cell/tissue components are associated with ageing and injury. Four different groups of PCLS were generated: (1) young, uninjured PCLS, (2) young, injured PCLS, (3) aged, uninjured PCLS, and (4) aged, injured PCLS (Fig. 1a). Injury was induced by briefly treating whole PCLS with 0.1 M HCl acid (whole-slice injury) (Fig. 2a). Young and aged PCLS were cultured for 48 hrs following acid injury and then compared to their uninjured counterparts.

**Figure 2.**
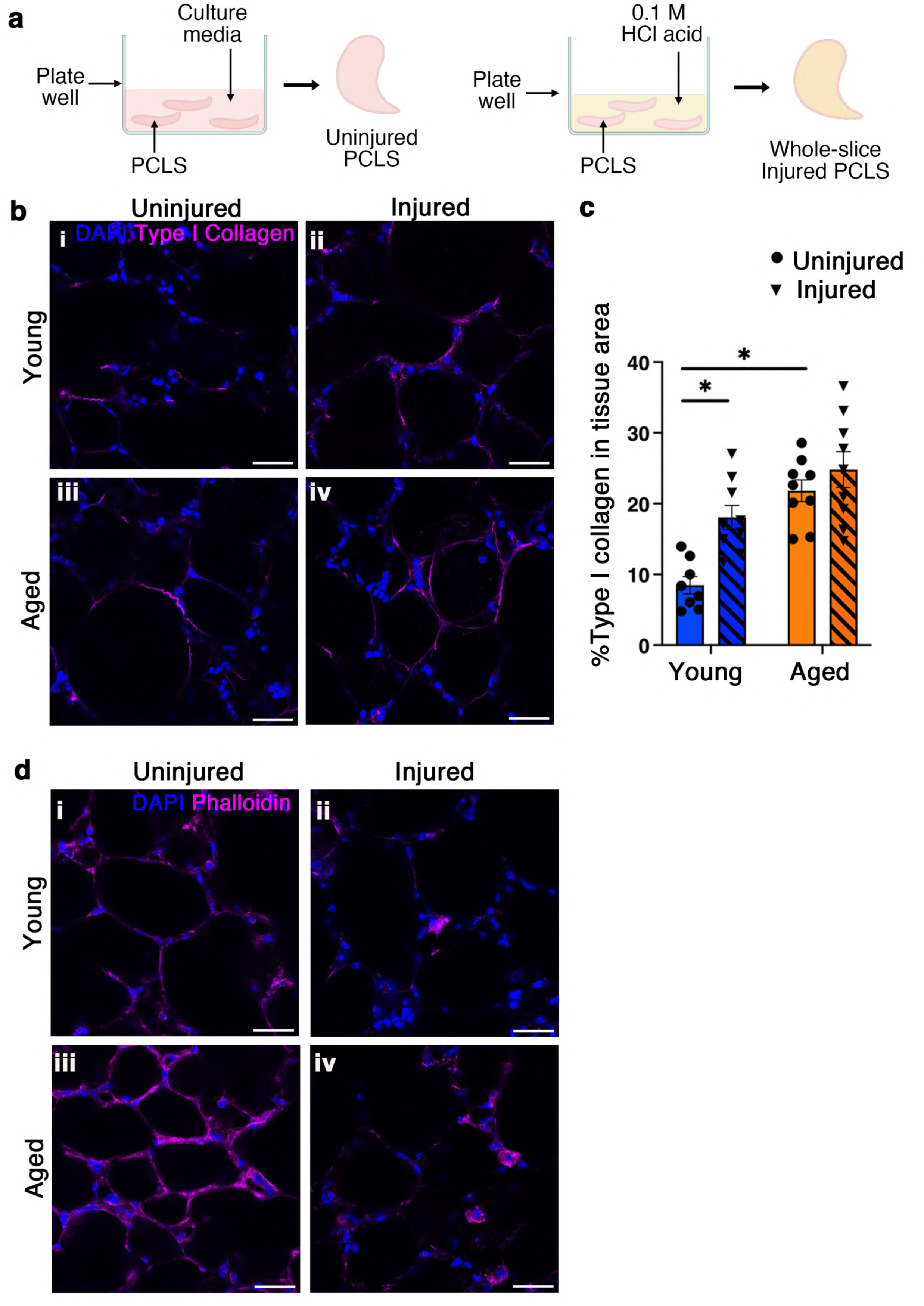
Whole-slice acid-induced injury alters type I collagen deposition in the ECM and the actin cytoskeleton organization in PCLS. (**a**) Whole-slice injury was induced by treating PCLS with 0.1 M HCl across the whole slice for 1 min. (**b, c**) Representative confocal images (**b**) demonstrate the deposition of type I collagen (magenta) in the extracellular matrix (ECM) of PCLS obtained from young and aged mouse lungs, subjected to whole-slice acid injury (“injured”) or no injury (“uninjured”), at 48 hours post-injury. DAPI (blue) stains nuclei. Type I collagen deposition was quantified as the percentage of total tissue area (**c**). (**d**) Representative confocal images show the organization of F-actin filaments (**d**, magenta). DAPI (blue) stains nuclei. Scale bars: 30 μm. Data are presented as mean ± S.E.M. Each dot represents one PCLS. *N* = 3, with 3 PCLS per biological replicate (except for the young uninjured PCLS group). Two-way ANOVA test with Tukey’s multiple comparisons test with *p* < 0.05 as significant.

We first compared the deposition of type I collagen, a key component of the lung interstitial matrix, in the ECM in young and aged PCLS with/without injury (Fig. 2b). Type I collagen content was identified and quantified as percentage total tissue area using a custom automated FIJI macro (Fig. 2c). Young, uninjured PCLS (Fig. 2b i) exhibited a significantly lower percentage area of type I collagen (8.47%) compared to aged, uninjured PCLS (Fig. 2b iii) (21.82%) (*p* = 0.0001). Following acid injury, young PCLS showed a 2.13-fold increase in type I collagen deposition, rising from 8.47% to 18.05% (*p* = 0.0055) (Fig. 2b ii). Aged PCLS showed only a marginal further increase in type I collagen deposition after injury (Fig. 2b iii, iv).

We then investigated whether cytoskeleton organisation was altered in aged and/or injured PCLS by labelling filamentous actin (F-actin), a major component of the actin cytoskeleton. Fig. 2d illustrates F-actin distribution among young and aged PCLS with/without injury. F-actin was visible throughout the cytoplasm in both young and aged uninjured PCLS (Fig. 2d i, iii), and it was more pronounced in aged PCLS (Fig. 2d iii). Upon injury, F-actin organisation was markedly different in both young and aged PCLS (Fig. 2d ii, iv). There was a pronounced loss of cortical actin and instead F-actin was predominantly present in the peri-nuclear region of cells in injured PCLS.

The morphological changes identified in F-actin and collagen I were consistent with known hallmarks of lung ageing and injury, confirming that changes to these read-outs could be detected in physiologically distinct groups of PCLS.

### Comparing early repair responses in young and aged adult PCLS

To determine whether the repair response differed in young and aged lung PCLS, we quantified the percentage of proSP-C^+^ ATII/progenitor cells and Ki67^+^ proliferating cells in uninjured PCLS compared to PCLS subjected to whole-slice injury, obtained from chronologically distinct lungs (Fig. 3a).

**Figure 3.**
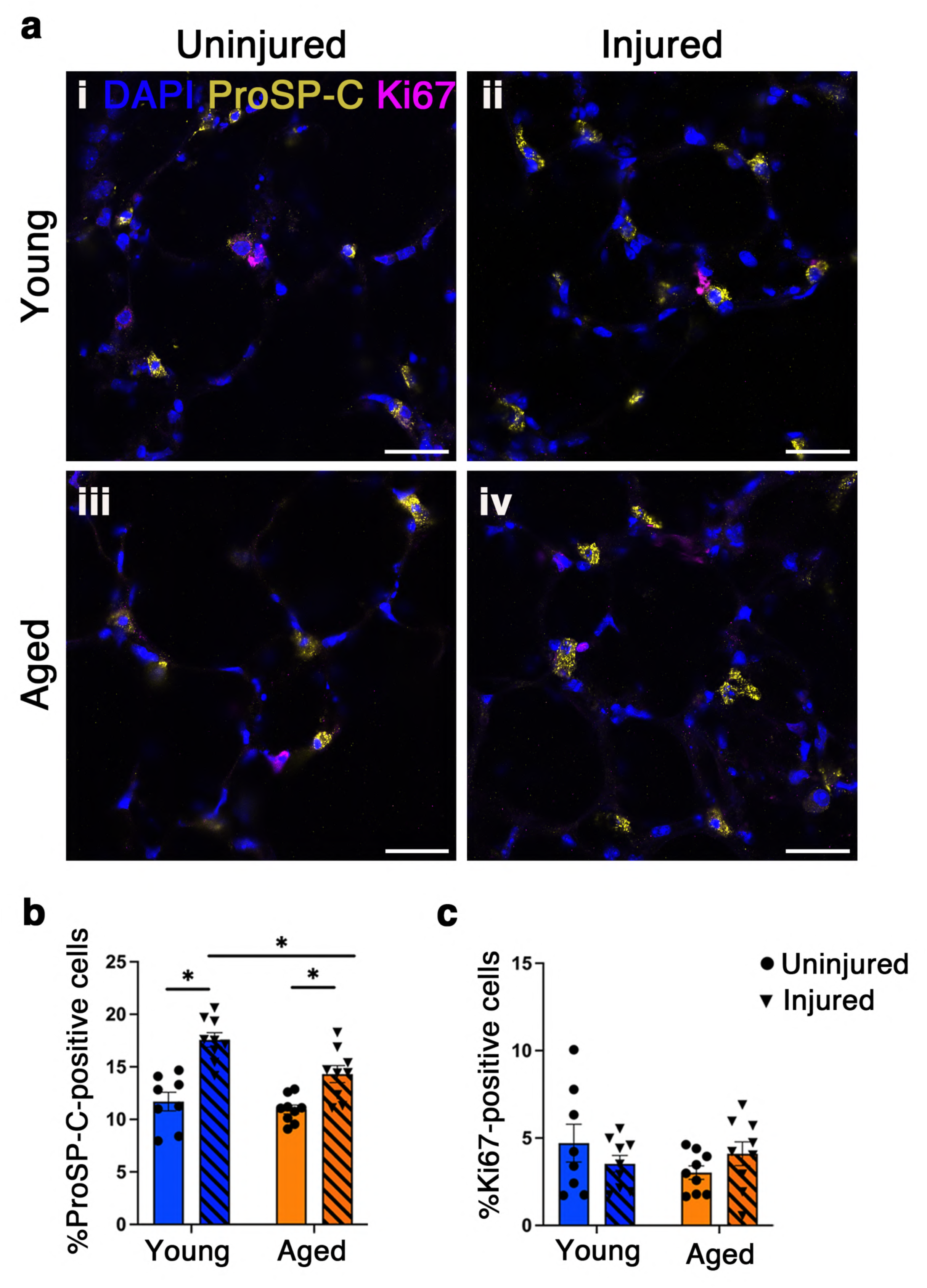
Alveolar progenitor cells increase in PCLS upon whole-slice acid-induced injury. (**a**) Representative confocal images demonstrate the distribution of cells positive for proSP-C (yellow) and Ki67 (magenta) in PCLS 48 hrs after injury. DAPI (blue) stains nuclei. Scale bars: 30 μm. (**b, c**) Percentages of alveolar type II (proSP-C+) cells (**b**) and proliferating (Ki67+) cells (**c**). Data are presented as mean ± S.E.M. Each dot represents one PCLS. *N* = 3, with 3 PCLS per biological replicate (except for the young uninjured PCLS group). Two-way ANOVA test with Tukey’s multiple comparisons test with *p* < 0.05 as significant.

Uninjured PCLS from both young and aged mice exhibited similar levels of proSP-C^+^ cells (11.69% in young vs. 10.93% in aged) (Fig. 3a i, iii, 3b). Following injury, proSP-C^+^ cells increased significantly in both young and aged PCLS (Fig. 3a ii, iv, 3b). The increase was 5.88% in young PCLS (*p* < 0.0001), while aged PCLS showed a smaller rise of 3.38% (*p* = 0.0095) (Fig. 3b). Notably, the percentage of proSP-C^+^ cells was significantly higher after injury in young PCLS than in aged PCLS (*p* = 0.0134).

The percentage of proliferating (Ki67^+^) cells was similar in young and aged PCLS, with no notable changes observed after injury (Fig. 3c).

### Treatment with EVs derived from young, uninjured PCLS (YU EVs) decreases apoptotic cells in both young and aged AIR-PCLS

Having established that tangible and quantifiable changes could be detected in aged and/or injured lungs, we sought to determine whether EVs obtained from young or aged PCLS exerted functional effects on spatially injured PCLS obtained from either young or aged mice. To do this, we utilised the acid injury and repair (AIR model) that we previously established^27^. In this model, spatially restricted acid-induced injury of PCLS leads to quantifiable pro-repair responses (Fig. 4a). This heterogeneous pattern of injured tissue adjacent to an uninjured area created in the AIR model (Fig. 4b-e) is highly representative of diseased lungs *in vivo*.

**Figure 4.**
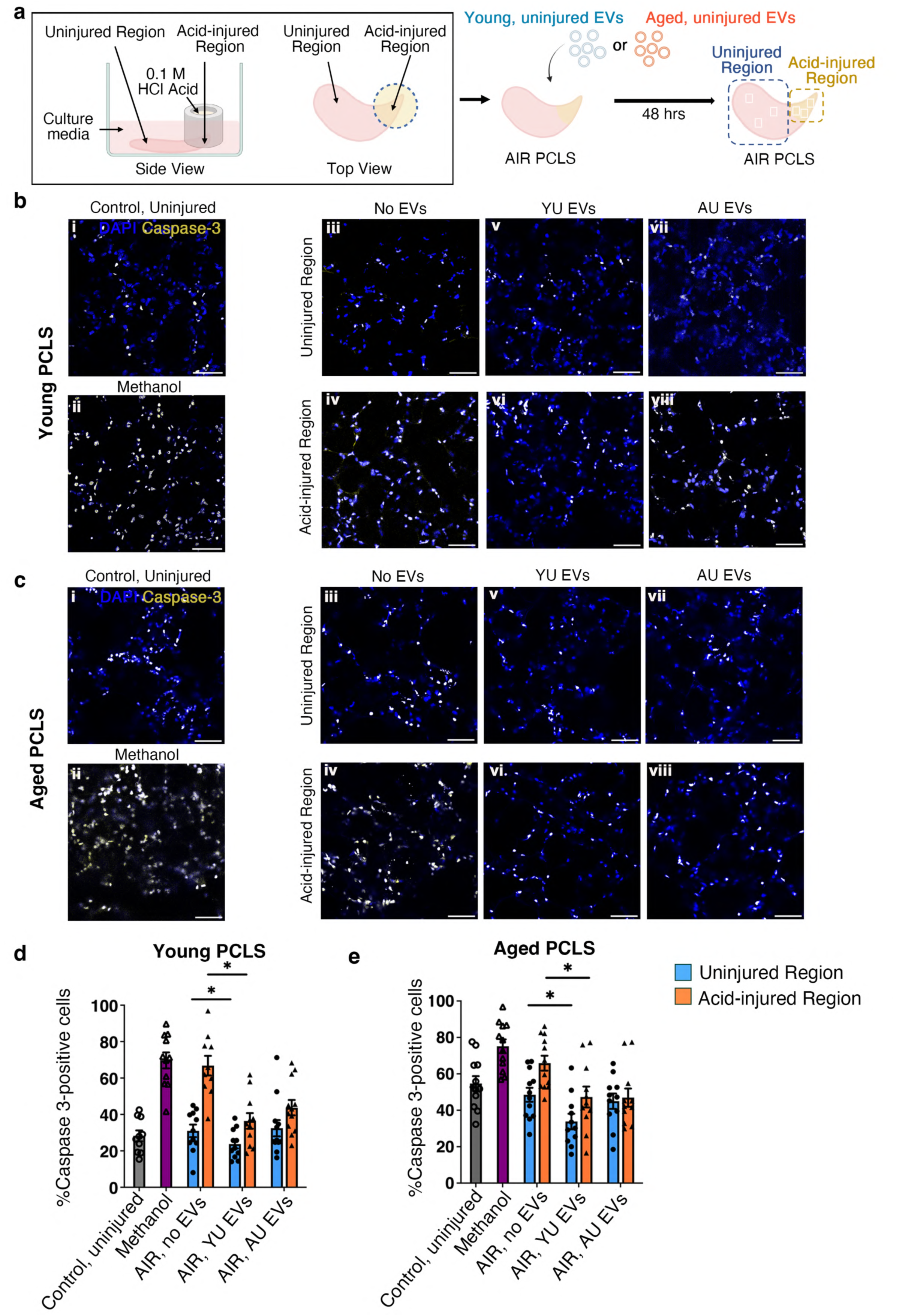
EVs isolated from young, uninjured PCLS decreases apoptosis in young and aged AIR PCLS. (**a**) The acid injury repair (AIR) model was established by injuring a spatially restricted region of PCLS with 0.1 M HCl for 1 min, creating an acid-injured region adjacent to an uninjured region. AIR PCLS were treated for YU EVs or AU EVs for 48 hrs, then stained and imaged. For each AIR PCLS, 3 separate fields of view were imaged in the uninjured region and 3 in the acid-injured region. (**b-c**) Representative confocal images show cleaved caspase 3^+^ cells (yellow) in injured and uninjured regions of young (**b**) or aged (**c**) AIR PCLS treated with no EVs, YU EVs or AU EVs. Uninjured PCLS or PCLS treated with methanol served as controls. DAPI (blue) stains nuclei. Scale bars: 50 µm. (**d-e**) Percentage of caspase 3^+^ cells in young (**d**) or aged (**e**) PCLS. Data are presented as mean ± S.E.M. Each dot represents one PCLS. *N* = 4, with 3 PCLS per biological replicate, except in some groups where two PCLS were used in one or two biological replicates. Two-way ANOVA test with Tukey’s multiple comparisons test with *p* < 0.05 as significant.

First, the baseline percentage of caspase 3^+^ cells were measured and quantified in young PCLS. Young control, uninjured PCLS contained an average of 28.19% caspase 3^+^ cells (Fig. 4b i, d), while methanol-treated PCLS (a positive control for dead cells) showed a significantly higher percentage of caspase 3^+^ cells, as expected (69.70%, Fig. 4b ii, d, *p* < 0.0001).

In young AIR-PCLS without any EV treatment the acid-injured region exhibited a markedly increased percentage of caspase 3^+^ cells (66.82%, Fig. 4b iv, d) compared to the uninjured region (30.92%, Fig. 4b iii, d, *p* < 0.0001). Treatment with YU EVs significantly reduced caspase 3^+^ cells in both the acid-injured (by 30.39%, Fig. 4b vi vs. iv) and uninjured (by 7.26%, Fig. 4b v vs. iii, d) regions (*p* = 0.0022 vs no EVs). Treatment with AU EVs did not alter the percentage of caspase 3^+^ cells in the uninjured region (Fig. 4b vii vs. iii) and although there was some reduction in the acid-injured region (Fig. 4b viii vs. iv), this was not statistically significant (Fig. 4d), highlighting a marked difference in the effects of YU and AU EV treatments.

In aged PCLS (Fig. 4c, e), control, uninjured PCLS exhibited a baseline level of 54.71% caspase 3^+^ (Fig. 4c i), significantly higher than that of young PCLS (*p* < 0.0001 vs. Fig. 4b i), but still lower than methanol-treated PCLS (75.14%, Fig. 4c ii, *p* = 0.0376 vs. Fig. 4c i). In aged AIR PCLS, the acid-injured region contained a higher percentage of caspase 3^+^ cells (65.87%, Fig. 4c iv) compared to the uninjured region (48.53%, Fig. 4c iii), though this difference was smaller than seen in young AIR-PCLS and was not statistically significant.

Treatment of aged PCLS with YU EVs significantly reduced caspase 3^+^ cells in both the acid-injured (by 18.57%, Fig. 4c vi vs. iv, *p* = 0.0105 vs no EVs) and uninjured (by 14.70%, Fig. 4c v vs. iii, *p* = 0.0105 vs. no EVs) regions of AIR-PCLS, mirroring their effect on young AIR-PCLS (Fig. 4b v, vi). AU EVs led to a small reduction in caspase 3^+^ cells in both the injured (Fig. 4c viii vs. iv) and uninjured (Fig. 4c vii vs. iii) regions, but this reduction was not statistically significant (Fig. 4e, *p* = 0.2596).

### Treatment with EVs derived from young, uninjured PCLS (YU EVs) increases alveolar progenitor cells in aged AIR-PCLS

We then assessed whether PCLS-derived EVs could enhance the early repair response after injury by treating AIR-PCLS with either YU or AU EVs and quantifying the number of ATII cells (proSP-C^+^) and proliferating cells (Ki67^+^) 48 hrs after treatment. Previously, we showed that acid injury results in an increased percentage of alveolar progenitor cells (proSP-C^+^) within the injured region of AIR-PCLS^27,28^.

Consistent with previous findings^27^, injury significantly increased the percentage of proSP-C^+^ cells in young AIR-PCLS (Fig. 5a i-iii, c). Young control, uninjured PCLS (Fig. 5a i) contained 7.52% proSP-C^+^ cells. After injury, the percentage of proSP-C^+^ cells rose significantly in the acid-injured region of AIR PCLS (17.44%, Fig. 5a iii) compared to the uninjured region (8.78%, Fig. 5a ii, *p* < 0.0001). However, treatment with either YU or AU EVs did not result in any further increase in proSP-C^+^ cells in the acid-injured region of young AIR-PCLS (Fig. 5 v vs. iii, vi vs. iii, c).

**Figure 5.**
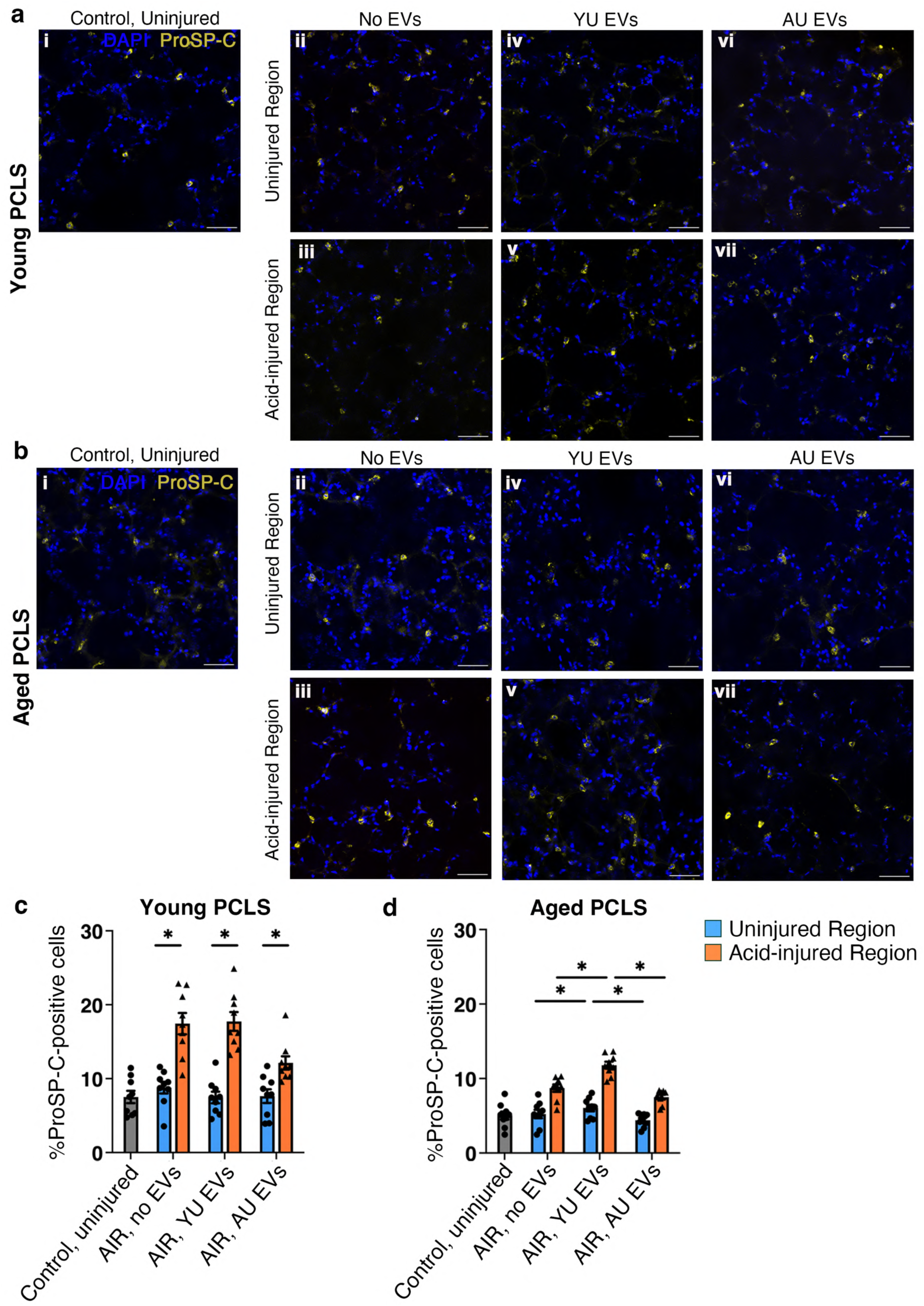
EVs isolated from young, uninjured PCLS increases the number of proSP-C^+^ cells in aged AIR PCLS. (**a-b**) Representative confocal images demonstrate the distribution of proSP-C^+^ cells (yellow; alveolar type II/progenitor cells) in injured and uninjured regions of young (**a**) and aged (**b**) AIR PCLS treated with no EVs, YU EVs or AU EVs for 48 hrs. DAPI (blue) stains nuclei. Uninjured PCLS served as a control. DAPI (blue) stains nuclei. Scale bars: 50 µm. (**c-d**) Percentages of proSP-C^+^ cells in young (**c**) or aged (**d**) PCLS. Data are presented as mean ± S.E.M. Each dot represents one PCLS. *N* = 3, with 3 PCLS per biological replicate. Two-way ANOVA test with Tukey’s multiple comparisons test with *p* < 0.05 as significant.

Aged PCLS exhibited a lower level of proSP-C^+^ cells across all groups (Fig. 5b, d) compared to young PCLS, reflecting the reduced repair capacity of aged lungs. Aged control, uninjured PCLS contained 5.03% proSP-C^+^ cells (Fig. 5b i), significantly lower than that in young control PCLS (*p* = 0.0315 vs. Fig. 5a i). In response to injury, aged AIR-PCLS showed a significant increase in proSP-C^+^ cells in the acid-injured region of AIR-PCLS (8.72%, Fig. 5b iii) compared to the uninjured region (5.23%, Fig. 5b ii, *p* < 0.0001). However, this increase (3.49%) was smaller than that observed in young AIR-PCLS (8.66%).

Notably, treatment with YU EVs further increased the number of proSP-C^+^ cells in aged AIR-PCLS, reaching 11.78% in the acid-injured region (Fig. 5b v) and 6.05% in the uninjured region (Fig. 5b iv). These levels were significantly higher than those in AIR-PCLS that received no EVs (Fig. 5b iii, ii, d, *p* = 0.0041) or treated with AU EVs (Fig. 5b vii, vi, d, *p* < 0.0001). In contrast, AU EVs did not further increase the percentage of proSP-C^+^ cells in either region of aged AIR-PCLS (Fig. 5b vii vs. iii, vi vs. ii, d).

### Treatment with EVs derived from aged, uninjured PCLS (AU EVs) alters cell proliferation in young AIR-PCLS

We then investigated whether YU or AU EVs can alter cell proliferation, which was previously shown to increase in the AIR-PCLS, following spatial acid injury^27^.

In young AIR-PCLS, the acid-injured region did not show a significantly altered percentage of proliferating cells (Ki67^+^) compared to control, uninjured PCLS (2.86% vs 3.21%) (Fig. 6a iii vs. i, c). Treatment with YU EVs did not alter the number of Ki67^+^ cells (Fig. 6a iv, v, c).

**Figure 6.**
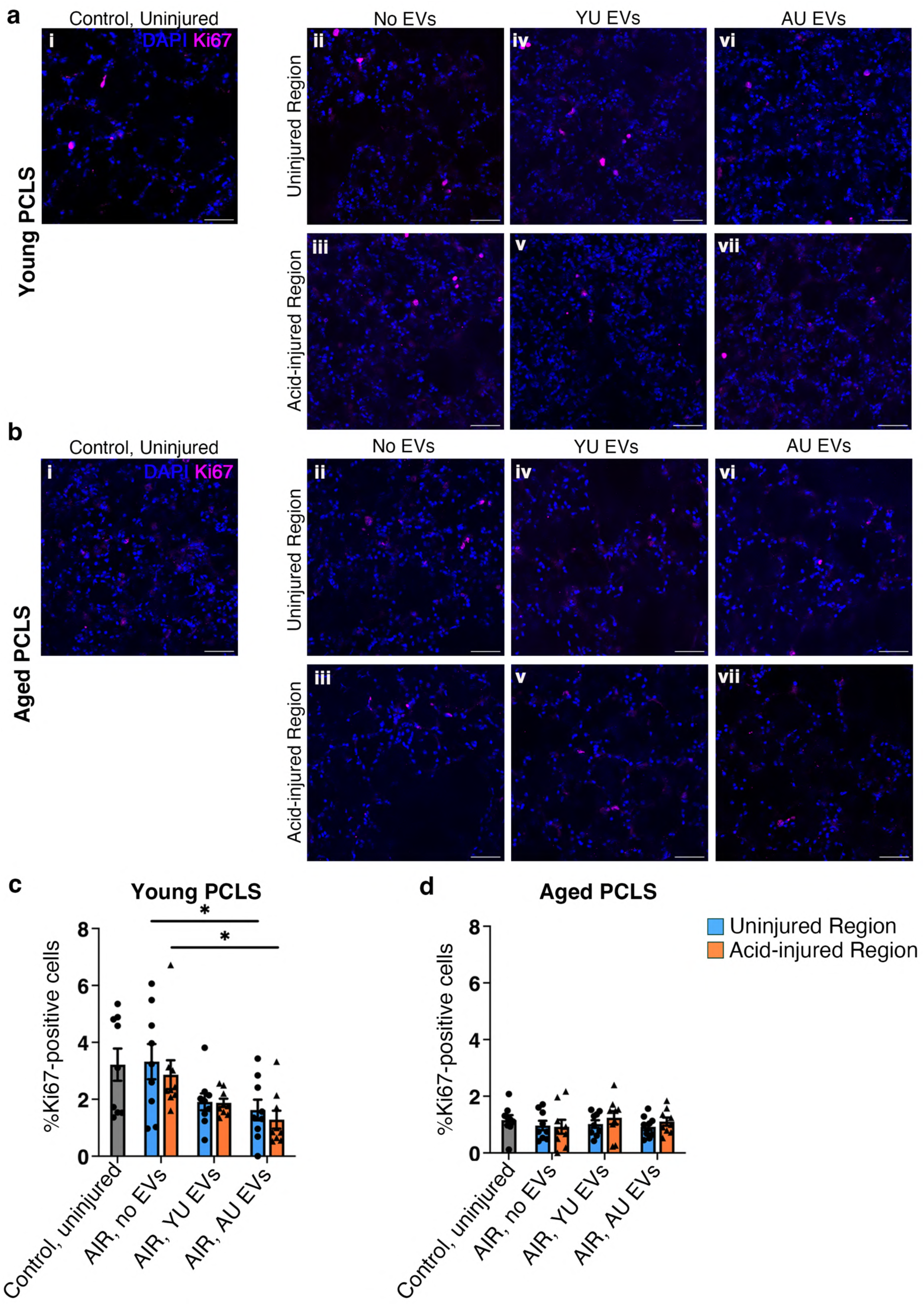
EVs isolated from aged, uninjured PCLS decreases cell proliferation in young AIR PCLS. **(a-b)** Representative confocal images demonstrate the distribution of Ki67^+^ cells (magenta; proliferating cells) in injured and uninjured regions of young (**a**) and aged (**b**) AIR PCLS treated with no EVs, YU EVs or AU EVs for 48 hrs. DAPI (blue) stains nuclei. Uninjured PCLS served as a control. DAPI (blue) stains nuclei. Scale bars: 50 µm. (**c-d**) Percentages of Ki67^+^ cells in young (**c**) or aged (**d**) PCLS. Data are presented as mean ± S.E.M. Each dot represents one PCLS. *N* = 3, with 3 PCLS per biological replicate. Two-way ANOVA test with Tukey’s multiple comparisons test with *p* < 0.05 as significant.

However, interestingly, treatment with AU EVs significantly reduced Ki67^+^ cells in both uninjured (1.62%, Fig. 6a vi, c) and acid-injured (1.29%, Fig. 6a vii, c) regions of the AIR-PCLS (*p* = 0.0064).

Compared to young PCLS, all groups of aged PCLS had a lower percentage of Ki67^+^ cells (Fig. 6b, d). For example, aged control, uninjured PCLS contained 1.16% Ki67^+^ cells (Fig 6b i), significantly lower than that in young control, uninjured PCLS (*p* < 0.0008 vs. Fig. 6a i).

Neither injury nor EV treatment altered the percentage of Ki67^+^ cells in aged PCLS.

### Type I collagen deposition in the ECM increases with ageing and upon injury in AIR-PCLS and is modified by YU EVs

To examine whether PCLS-derived EVs could modify the ECM in an injured tissue environment, we first quantified type I collagen in both the injured and uninjured regions of young and aged AIR-PCLS after treatment with either YU or AU EVs for 48 hrs.

Following injury, type I collagen deposition increased significantly in both young and aged PCLS (Fig. 7). Young control, uninjured PCLS contained a percentage area covered by type I collagen of 9.25% (Fig. 7a i, c). After injury, type I collagen accumulation markedly increased in the acid-injured region (35.83%, Fig. 7a iii) compared to the uninjured region in young AIR-PCLS (12.71%, Fig. 7a ii, c, *p* < 0.0001). As anticipated, aged PCLS exhibited higher baseline levels of type I collagen deposition compared to young PCLS. In aged control, uninjured PCLS, collagen covered 23.37% of the tissue area, significantly higher than in young PCLS (Fig. 7b i, d vs. Fig. 7a i, c, *p* = 0.0008). Similarly, acid injury led to a significant increase in type I collagen deposition in the acid-injured region of aged AIR-PCLS (36.82%, Fig. 7b iii) compared to the uninjured region (19.82%, Fig. 7b ii, d, *p* < 0.0001).

**Figure 7.**
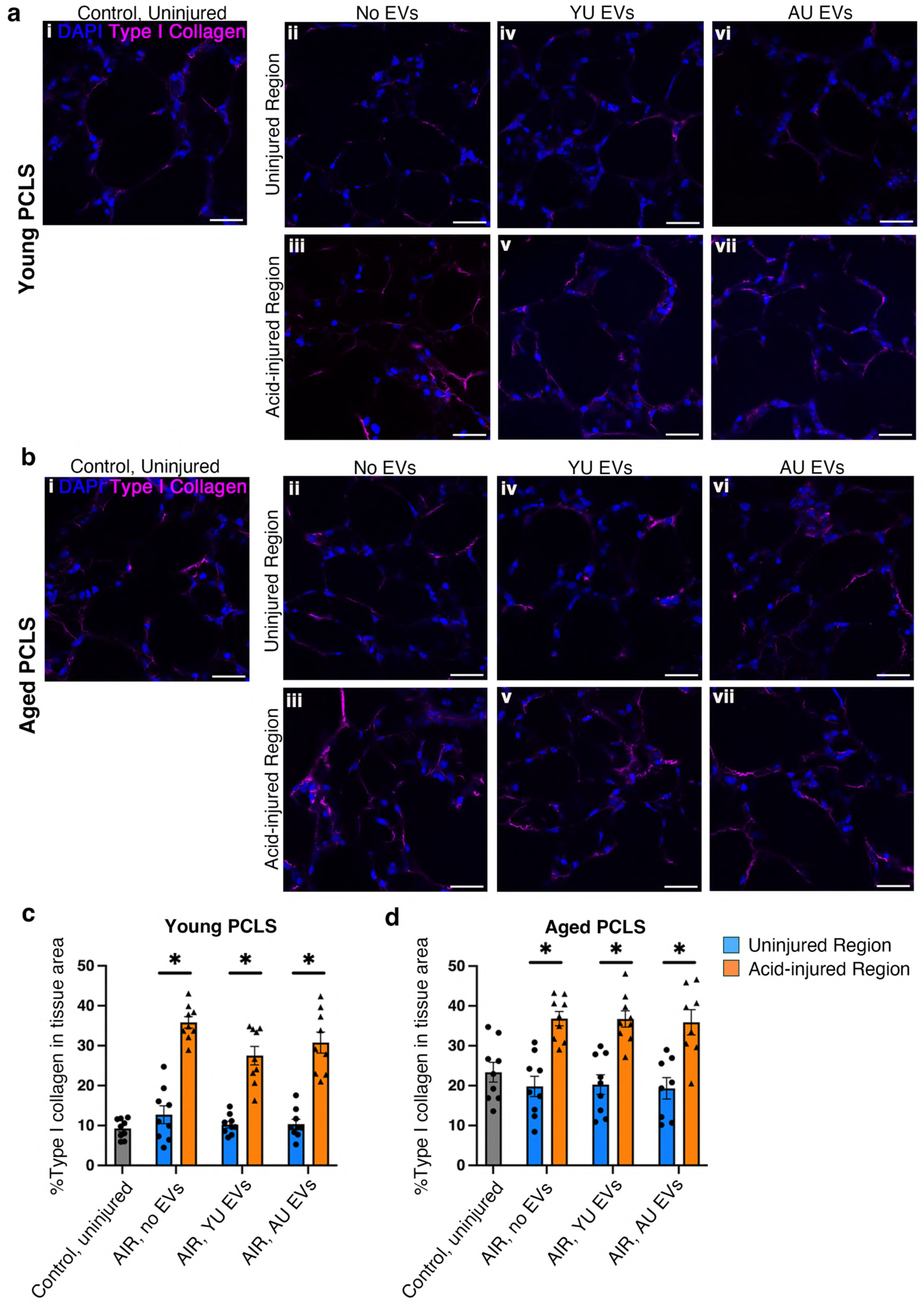
Type I collagen deposition in the ECM increases with ageing and upon injury, and EVs isolated from young, uninjured PCLS shows a trend to reducing post-injury collagen deposition in young PCLS. (**a-b**) Representative confocal images demonstrate the deposition of type I collagen (magenta) in the ECM in in injured and uninjured regions of young (**a**) and aged (**b**) AIR PCLS treated with no EVs, YU EVs or AU EVs for 48 hrs. DAPI (blue) stains nuclei. Uninjured PCLS served as a control. DAPI (blue) stains nuclei. Scale bars: 50 µm. (**c-d**) Type I collagen deposition in the ECM quantified as a percentage of tissue area in young (**c**) or aged (**d**) PCLS. Data are presented as mean ± S.E.M. Each dot represents one PCLS. *N* = 3 biological replicates, with 2-3 PCLS per biological replicaten. Two-way ANOVA test with Tukey’s multiple comparisons test with *p* < 0.05 as significant.

Additionally, groups of young or aged AIR-PCLS were treated with either YU or AU EVs for 48 hrs before quantifying type I collagen. In young AIR-PCLS treated with YU EVs, collagen accumulation in the injured region was 27.51% (Fig. 7a v), which was 8.32% lower than in AIR-PCLS with no EV treatment (Fig. 7a iii, *p* = 0.0680). Collagen deposition in the injured region of young AIR-PCLS treated with AU EVs was only 3.45% lower (Fig. 7a vii vs. iii, *p* = 0.4412). However, these differences were not statistically significant.

In aged AIR-PCLS, there was no effect on type I collagen following treatment with either YU or AU EVs (Fig. 7b, d, *p* > 0.9999).

### PCLS secrete EVs with altered miRNA contents with ageing and upon injury

Functional experiments showed differing biological responses of PCLS to EVs isolated from aged or young lungs. EVs contain a variety of bioactive molecules, existing research has identified microRNA (miRNA) and protein content of EVs as important mediators of their function^29^. Here we focused on their miRNA content. RNA was isolated from the following four groups of EVs: (1) young, uninjured PCLS (“YU EVs”), (2) young, injured PCLS (“YI EVs”), (3) aged, uninjured PCLS (“AU EVs”), and (4) aged, injured PCLS (“AI EVs”). Small RNA sequencing was performed and miRNAs that were differentially expressed among the EV groups were identified. For each miRNA that reached the significance threshold (adjusted p < 0.05 and |log_2_(fold change)| ≥ 1), further analysis was conducted. First, experimentally validated target genes were retrieved from existing databases (Supplementary Data 1, 2). Then, PANTHER pathway enrichment analysis was conducted to identify overrepresented signalling pathways among the target genes (Supplementary 3, 4), followed by gene network analysis to reveal interactions across enriched pathways.

We first compared AU EVs to YU EVs. The expression level of miR-203-3p, was down-regulated by approximately 15.56-fold (log_2_[fold change] ≈ −3.96, adjusted *p* = 0.004) in AU EVs (Fig. 8a). MiRNA target analysis identified 120 genes that were previously experimentally validated targets of miR-203-3p (Supplementary Data 1, 2). Three signalling pathways were significantly overrepresented in the target genes of miR-203-3p: Ras pathway (cell survival and proliferation, and cytoskeletal reorganization), CCKR signalling map (cell survival, proliferation, migration and adhesion) and Integrin signalling pathway (actin reorganization), involving 10 of its target genes in total (Fig. 8b). Network analysis (Fig. 8c) revealed extensive connections across the pathways through shared genes, especially between Ras and CCKR pathways.

**Figure 8.**
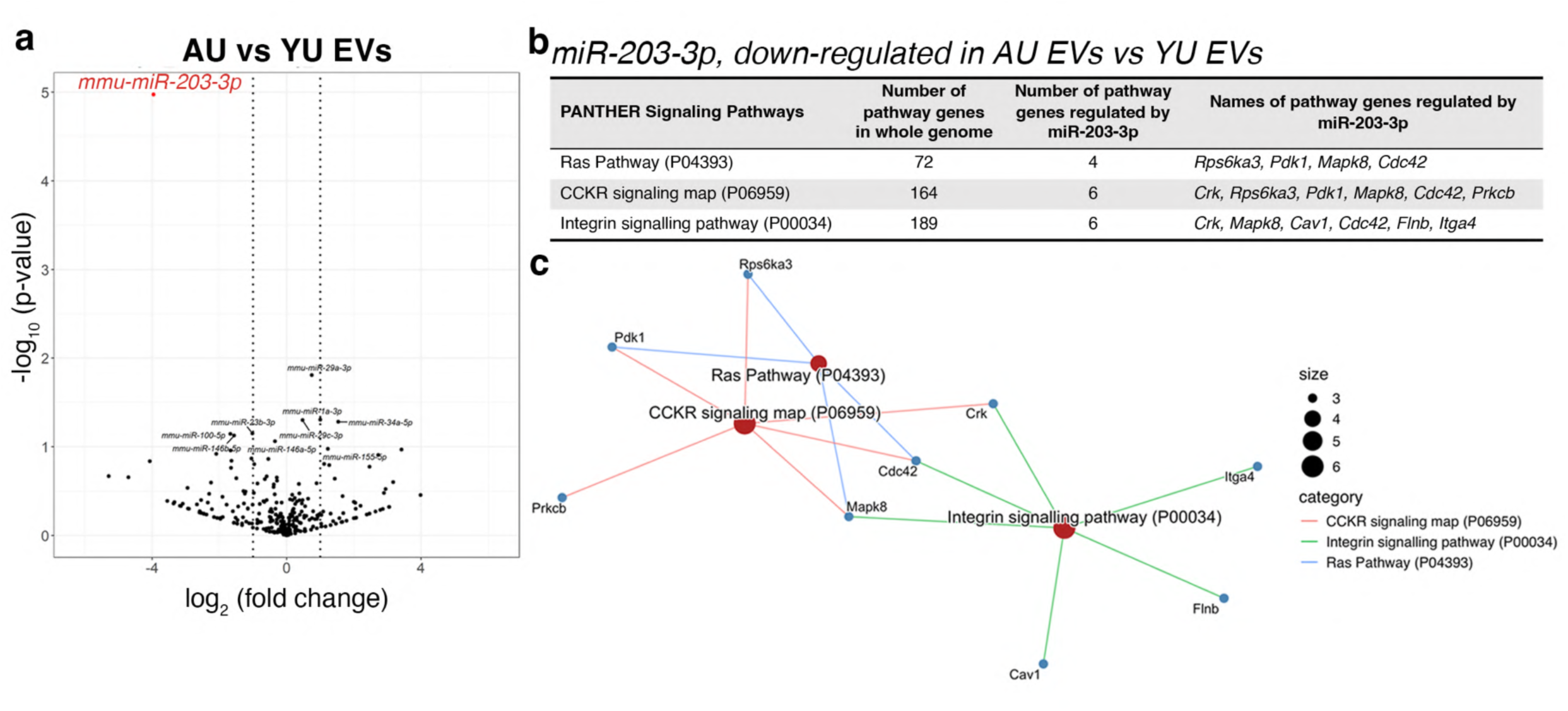
MiRNA miR-203-3p is down-regulated in EVs isolated from aged, uninjured PCLS compared to EVs isolated from young, uninjured PCLS. (**a**) Volcano plot of differentially expressed micro RNAs (miRNAs) in AU EVs in relative to YU EVs. Each dot represents one miRNA. Significantly differentially expressed miRNA (fold change > 2, adjusted *p*-value < 0.05) is highlighted in red. *N* = 3. (**b**) All PANTHER signalling pathways overrepresented (FDR < 0.05 from Fisher’s exact test) in target genes of miR-203-3p, and miR-203-3p target genes associated with each overrepresented pathway. (**c**) Network analysis of genes associated with all overrepresented signalling pathways in target genes of miR-203-3p.

Comparison of YI EVs to YU EVs revealed that miR-150-5p was down-regulated by 2.08-fold (log_2_[fold change] ≈ −1.06, adjusted *p* = 0.048) in YI EVs (Fig. 9a). miRNA target analysis identified 327 experimentally validated target genes of miR-150-5p (Supplementary Data 1, 2). Nine signalling pathways were overrepresented, involving 33 miR-150-5p target genes (Fig. 9b, Supplementary Data 3, 4). Of those, 3 pathways are important for cell survival and proliferation (EGF receptor, PDGF, and CCKR signalling pathways). Notably, pathways associated with tissue repair, angiogenesis and/or lung development are overrepresented in the target genes of miR-150-5p: EGF receptor signalling pathway (lung development and repair), PDGF signalling pathway (lung development, angiogenesis), Hedgehog signalling pathway (lung development and repair), angiogenesis and FGF signalling pathway (angiogenesis, lung development and repair). Among the overrepresented pathways, the Hedgehog signalling pathway had the highest percentage of genes regulated by miR-150-5p (20.00%). The percentage was approximately double that of the second and third highest pathways, insulin/IGF pathway-protein kinase B signalling cascade (10.53%) and interleukin signalling pathway genes (9.57%). Network analysis (Fig. 9c) showed considerable gene interconnections across multiple pathways, namely angiogenesis, EGF receptor, FGF, PDGF, CCKR and interleukin signalling pathways, indicating strong crosstalk between these pathways. In contrast, Hedgehog signalling pathway displayed limited connections with the other pathways.

**Figure 9.**
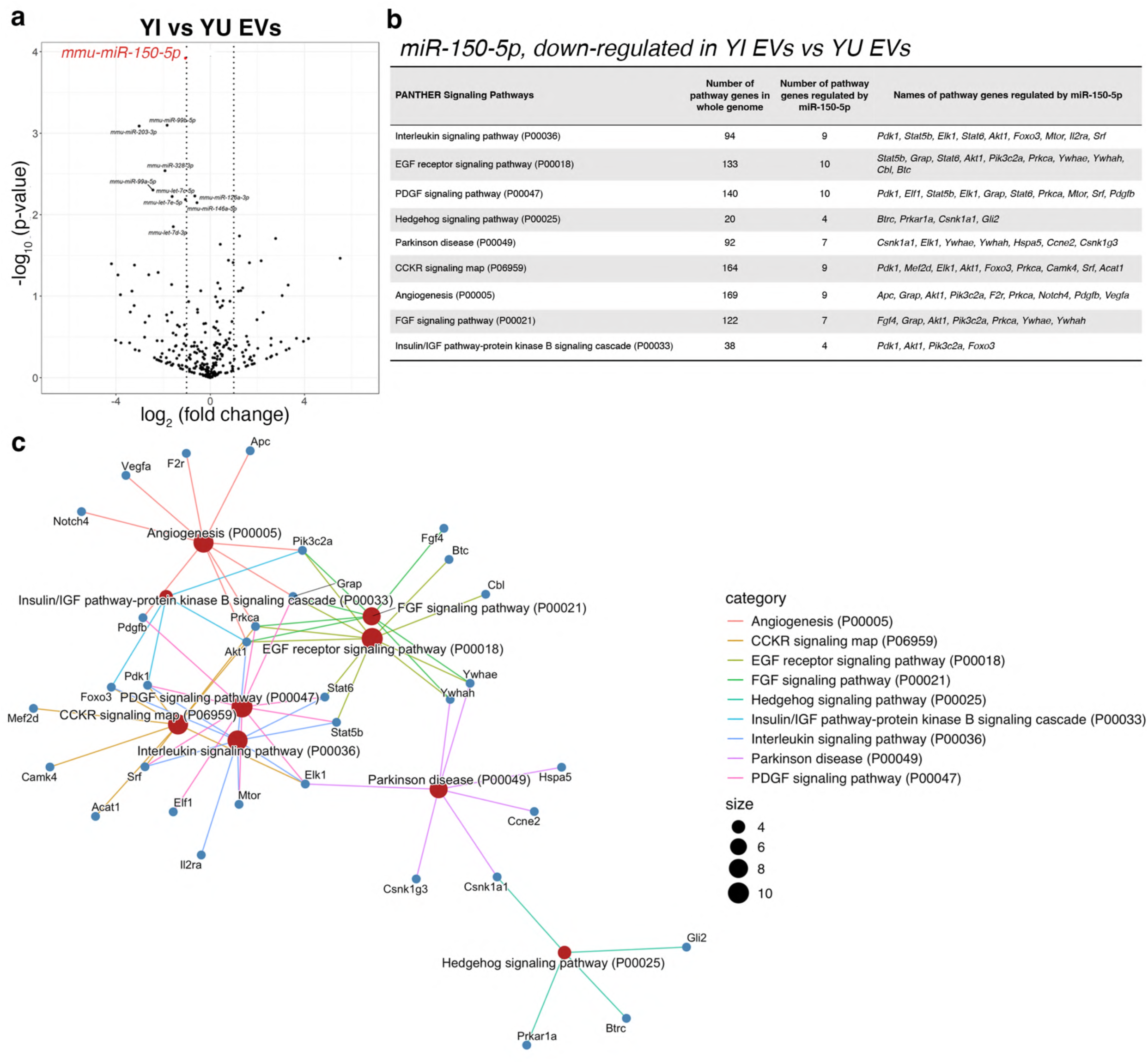
MiRNA miR-150-5p is down-regulated in EVs isolated from young, injured PCLS compared to EVs isolated from young, uninjured PCLS. (**a**) Volcano plot of differentially expressed miRNAs in YI EVs in relative to YU EVs. Each dot represents one miRNA. Significantly differentially expressed miRNA (fold change > 2, adjusted *p*-value < 0.05) is highlighted in red. *N* = 3. (**b**) All PANTHER signalling pathways overrepresented (FDR < 0.05 from Fisher’s exact test) in target genes of miR-150-5p, and miR-150-5p target genes associated with each overrepresented pathway. (**c**) Network analysis of genes associated with all overrepresented signalling pathways in target genes of miR-150-5p.

Upon injury, aged PCLS secreted EVs with significantly reduced levels of 3 miRNAs: let-7e-5p (down-regulated by 2.93-fold, log2[fold change] ≈ −1.55, adjusted *p* = 0.010), miR-328-3p (down-regulated by 5.72-fold, log2[fold change] ≈ −2.52, adjusted *p* = 0.010) and miR-151-3p (down-regulated by 15.00-fold, log2[fold change] ≈ −3.91, adjusted *p* = 0.010) (Fig. 10a).

**Figure 10.**
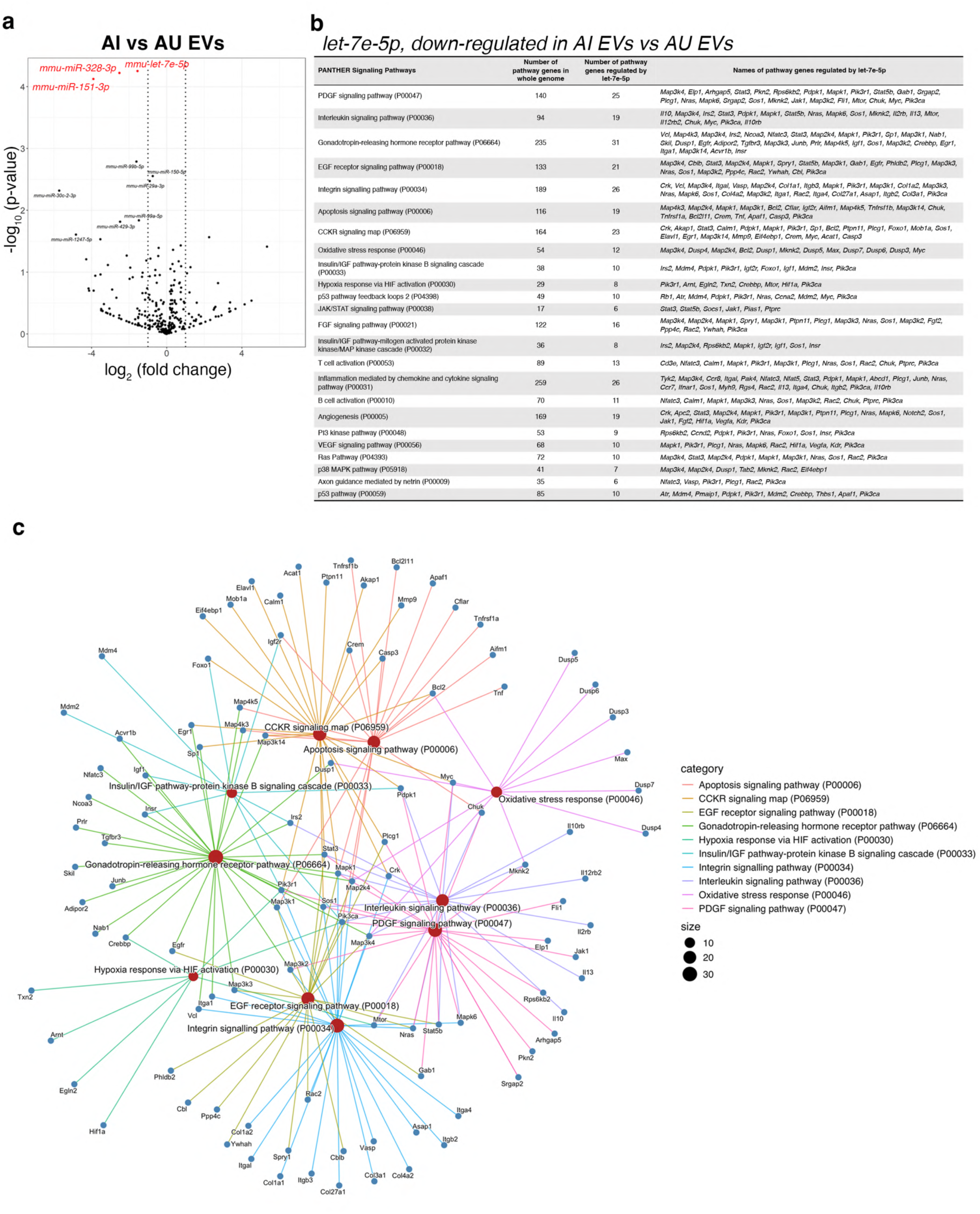
MiRNAs let-7e-5p, miR-328-3p and miR-151-3p are down-regulated in EVs isolated from aged, injured PCLS compared to EVs isolated from aged, uninjured PCLS. (**a**) Volcano plot of differentially expressed miRNAs between AI EVs in relative to AU PCLS. Each dot represents one miRNA. Significantly differentially expressed miRNAs (fold change > 2, adjusted *p*-value < 0.05) are highlighted in red. *N* = 3. (**b**) All PANTHER signalling pathways overrepresented (FDR < 0.05 from Fisher’s exact test) in target genes of let-7e-5p and let-7e-5p target genes associated with each overrepresented pathway. **(c)** Network analysis of genes associated with top 10 overrepresented PANTHER signalling pathways in target genes of let-7e-5p.

MiRNAs miR-328-3p and miR-151-3p had 247 and 92 experimentally validated target genes, respectively (Supplementary Data 1, 2); however, no signalling pathways were significantly represented among their target genes. In contrast, let-7e-5p had 1028 validated target genes among which 24 signalling pathways were overrepresented, involving 130 of let-7e-5p targets (Fig. 10b, Supplementary Data 1-4). These enriched pathways included those associated with cell survival and proliferation, including CCKR, PDGF and EGF receptor pathways, and those critical for tissue repair and lung development, namely angiogenesis, PDGF, EGF receptor, FGF and VEGF signalling pathways. Additionally, several pathways related cellular response to stress and immune responses were also overrepresented in let-7e-5p targets, such as oxidative stress response, p53/MAPK and apoptosis pathways.

Network analysis of the top 10 overrepresented pathways (Fig. 10c) highlighted substantial gene interactions, suggesting pathway crosstalk via genes regulated by let-7e-5p.

### Treatment with EVs isolated from young, injured lung tissue inhibit significant increase in type I collagen post-injury

Our previous functional experiments did not include EVs derived from young, injured lung tissue. Having identified that MiR-150-5p was differentially expressed between YU EVs and YI EVs, we treated young AIR-PCLS with YI EVs and then compared their effects to those of YU EVs in two aspects: their ability to alter the number of alveolar progenitor cells, an indication of a repair response following injury, and their capacity to attenuate post-injury type I collagen accumulation in the ECM (Fig. 11a).

**Figure 11.**
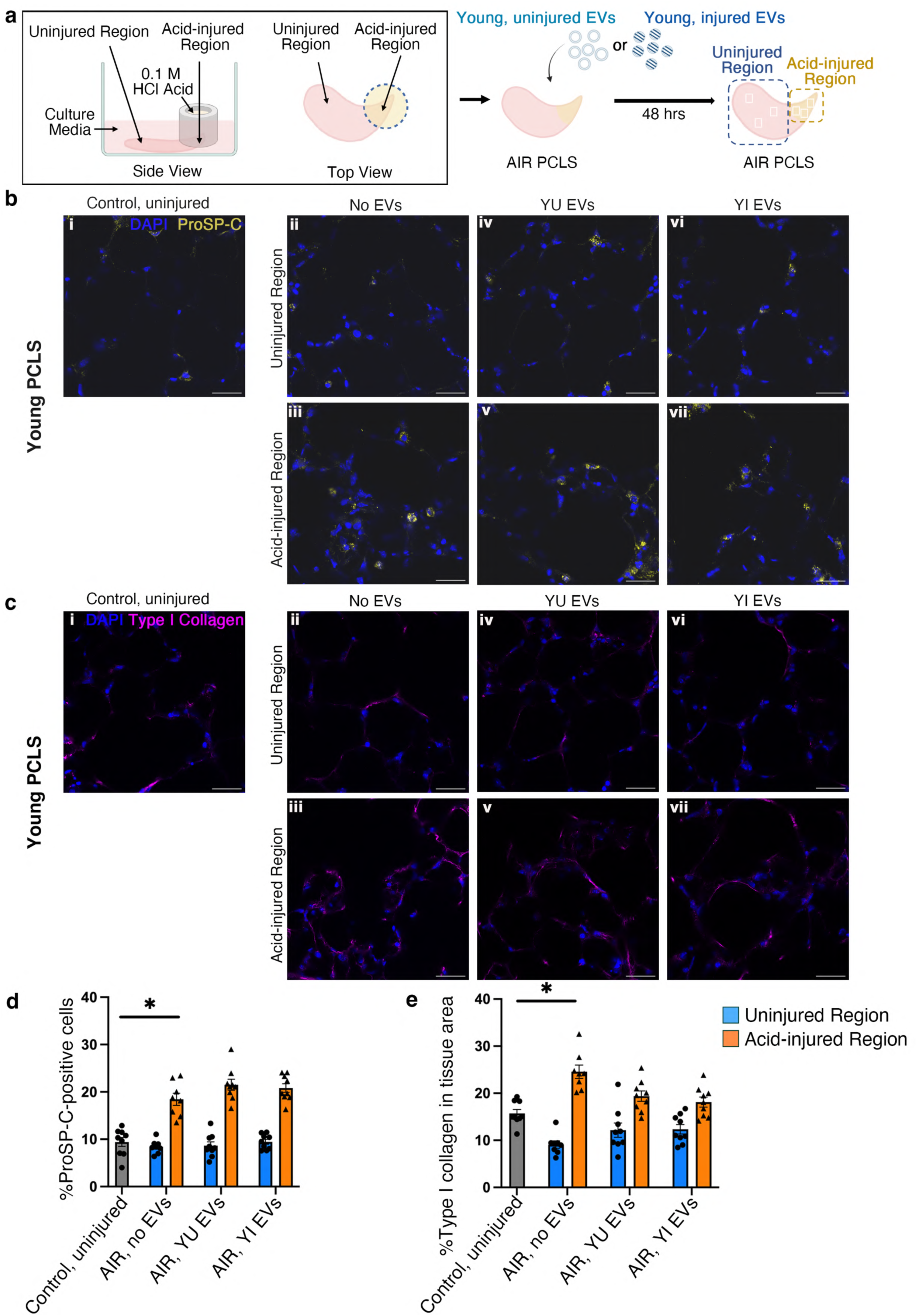
EVs isolated from young, injured PCLS shows a trend to reducing post-injury collagen deposition in young PCLS. (**a-b**) Representative confocal images demonstrate proSP-C^+^ cells (**a**, yellow) and the deposition of type I collagen (**b**, magenta) in the ECM in injured and uninjured regions of young AIR PCLS that received no EVs, YU EVs or YI EVs. Uninjured PCLS served as a control. DAPI (blue) stains nuclei. Scale bars: 30 µm. (**c, d**) Percentage of proSP-C^+^ cells (c) and type I collagen deposition quantified as a percentage of tissue area (**d**) in young PCLS. Data are presented as mean ± S.E.M. Each dot represents one PCLS. *N* = 3, with 3 PCLS per biological replicate (except for AIR PCLS treated with no EVs, 2 PCLS in 1 of the biological replicates). Two-way ANOVA test with Tukey’s multiple comparisons test with *p* < 0.05 as significant.

As expected, after injury, proSP-C^+^ cells were significantly increased in the acid-injured region (18.45%, Fig. 11b iii) of AIR-PCLS compared to the uninjured region (8.40%, Fig. 11b ii, *p* < 0.0001). Treatment with either YU EVs or with YI EVs marginally increased the percentage of proSP-C^+^ cells in the acid injured region of AIR-PCLS by approximately 2%, YU EVs (21.52% proSP-C^+^, Fig. 11b v) or YI EVs (20.82% proSP-C^+^, Fig. 11b vii), but this difference was not statistically significant (Fig. 11d).

Finally, we evaluated whether YI EVs could alter type I collagen deposition in the ECM (Fig. 11c, e). In agreement with previous data (Fig. 7), acid injury increased collagen accumulation in the acid-injured region of AIR-PCLS by 8.86%, compared to control, uninjured PCLS (from 15.71% to 24.57%, Fig. 11c i, iii, 11e, *p* = 0.0255).

Treatment with EVs led to an inhibition of any significant increase in collagen deposition in the acid injured regions. In the YU EV-treated group, there was only a 3.66% increase in collagen area in response to injury compared to control, uninjured PCLS (from 15.71% to 19.37%, Fig. 11c i,v, 11e, *p* = 0.1100). In the YI EV-treated group, this increase was only 2.41% (from 15.71% to 18.12%, Fig. 11c i,vii, 11e, *p* = 0.2163).

## Discussion

EVs have received much attention for their potential as cell-free mediators of repair^2–4^. A variety of clinical trials have been conducted to test their potential to treat lung diseases from BPD to ARDS^30–32^. In early studies, EVs derived from mesenchymal stromal/stem cells (MSCs) were widely used, rationalising that stem cell derived EVs would have high regenerative capacity.

More recently, emerging evidence that the contents and effects of EVs are modulated by the source and physiological state of tissue they are derived from, has turned attention to tissue derived EVs. This shift from MSC to tissue derived EVs led to the discovery that EVs can sometimes drive disease by transferring pathogenic content to recipient cells/tissue^4,33^. It is equally clear however, that tissue derived EVs can alleviate lung disease^34^.

Most studies of EVs’ functional effects, have looked at their capacity to improve survival and/or to attenuate inflammation^35,36^ rather than testing their regenerative capacity. Aged lungs are significantly less capable of tissue repair and regeneration than young adult lungs. In this manuscript, we investigated the content and repair capacity of lung EVs derived from these two age groups, and we used precision-cut lung slice-derived EVs as tool to explore potential underlying factors responsible for the decreased repair capacity of aged lungs.

The characteristics of aged lungs, including altered ECM composition, stiffness and structure were clearly present in the aged PCLS used in this study^18,37,38^.

Assessment of the size and quantity of EVs produced from young and aged lung tissue showed that they were similar, regardless of age or ex-vivo acid injury. We also measured the mechanical properties of individual EVs using AFM; probing EV biomechanics is an emerging area of research for disease diagnosis and therapeutic strategies ^39,40^. The mechanical properties of EVs are known to be heterogeneous due to the differences in size, cargo and origin^40^. For example, whilst the presence of transmembrane proteins help maintain membrane structure and resistance to external pressure, membrane-associated proteins (e.g. Alix and Tsg101) can form networks or complexes on the membrane, increasing the stiffness of EVs^39^. It has also been shown that the stiffness of EVs obtained from healthy and injured cells varies^41^. In our study, we found that larger EVs tended to exhibit lower stiffness levels than smaller EVs; interestingly, this difference was most obvious in EVs obtained from both young and old injured PCLS. This data not only highlights the difference in biomechanical properties of larger and smaller EVs but also identified that the magnitude of these differences varies depending on the age of the source tissue and the presence of injury, emphasising the utility of measuring the biophysical properties of EVs.

We examined the effects of EVs obtained from young adult or aged lungs on cell survival. To investigate the repair capacity of EVs, we employed the acid injury and repair (AIR) model, in which the repair response of lung tissue to injury can be tracked and quantified in precision-cut lung slices, following administration of a spatially restricted area of injury ^27,28^ and we also analysed the small RNA content of each tissue derived EV group. From this analysis, we identified 5 miRNAs that were differentially regulated between young and aged EVs and subsequent bioinformatic analysis of the genes regulated by these miRNAs identified key signalling pathways and biological processes that these miRNAs can regulate.

Our data show that PCLS closely reflect the chronological age-related biology of *in vivo* lungs. Comparing whole PCLS that were injured to uninjured, control PCLS, we found that proSP-C^+^ ATII progenitor cells significantly increase in both young and aged PCLS, but the increase is smaller in aged lungs indicating a less robust repair response. The percentage of collagen positive staining was greater in aged lungs compared to young lungs and F-actin distribution was markedly different in aged and young adult PCLS, following acid injury, again reflecting known changes in aged and/or injured lungs from *in vivo* studies^37,42–46^.

Treatment with YU EVs invoked a marked reduction in apoptosis in both young adult and aged PCLS whereas AU EVs did not significantly reduce apoptosis in either young adult or aged PCLS. This indicates a functional difference in the effects of EVs derived from young adult or aged lung tissue, that likely reflects the altered composition of the lung tissue. We also found that the difference in percentage apoptosis between injured and uninjured regions of AIR-PCLS is less in aged PCLS than in young PCLS, perhaps as a result of the already higher baseline levels of apoptosis in aged tissue.

As expected, in the injured region of AIR-PCLS, the overall increase in proSP-C^+^ cells was much higher in young adult tissue than in aged lung tissue, consistent with diminished repair capacity of ageing lungs. In addition, collagen content was increased both in aged compared to young AIR-PCLS and in the injured regions compared to uninjured regions of AIR-PCLS, again, reflecting in vivo data showing that both ageing and disease frequently lead to increased collagen content^38,43,47^.

Investigation of the repair capacity of EVs using the AIR-PCLS model showed that EVs derived from young uninjured lungs had a significant pro-repair effect on aged lung tissue whereas treatment with YU EVs did not significantly increase the percentage of progenitor cells that were induced in the injured region of young adult AIR-PCLS. This indicates the contents of young EVs can enhance repair of aged lung tissue, in which the endogenous capacity for repair is diminished.

An interesting finding from examination of AIR-PCLS to compare the function of young uninjured EVs and aged uninjured EVs was that AU EVs caused a significant reduction in the percentage of proliferation in young adult AIR-PCLS only. This suggests either the presence of a factor/s that inhibit proliferation or the absence of a factor/s that enhances proliferation in AU EVs.

Having identified functional differences between EV groups, we went on to conduct a differential analysis of the miRNA content in young adult and aged lung derived EVs. We focused on miRNAs to obtain a broad overview of the programmes of gene changes that were altered in EVs secreted by aged vs young lungs^48,49^. Because miRNAs suppress expression of multiple gene targets, we reasoned that identifying whether any miRNAs were significantly altered in different EV populations and then analysing the genes that these miRNAs regulate, would provide a broad overview of genes and pathways driving the functional effects of different EV groups.

We consistently observed a greater effect on PCLS of treatment with YU EVs compared to AU EVs. For example, a greater reduction in collagen content in the injured region of young AIR-PCLS (Fig. 7c) and increased induction of progenitor cells in the injured region of aged AIR-PCLS (Fig. 5d). MiR-203-3p was downregulated in AU EVs, and this miRNA targets a number of genes important for ECM and actin cytoskeleton regulation such as Itga4, Cdc42 and Flnb, which may relate to the ECM and actin cytoskeleton changes present in aged lung tissue (Fig.2b, d).

Comparison of YI vs YU EVs revealed young PCLS secrete EVs with significantly reduced levels of miR-150-5p in response to injury. PANTHER pathway enrichment analysis of experimentally validated targets of miR-150-5p found nine significantly enriched signalling pathways. Three of these regulate cell survival and proliferation and the remaining pathways regulate lung development and repair, suggesting that some of the gene targets of miR-150-5p could be potential pro-repair candidates. Of particular interest is the overrepresentation of the Hedgehog signalling pathway genes among miR-150-5p targets. The Hedgehog pathway is critical for all stages of lung development including alveolar formation^50–52^. Many studies have also highlighted an important role for Shh in adult lung homeostasis and repair, though the exact nature of its role in the complex environment of the *in vivo* lungs is still not fully understood^53–56^. Moreover, Hedgeghog interacting protein (HHIP), an inhibitor of the hedgehog pathway, was found to be associated with susceptibility to COPD in a genome-wide association study (GWAS) comparing lung function in heavy smokers^57^.

The finding that miR-150-5p, which regulates multiple genes and pathways important for lung repair, was significantly down-regulated in EVs derived from young injured PCLS (YI EVs) compared to YU EVs, led us to further experiments. We assessed whether treatment with YI EVs, which carry different cargo to YU EVs, might show more potent effects on the induction of progenitor cells or reduction of collagen content of AIR-PCLS from young adult lungs. YI EVs had not been included in our earlier functional experiments (Figs 4-7) because our focus was on ageing. There was no significant difference between the effect of YI EVs and YU EVs although the effects of YI EVs on inhibiting the significant increase of collagen, in injured regions of AIR-PCLS, was marginally greater (Fig.11e). It could be that YI EV treatment would lead to altered effects on aged lung tissue, however, due to a lack of available mice, we did not include tissue from aged lungs in these later experiments.

Comparison of AI vs AU EVs revealed three miRNAs significantly down-regulated in AI EVs compared to AU EVs, but only let-7e-5p showed significant pathway enrichment among its target genes. Interestingly, like miR-150-5p targets, let-7e-5p targets were enriched for pathways involved in lung development and repair, including: PDGF, FGF and VEGF signalling pathways.

## Limitations

We identified differentially expressed miRNAs by sequencing RNAs isolated from the entire population of PCLS-derived EVs. The use of PCLS enabled us to study pooled EVs secreted by resident lung cell populations and ECM, closely mimicking the combination of EV-mediated signals found *in vivo*. However, analysing pooled EVs inevitably averages any potential particle-to-particle variability, and rarer miRNAs present only in some specific EVs may therefore have been masked. More advanced sequencing technology is needed to profile EV-RNA content at a single particle resolution. In addition, EVs from tissue could contain a mixture of ‘good/pro-repair’ and ‘harmful’ biological signals, especially from diseased/injured tissue. With the current technology, we cannot separate them but profiling the biological content of EVs at the single particle resolution would help identify the cargo that may be beneficial to drive tissue repair.

In this study, EV were isolated from PCLS-conditioned medium using size exclusion chromatography, a technique increasingly adopted for its higher EV yield, lower protein contamination and better preservation of EV structure and composition than traditional EV isolation method^58^. Nevertheless, it can co-isolate similarly sized particles such as lipoproteins and dilute the sample, requiring downstream concentration step through which we can lose EVs. Future studies will consider tangential flow filtration method for isolation of EVs from PCLS for higher yield of EVs, enabling higher concentrations of EVs used for treatment.

Based on the bioinformatic findings on miR-150-5p, we anticipated greater pro-repair effects of YI EVs than YU EVs; however, although both these EV groups reduced collagen deposition, neither reached a level of statistical significance. Future studies should examine whether treatment with higher EV concentrations might result in a stronger effect that leads to statistically significant reduction in collagen deposition. This might also reveal functional differences between injured and uninjured EVs.

In conclusion, this study shows that EVs derived from PCLS can be used as a tool to investigate the cell-to-cell signals that are released from lung tissue of different age or physiological state. We have shown that the physiology and function of EVs is significantly different in aged lungs, reflecting the biological differences in aged lung tissue. Interestingly, our data shows that EVs derived from young lung tissue have the capacity to enhance repair of aged lung tissue revealing that although aged lung tissue has a diminished repair capacity, it can mount a repair response if provided with the right biological cues. This finding has important implications for future therapeutic strategies to repair or regenerate aged lungs.

## Methods

### Sex as a biological variable

Both male and female mice were used to obtain lung samples. However, sex was not considered as a biological variable.

### PCLS generation and in vitro culture

Lungs were harvested from C57BL/6 mice, aged 10-12 weeks (“young”) or 18 months (“aged”) and inflated with 2% (w/v) low gelling point agarose solution (Sigma, A9414), as previously described^59^. Harvested lungs were kept in ice-cold Hank’s Balanced Salt Solution (HBBS; ThermoFisher Scientific, 14025-100) containing 1% 4-(2-hydroxyethyl)-1-piperazineethanesulfonic acid (HEPES; ThermoFisher Scientific) until slicing. Individual lung lobes were separated and sliced transversely at a thickness of 300 µm using an automated vibratome (Compresstome® VF-300-0, Precisionary Instruments) in HBBS/HEPES. PCLS of similar sizes were collected, to facilitate downstream analysis, and transferred to ice-cold serum-free media (Dulbecco’s Modified Eagle Medium [SF-DMEM; Sigma, 31966-201] with 1% penicillin-streptomycin [Merck Life Science, P0781-100ML). PCLS were washed three times with pre-warmed SF-DMEM and cultured in a humidified incubator at 37 ° C, 5% CO_2_ for up to 48 hrs.

### Whole-slice injury in mouse PCLS

Whole-slice acid-induced injury was performed on the same day as PCLS generation. After 2 hr incubation at 37°C, PCLS were washed three times with pre-warmed SF-DMEM. Immediately prior to use, 0.1 M hydrochloric acid (HCl) was prepared with ice-cold SF-DMEM. Five PCLS were incubated with 0.1 M HCl (500 μl/well) for 1 min. HCl acid was removed immediately after 1 min acid incubation and injured PCLS were then washed five times with pre-warmed SF-DMEM to eliminate any residual acid. Five uninjured PCLS per well were included as a control. Injured and uninjured PCLS were cultured in 12-well plates containing 5 PCLS per well in 2 ml pre-warmed EV isolation media (0.5% [w/v] bovine serum albumin [BSA; Sigma-Aldrich, A7030-100G] in SF-DMEM containing 1% penicillin-streptomycin; 2 mL/well) for 48 hrs at 37°C, 5% CO2.

### Extracellular vesicle isolation

Following the 48-hr incubation of PCLS, EV isolation media was collected from 30 PCLS (5 per well) for each physiological condition (aged/young, injured/uninjured). For each condition, media from all corresponding wells (2 ml/well) were combined to give a total of 12 mL per condition. All media samples were then centrifuged at 1,500 × *g* for 5 min at 4°C, followed by a second centrifugation at 5,000 × *g* for 15 min to remove cells, debris, and large particles. The supernatants were then transferred into a Vivaspin® 15R, 30,000 MWCO spin column (Sartorius, VS15RH22) and centrifuged at 5,000 x *g* for 20 min to concentrate the samples to < 500 μL. Prior to EV isolation by size exclusion chromatography, qEV original columns (IZON, 35 nm Gen 2, ICO-35) were washed by allowing 18 mL of 1× RNase-free PBS to run through the column matrix. The concentrated samples were added to the qEV original column and topped up with 1× DNase/RNase-free PBS once the samples entered the matrix of the column. After 2.5 mL of void volume, the first 1.5 mL of the flow-through was collected and transferred to a Amicon® Ultra-4 Centrifugal Filter (Sigma-Aldrich, UFC803024) column and centrifuged at 5,000 × *g* for 10 min to concentrate to < 100 μL. The concentrated EV samples were diluted in 1× RNase-free PBS to make up a total volume of 200 μL and stored at −80°C until further use.

### Nanoscale flow cytometry

Nanoscale flow cytometry was performed using the NanoFCM NanoAnalyzer to compare the size and concentration of EVs isolated from different groups of PCLS. Briefly, 0.25 μM fluorescent silica nanospheres of known concentration and silica nanosphere cocktail (68-155 nm) were used as concentration and size standards for parameter calibration as per manufacturer’s instructions. Then, 5 μL of EV samples were diluted in 1× DNase/RNase-free PBS at a ratio of 1:2 or 1:8 to meet the recommended concentration range of 2,000-20,000 particles/min. Each sample was measured for 1 min at 10 mW laser power, 10% SS decay and 1 kPa sampling pressure. The concentration of EV samples was determined at a gating of 40-140 nm. Two measurements were taken for each sample and averaged to obtain the final concentration and particle size.

### Atomic force microscopy (AFM)

Atomic force microscopy (AFM) was used to measure the size and mechanical properties of EVs, using a NanoWizard 4 Nanoscience AFM (JPK Bruker, Berlin, Germany). Undiluted EV samples (20 µl) were placed on freshly cleaved mica coated with 0.01% poly-L-lysine solution (Sigma-Aldrich, cat no. P8920). After 30 min incubation, mica was gently washed twice with PBS then samples were measured in liquid using the Qi™ mode, with a paraboloid-shaped cantilever (Nanosensors qp-BioAC-10) with spring constant 107.5 mN/m and sensitivity 9.477 nm/v, and setpoint 0.2 nN was used. Images were taken at 256 × 256 pixels with scan size ranging from 400 nm to 1.5 μm. JPK Data Processing software was used to analyse data and Young’s modulus micrographs were generated by batch processing with Hertz-fit and the stiffness of EVs were compared amongst similar sized EVs. The size of EVs was quantified by measuring the diameter using the cross-section tool through the middle of each particle. For each micrograph, one to ten EVs in the range of 40 – 350 nm in diameter were analysed by averaging Young’s modulus in the middle region of EVs covering 20 nm^2^ for EVs sized less than 100 nm, 60 nm^2^ for EVs sized between 100-200 nm, and 115 nm^2^ for EVs sized greater than 200 nm in diameter. Young’s modulus data from three to five micrographs were averaged for each replicate experiment and data from four separate experiments were analysed. Single EVs in scan size 400 nm were presented as representative image for each type of EVs.

### Isolation of RNA content from EVs (EV-RNA)

EVs derived from 30 PCLS for each condition were isolated and concentrated as described above. Total RNA was extracted from EVs with Direct-zol™ RNA Microprep kit (Zymo Research, R2060) as per manufacturer’s instructions. Briefly, concentrated EVs were mixed with 400 μL of Trizol and 500 μL of 100% ethanol, then centrifuged at 13,000 × *g* in a Zymo-Spin™ IC Column for 30 s. Next, samples were washed with RNA Wash Buffer and centrifuged at 13,000 × *g* for 30 s. The flow-through was discarded. Samples were then treated with 30 IU DNase I diluted in DNA Digestion Buffer for 15 mins at room temperature (RT). Samples were then washed twice with Direct-zol™ RNA PreWash buffer and once with RNA Wash Buffer, centrifuging and discarding the flow-through after each wash. Finally, RNA was eluted from the Zymo-Spin™ IC Column membrane by adding 10 μL of DNase/RNase-Free Water and centrifuging at 13,000 × *g* for 30 s. To maximise RNA collection, the RNA eluate was reapplied to the Zymo-Spin™ IC Column membrane and centrifuged again at 13,000 × *g* for 30 s. Total RNA from EVs derived from young and aged PCLS with/without acid injury was quantified using the Agilent 2200 High Sensitivity RNA ScreenTape system (Agilent technologies). RNA samples were stored at −80°C until sending to Omiics [https://omiics.com/] for small RNA-sequencing.

### Generation of the AIR model in mouse PCLS and EV treatment

One day prior to injury, 30% (w/v) Pluronic F-127 gel (Sigma, P2443-250G) was dissolved in ice-cold SF-DMEM with 1% penicillin-streptomycin overnight at 4°C. On the day of PCLS generation, 12M stock HCl was diluted in ice-cold SF-DMEM and Pluronic gel to obtain a final concentration of 0.1 M HCl containing 15% (w/v) Pluronic gel. PCLS were first washed three times with SF-DMEM. A single PCLS was positioned at the centre one well of a 24-well plate. A Pyrex® cloning cylinder (Corning, 3166-8, 8 × 8 mm) was coated on one side of the rim with Dow Corning® high-vacuum silicone grease (Sigma-Aldrich, Z273554) and placed onto the narrower end of a PCLS to create an isolated region of tissue. Appropriate force was applied to properly seal the region without damaging the tissue. Next, 500 μL of SF-DMEM was added to fill the outer area of the well to confirm that the seal was intact and no media entered the cylinder. Then 100 μL of 0.1 M HCl was added to the isolated region of PCLS within the cylinder and incubated for 1 min. During incubation, the colour of the media outside the cylinder was monitored to ensure no acid leakage. Immediately after injury, HCl acid was removed and the isolated region inside the cylinder was washed five times with SF-DMEM. The cylinder was then removed and the PCLS with spatially restricted acid-induced injury (“AIR-PCLS”) was transferred to a 48-well plate containing fresh SF-DMEM. Uninjured PCLS were also included as a control. Injured and uninjured PCLS were then treated with EVs as follows: EV treatments at a final concentration of 1 × 10^6^ particles/ml were made up in SF-DMEM. Individual injured and uninjured PCLS were incubated either in 150 μL of EVs (1 × 10^6^ particles/ml) or media containing no EVs (as a control) in a humidified incubator at 37°C, 5% CO2 for 48 hrs.

### Caspase-3 assay

After 48 hours of EV treatment, apoptotic cells in AIR PCLS were fluorescently labelled with NucView® 488 Caspase-3 Assay Kit (Biotium, 30029). Uninjured PCLS and PCLS treated with 70% methanol were included as controls. PCLS were incubated with 5 μM NucView 488 substrate in PBS in the dark at RT for 30 mins. After washing three times with PBS, slices were fixed with 4% PFA (ThermoFisher Scientific, 28908) for 15 min at RT and washed three times with PBS, before staining with DAPI (20 μL/mL, ThermoFisher Scientific, 62248) for 15 min at RT. PCLS were then washed once with PBS and mounted on glass slides using ProLong™ Gold Antifade Mountant (ThermoFisher Scientific, P36930). Slices were imaged within 24 hr using a Leica STELLARIS 5 inverted microscope with a 40 × oil immersion objective and LAS X software. Data were collected from three separate fields of view per injured and uninjured regions of each AIR-PCLS and three fields of view per control PCLS. Each independent experiment included three PCLS per condition (unless otherwise stated), with a total of four biological replicates (*n* = 4).

### Immunofluorescence staining of PCLS

PCLS were collected for immunostaining: (a) PCLS with/without whole-slice injury after removal of EV isolation media, and (b) control and AIR PCLS at the 48-hr time point after spatially restricted acid injury and/or EV treatment. PCLS were fixed with 4% PFA for 15 mins at RT and washed three times with PBS, 5 mins/time. Slices were then permeabilised by incubating with 0.5% Triton X-100 (Sigma-Aldrich, RES9690T-A101X) for 30 mins at RT with gentle rocking and washed three times as described above. Next, PCLS were incubated in PBS-BT (1% BSA, 0.2% Triton X-100 in PBS) for 1 hr at RT with gentle rocking to avoid non-specific binding. Subsequently, PCLS were incubated overnight at 4°C with anti-type I collagen-UNLB polyclonal (Southern Biotech, 1310-01, goat, 1:200) or a combination of anti-prosurfactant protein C polyclonal (proSP-C; Merck Millipore, AB3786, rabbit, 1:500) and Ki67 monoclonal (SolA15, ThermoFisher Scientific, 14-5698-82, rat, 1:500) primary antibodies in PBS-BT. For staining of F-actin, PCLS were incubated phalloidin-rhodamine pre-conjugated with Alexa Fluor 555 (ThermoFisher Scientific, A34055) 2 hrs at RT in the dark with gentle rocking. PCLS were then washed three times with PBS-BT. PCLS were incubated with secondary antibodies anti-rabbit Alexa-488 (ThermoFisher Scientific, A11008, goat) for proSP-C, anti-rat Alexa-568 (ThermoFisher Scientific, A11077, goat) for Ki67, or anti-goat Alexa-594 (ThermoFisher Scientific, A11058, donkey) for collagen, diluted 1:500 in PBS-BT, for 2 hrs at RT in the dark with gentle rocking. For staining of F-actin, PCLS were incubated phalloidin-rhodamine conjugated with Alexa Fluor 555 conjugates (ThermoFisher Scientific, A34055, 1:500) for 2 hrs at RT in the dark with gentle rocking.

Slices were washed three times with PBS-BT, 5 mins per wash, before staining with DAPI (2 μg/mL) for 15 mins in the dark at RT with gentle rocking. Finally, PCLS were washed twice with 1X PBS and mounted on glass slides using ProLong™ Gold Antifade Mountant.

Mounted slices were left overnight to set at RT in the dark and kept at 4°C until imaging.

### Imaging and quantification of immunofluorescence staining in PCLS

Images of PCLS were taken using a Leica STELLARIS 5 inverted microscope with a 40 × oil immersion objective or a 63 × oil immersion objective and LAS X software, or a Zeiss Axio Observer widefield microscope using a 40 × air objective with ApoTome and Zen software (blue edition). Data were collected from three separate fields of view per slice for each control/uninjured PCLS and PCLS with whole-slice injury, and from three separate fields of view per injured and uninjured region of each AIR PCLS. Each independent experiment included three PCLS per condition (unless otherwise stated), with a total of three biological replicates (*n* = 3).

Percentages of proSP-C^+^ and Ki67^+^ cells in images taken at 40 × magnification were manually quantified using ImageJ. Area of type I collagen deposition was quantified using an in-house ImageJ macro designed by Steve Rothery (Facility for Imaging by Light Microscopy, Imperial College London) to quantify percentage collagen area relative to total tissue area. The percentage area of type I collagen in total tissue area was calculated in images taken at either 40 × magnification (for AIR PCLS) or 63 × magnification (for PCLS with whole-slice injury).

### Small RNA-sequencing and bioinformatic analysis

Small RNA-sequencing, differential expression analysis and miRNA targeting analysis were performed externally by Omiics. MiRNAs were considered significantly differentially expressed if their adjusted *p* < 0.05 and if their absolute log2 (fold change) ≥ 1.

Experimentally validated target genes of each differentially expressed miRNAs were obtained by miRNA targeting analysis using *multimir* package based on miRNA-target interaction databases miRecords^60^, miRTarBase^61^ and TarBase^62^ with a cutoff of adjusted *p* value < 0.05. PANTHER pathway overrepresentation analysis was performed on pantherdb.org (ver 19.0)^63^ to identify signaling pathways overrepresented in target genes of each differentially expressed miRNA. The complete *Mus musculus* genome was used as reference. Fisher’s exact test was applied, followed by the Benjamini-Hochberg procedure, and pathways with FDR < 0.05 were considered to be significantly overrepresented. Network plots were generated for top 10 or all (if total number of overrepresented pathways is less than 10) overrepresented pathways using *ggplot2* and *ClusterProfiler4.0* packages in RStudio (ver 4.3.1).

### Statistical analysis

Data were analysed using GraphPad Prism Version 10.0.3 (217). Data were presented as mean value of each technical replicate (3 per biological replicate/experiment repeat) ± the standard error of mean (S.E.M.). Statistical significance was determined with either Mann-Whitney test or two-way ANOVA test followed by Tukey’s multiple comparison test. *P* value < 0.05 was considered as statistically significant. The number of biological replicate (n) is specified in each respective figure legend.

### Study approval

All animal procedures were conducted in compliance with the approval obtained from the South Kensington Animal Welfare and Ethical Review Board committee at Imperial College London.

### Data Availability

The data supporting the findings of this study are available in the Supporting Data Values file.

## Author Contributions

Q.C., P.T., and S.K. conducted the experiments and performed data analyses. Q.C., S.K., and C.H.D. prepared figures and wrote the manuscript. C.D.N. and F.J.C. provided materials for experiments with aged mouse lung tissue. S.K. and C.H.D. conceived and supervised the project and provided funds. All authors have reviewed and approved the final manuscript.

