## Supplementary data for "Lung derived extracellular vesicles have distinct pro-repair effects, depending on the age of their source tissue"

### Supplementary Data 1

| Downregulated in Aged Control compared to Young Control | | | | | | | | | |
|---|---|---|---|---|---|---|---|---|---|
| miR-203-3p targets | | | | | | | | | |
| database | mature_mirna_acc | mature_mirna_id | target_symbol | target_entrez | target_ensembl | experiment | support_type | pubmed_id | type |
| mirtarbase | MIMAT0000236 | mmu-miR-203-3p | ZNF148 | 7707 | ENSG00000163848 | Luciferase reporter assay//Microarray//qRT-PCR//Western blot | Functional MTI | 22842794 | validated |
| mirtarbase | MIMAT0000236 | mmu-miR-203-3p | Cav1 | 12389 | ENSMUSG00000007655 | Immunoblot//Luciferase reporter assay//qRT-PCR | Functional MTI | 22421148 | validated |
| mirtarbase | MIMAT0000236 | mmu-miR-203-3p | Cp | 12870 | ENSMUSG00000003617 | HITS-CLIP | Functional MTI (Weak) | 23142080 | validated |
| mirtarbase | MIMAT0000236 | mmu-miR-203-3p | Cpox | 12892 | ENSMUSG00000022742 | HITS-CLIP | Functional MTI (Weak) | 25083871 | validated |
| mirtarbase | MIMAT0000236 | mmu-miR-203-3p | Itga4 | 16401 | ENSMUSG00000027009 | HITS-CLIP | Functional MTI (Weak) | 25083871 | validated |
| mirtarbase | MIMAT0000236 | mmu-miR-203-3p | Myd88 | 17874 | ENSMUSG00000032508 | Luciferase reporter assay | Functional MTI | 25723469 | validated |
| mirtarbase | MIMAT0000236 | mmu-miR-203-3p | Prkcb | 18751 | ENSMUSG00000052889 | HITS-CLIP | Functional MTI (Weak) | 21258322 | validated |
| mirtarbase | MIMAT0000236 | mmu-miR-203-3p | Uri1 | 19777 | ENSMUSG00000030421 | HITS-CLIP | Functional MTI (Weak) | 25083871 | validated |
| mirtarbase | MIMAT0000236 | mmu-miR-203-3p | Rs1 | 20147 | ENSMUSG00000031293 | HITS-CLIP | Functional MTI (Weak) | 21258322 | validated |
| mirtarbase | MIMAT0000236 | mmu-miR-203-3p | Rs1 | 20147 | ENSMUSG00000031293 | HITS-CLIP | Functional MTI (Weak) | 19536157 | validated |
| mirtarbase | MIMAT0000236 | mmu-miR-203-3p | Stxbp4 | 20913 | ENSMUSG00000020546 | HITS-CLIP | Functional MTI (Weak) | 25083871 | validated |
| mirtarbase | MIMAT0000236 | mmu-miR-203-3p | Tlr4 | 21898 | | HITS-CLIP | Functional MTI (Weak) | 23142080 | validated |
| mirtarbase | MIMAT0000236 | mmu-miR-203-3p | Trp63 | 22061 | ENSMUSG00000022510 | Luciferase reporter assay//qRT-PCR//Western blot | Functional MTI | 18311128 | validated |
| mirtarbase | MIMAT0000236 | mmu-miR-203-3p | Trp63 | 22061 | ENSMUSG00000022510 | Luciferase reporter assay//Western blot | Functional MTI | 20216554 | validated |
| mirtarbase | MIMAT0000236 | mmu-miR-203-3p | Trp63 | 22061 | ENSMUSG00000022510 | Immunoblot//Luciferase reporter assay//qRT-PCR | Functional MTI | 22421148 | validated |
| mirtarbase | MIMAT0000236 | mmu-miR-203-3p | Ung | 22256 | ENSMUSG00000029591 | HITS-CLIP | Functional MTI (Weak) | 25083871 | validated |

| | | | | | | | | | |
|---|---|---|---|---|---|---|---|---|---|
| mirtarbase | MIMAT0000236 | mmu-miR-203-3p | Ung | 22256 | ENSMUSG00000029591 | HITS-CLIP | Functional MTI (Weak) | 23597149 | validated |
| mirtarbase | MIMAT0000236 | mmu-miR-203-3p | Ung | 22256 | ENSMUSG00000029591 | HITS-CLIP | Functional MTI (Weak) | 23142080 | validated |
| mirtarbase | MIMAT0000236 | mmu-miR-203-3p | Mapk8 | 26419 | ENSMUSG00000021936 | HITS-CLIP | Functional MTI (Weak) | 25083871 | validated |
| mirtarbase | MIMAT0000236 | mmu-miR-203-3p | Commd 10 | 69456 | ENSMUSG00000042705 | HITS-CLIP | Functional MTI (Weak) | 25083871 | validated |
| mirtarbase | MIMAT0000236 | mmu-miR-203-3p | Gtpbp3 | 70359 | ENSMUSG00000007610 | HITS-CLIP | Functional MTI (Weak) | 21258322 | validated |
| mirtarbase | MIMAT0000236 | mmu-miR-203-3p | Ces2g | 72361 | ENSMUSG00000031877 | HITS-CLIP | Functional MTI (Weak) | 25083871 | validated |
| mirtarbase | MIMAT0000236 | mmu-miR-203-3p | Cyld | 74256 | ENSMUSG00000036712 | HITS-CLIP | Functional MTI (Weak) | 23142080 | validated |
| mirtarbase | MIMAT0000236 | mmu-miR-203-3p | Fgf16 | 80903 | | HITS-CLIP | Functional MTI (Weak) | 25083871 | validated |
| mirtarbase | MIMAT0000236 | mmu-miR-203-3p | Pcdhb1 9 | 93890 | | HITS-CLIP | Functional MTI (Weak) | 23597149 | validated |
| mirtarbase | MIMAT0000236 | mmu-miR-203-3p | Snora6 2 | 104433 | | Luciferase reporter assay//qRT-PCR | Functional MTI | 23175673 | validated |
| mirtarbase | MIMAT0000236 | mmu-miR-203-3p | Thsd7b | 210417 | ENSMUSG00000042581 | HITS-CLIP | Functional MTI (Weak) | 25083871 | validated |
| mirtarbase | MIMAT0000236 | mmu-miR-203-3p | Ccdc11 0 | 212392 | ENSMUSG00000071104 | HITS-CLIP | Functional MTI (Weak) | 23142080 | validated |
| mirtarbase | MIMAT0000236 | mmu-miR-203-3p | Srek1 | 218543 | ENSMUSG00000032621 | HITS-CLIP | Functional MTI (Weak) | 25083871 | validated |
| mirtarbase | MIMAT0000236 | mmu-miR-203-3p | Zfp281 | 226442 | ENSMUSG00000041483 | Luciferase reporter assay | Functional MTI | 18311128 | validated |
| mirtarbase | MIMAT0000236 | mmu-miR-203-3p | Katnal1 | 231912 | ENSMUSG00000041298 | HITS-CLIP | Functional MTI (Weak) | 25083871 | validated |
| mirtarbase | MIMAT0000236 | mmu-miR-203-3p | Frmd3 | 242506 | ENSMUSG00000049122 | HITS-CLIP | Functional MTI (Weak) | 25083871 | validated |
| mirtarbase | MIMAT0000236 | mmu-miR-203-3p | Frmd3 | 242506 | ENSMUSG00000049122 | HITS-CLIP | Functional MTI (Weak) | 23597149 | validated |
| mirtarbase | MIMAT0000236 | mmu-miR-203-3p | Frmd3 | 242506 | ENSMUSG00000049122 | HITS-CLIP | Functional MTI (Weak) | 23142080 | validated |
| mirtarbase | MIMAT0000236 | mmu-miR-203-3p | Frmd3 | 242506 | ENSMUSG00000049122 | HITS-CLIP | Functional MTI (Weak) | 21258322 | validated |

| | | | | | | | | | |
|---|---|---|---|---|---|---|---|---|---|
| mirtarbase | MIMAT0000236 | mmu-miR-203-3p | Bnc2 | 242509 | ENSMUSG00000028487 | HITS-CLIP | Functional MTI (Weak) | 25083871 | validated |
| mirtarbase | MIMAT0000236 | mmu-miR-203-3p | Zfp300 | 245368 | ENSMUSG00000031079 | HITS-CLIP | Functional MTI (Weak) | 25083871 | validated |
| mirtarbase | MIMAT0000236 | mmu-miR-203-3p | Alkbh5 | 268420 | ENSMUSG00000042650 | HITS-CLIP | Functional MTI (Weak) | 25083871 | validated |
| mirtarbase | MIMAT0000236 | mmu-miR-203-3p | Srsf12 | 272009 | ENSMUSG00000054679 | HITS-CLIP | Functional MTI (Weak) | 25083871 | validated |
| mirtarbase | MIMAT0000236 | mmu-miR-203-3p | H2bc21 | 319190 | ENSMUSG00000068854 | HITS-CLIP | Functional MTI (Weak) | 25083871 | validated |
| mirtarbase | MIMAT0000236 | mmu-miR-203-3p | Zmiz1 | 328365 | ENSMUSG00000007817 | HITS-CLIP | Functional MTI (Weak) | 23142080 | validated |
| mirtarbase | MIMAT0000236 | mmu-miR-203-3p | Rbm44 | 329207 | ENSMUSG00000070732 | Luciferase reporter assay//qRT-PCR//Western blot | Functional MTI | 22909339 | validated |
| mirtarbase | MIMAT0000236 | mmu-miR-203-3p | Ptchd4 | 627626 | ENSMUSG00000042256 | HITS-CLIP | Functional MTI (Weak) | 25083871 | validated |
| mirtarbase | MIMAT0000236 | mmu-miR-203-3p | Ptchd4 | 627626 | ENSMUSG00000042256 | HITS-CLIP | Functional MTI (Weak) | 23142080 | validated |
| tarbase | MIMAT0000236 | mmu-miR-203-3p | Ndufa9 | 66108 | ENSMUSG00000000399 | Degradome sequencing | positive | | validated |
| tarbase | MIMAT0000236 | mmu-miR-203-3p | Kdelr1 | 68137 | ENSMUSG00000002778 | Degradome sequencing | positive | | validated |
| tarbase | MIMAT0000236 | mmu-miR-203-3p | Cdip1 | 66626 | ENSMUSG00000004071 | Degradome sequencing | positive | | validated |
| tarbase | MIMAT0000236 | mmu-miR-203-3p | Pgrmc1 | 53328 | ENSMUSG00000006373 | Degradome sequencing | positive | | validated |
| tarbase | MIMAT0000236 | mmu-miR-203-3p | Pdk1 | 228026 | ENSMUSG00000006494 | Degradome sequencing | positive | | validated |
| tarbase | MIMAT0000236 | mmu-miR-203-3p | Cdc42 | 12540 | ENSMUSG00000006699 | Degradome sequencing//Degradome sequencing | positive | | validated |
| tarbase | MIMAT0000236 | mmu-miR-203-3p | Tgfbr1 | 21812 | ENSMUSG00000007613 | Degradome sequencing | positive | | validated |
| tarbase | MIMAT0000236 | mmu-miR-203-3p | Slc38a3 | 76257 | ENSMUSG00000010064 | Degradome sequencing//Degradome sequencing//Degradome sequencing | positive | | validated |
| tarbase | MIMAT0000236 | mmu-miR-203-3p | Steap4 | 117167 | ENSMUSG00000012428 | Degradome sequencing | positive | | validated |
| tarbase | MIMAT0000236 | mmu-miR-203-3p | Hspa8 | 15481 | ENSMUSG00000015656 | Degradome sequencing | positive | | validated |
| tarbase | MIMAT0000236 | mmu-miR-203-3p | Lamp2 | 16784 | ENSMUSG00000016534 | Degradome sequencing//Degradome sequencing//Degradome sequencing | positive | | validated |

| | | | | | | | | | |
|---|---|---|---|---|---|---|---|---|---|
| tarbase | MIMAT0000236 | mmu-miR-203-3p | Crk | 12928 | ENSMUSG00000017776 | Degradome sequencing//Degradome sequencing//Degradome sequencing//Degradome sequencing | positive | | validated |
| tarbase | MIMAT0000236 | mmu-miR-203-3p | Sparc | 20692 | ENSMUSG00000018593 | Degradome sequencing | positive | | validated |
| tarbase | MIMAT0000236 | mmu-miR-203-3p | Acsl1 | 14081 | ENSMUSG00000018796 | Degradome sequencing | positive | | validated |
| tarbase | MIMAT0000236 | mmu-miR-203-3p | Psen1 | 19164 | ENSMUSG00000019969 | Degradome sequencing | positive | | validated |
| tarbase | MIMAT0000236 | mmu-miR-203-3p | Nedd1 | 17997 | ENSMUSG00000019988 | Degradome sequencing | positive | | validated |
| tarbase | MIMAT0000236 | mmu-miR-203-3p | Xpo1 | 103573 | ENSMUSG00000020290 | Degradome sequencing | positive | | validated |
| tarbase | MIMAT0000236 | mmu-miR-203-3p | Ppp2ca | | ENSMUSG00000020349 | Degradome sequencing | positive | | validated |
| tarbase | MIMAT0000236 | mmu-miR-203-3p | Canx | 12330 | ENSMUSG00000020368 | Degradome sequencing | positive | | validated |
| tarbase | MIMAT0000236 | mmu-miR-203-3p | Lpin1 | 14245 | ENSMUSG00000020593 | Degradome sequencing | positive | | validated |
| tarbase | MIMAT0000236 | mmu-miR-203-3p | Rffl | 67338 | ENSMUSG00000020696 | Degradome sequencing | positive | | validated |
| tarbase | MIMAT0000236 | mmu-miR-203-3p | Susd6 | 217684 | ENSMUSG00000021133 | Degradome sequencing | positive | | validated |
| tarbase | MIMAT0000236 | mmu-miR-203-3p | Lgmn | 19141 | ENSMUSG00000021190 | Degradome sequencing | positive | | validated |
| tarbase | MIMAT0000236 | mmu-miR-203-3p | Yy1 | 22632 | ENSMUSG00000021264 | Degradome sequencing | positive | | validated |
| tarbase | MIMAT0000236 | mmu-miR-203-3p | Nsd1 | 18193 | ENSMUSG00000021488 | Degradome sequencing | positive | | validated |
| tarbase | MIMAT0000236 | mmu-miR-203-3p | Samd8 | 67630 | ENSMUSG00000021770 | Degradome sequencing | positive | | validated |
| tarbase | MIMAT0000236 | mmu-miR-203-3p | Parg | | ENSMUSG00000021911 | Degradome sequencing | positive | | validated |
| tarbase | MIMAT0000236 | mmu-miR-203-3p | Atad2 | 70472 | ENSMUSG00000022360 | Degradome sequencing | positive | | validated |
| tarbase | MIMAT0000236 | mmu-miR-203-3p | Slc38a4 | 69354 | ENSMUSG00000022464 | Degradome sequencing//Degradome sequencing | positive | | validated |
| tarbase | MIMAT0000236 | mmu-miR-203-3p | Atp13a3 | 224088 | ENSMUSG00000022533 | Degradome sequencing | positive | | validated |
| tarbase | MIMAT0000236 | mmu-miR-203-3p | Son | 20658 | ENSMUSG00000022961 | Degradome sequencing | positive | | validated |
| tarbase | MIMAT0000236 | mmu-miR-203-3p | Lpin2 | 64898 | ENSMUSG00000024052 | Degradome sequencing | positive | | validated |
| tarbase | MIMAT0000236 | mmu-miR-203-3p | Vapa | 30960 | ENSMUSG00000024091 | Degradome sequencing | positive | | validated |
| tarbase | MIMAT0000236 | mmu-miR-203-3p | Tcf7l2 | 21416 | ENSMUSG00000024985 | Degradome sequencing | positive | | validated |
| tarbase | MIMAT0000236 | mmu-miR-203-3p | Hhex | 15242 | ENSMUSG00000024986 | Degradome sequencing//Degradome sequencing//Degradome sequencing//Degradome sequencing | positive | | validated |
| tarbase | MIMAT0000236 | mmu-miR-203-3p | Flnb | 286940 | ENSMUSG00000025278 | Degradome sequencing | positive | | validated |
| tarbase | MIMAT0000236 | mmu-miR-203-3p | Prim1 | 19075 | ENSMUSG00000025395 | Degradome sequencing | positive | | validated |
| tarbase | MIMAT0000236 | mmu-miR-203-3p | Shisa5 | 66940 | ENSMUSG00000025647 | Degradome sequencing | positive | | validated |
| tarbase | MIMAT0000236 | mmu-miR-203-3p | Idh1 | 15926 | ENSMUSG00000025950 | Degradome sequencing | positive | | validated |
| tarbase | MIMAT0000236 | mmu-miR-203-3p | Slc40a1 | 53945 | ENSMUSG00000025993 | Degradome sequencing | positive | | validated |

| | | | | | | | | | |
|---|---|---|---|---|---|---|---|---|---|
| tarbase | MIMAT0000236 | mmu-miR-203-3p | Tsn | | ENSMUSG00000026374 | Degradome sequencing | positive | | validated |
| tarbase | MIMAT0000236 | mmu-miR-203-3p | Slc25a25 | 227731 | ENSMUSG00000026819 | Degradome sequencing | positive | | validated |
| tarbase | MIMAT0000236 | mmu-miR-203-3p | Rbms1 | 56878 | ENSMUSG00000026970 | Degradome sequencing//Degradome sequencing | positive | | validated |
| tarbase | MIMAT0000236 | mmu-miR-203-3p | Cat | 12359 | ENSMUSG00000027187 | Degradome sequencing | positive | | validated |
| tarbase | MIMAT0000236 | mmu-miR-203-3p | Trpm7 | 58800 | ENSMUSG00000027365 | Degradome sequencing//Degradome sequencing//Degradome sequencing | positive | | validated |
| tarbase | MIMAT0000236 | mmu-miR-203-3p | Stx16 | 228960 | ENSMUSG00000027522 | Degradome sequencing | positive | | validated |
| tarbase | MIMAT0000236 | mmu-miR-203-3p | Kcnq5 | 226922 | ENSMUSG00000028033 | Degradome sequencing | positive | | validated |
| tarbase | MIMAT0000236 | mmu-miR-203-3p | Adh5 | 11532 | ENSMUSG00000028138 | Degradome sequencing | positive | | validated |
| tarbase | MIMAT0000236 | mmu-miR-203-3p | Ptbp3 | 230257 | ENSMUSG00000028382 | Degradome sequencing | positive | | validated |
| tarbase | MIMAT0000236 | mmu-miR-203-3p | Plpp3 | 67916 | ENSMUSG00000028517 | Degradome sequencing//Degradome sequencing | positive | | validated |
| tarbase | MIMAT0000236 | mmu-miR-203-3p | Sepsecs | 211006 | ENSMUSG00000029173 | Degradome sequencing | positive | | validated |
| tarbase | MIMAT0000236 | mmu-miR-203-3p | Paics | 67054 | ENSMUSG00000029247 | Degradome sequencing | positive | | validated |
| tarbase | MIMAT0000236 | mmu-miR-203-3p | Foxp2 | 114142 | ENSMUSG00000029563 | Degradome sequencing | positive | | validated |
| tarbase | MIMAT0000236 | mmu-miR-203-3p | Aass | 30956 | ENSMUSG00000029695 | Degradome sequencing | positive | | validated |
| tarbase | MIMAT0000236 | mmu-miR-203-3p | Pon3 | 269823 | ENSMUSG00000029759 | Degradome sequencing | positive | | validated |
| tarbase | MIMAT0000236 | mmu-miR-203-3p | Lrig1 | 16206 | ENSMUSG00000030029 | Degradome sequencing//Degradome sequencing//Degradome sequencing | positive | | validated |
| tarbase | MIMAT0000236 | mmu-miR-203-3p | Rps6ka3 | 110651 | ENSMUSG00000031309 | Degradome sequencing | positive | | validated |
| tarbase | MIMAT0000236 | mmu-miR-203-3p | Sorbs2 | 234214 | ENSMUSG00000031626 | Degradome sequencing | positive | | validated |
| tarbase | MIMAT0000236 | mmu-miR-203-3p | Dnaja2 | 56445 | ENSMUSG00000031701 | Degradome sequencing | positive | | validated |
| tarbase | MIMAT0000236 | mmu-miR-203-3p | Elovl5 | 68801 | ENSMUSG00000032349 | Degradome sequencing | positive | | validated |
| tarbase | MIMAT0000236 | mmu-miR-203-3p | Tomm6 | | ENSMUSG00000033475 | Degradome sequencing | positive | | validated |
| tarbase | MIMAT0000236 | mmu-miR-203-3p | Slc16a2 | 20502 | ENSMUSG00000033965 | Degradome sequencing | positive | | validated |
| tarbase | MIMAT0000236 | mmu-miR-203-3p | Pds5b | 100710 | ENSMUSG00000034021 | Degradome sequencing | positive | | validated |
| tarbase | MIMAT0000236 | mmu-miR-203-3p | Ctnnd1 | 12388 | ENSMUSG00000034101 | Degradome sequencing | positive | | validated |

| | | | | | | | | | |
|---|---|---|---|---|---|---|---|---|---|
| tarbase | MIMAT0000236 | mmu-miR-203-3p | Arhgap 5 | 11855 | ENSMUSG00000035133 | Degradome sequencing | positive | | validated |
| tarbase | MIMAT0000236 | mmu-miR-203-3p | Hykk | 235386 | ENSMUSG00000035878 | Degradome sequencing | positive | | validated |
| tarbase | MIMAT0000236 | mmu-miR-203-3p | Rnf139 | 75841 | ENSMUSG00000037075 | Degradome sequencing | positive | | validated |
| tarbase | MIMAT0000236 | mmu-miR-203-3p | 4932438A13Rik | 229227 | ENSMUSG00000037270 | Degradome sequencing | positive | | validated |
| tarbase | MIMAT0000236 | mmu-miR-203-3p | Fam13a | | ENSMUSG00000037709 | Degradome sequencing | positive | | validated |
| tarbase | MIMAT0000236 | mmu-miR-203-3p | Rdh7 | 54150 | ENSMUSG00000040134 | Degradome sequencing | positive | | validated |
| tarbase | MIMAT0000236 | mmu-miR-203-3p | Arhgap 32 | 330914 | ENSMUSG00000041444 | Degradome sequencing | positive | | validated |
| tarbase | MIMAT0000236 | mmu-miR-203-3p | Alkbh5 | 268420 | ENSMUSG00000042650 | Degradome sequencing | positive | | validated |
| tarbase | MIMAT0000236 | mmu-miR-203-3p | Insig1 | 231070 | ENSMUSG00000045294 | Degradome sequencing//Degradome sequencing//Degradome sequencing | positive | | validated |
| tarbase | MIMAT0000236 | mmu-miR-203-3p | Slc25a2 3 | 66972 | ENSMUSG00000046329 | Degradome sequencing | positive | | validated |
| tarbase | MIMAT0000236 | mmu-miR-203-3p | Aff4 | 93736 | ENSMUSG00000049470 | Degradome sequencing//Degradome sequencing//Degradome sequencing | positive | | validated |
| tarbase | MIMAT0000236 | mmu-miR-203-3p | Maml1 | 103806 | ENSMUSG00000050567 | Degradome sequencing | positive | | validated |
| tarbase | MIMAT0000236 | mmu-miR-203-3p | Pcbp1 | 23983 | ENSMUSG00000051695 | Degradome sequencing | positive | | validated |
| tarbase | MIMAT0000236 | mmu-miR-203-3p | Hbb-bs | 15129 | ENSMUSG00000052305 | Degradome sequencing | positive | | validated |
| tarbase | MIMAT0000236 | mmu-miR-203-3p | Gckr | 231103 | ENSMUSG00000059434 | Degradome sequencing | positive | | validated |
| tarbase | MIMAT0000236 | mmu-miR-203-3p | Nup98 | 269966 | ENSMUSG00000063550 | Degradome sequencing | positive | | validated |
| tarbase | MIMAT0000236 | mmu-miR-203-3p | Cyp4a1 2a | 277753 | ENSMUSG00000066071 | Degradome sequencing | positive | | validated |
| tarbase | MIMAT0000236 | mmu-miR-203-3p | Irgm2 | 54396 | ENSMUSG00000069874 | Degradome sequencing | positive | | validated |
| tarbase | MIMAT0000236 | mmu-miR-203-3p | Rbm47 | 245945 | ENSMUSG00000070780 | Degradome sequencing | positive | | validated |
| tarbase | MIMAT0000236 | mmu-miR-203-3p | Ganab | 14376 | ENSMUSG00000071650 | Degradome sequencing | positive | | validated |
| tarbase | MIMAT0000236 | mmu-miR-203-3p | Gm587 8 | 545861 | ENSMUSG00000072952 | Degradome sequencing | positive | | validated |
| tarbase | MIMAT0000236 | mmu-miR-203-3p | Trim71 | | ENSMUSG00000079259 | Degradome sequencing | positive | | validated |

| Downregulated in Young Injured compared to Young Control | | | | | | | | | |
|---|---|---|---|---|---|---|---|---|---|
| miR-150-5p targets | | | | | | | | | |
| database | mature_mirna_acc | mature_mirna_id | target_symbol | target_entrez | target_ensembl | experiment | support_type | pubmed_id | type |
| mirecords | MIMAT0000160 | mmu-miR-150-5p | Myb | 17863 | ENSMUSG00000019982 | activity assay//Luciferase activity assay | | 17923094 | validated |
| mirecords | MIMAT0000160 | mmu-miR-150-5p | Myb | 17863 | ENSMUSG00000019982 | activity assay//Luciferase activity assay | | 17923094 | validated |
| mirecords | MIMAT0000160 | mmu-miR-150-5p | Vegfa | 22339 | ENSMUSG00000023951 | Western blot | | 18500251 | validated |
| mirecords | MIMAT0000160 | mmu-miR-150-5p | Pdgfb | 18591 | ENSMUSG00000000489 | Western blot | | 18500251 | validated |
| mirecords | MIMAT0000160 | mmu-miR-150-5p | Cxcr4 | 12767 | ENSMUSG00000045382 | Western blot//Luciferase activity assay | | 22039399 | validated |
| mirtarbase | MIMAT0000160 | mmu-miR-150-5p | Abl2 | 11352 | ENSMUSG00000026596 | HITS-CLIP | Functional MTI (Weak) | 21258322 | validated |
| mirtarbase | MIMAT0000160 | mmu-miR-150-5p | Akt1 | 11651 | ENSMUSG00000001729 | Luciferase reporter assay//qRT-PCR//Western blot | Functional MTI | 26196694 | validated |
| mirtarbase | MIMAT0000160 | mmu-miR-150-5p | Zfp36l1 | 12192 | ENSMUSG00000021127 | Luciferase reporter assay//qRT-PCR | Functional MTI | 26833392 | validated |
| mirtarbase | MIMAT0000160 | mmu-miR-150-5p | Chic1 | 12212 | ENSMUSG00000031327 | HITS-CLIP | Functional MTI (Weak) | 21258322 | validated |
| mirtarbase | MIMAT0000160 | mmu-miR-150-5p | Btc | 12223 | ENSMUSG00000082361 | HITS-CLIP | Functional MTI (Weak) | 21258322 | validated |
| mirtarbase | MIMAT0000160 | mmu-miR-150-5p | Btrc | 12234 | ENSMUSG00000025217 | HITS-CLIP | Functional MTI (Weak) | 21258322 | validated |
| mirtarbase | MIMAT0000160 | mmu-miR-150-5p | Camk4 | 12326 | ENSMUSG00000038128 | HITS-CLIP | Functional MTI (Weak) | 25083871 | validated |
| mirtarbase | MIMAT0000160 | mmu-miR-150-5p | Camk4 | 12326 | ENSMUSG00000038128 | HITS-CLIP | Functional MTI (Weak) | 21258322 | validated |
| mirtarbase | MIMAT0000160 | mmu-miR-150-5p | Cbl | 12402 | ENSMUSG00000034342 | Flow//Luciferase reporter assay//Microarray//qRT-PCR | Functional MTI | 23604034 | validated |
| mirtarbase | MIMAT0000160 | mmu-miR-150-5p | Cxcr4 | 12767 | ENSMUSG00000045382 | ELISA//Western blot//Luciferase reporter assay//qRT-PCR//Immunohistochemistry | Functional MTI | 25466411 | validated |
| mirtarbase | MIMAT0000160 | mmu-miR-150-5p | Col4a5 | 12830 | ENSMUSG00000031274 | HITS-CLIP | Functional MTI (Weak) | 25083871 | validated |
| mirtarbase | MIMAT0000160 | mmu-miR-150-5p | Cycs | 13063 | ENSMUSG00000063694 | HITS-CLIP | Functional MTI (Weak) | 25083871 | validated |
| mirtarbase | MIMAT0000160 | mmu-miR-150-5p | Egr2 | 13654 | ENSMUSG00000037868 | Flow//Luciferase reporter assay//Microarray//qRT-PCR | Functional MTI | 23604034 | validated |
| mirtarbase | MIMAT0000160 | mmu-miR-150-5p | Egr2 | 13654 | ENSMUSG00000116458 | Flow//Luciferase reporter assay//Microarray//qRT-PCR | Functional MTI | 23604034 | validated |
| mirtarbase | MIMAT0000160 | mmu-miR-150-5p | Eif4g2 | 13690 | ENSMUSG00000005610 | HITS-CLIP | Functional MTI (Weak) | 21258322 | validated |
| mirtarbase | MIMAT0000160 | mmu-miR-150-5p | Elf1 | 13709 | ENSMUSG00000036461 | HITS-CLIP | Functional MTI (Weak) | 21258322 | validated |
| mirtarbase | MIMAT0000160 | mmu-miR-150-5p | Elk1 | 13712 | ENSMUSG00000009406 | Luciferase reporter assay//qRT-PCR | Functional MTI | 26833392 | validated |

| | | | | | | | | | |
|---|---|---|---|---|---|---|---|---|---|
| mirtarbase | MIMAT0000160 | mmu-miR-150-5p | F2r | 14062 | ENSMUSG00000048376 | HITS-CLIP | Functional MTI (Weak) | 21258322 | validated |
| mirtarbase | MIMAT0000160 | mmu-miR-150-5p | Gad2 | 14417 | ENSMUSG00000026787 | HITS-CLIP | Functional MTI (Weak) | 25083871 | validated |
| mirtarbase | MIMAT0000160 | mmu-miR-150-5p | Gad2 | 14417 | ENSMUSG00000026787 | HITS-CLIP | Functional MTI (Weak) | 21258322 | validated |
| mirtarbase | MIMAT0000160 | mmu-miR-150-5p | Gpr12 | 14738 | ENSMUSG00000041468 | HITS-CLIP | Functional MTI (Weak) | 25083871 | validated |
| mirtarbase | MIMAT0000160 | mmu-miR-150-5p | Grm1 | 14816 | ENSMUSG00000019828 | HITS-CLIP | Functional MTI (Weak) | 21258322 | validated |
| mirtarbase | MIMAT0000160 | mmu-miR-150-5p | H2-T3 | 15043 | ENSMUSG00000054128 | HITS-CLIP | Functional MTI (Weak) | 21258322 | validated |
| mirtarbase | MIMAT0000160 | mmu-miR-150-5p | Hyal1 | 15586 | ENSMUSG00000010051 | HITS-CLIP | Functional MTI (Weak) | 21258322 | validated |
| mirtarbase | MIMAT0000160 | mmu-miR-150-5p | Ifngr2 | 15980 | | HITS-CLIP | Functional MTI (Weak) | 21258322 | validated |
| mirtarbase | MIMAT0000160 | mmu-miR-150-5p | Kif5b | 16573 | ENSMUSG00000006740 | HITS-CLIP | Functional MTI (Weak) | 21258322 | validated |
| mirtarbase | MIMAT0000160 | mmu-miR-150-5p | Mc1r | 17199 | ENSMUSG00000074037 | HITS-CLIP | Functional MTI (Weak) | 21258322 | validated |
| mirtarbase | MIMAT0000160 | mmu-miR-150-5p | Mpv17 | 17527 | ENSMUSG00000107283 | HITS-CLIP | Functional MTI (Weak) | 21258322 | validated |
| mirtarbase | MIMAT0000160 | mmu-miR-150-5p | Myb | 17863 | ENSMUSG00000019982 | Luciferase reporter assay | Functional MTI | 17923094 | validated |
| mirtarbase | MIMAT0000160 | mmu-miR-150-5p | Myb | 17863 | ENSMUSG00000019982 | Luciferase reporter assay//qRT-PCR | Functional MTI | 23604034 | validated |
| mirtarbase | MIMAT0000160 | mmu-miR-150-5p | Myb | 17863 | ENSMUSG00000019982 | Luciferase reporter assay//qRT-PCR | Functional MTI | 26833392 | validated |
| mirtarbase | MIMAT0000160 | mmu-miR-150-5p | Myd88 | 17874 | ENSMUSG00000032508 | Immunoblot//Immunohistochemistry//Luciferase reporter assay//Microarray//qRT-PCR | Functional MTI | 23166298 | validated |
| mirtarbase | MIMAT0000160 | mmu-miR-150-5p | Nfic | 18029 | ENSMUSG00000055053 | HITS-CLIP | Functional MTI (Weak) | 21258322 | validated |
| mirtarbase | MIMAT0000160 | mmu-miR-150-5p | Emc8 | 18117 | ENSMUSG00000031819 | HITS-CLIP | Functional MTI (Weak) | 21258322 | validated |
| mirtarbase | MIMAT0000160 | mmu-miR-150-5p | Notch4 | 18132 | ENSMUSG00000015468 | Luciferase reporter assay//Western blot | Non-Functional MTI | 18500251 | validated |
| mirtarbase | MIMAT0000160 | mmu-miR-150-5p | Npy1r | 18166 | ENSMUSG00000036437 | HITS-CLIP | Functional MTI (Weak) | 21258322 | validated |
| mirtarbase | MIMAT0000160 | mmu-miR-150-5p | Ogdh | 18293 | ENSMUSG00000020456 | HITS-CLIP | Functional MTI (Weak) | 21258322 | validated |
| mirtarbase | MIMAT0000160 | mmu-miR-150-5p | Pdgfb | 18591 | ENSMUSG00000000489 | Western blot//Reporter assay//Luciferase reporter assay | Functional MTI | 18500251 | validated |

| | | | | | | | | | |
|---|---|---|---|---|---|---|---|---|---|
| mirtarbase | MIMAT0000160 | mmu-miR-150-5p | Pik3c2a | 18704 | ENSMUSG00000030660 | HITS-CLIP | Functional MTI (Weak) | 21258322 | validated |
| mirtarbase | MIMAT0000160 | mmu-miR-150-5p | Pkd2 | 18764 | ENSMUSG00000034462 | HITS-CLIP | Functional MTI (Weak) | 21258322 | validated |
| mirtarbase | MIMAT0000160 | mmu-miR-150-5p | Pla2g2d | 18782 | ENSMUSG00000041202 | HITS-CLIP | Functional MTI (Weak) | 21258322 | validated |
| mirtarbase | MIMAT0000160 | mmu-miR-150-5p | Ppargc1a | 19017 | ENSMUSG00000029167 | Immunoblot//Immunohistochemistry//Immunoprecipitaion//Luciferase reporter assay//Microarray//qRT-PCR | Functional MTI | 24722250 | validated |
| mirtarbase | MIMAT0000160 | mmu-miR-150-5p | Ppp2r3d | 19054 | | HITS-CLIP | Functional MTI (Weak) | 21258322 | validated |
| mirtarbase | MIMAT0000160 | mmu-miR-150-5p | Mob4 | 19070 | ENSMUSG00000025979 | HITS-CLIP | Functional MTI (Weak) | 21258322 | validated |
| mirtarbase | MIMAT0000160 | mmu-miR-150-5p | Rhag | 19743 | ENSMUSG00000023926 | HITS-CLIP | Functional MTI (Weak) | 21258322 | validated |
| mirtarbase | MIMAT0000160 | mmu-miR-150-5p | Slc6a2 | 20538 | ENSMUSG00000055368 | HITS-CLIP | Functional MTI (Weak) | 21258322 | validated |
| mirtarbase | MIMAT0000160 | mmu-miR-150-5p | Srf | 20807 | ENSMUSG00000015605 | qRT-PCR//Western blot | Functional MTI | 25639779 | validated |
| mirtarbase | MIMAT0000160 | mmu-miR-150-5p | Stxbp4 | 20913 | ENSMUSG00000020546 | HITS-CLIP | Functional MTI (Weak) | 21258322 | validated |
| mirtarbase | MIMAT0000160 | mmu-miR-150-5p | Cntn2 | 21367 | ENSMUSG00000053024 | HITS-CLIP | Functional MTI (Weak) | 21258322 | validated |
| mirtarbase | MIMAT0000160 | mmu-miR-150-5p | Vegfa | 22339 | ENSMUSG00000023951 | Western blot//Reporter assay//Luciferase reporter assay | Functional MTI | 18500251 | validated |
| mirtarbase | MIMAT0000160 | mmu-miR-150-5p | Vegfa | 22339 | ENSMUSG00000023951 | Immunoblot//qRT-PCR | Functional MTI (Weak) | 23362149 | validated |
| mirtarbase | MIMAT0000160 | mmu-miR-150-5p | Vegfa | 22339 | ENSMUSG00000023951 | Luciferase reporter assay//qRT-PCR//Western blot | Functional MTI | 27391072 | validated |
| mirtarbase | MIMAT0000160 | mmu-miR-150-5p | Map3k12 | 26404 | ENSMUSG00000023050 | HITS-CLIP | Functional MTI (Weak) | 21258322 | validated |
| mirtarbase | MIMAT0000160 | mmu-miR-150-5p | Ercc4 | 50505 | | HITS-CLIP | Functional MTI (Weak) | 21258322 | validated |
| mirtarbase | MIMAT0000160 | mmu-miR-150-5p | Fbxl3 | 50789 | ENSMUSG00000022124 | HITS-CLIP | Functional MTI (Weak) | 21258322 | validated |
| mirtarbase | MIMAT0000160 | mmu-miR-150-5p | Chek2 | 50883 | ENSMUSG00000029521 | HITS-CLIP | Functional MTI (Weak) | 25083871 | validated |
| mirtarbase | MIMAT0000160 | mmu-miR-150-5p | Chek2 | 50883 | ENSMUSG00000029521 | HITS-CLIP | Functional MTI (Weak) | 21258322 | validated |
| mirtarbase | MIMAT0000160 | mmu-miR-150-5p | Tmem38b | 52076 | ENSMUSG00000028420 | HITS-CLIP | Functional MTI (Weak) | 26203562 | validated |
| mirtarbase | MIMAT0000160 | mmu-miR-150-5p | Tmem38b | 52076 | ENSMUSG00000028420 | HITS-CLIP | Functional MTI (Weak) | 23597149 | validated |
| mirtarbase | MIMAT0000160 | mmu-miR-150-5p | Tmem38b | 52076 | ENSMUSG00000028420 | HITS-CLIP | Functional MTI (Weak) | 21258322 | validated |

| | | | | | | | | | |
|---|---|---|---|---|---|---|---|---|---|
| mirtarbase | MIMAT0000160 | mmu-miR-150-5p | Kcnk6 | 52150 | ENSMUSG00000046410 | HITS-CLIP | Functional MTI (Weak) | 21258322 | validated |
| mirtarbase | MIMAT0000160 | mmu-miR-150-5p | Tet1 | 52463 | ENSMUSG00000047146 | HITS-CLIP | Functional MTI (Weak) | 21258322 | validated |
| mirtarbase | MIMAT0000160 | mmu-miR-150-5p | Angel2 | 52477 | ENSMUSG00000026634 | HITS-CLIP | Functional MTI (Weak) | 21258322 | validated |
| mirtarbase | MIMAT0000160 | mmu-miR-150-5p | Diaph2 | 54004 | ENSMUSG00000034480 | HITS-CLIP | Functional MTI (Weak) | 21258322 | validated |
| mirtarbase | MIMAT0000160 | mmu-miR-150-5p | Cxcl11 | 56066 | | HITS-CLIP | Functional MTI (Weak) | 21258322 | validated |
| mirtarbase | MIMAT0000160 | mmu-miR-150-5p | Car5b | 56078 | ENSMUSG00000031373 | HITS-CLIP | Functional MTI (Weak) | 21258322 | validated |
| mirtarbase | MIMAT0000160 | mmu-miR-150-5p | Mrpl19 | 56284 | ENSMUSG00000030045 | HITS-CLIP | Functional MTI (Weak) | 21258322 | validated |
| mirtarbase | MIMAT0000160 | mmu-miR-150-5p | Arhgap23 | 58996 | ENSMUSG00000049807 | HITS-CLIP | Functional MTI (Weak) | 21258322 | validated |
| mirtarbase | MIMAT0000160 | mmu-miR-150-5p | Ing5 | 66262 | ENSMUSG00000026283 | HITS-CLIP | Functional MTI (Weak) | 25083871 | validated |
| mirtarbase | MIMAT0000160 | mmu-miR-150-5p | Sorcs3 | 66673 | ENSMUSG00000063434 | HITS-CLIP | Functional MTI (Weak) | 23142080 | validated |
| mirtarbase | MIMAT0000160 | mmu-miR-150-5p | Magt1 | 67075 | ENSMUSG00000031232 | HITS-CLIP | Functional MTI (Weak) | 21258322 | validated |
| mirtarbase | MIMAT0000160 | mmu-miR-150-5p | Lrp2bp | 67620 | ENSMUSG00000031637 | HITS-CLIP | Functional MTI (Weak) | 21258322 | validated |
| mirtarbase | MIMAT0000160 | mmu-miR-150-5p | Rgs8 | 67792 | ENSMUSG00000042671 | Immunoblot//qRT-PCR | Functional MTI (Weak) | 23362149 | validated |
| mirtarbase | MIMAT0000160 | mmu-miR-150-5p | Slc25a39 | 68066 | ENSMUSG00000018677 | HITS-CLIP | Functional MTI (Weak) | 21258322 | validated |
| mirtarbase | MIMAT0000160 | mmu-miR-150-5p | Tprkb | 69786 | ENSMUSG00000054226 | HITS-CLIP | Functional MTI (Weak) | 25083871 | validated |
| mirtarbase | MIMAT0000160 | mmu-miR-150-5p | Tprkb | 69786 | ENSMUSG00000054226 | HITS-CLIP | Functional MTI (Weak) | 21258322 | validated |
| mirtarbase | MIMAT0000160 | mmu-miR-150-5p | Prdm16 | 70673 | ENSMUSG00000039410 | Immunoblot//Immunohistochemistry//Immunoprecipitaion//Luciferase reporter assay//Microarray//qRT-PCR | Functional MTI | 24722250 | validated |
| mirtarbase | MIMAT0000160 | mmu-miR-150-5p | Pwwp2a | 70802 | ENSMUSG00000044950 | HITS-CLIP | Functional MTI (Weak) | 21258322 | validated |
| mirtarbase | MIMAT0000160 | mmu-miR-150-5p | Grap | 71520 | ENSMUSG00000004837 | HITS-CLIP | Functional MTI (Weak) | 25083871 | validated |
| mirtarbase | MIMAT0000160 | mmu-miR-150-5p | Grap | 71520 | ENSMUSG00000004837 | HITS-CLIP | Functional MTI (Weak) | 23597149 | validated |
| mirtarbase | MIMAT0000160 | mmu-miR-150-5p | Grap | 71520 | ENSMUSG00000004837 | HITS-CLIP | Functional MTI (Weak) | 21258322 | validated |
| mirtarbase | MIMAT0000160 | mmu-miR-150-5p | Mau2 | 74549 | ENSMUSG00000031858 | HITS-CLIP | Functional MTI (Weak) | 21258322 | validated |

| | | | | | | | | | |
|---|---|---|---|---|---|---|---|---|---|
| mirtarbase | MIMAT0000160 | mmu-miR-150-5p | Asxl2 | 75302 | ENSMUSG00000037486 | HITS-CLIP | Functional MTI (Weak) | 21258322 | validated |
| mirtarbase | MIMAT0000160 | mmu-miR-150-5p | Mpp7 | 75739 | ENSMUSG00000057440 | HITS-CLIP | Functional MTI (Weak) | 21258322 | validated |
| mirtarbase | MIMAT0000160 | mmu-miR-150-5p | Bicd2 | 76895 | ENSMUSG00000037933 | HITS-CLIP | Functional MTI (Weak) | 23142080 | validated |
| mirtarbase | MIMAT0000160 | mmu-miR-150-5p | Usp45 | 77593 | ENSMUSG00000040455 | HITS-CLIP | Functional MTI (Weak) | 21258322 | validated |
| mirtarbase | MIMAT0000160 | mmu-miR-150-5p | H2az2 | 77605 | ENSMUSG00000041126 | HITS-CLIP | Functional MTI (Weak) | 21258322 | validated |
| mirtarbase | MIMAT0000160 | mmu-miR-150-5p | Rab27b | 80718 | ENSMUSG00000024511 | HITS-CLIP | Functional MTI (Weak) | 21258322 | validated |
| mirtarbase | MIMAT0000160 | mmu-miR-150-5p | Kcnn1 | 84036 | ENSMUSG00000002908 | HITS-CLIP | Functional MTI (Weak) | 21258322 | validated |
| mirtarbase | MIMAT0000160 | mmu-miR-150-5p | Kcnn1 | 84036 | ENSMUSG00000111706 | HITS-CLIP | Functional MTI (Weak) | 21258322 | validated |
| mirtarbase | MIMAT0000160 | mmu-miR-150-5p | Rabif | 98710 | | HITS-CLIP | Functional MTI (Weak) | 21258322 | validated |
| mirtarbase | MIMAT0000160 | mmu-miR-150-5p | Unc5b | 107449 | ENSMUSG00000020099 | HITS-CLIP | Functional MTI (Weak) | 26203562 | validated |
| mirtarbase | MIMAT0000160 | mmu-miR-150-5p | Unc5b | 107449 | ENSMUSG00000020099 | HITS-CLIP | Functional MTI (Weak) | 25083871 | validated |
| mirtarbase | MIMAT0000160 | mmu-miR-150-5p | Unc5b | 107449 | ENSMUSG00000020099 | HITS-CLIP | Functional MTI (Weak) | 23597149 | validated |
| mirtarbase | MIMAT0000160 | mmu-miR-150-5p | Unc5b | 107449 | ENSMUSG00000020099 | HITS-CLIP | Functional MTI (Weak) | 23142080 | validated |
| mirtarbase | MIMAT0000160 | mmu-miR-150-5p | Unc5b | 107449 | ENSMUSG00000020099 | HITS-CLIP | Functional MTI (Weak) | 21258322 | validated |
| mirtarbase | MIMAT0000160 | mmu-miR-150-5p | Unc5b | 107449 | ENSMUSG00000020099 | HITS-CLIP | Functional MTI (Weak) | 19536157 | validated |
| mirtarbase | MIMAT0000160 | mmu-miR-150-5p | Ssr1 | 107513 | ENSMUSG00000021427 | HITS-CLIP | Functional MTI (Weak) | 21258322 | validated |
| mirtarbase | MIMAT0000160 | mmu-miR-150-5p | Chrnb4 | 108015 | ENSMUSG00000035200 | HITS-CLIP | Functional MTI (Weak) | 21258322 | validated |
| mirtarbase | MIMAT0000160 | mmu-miR-150-5p | Acat1 | 110446 | ENSMUSG00000032047 | HITS-CLIP | Functional MTI (Weak) | 21258322 | validated |
| mirtarbase | MIMAT0000160 | mmu-miR-150-5p | Adarb1 | 110532 | ENSMUSG00000020262 | HITS-CLIP | Functional MTI (Weak) | 21258322 | validated |
| mirtarbase | MIMAT0000160 | mmu-miR-150-5p | Cds2 | 110911 | ENSMUSG00000058793 | HITS-CLIP | Functional MTI (Weak) | 25083871 | validated |
| mirtarbase | MIMAT0000160 | mmu-miR-150-5p | Cds2 | 110911 | ENSMUSG00000058793 | HITS-CLIP | Functional MTI (Weak) | 21258322 | validated |
| mirtarbase | MIMAT0000160 | mmu-miR-150-5p | Zfp607b | 112415 | ENSMUSG00000057093 | HITS-CLIP | Functional MTI (Weak) | 21258322 | validated |

| | | | | | | | | | |
|---|---|---|---|---|---|---|---|---|---|
| mirtarbase | MIMAT0000160 | mmu-miR-150-5p | Tmlhe | 192289 | ENSMUSG00000079834 | HITS-CLIP | Functional MTI (Weak) | 21258322 | validated |
| mirtarbase | MIMAT0000160 | mmu-miR-150-5p | Ripk2 | 192656 | ENSMUSG00000041135 | Immunoblot//Luciferase reporter assay//qRT-PCR | Functional MTI | 26391398 | validated |
| mirtarbase | MIMAT0000160 | mmu-miR-150-5p | Gabpb2 | 213054 | ENSMUSG00000038766 | HITS-CLIP | Functional MTI (Weak) | 21258322 | validated |
| mirtarbase | MIMAT0000160 | mmu-miR-150-5p | Nav1 | 215690 | ENSMUSG00000009418 | HITS-CLIP | Functional MTI (Weak) | 21258322 | validated |
| mirtarbase | MIMAT0000160 | mmu-miR-150-5p | Ybey | 216119 | ENSMUSG00000033126 | HITS-CLIP | Functional MTI (Weak) | 21258322 | validated |
| mirtarbase | MIMAT0000160 | mmu-miR-150-5p | Heatr6 | 217026 | ENSMUSG00000000976 | HITS-CLIP | Functional MTI (Weak) | 21258322 | validated |
| mirtarbase | MIMAT0000160 | mmu-miR-150-5p | Tns4 | 217169 | ENSMUSG00000017607 | HITS-CLIP | Functional MTI (Weak) | 21258322 | validated |
| mirtarbase | MIMAT0000160 | mmu-miR-150-5p | Cd300a | 217303 | ENSMUSG00000034652 | HITS-CLIP | Functional MTI (Weak) | 21258322 | validated |
| mirtarbase | MIMAT0000160 | mmu-miR-150-5p | Nol10 | 217431 | ENSMUSG00000061458 | HITS-CLIP | Functional MTI (Weak) | 21258322 | validated |
| mirtarbase | MIMAT0000160 | mmu-miR-150-5p | Mlh3 | 217716 | ENSMUSG00000021245 | HITS-CLIP | Functional MTI (Weak) | 21258322 | validated |
| mirtarbase | MIMAT0000160 | mmu-miR-150-5p | Cdc14b | 218294 | ENSMUSG00000033102 | HITS-CLIP | Functional MTI (Weak) | 21258322 | validated |
| mirtarbase | MIMAT0000160 | mmu-miR-150-5p | Rprd1a | 225283 | ENSMUSG00000040446 | HITS-CLIP | Functional MTI (Weak) | 21258322 | validated |
| mirtarbase | MIMAT0000160 | mmu-miR-150-5p | Pbxip1 | 229534 | ENSMUSG00000042613 | HITS-CLIP | Functional MTI (Weak) | 21258322 | validated |
| mirtarbase | MIMAT0000160 | mmu-miR-150-5p | Sdad1 | 231452 | ENSMUSG00000029415 | HITS-CLIP | Functional MTI (Weak) | 21258322 | validated |
| mirtarbase | MIMAT0000160 | mmu-miR-150-5p | Glyctk | 235582 | ENSMUSG00000020258 | HITS-CLIP | Functional MTI (Weak) | 21258322 | validated |
| mirtarbase | MIMAT0000160 | mmu-miR-150-5p | Zfp811 | 240063 | ENSMUSG00000055202 | HITS-CLIP | Functional MTI (Weak) | 21258322 | validated |
| mirtarbase | MIMAT0000160 | mmu-miR-150-5p | Tmem245 | 242474 | ENSMUSG00000055296 | HITS-CLIP | Functional MTI (Weak) | 21258322 | validated |
| mirtarbase | MIMAT0000160 | mmu-miR-150-5p | Zfp568 | 243905 | ENSMUSG00000074221 | HITS-CLIP | Functional MTI (Weak) | 25083871 | validated |
| mirtarbase | MIMAT0000160 | mmu-miR-150-5p | Dkc1 | 245474 | ENSMUSG00000031403 | HITS-CLIP | Functional MTI (Weak) | 21258322 | validated |
| mirtarbase | MIMAT0000160 | mmu-miR-150-5p | AY07488 | 246735 | | HITS-CLIP | Functional MTI (Weak) | 21258322 | validated |
| mirtarbase | MIMAT0000160 | mmu-miR-150-5p | Depdc5 | 277854 | ENSMUSG00000037426 | HITS-CLIP | Functional MTI (Weak) | 21258322 | validated |
| mirtarbase | MIMAT0000160 | mmu-miR-150-5p | Zfyve27 | 319740 | ENSMUSG00000018820 | HITS-CLIP | Functional MTI | 21258322 | validated |
| mirtarbase | MIMAT0000160 | mmu-miR-150-5p | A830018L | 320492 | ENSMUSG00000057715 | HITS-CLIP | Functional MTI | 21258322 | validated |
| mirtarbase | MIMAT0000160 | mmu-miR-150-5p | Gm14325 | 329575 | | HITS-CLIP | Functional MTI | 21258322 | validated |

| | | | | | | | | | |
|---|---|---|---|---|---|---|---|---|---|
| mirtarbase | MIMAT0000160 | mmu-miR-150-5p | Zfp866 | 330788 | | HITS-CLIP | Functional MTI (Weak) | 21258322 | validated |
| mirtarbase | MIMAT0000160 | mmu-miR-150-5p | Dixdc1 | 330938 | ENSMUSG00000032064 | HITS-CLIP | Functional MTI (Weak) | 21258322 | validated |
| mirtarbase | MIMAT0000160 | mmu-miR-150-5p | Zfp882 | 382019 | | HITS-CLIP | Functional MTI (Weak) | 21258322 | validated |
| mirtarbase | MIMAT0000160 | mmu-miR-150-5p | Zfp488 | 382867 | ENSMUSG00000044519 | HITS-CLIP | Functional MTI (Weak) | 21258322 | validated |
| mirtarbase | MIMAT0000160 | mmu-miR-150-5p | Mettl21e | 403183 | ENSMUSG00000046828 | HITS-CLIP | Functional MTI (Weak) | 21258322 | validated |
| mirtarbase | MIMAT0000160 | mmu-miR-150-5p | Gm14430 | 627914 | | HITS-CLIP | Functional MTI (Weak) | 21258322 | validated |
| mirtarbase | MIMAT0000160 | mmu-miR-150-5p | Zfp970 | 628308 | | HITS-CLIP | Functional MTI (Weak) | 21258322 | validated |
| mirtarbase | MIMAT0000160 | mmu-miR-150-5p | Gm14326 | 665211 | ENSMUSG00000078862 | HITS-CLIP | Functional MTI (Weak) | 21258322 | validated |
| mirtarbase | MIMAT0000160 | mmu-miR-150-5p | Gm4631 | 668039 | ENSMUSG00000078899 | HITS-CLIP | Functional MTI (Weak) | 21258322 | validated |
| mirtarbase | MIMAT0000160 | mmu-miR-150-5p | Gm4631 | 1E+08 | ENSMUSG00000078899 | HITS-CLIP | Functional MTI (Weak) | 21258322 | validated |
| tarbase | MIMAT0000160 | mmu-miR-150-5p | Mef2d | 17261 | ENSMUSG00000001419 | Degradome sequencing | positive | | validated |
| tarbase | MIMAT0000160 | mmu-miR-150-5p | Kmt2a | 214162 | ENSMUSG00000002028 | Degradome sequencing | positive | | validated |
| tarbase | MIMAT0000160 | mmu-miR-150-5p | Stat6 | 20852 | ENSMUSG00000002147 | Degradome sequencing | positive | | validated |
| tarbase | MIMAT0000160 | mmu-miR-150-5p | Cnot11 | 52846 | ENSMUSG00000003135 | Degradome sequencing | positive | | validated |
| tarbase | MIMAT0000160 | mmu-miR-150-5p | Slc2a3 | 20527 | ENSMUSG00000003153 | Degradome sequencing//Degradome sequencing | positive | | validated |
| tarbase | MIMAT0000160 | mmu-miR-150-5p | Mmd | 67468 | ENSMUSG00000003948 | Degradome sequencing | positive | | validated |
| tarbase | MIMAT0000160 | mmu-miR-150-5p | Hdgf | 15191 | ENSMUSG00000004897 | Degradome sequencing | positive | | validated |
| tarbase | MIMAT0000160 | mmu-miR-150-5p | Metap1 | 75624 | ENSMUSG00000005813 | Degradome sequencing | positive | | validated |
| tarbase | MIMAT0000160 | mmu-miR-150-5p | Apc | 11789 | ENSMUSG00000005871 | Degradome sequencing//Degradome sequencing | positive | | validated |
| tarbase | MIMAT0000160 | mmu-miR-150-5p | Pdk1 | 228026 | ENSMUSG00000006494 | Degradome sequencing//Degradome sequencing | positive | | validated |
| tarbase | MIMAT0000160 | mmu-miR-150-5p | Mark3 | 17169 | ENSMUSG00000007411 | Degradome sequencing | positive | | validated |
| tarbase | MIMAT0000160 | mmu-miR-150-5p | Tgfbr1 | 21812 | ENSMUSG00000007613 | Degradome sequencing | positive | | validated |
| tarbase | MIMAT0000160 | mmu-miR-150-5p | Snrpb2 | | ENSMUSG00000008333 | Degradome sequencing | positive | | validated |
| tarbase | MIMAT0000160 | mmu-miR-150-5p | Cbx5 | 12419 | ENSMUSG00000009575 | Degradome sequencing | positive | | validated |
| tarbase | MIMAT0000160 | mmu-miR-150-5p | Ptprs | 19280 | ENSMUSG00000013236 | Degradome sequencing | positive | | validated |
| tarbase | MIMAT0000160 | mmu-miR-150-5p | Rnf168 | 70238 | ENSMUSG00000014074 | Degradome sequencing | positive | | validated |
| tarbase | MIMAT0000160 | mmu-miR-150-5p | Anapc1 | 17222 | ENSMUSG00000014355 | Degradome sequencing//Degradome sequencing | positive | | validated |
| tarbase | MIMAT0000160 | mmu-miR-150-5p | Capza2 | 12343 | ENSMUSG00000015733 | Degradome sequencing | positive | | validated |
| tarbase | MIMAT0000160 | mmu-miR-150-5p | Nfe2l2 | 18024 | ENSMUSG00000015839 | Degradome sequencing | positive | | validated |
| tarbase | MIMAT0000160 | mmu-miR-150-5p | Cd274 | 60533 | ENSMUSG00000016496 | Degradome sequencing | positive | | validated |

| | | | | | | | | | |
|---|---|---|---|---|---|---|---|---|---|
| tarbase | MIMAT0000160 | mmu-miR-150-5p | Zfp207 | 22680 | ENSMUSG00000017421 | Degradome sequencing//Degradome sequencing | positive | | validated |
| tarbase | MIMAT0000160 | mmu-miR-150-5p | Top2b | 21974 | ENSMUSG00000017485 | Degradome sequencing | positive | | validated |
| tarbase | MIMAT0000160 | mmu-miR-150-5p | Mybl2 | 17865 | ENSMUSG00000017861 | Degradome sequencing | positive | | validated |
| tarbase | MIMAT0000160 | mmu-miR-150-5p | Srsf1 | 110809 | ENSMUSG00000018379 | Degradome sequencing//Degradome sequencing//Degradome sequencing | positive | | validated |
| tarbase | MIMAT0000160 | mmu-miR-150-5p | Ikzf1 | 22778 | ENSMUSG00000018654 | Degradome sequencing | positive | | validated |
| tarbase | MIMAT0000160 | mmu-miR-150-5p | Ywhah | 22629 | ENSMUSG00000018965 | Degradome sequencing | positive | | validated |
| tarbase | MIMAT0000160 | mmu-miR-150-5p | Zfp687 | 78266 | ENSMUSG00000019338 | Degradome sequencing//Degradome sequencing | positive | | validated |
| tarbase | MIMAT0000160 | mmu-miR-150-5p | Utrn | 22288 | ENSMUSG00000019820 | Degradome sequencing | positive | | validated |
| tarbase | MIMAT0000160 | mmu-miR-150-5p | Prep | | ENSMUSG00000019849 | Degradome sequencing | positive | | validated |
| tarbase | MIMAT0000160 | mmu-miR-150-5p | Atp2b1 | 67972 | ENSMUSG00000019943 | Degradome sequencing | positive | | validated |
| tarbase | MIMAT0000160 | mmu-miR-150-5p | Myb | 17863 | ENSMUSG00000019982 | Degradome sequencing//Degradome sequencing | positive | | validated |
| tarbase | MIMAT0000160 | mmu-miR-150-5p | Ccar1 | 67500 | ENSMUSG00000020074 | Degradome sequencing | positive | | validated |
| tarbase | MIMAT0000160 | mmu-miR-150-5p | Vps54 | 245944 | ENSMUSG00000020128 | Degradome sequencing//Degradome sequencing | positive | | validated |
| tarbase | MIMAT0000160 | mmu-miR-150-5p | Ahsa2 | 268390 | ENSMUSG00000020288 | Degradome sequencing | positive | | validated |
| tarbase | MIMAT0000160 | mmu-miR-150-5p | Hnrnpab | 15384 | ENSMUSG00000020358 | Degradome sequencing | positive | | validated |
| tarbase | MIMAT0000160 | mmu-miR-150-5p | Patz1 | 56218 | ENSMUSG00000020453 | Degradome sequencing | positive | | validated |
| tarbase | MIMAT0000160 | mmu-miR-150-5p | Pum2 | 80913 | ENSMUSG00000020594 | Degradome sequencing | positive | | validated |
| tarbase | MIMAT0000160 | mmu-miR-150-5p | Prkar1a | 19084 | ENSMUSG00000020612 | Degradome sequencing//Degradome sequencing | positive | | validated |
| tarbase | MIMAT0000160 | mmu-miR-150-5p | Tlk2 | 24086 | ENSMUSG00000020694 | Degradome sequencing | positive | | validated |
| tarbase | MIMAT0000160 | mmu-miR-150-5p | Cluh | 74148 | ENSMUSG00000020741 | Degradome sequencing | positive | | validated |
| tarbase | MIMAT0000160 | mmu-miR-150-5p | Blmh | 104184 | ENSMUSG00000020840 | Degradome sequencing//Degradome sequencing | positive | | validated |
| tarbase | MIMAT0000160 | mmu-miR-150-5p | Ywhae | 22627 | ENSMUSG00000020849 | Degradome sequencing//Degradome sequencing | positive | | validated |
| tarbase | MIMAT0000160 | mmu-miR-150-5p | Prpf8 | 192159 | ENSMUSG00000020850 | Degradome sequencing//Degradome sequencing | positive | | validated |
| tarbase | MIMAT0000160 | mmu-miR-150-5p | Top2a | 21973 | ENSMUSG00000020914 | Degradome sequencing | positive | | validated |
| tarbase | MIMAT0000160 | mmu-miR-150-5p | Stat5b | 20851 | ENSMUSG00000020919 | Degradome sequencing | positive | | validated |
| tarbase | MIMAT0000160 | mmu-miR-150-5p | Zfyve21 | 68520 | ENSMUSG00000021286 | Degradome sequencing//Degradome sequencing | positive | | validated |
| tarbase | MIMAT0000160 | mmu-miR-150-5p | Irf4 | 16364 | ENSMUSG00000021356 | Degradome sequencing//Degradome sequencing | positive | | validated |
| tarbase | MIMAT0000160 | mmu-miR-150-5p | Nsd1 | 18193 | ENSMUSG00000021488 | Degradome sequencing | positive | | validated |
| tarbase | MIMAT0000160 | mmu-miR-150-5p | Kat6b | 54169 | ENSMUSG00000021767 | Degradome sequencing | positive | | validated |
| tarbase | MIMAT0000160 | mmu-miR-150-5p | Arhgef3 | 71704 | ENSMUSG00000021895 | Degradome sequencing//Degradome sequencing | positive | | validated |
| tarbase | MIMAT0000160 | mmu-miR-150-5p | Rnf19a | 30945 | ENSMUSG00000022280 | Degradome sequencing//Degradome sequencing | positive | | validated |

| | | | | | | | | | |
|---|---|---|---|---|---|---|---|---|---|
| tarbase | MIMAT0000160 | mmu-miR-150-5p | Twf1 | 19230 | ENSMUSG00000022451 | Degradome sequencing//Degradome sequencing | positive | | validated |
| tarbase | MIMAT0000160 | mmu-miR-150-5p | Nde1 | 67203 | ENSMUSG00000022678 | Degradome sequencing | positive | | validated |
| tarbase | MIMAT0000160 | mmu-miR-150-5p | Tfrc | 22042 | ENSMUSG00000022797 | Degradome sequencing | positive | | validated |
| tarbase | MIMAT0000160 | mmu-miR-150-5p | Iqcb1 | 320299 | ENSMUSG00000022837 | Degradome sequencing | positive | | validated |
| tarbase | MIMAT0000160 | mmu-miR-150-5p | Brwd1 | 93871 | ENSMUSG00000022914 | Degradome sequencing | positive | | validated |
| tarbase | MIMAT0000160 | mmu-miR-150-5p | Map3k12 | 26404 | ENSMUSG00000023050 | Degradome sequencing | positive | | validated |
| tarbase | MIMAT0000160 | mmu-miR-150-5p | Pim1 | 18712 | ENSMUSG00000024014 | Degradome sequencing//Degradome sequencing | positive | | validated |
| tarbase | MIMAT0000160 | mmu-miR-150-5p | Lbh | 77889 | ENSMUSG00000024063 | Degradome sequencing | positive | | validated |
| tarbase | MIMAT0000160 | mmu-miR-150-5p | Birc6 | 12211 | ENSMUSG00000024073 | Degradome sequencing | positive | | validated |
| tarbase | MIMAT0000160 | mmu-miR-150-5p | Etf1 | 225363 | ENSMUSG00000024360 | Degradome sequencing//Degradome sequencing | positive | | validated |
| tarbase | MIMAT0000160 | mmu-miR-150-5p | Gabbr1 | 54393 | ENSMUSG00000024462 | Degradome sequencing | positive | | validated |
| tarbase | MIMAT0000160 | mmu-miR-150-5p | Dcp2 | 70640 | ENSMUSG00000024472 | Degradome sequencing | positive | | validated |
| tarbase | MIMAT0000160 | mmu-miR-150-5p | Tcerg1 | 56070 | ENSMUSG00000024498 | Degradome sequencing | positive | | validated |
| tarbase | MIMAT0000160 | mmu-miR-150-5p | Csnk1a1 | 93687 | ENSMUSG00000024576 | Degradome sequencing//Degradome sequencing | positive | | validated |
| tarbase | MIMAT0000160 | mmu-miR-150-5p | Nars | 70223 | ENSMUSG00000024587 | Degradome sequencing | positive | | validated |
| tarbase | MIMAT0000160 | mmu-miR-150-5p | Zfp91 | 109910 | ENSMUSG00000024695 | Degradome sequencing | positive | | validated |
| tarbase | MIMAT0000160 | mmu-miR-150-5p | Cdc37l1 | 67072 | ENSMUSG00000024780 | Degradome sequencing | positive | | validated |
| tarbase | MIMAT0000160 | mmu-miR-150-5p | Tnks2 | 74493 | ENSMUSG00000024811 | Degradome sequencing//Degradome sequencing | positive | | validated |
| tarbase | MIMAT0000160 | mmu-miR-150-5p | Vldlr | 22359 | ENSMUSG00000024924 | Degradome sequencing//Degradome sequencing | positive | | validated |
| tarbase | MIMAT0000160 | mmu-miR-150-5p | Lcor | 212391 | ENSMUSG00000025019 | Degradome sequencing//Degradome sequencing | positive | | validated |
| tarbase | MIMAT0000160 | mmu-miR-150-5p | Add3 | 27360 | ENSMUSG00000025026 | Degradome sequencing | positive | | validated |
| tarbase | MIMAT0000160 | mmu-miR-150-5p | Scd2 | 20250 | ENSMUSG00000025203 | Degradome sequencing//Degradome sequencing | positive | | validated |
| tarbase | MIMAT0000160 | mmu-miR-150-5p | Pikfyve | 18711 | ENSMUSG00000025949 | Degradome sequencing | positive | | validated |
| tarbase | MIMAT0000160 | mmu-miR-150-5p | Cd28 | 12487 | ENSMUSG00000026012 | Degradome sequencing//Degradome sequencing | positive | | validated |
| tarbase | MIMAT0000160 | mmu-miR-150-5p | Acbd3 | 170760 | ENSMUSG00000026499 | Degradome sequencing | positive | | validated |
| tarbase | MIMAT0000160 | mmu-miR-150-5p | Dcaf6 | 74106 | ENSMUSG00000026571 | Degradome sequencing | positive | | validated |
| tarbase | MIMAT0000160 | mmu-miR-150-5p | Abl2 | 11352 | ENSMUSG00000026596 | Degradome sequencing | positive | | validated |
| tarbase | MIMAT0000160 | mmu-miR-150-5p | Il2ra | 16184 | ENSMUSG00000026770 | Degradome sequencing | positive | | validated |
| tarbase | MIMAT0000160 | mmu-miR-150-5p | Hspa5 | 14828 | ENSMUSG00000026864 | Degradome sequencing//Degradome sequencing | positive | | validated |
| tarbase | MIMAT0000160 | mmu-miR-150-5p | Hipk3 | 15259 | ENSMUSG00000027177 | Degradome sequencing | positive | | validated |
| tarbase | MIMAT0000160 | mmu-miR-150-5p | Fbxo3 | 57443 | ENSMUSG00000027180 | Degradome sequencing | positive | | validated |
| tarbase | MIMAT0000160 | mmu-miR-150-5p | Caprin1 | 53872 | ENSMUSG00000027184 | Degradome sequencing | positive | | validated |
| tarbase | MIMAT0000160 | mmu-miR-150-5p | Pdia3 | 14827 | ENSMUSG00000027248 | Degradome sequencing//Degradome sequencing//Degradome sequencing | positive | | validated |
| tarbase | MIMAT0000160 | mmu-miR-150-5p | Snx5 | 69178 | ENSMUSG00000027423 | Degradome sequencing | positive | | validated |
| tarbase | MIMAT0000160 | mmu-miR-150-5p | Gnas | 14683 | ENSMUSG00000027523 | Degradome sequencing | positive | | validated |

| | | | | | | | | | |
|---|---|---|---|---|---|---|---|---|---|
| tarbase | MIMAT0000160 | mmu-miR-150-5p | Mbnl1 | 56758 | ENSMUSG00000027763 | Degradome sequencing | positive | | validated |
| tarbase | MIMAT0000160 | mmu-miR-150-5p | Tpm3 | 59069 | ENSMUSG00000027940 | Degradome sequencing | positive | | validated |
| tarbase | MIMAT0000160 | mmu-miR-150-5p | Hadh | 15107 | ENSMUSG00000027984 | Degradome sequencing | positive | | validated |
| tarbase | MIMAT0000160 | mmu-miR-150-5p | Ccne2 | 12448 | ENSMUSG00000028212 | Degradome sequencing//Degradome sequencing | positive | | validated |
| tarbase | MIMAT0000160 | mmu-miR-150-5p | Tmeff1 | 230157 | ENSMUSG00000028347 | Degradome sequencing | positive | | validated |
| tarbase | MIMAT0000160 | mmu-miR-150-5p | Zdhhc21 | 68268 | ENSMUSG00000028403 | Degradome sequencing | positive | | validated |
| tarbase | MIMAT0000160 | mmu-miR-150-5p | Rad23b | 19359 | ENSMUSG00000028426 | Degradome sequencing | positive | | validated |
| tarbase | MIMAT0000160 | mmu-miR-150-5p | Macf1 | 11426 | ENSMUSG00000028649 | Degradome sequencing | positive | | validated |
| tarbase | MIMAT0000160 | mmu-miR-150-5p | Nasp | 50927 | ENSMUSG00000028693 | Degradome sequencing | positive | | validated |
| tarbase | MIMAT0000160 | mmu-miR-150-5p | AU04032( | 100317 | ENSMUSG00000028830 | Degradome sequencing//Degradome sequencing | positive | | validated |
| tarbase | MIMAT0000160 | mmu-miR-150-5p | Mtor | 56717 | ENSMUSG00000028991 | Degradome sequencing | negative | | validated |
| tarbase | MIMAT0000160 | mmu-miR-150-5p | Rnf4 | 19822 | ENSMUSG00000029110 | Degradome sequencing//Degradome sequencing | positive | | validated |
| tarbase | MIMAT0000160 | mmu-miR-150-5p | Gfi1 | 14581 | ENSMUSG00000029275 | Degradome sequencing | positive | | validated |
| tarbase | MIMAT0000160 | mmu-miR-150-5p | Ccng2 | | ENSMUSG00000029385 | Degradome sequencing//Degradome sequencing | positive | | validated |
| tarbase | MIMAT0000160 | mmu-miR-150-5p | Abcb9 | 56325 | ENSMUSG00000029408 | Degradome sequencing//Degradome sequencing | positive | | validated |
| tarbase | MIMAT0000160 | mmu-miR-150-5p | Sppl3 | 74585 | ENSMUSG00000029550 | Degradome sequencing | positive | | validated |
| tarbase | MIMAT0000160 | mmu-miR-150-5p | Exoc4 | 20336 | ENSMUSG00000029763 | Degradome sequencing | positive | | validated |
| tarbase | MIMAT0000160 | mmu-miR-150-5p | Mtpn | 14489 | ENSMUSG00000029840 | Degradome sequencing | positive | | validated |
| tarbase | MIMAT0000160 | mmu-miR-150-5p | Mkrn1 | 54484 | ENSMUSG00000029922 | Degradome sequencing | positive | | validated |
| tarbase | MIMAT0000160 | mmu-miR-150-5p | Isy1 | | ENSMUSG00000030056 | Degradome sequencing | positive | | validated |
| tarbase | MIMAT0000160 | mmu-miR-150-5p | Foxp1 | 108655 | ENSMUSG00000030067 | Degradome sequencing | positive | | validated |
| tarbase | MIMAT0000160 | mmu-miR-150-5p | Bhlhe40 | 20893 | ENSMUSG00000030103 | Degradome sequencing//Degradome sequencing | positive | | validated |
| tarbase | MIMAT0000160 | mmu-miR-150-5p | Adipor2 | 68465 | ENSMUSG00000030168 | Degradome sequencing | positive | | validated |
| tarbase | MIMAT0000160 | mmu-miR-150-5p | Arhgdib | 11857 | ENSMUSG00000030220 | Degradome sequencing//Degradome sequencing | positive | | validated |
| tarbase | MIMAT0000160 | mmu-miR-150-5p | Sec13 | 110379 | ENSMUSG00000030298 | Degradome sequencing | positive | | validated |
| tarbase | MIMAT0000160 | mmu-miR-150-5p | Pagr1a | | ENSMUSG00000030680 | Degradome sequencing | positive | | validated |
| tarbase | MIMAT0000160 | mmu-miR-150-5p | Il21r | 60504 | ENSMUSG00000030745 | Degradome sequencing | positive | | validated |
| tarbase | MIMAT0000160 | mmu-miR-150-5p | Ash2l | 23808 | ENSMUSG00000031575 | Degradome sequencing | positive | | validated |
| tarbase | MIMAT0000160 | mmu-miR-150-5p | Tnpo2 | 212999 | ENSMUSG00000031691 | Degradome sequencing | positive | | validated |
| tarbase | MIMAT0000160 | mmu-miR-150-5p | Ccnb2 | 12442 | ENSMUSG00000032218 | Degradome sequencing | positive | | validated |
| tarbase | MIMAT0000160 | mmu-miR-150-5p | Anp32a | 11737 | ENSMUSG00000032249 | Degradome sequencing | positive | | validated |
| tarbase | MIMAT0000160 | mmu-miR-150-5p | Pkm | 18746 | ENSMUSG00000032294 | Degradome sequencing//Degradome sequencing | positive | | validated |
| tarbase | MIMAT0000160 | mmu-miR-150-5p | Ulk3 | 71742 | ENSMUSG00000032308 | Degradome sequencing | positive | | validated |
| tarbase | MIMAT0000160 | mmu-miR-150-5p | Pdcd6ip | 18571 | ENSMUSG00000032504 | Degradome sequencing//Degradome se( | positive | | validated |
| tarbase | MIMAT0000160 | mmu-miR-150-5p | Topbp1 | 235559 | ENSMUSG00000032555 | Degradome sequencing | positive | | validated |
| tarbase | MIMAT0000160 | mmu-miR-150-5p | Cish | 12700 | ENSMUSG00000032578 | Degradome sequencing//Degradome se( | positive | | validated |
| tarbase | MIMAT0000160 | mmu-miR-150-5p | Cip2a | 224171 | ENSMUSG00000033031 | Degradome sequencing | positive | | validated |
| tarbase | MIMAT0000160 | mmu-miR-150-5p | Atg9a | 245860 | ENSMUSG00000033124 | Degradome sequencing | positive | | validated |

| | | | | | | | | | |
|---|---|---|---|---|---|---|---|---|---|
| tarbase | MIMAT0000160 | mmu-miR-150-5p | Szt2 | 230676 | ENSMUSG00000033253 | Degradome sequencing//Degradome se | positive | | validated |
| tarbase | MIMAT0000160 | mmu-miR-150-5p | Mif | 17319 | ENSMUSG00000033307 | Degradome sequencing | positive | | validated |
| tarbase | MIMAT0000160 | mmu-miR-150-5p | Fem1c | 240263 | ENSMUSG00000033319 | Degradome sequencing//Degradome sequencing | positive | | validated |
| tarbase | MIMAT0000160 | mmu-miR-150-5p | Mapkbp1 | 26390 | ENSMUSG00000033902 | Degradome sequencing | positive | | validated |
| tarbase | MIMAT0000160 | mmu-miR-150-5p | Mga | 29808 | ENSMUSG00000033943 | Degradome sequencing//Degradome sequencing | positive | | validated |
| tarbase | MIMAT0000160 | mmu-miR-150-5p | Setd5 | 72895 | ENSMUSG00000034269 | Degradome sequencing | positive | | validated |
| tarbase | MIMAT0000160 | mmu-miR-150-5p | Smc4 | 70099 | ENSMUSG00000034349 | Degradome sequencing | positive | | validated |
| tarbase | MIMAT0000160 | mmu-miR-150-5p | Zfp395 | 380912 | ENSMUSG00000034522 | Degradome sequencing | positive | | validated |
| tarbase | MIMAT0000160 | mmu-miR-150-5p | Bmp2k | 140780 | ENSMUSG00000034663 | Degradome sequencing | positive | | validated |
| tarbase | MIMAT0000160 | mmu-miR-150-5p | Tet3 | 194388 | ENSMUSG00000034832 | Degradome sequencing//Degradome sequencing | positive | | validated |
| tarbase | MIMAT0000160 | mmu-miR-150-5p | Ap2b1 | 71770 | ENSMUSG00000035152 | Degradome sequencing | positive | | validated |
| tarbase | MIMAT0000160 | mmu-miR-150-5p | Rabgap1 | 227800 | ENSMUSG00000035437 | Degradome sequencing | positive | | validated |
| tarbase | MIMAT0000160 | mmu-miR-150-5p | Rsf1 | 233532 | ENSMUSG00000035623 | Degradome sequencing | positive | | validated |
| tarbase | MIMAT0000160 | mmu-miR-150-5p | Ggta1 | 14594 | ENSMUSG00000035778 | Degradome sequencing | positive | | validated |
| tarbase | MIMAT0000160 | mmu-miR-150-5p | Pip4p1 | 219024 | ENSMUSG00000035953 | Degradome sequencing | positive | | validated |
| tarbase | MIMAT0000160 | mmu-miR-150-5p | Ube2r2 | 67615 | ENSMUSG00000036241 | Degradome sequencing | positive | | validated |
| tarbase | MIMAT0000160 | mmu-miR-150-5p | Oxsr1 | 108737 | ENSMUSG00000036737 | Degradome sequencing | positive | | validated |
| tarbase | MIMAT0000160 | mmu-miR-150-5p | Map11 | 231807 | ENSMUSG00000036948 | Degradome sequencing//Degradome sequencing | positive | | validated |
| tarbase | MIMAT0000160 | mmu-miR-150-5p | Slc12a9 | 83704 | ENSMUSG00000037344 | Degradome sequencing | positive | | validated |
| tarbase | MIMAT0000160 | mmu-miR-150-5p | Cd81 | 12520 | ENSMUSG00000037706 | Degradome sequencing | positive | | validated |
| tarbase | MIMAT0000160 | mmu-miR-150-5p | Nufip2 | 68564 | ENSMUSG00000037857 | Degradome sequencing//Degradome sequencing | positive | | validated |
| tarbase | MIMAT0000160 | mmu-miR-150-5p | Egr2 | 13654 | ENSMUSG00000037868 | Degradome sequencing//Degradome sequencing | positive | | validated |
| tarbase | MIMAT0000160 | mmu-miR-150-5p | Stk35 | 67333 | ENSMUSG00000037885 | Degradome sequencing | positive | | validated |
| tarbase | MIMAT0000160 | mmu-miR-150-5p | Bri3bp | 76809 | ENSMUSG00000037905 | Degradome sequencing | positive | | validated |
| tarbase | MIMAT0000160 | mmu-miR-150-5p | Rara | 19401 | ENSMUSG00000037992 | Degradome sequencing//Degradome sequencing | positive | | validated |
| tarbase | MIMAT0000160 | mmu-miR-150-5p | Ccdc50 | 67501 | ENSMUSG00000038127 | Degradome sequencing | positive | | validated |
| tarbase | MIMAT0000160 | mmu-miR-150-5p | Ttc39b | 69863 | ENSMUSG00000038172 | Degradome sequencing | positive | | validated |
| tarbase | MIMAT0000160 | mmu-miR-150-5p | Mllt6 | 246198 | ENSMUSG00000038437 | Degradome sequencing | positive | | validated |
| tarbase | MIMAT0000160 | mmu-miR-150-5p | Jarid2 | 16468 | ENSMUSG00000038518 | Degradome sequencing | positive | | validated |
| tarbase | MIMAT0000160 | mmu-miR-150-5p | Rpl12 | | ENSMUSG00000038900 | Degradome sequencing | positive | | validated |
| tarbase | MIMAT0000160 | mmu-miR-150-5p | Baz2a | 116848 | ENSMUSG00000040054 | Degradome sequencing//Degradome sequencing | positive | | validated |
| tarbase | MIMAT0000160 | mmu-miR-150-5p | Gramd1b | 235283 | ENSMUSG00000040111 | Degradome sequencing | positive | | validated |
| tarbase | MIMAT0000160 | mmu-miR-150-5p | Bach2 | 12014 | ENSMUSG00000040270 | Degradome sequencing | positive | | validated |
| tarbase | MIMAT0000160 | mmu-miR-150-5p | Phc1 | 13619 | ENSMUSG00000040669 | Degradome sequencing | positive | | validated |
| tarbase | MIMAT0000160 | mmu-miR-150-5p | Fam117b | 72750 | ENSMUSG00000041040 | Degradome sequencing | positive | | validated |
| tarbase | MIMAT0000160 | mmu-miR-150-5p | Ripk2 | 192656 | ENSMUSG00000041135 | Degradome sequencing | positive | | validated |
| tarbase | MIMAT0000160 | mmu-miR-150-5p | Ppp4r3a | 68734 | ENSMUSG00000041846 | Degradome sequencing | positive | | validated |
| tarbase | MIMAT0000160 | mmu-miR-150-5p | Kdm5b | 75605 | ENSMUSG00000042207 | Degradome sequencing | positive | | validated |

| | | | | | | | | | |
|---|---|---|---|---|---|---|---|---|---|
| tarbase | MIMAT0000160 | mmu-miR-150-5p | Zmym4 | 67785 | ENSMUSG00000042446 | Degradome sequencing | positive | | validated |
| tarbase | MIMAT0000160 | mmu-miR-150-5p | Sem1 | | ENSMUSG00000042541 | Degradome sequencing | positive | | validated |
| tarbase | MIMAT0000160 | mmu-miR-150-5p | Smg7 | 226517 | ENSMUSG00000042772 | Degradome sequencing | positive | | validated |
| tarbase | MIMAT0000160 | mmu-miR-150-5p | Hilpda | 69573 | ENSMUSG00000043421 | Degradome sequencing | positive | | validated |
| tarbase | MIMAT0000160 | mmu-miR-150-5p | Zbtb37 | 240869 | ENSMUSG00000043467 | Degradome sequencing | positive | | validated |
| tarbase | MIMAT0000160 | mmu-miR-150-5p | Swi5 | 72931 | ENSMUSG00000044627 | Degradome sequencing | positive | | validated |
| tarbase | MIMAT0000160 | mmu-miR-150-5p | Kmt5b | 225888 | ENSMUSG00000045098 | Degradome sequencing//Degradome sequencing | positive | | validated |
| tarbase | MIMAT0000160 | mmu-miR-150-5p | Zfp36l2 | 12193 | ENSMUSG00000045817 | Degradome sequencing//Degradome sequencing | positive | | validated |
| tarbase | MIMAT0000160 | mmu-miR-150-5p | Atxn1 | 20238 | ENSMUSG00000046876 | Degradome sequencing//Degradome sequencing | positive | | validated |
| tarbase | MIMAT0000160 | mmu-miR-150-5p | Cltc | 67300 | ENSMUSG00000047126 | Degradome sequencing | positive | | validated |
| tarbase | MIMAT0000160 | mmu-miR-150-5p | Tnrc6b | 213988 | ENSMUSG00000047888 | Degradome sequencing | positive | | validated |
| tarbase | MIMAT0000160 | mmu-miR-150-5p | Gli2 | 14633 | ENSMUSG00000048402 | Degradome sequencing | positive | | validated |
| tarbase | MIMAT0000160 | mmu-miR-150-5p | Mlec | 109154 | ENSMUSG00000048578 | Degradome sequencing | positive | | validated |
| tarbase | MIMAT0000160 | mmu-miR-150-5p | Foxo3 | 56484 | ENSMUSG00000048756 | Degradome sequencing | positive | | validated |
| tarbase | MIMAT0000160 | mmu-miR-150-5p | Smcr8 | 237782 | ENSMUSG00000049323 | Degradome sequencing | positive | | validated |
| tarbase | MIMAT0000160 | mmu-miR-150-5p | Aff4 | 93736 | ENSMUSG00000049470 | Degradome sequencing//Degradome sequencing | positive | | validated |
| tarbase | MIMAT0000160 | mmu-miR-150-5p | Tmsb4x | 19241 | ENSMUSG00000049775 | Degradome sequencing | positive | | validated |
| tarbase | MIMAT0000160 | mmu-miR-150-5p | Fgf4 | 14175 | ENSMUSG00000050917 | Degradome sequencing | positive | | validated |
| tarbase | MIMAT0000160 | mmu-miR-150-5p | Prkca | 18750 | ENSMUSG00000050965 | Degradome sequencing//Degradome sequencing | positive | | validated |
| tarbase | MIMAT0000160 | mmu-miR-150-5p | Irf2bp2 | 270110 | ENSMUSG00000051495 | Degradome sequencing | positive | | validated |
| tarbase | MIMAT0000160 | mmu-miR-150-5p | Mafg | 17134 | ENSMUSG00000051510 | Degradome sequencing | positive | | validated |
| tarbase | MIMAT0000160 | mmu-miR-150-5p | Tnrc6a | 233833 | ENSMUSG00000052707 | Degradome sequencing//Degradome sequencing | positive | | validated |
| tarbase | MIMAT0000160 | mmu-miR-150-5p | Cbx7 | 52609 | ENSMUSG00000053411 | Degradome sequencing | positive | | validated |
| tarbase | MIMAT0000160 | mmu-miR-150-5p | Parp4 | 328417 | ENSMUSG00000054509 | Degradome sequencing | positive | | validated |
| tarbase | MIMAT0000160 | mmu-miR-150-5p | Kdm2a | 225876 | ENSMUSG00000054611 | Degradome sequencing | positive | | validated |
| tarbase | MIMAT0000160 | mmu-miR-150-5p | Nfil3 | 18030 | ENSMUSG00000056749 | Degradome sequencing//Degradome sequencing | positive | | validated |
| tarbase | MIMAT0000160 | mmu-miR-150-5p | Npm1 | | ENSMUSG00000057113 | Degradome sequencing//Degradome sequencing | positive | | validated |
| tarbase | MIMAT0000160 | mmu-miR-150-5p | Slc44a2 | 68682 | ENSMUSG00000057193 | Degradome sequencing//Degradome sequencing//Degradome sequencing | positive | | validated |
| tarbase | MIMAT0000160 | mmu-miR-150-5p | Cep170 | 545389 | ENSMUSG00000057335 | Degradome sequencing//Degradome sequencing | positive | | validated |
| tarbase | MIMAT0000160 | mmu-miR-150-5p | Zfp329 | 67230 | ENSMUSG00000057894 | Degradome sequencing | positive | | validated |
| tarbase | MIMAT0000160 | mmu-miR-150-5p | Marf1 | 223989 | ENSMUSG00000060657 | Degradome sequencing | positive | | validated |
| tarbase | MIMAT0000160 | mmu-miR-150-5p | H2-K1 | 14972 | ENSMUSG00000061232 | Degradome sequencing | positive | | validated |
| tarbase | MIMAT0000160 | mmu-miR-150-5p | Cfl2 | 12632 | ENSMUSG00000062929 | Degradome sequencing//Degradome sequencing | positive | | validated |
| tarbase | MIMAT0000160 | mmu-miR-150-5p | Zfp26 | 22688 | ENSMUSG00000063108 | Degradome sequencing | positive | | validated |

| | | | | | | | | | |
|---|---|---|---|---|---|---|---|---|---|
| tarbase | MIMAT0000160 | mmu-miR-150-5p | Ldha | 16828 | ENSMUSG00000063229 | Degradome sequencing//Degradome sequencing | positive | | validated |
| tarbase | MIMAT0000160 | mmu-miR-150-5p | Arih2 | 23807 | ENSMUSG00000064145 | Degradome sequencing | positive | | validated |
| tarbase | MIMAT0000160 | mmu-miR-150-5p | Arfgef1 | 211673 | ENSMUSG00000067851 | Degradome sequencing | positive | | validated |
| tarbase | MIMAT0000160 | mmu-miR-150-5p | Tcp1 | 21454 | ENSMUSG00000068039 | Degradome sequencing//Degradome sequencing | positive | | validated |
| tarbase | MIMAT0000160 | mmu-miR-150-5p | Rnf121 | 75212 | ENSMUSG00000070426 | Degradome sequencing | positive | | validated |
| tarbase | MIMAT0000160 | mmu-miR-150-5p | Ptgr2 | 77219 | ENSMUSG00000072946 | Degradome sequencing | positive | | validated |
| tarbase | MIMAT0000160 | mmu-miR-150-5p | Csnk1g3 | 70425 | ENSMUSG00000073563 | Degradome sequencing//Degradome sequencing | positive | | validated |
| tarbase | MIMAT0000160 | mmu-miR-150-5p | Sox2 | | ENSMUSG00000074637 | Degradome sequencing | positive | | validated |
| tarbase | MIMAT0000160 | mmu-miR-150-5p | Pdzd8 | | ENSMUSG00000074746 | Degradome sequencing//Degradome sequencing | positive | | validated |
| tarbase | MIMAT0000160 | mmu-miR-150-5p | Wipf1 | 215280 | ENSMUSG00000075284 | Degradome sequencing | positive | | validated |
| tarbase | MIMAT0000160 | mmu-miR-150-5p | Selenot | | ENSMUSG00000075700 | Degradome sequencing | positive | | validated |
| tarbase | MIMAT0000160 | mmu-miR-150-5p | Ddi2 | 68817 | ENSMUSG00000078515 | Degradome sequencing | positive | | validated |
| tarbase | MIMAT0000160 | mmu-miR-150-5p | Ube2d3 | 66105 | ENSMUSG00000078578 | Degradome sequencing | positive | | validated |
| tarbase | MIMAT0000160 | mmu-miR-150-5p | Chd2 | 244059 | ENSMUSG00000078671 | Degradome sequencing//Degradome sequencing | positive | | validated |
| tarbase | MIMAT0000160 | mmu-miR-150-5p | Alkbh1 | | ENSMUSG00000079036 | Degradome sequencing | positive | | validated |
| tarbase | MIMAT0000160 | mmu-miR-150-5p | Ccr5 | 12774 | ENSMUSG00000079227 | Degradome sequencing | positive | | validated |
| tarbase | MIMAT0000160 | mmu-miR-150-5p | Hmgcs1 | 208715 | ENSMUSG00000093930 | Degradome sequencing | positive | | validated |
| tarbase | MIMAT0000160 | mmu-miR-150-5p | H1f0 | | ENSMUSG00000096210 | Degradome sequencing | positive | | validated |
| tarbase | MIMAT0000160 | mmu-miR-150-5p | Psmb9 | 16912 | ENSMUSG00000096727 | Degradome sequencing | positive | | validated |
| tarbase | MIMAT0000160 | mmu-miR-150-5p | | | ENSMUSG00000099515 | Degradome sequencing | positive | | validated |

| Downregulated in Aged Injured compared to Aged Control | | | | | | | | | |
|---|---|---|---|---|---|---|---|---|---|
| let-7e-5p targets | | | | | | | | | |
| **database** | **mature_mirna_acc** | **mature_mirna_id** | **target_symbol** | **target_entrez** | **target_ensembl** | **experiment** | **support_type** | **pubmed_id** | **type** |
| mirecords | MIMAT0000524 | mmu-let-7e-5p | Tlr4 | 21898 | ENSMUSG00000039005 | Luciferase activity assay | | 19699171 | validated |
| mirtarbase | MIMAT0000524 | mmu-let-7e-5p | Atp2b2 | 11941 | ENSMUSG00000030302 | HITS-CLIP | Functional MTI (Weak) | 21258322 | validated |
| mirtarbase | MIMAT0000524 | mmu-let-7e-5p | Col3a1 | 12825 | ENSMUSG00000026043 | qRT-PCR//Western blot | Functional MTI | 26272747 | validated |
| mirtarbase | MIMAT0000524 | mmu-let-7e-5p | Col1a1 | 12842 | ENSMUSG00000001506 | qRT-PCR//Western blot | Functional MTI | 26272747 | validated |
| mirtarbase | MIMAT0000524 | mmu-let-7e-5p | Col1a2 | 12843 | ENSMUSG00000029661 | qRT-PCR//Western blot | Functional MTI | 26272747 | validated |
| mirtarbase | MIMAT0000524 | mmu-let-7e-5p | Ifnar1 | 15975 | | HITS-CLIP | Functional MTI (Weak) | 21258322 | validated |
| mirtarbase | MIMAT0000524 | mmu-let-7e-5p | Il10 | 16153 | ENSMUSG00000016529 | Flow//Luciferase reporter assay//Microarray//qRT-PCR | Functional MTI | 23079871 | validated |
| mirtarbase | MIMAT0000524 | mmu-let-7e-5p | Il13 | 16163 | ENSMUSG00000020383 | Flow | Functional MTI (Weak) | 23079871 | validated |
| mirtarbase | MIMAT0000524 | mmu-let-7e-5p | Itga4 | 16401 | ENSMUSG00000027009 | qRT-PCR//Western blot | Functional MTI | 26272747 | validated |
| mirtarbase | MIMAT0000524 | mmu-let-7e-5p | Meis2 | 17536 | ENSMUSG00000027210 | HITS-CLIP | Functional MTI (Weak) | 23142080 | validated |
| mirtarbase | MIMAT0000524 | mmu-let-7e-5p | Nf2 | 18016 | ENSMUSG00000009073 | HITS-CLIP | Functional MTI (Weak) | 25083871 | validated |
| mirtarbase | MIMAT0000524 | mmu-let-7e-5p | Nf2 | 18016 | ENSMUSG00000009073 | HITS-CLIP | Functional MTI (Weak) | 23597149 | validated |
| mirtarbase | MIMAT0000524 | mmu-let-7e-5p | Scd1 | 20249 | ENSMUSG00000037071 | qRT-PCR//Western blot | Functional MTI | 26272747 | validated |
| mirtarbase | MIMAT0000524 | mmu-let-7e-5p | Thbs1 | 21825 | ENSMUSG00000040152 | qRT-PCR//Western blot | Functional MTI | 26272747 | validated |
| mirtarbase | MIMAT0000524 | mmu-let-7e-5p | Tlr4 | 21898 | | Luciferase reporter assay//qRT-PCR | Functional MTI | 19699171 | validated |
| mirtarbase | MIMAT0000524 | mmu-let-7e-5p | Apc2 | 23805 | ENSMUSG00000020135 | HITS-CLIP | Functional MTI (Weak) | 25083871 | validated |
| mirtarbase | MIMAT0000524 | mmu-let-7e-5p | Apc2 | 23805 | ENSMUSG00000020135 | HITS-CLIP | Functional MTI (Weak) | 23142080 | validated |
| mirtarbase | MIMAT0000524 | mmu-let-7e-5p | Apc2 | 23805 | ENSMUSG00000020135 | HITS-CLIP | Functional MTI (Weak) | 21258322 | validated |
| mirtarbase | MIMAT0000524 | mmu-let-7e-5p | Cts8 | 56094 | ENSMUSG00000057446 | HITS-CLIP | Functional MTI (Weak) | 25083871 | validated |
| mirtarbase | MIMAT0000524 | mmu-let-7e-5p | Cts8 | 56094 | ENSMUSG00000057446 | HITS-CLIP | Functional MTI (Weak) | 21258322 | validated |
| mirtarbase | MIMAT0000524 | mmu-let-7e-5p | Col24a1 | 71355 | ENSMUSG00000028197 | qRT-PCR//Western blot | Functional MTI | 26272747 | validated |
| mirtarbase | MIMAT0000524 | mmu-let-7e-5p | Lins1 | 72635 | ENSMUSG00000053091 | HITS-CLIP | Functional MTI (Weak) | 25083871 | validated |
| mirtarbase | MIMAT0000524 | mmu-let-7e-5p | Zfp444 | 72667 | ENSMUSG00000044876 | HITS-CLIP | Functional MTI (Weak) | 21258322 | validated |
| mirtarbase | MIMAT0000524 | mmu-let-7e-5p | Msi2 | 76626 | ENSMUSG00000069769 | HITS-CLIP | Functional MTI (Weak) | 25083871 | validated |
| mirtarbase | MIMAT0000524 | mmu-let-7e-5p | Eif3j1 | 78655 | | Reporter assay | Non-Functional MTI | 15131085 | validated |
| mirtarbase | MIMAT0000524 | mmu-let-7e-5p | Fbxl14 | 101358 | | HITS-CLIP | Functional MTI (Weak) | 25083871 | validated |
| mirtarbase | MIMAT0000524 | mmu-let-7e-5p | Itga1 | 109700 | ENSMUSG00000042284 | qRT-PCR//Western blot | Functional MTI | 26272747 | validated |

| | | | | | | | | | |
|---|---|---|---|---|---|---|---|---|---|
| mirtarbase | MIMAT0000524 | mmu-let-7e-5p | Armc2 | 213402 | ENSMUSG00000071324 | HITS-CLIP | Functional MTI (Weak) | 23597149 | validated |
| mirtarbase | MIMAT0000524 | mmu-let-7e-5p | Armc2 | 213402 | ENSMUSG00000071324 | HITS-CLIP | Functional MTI (Weak) | 21258322 | validated |
| mirtarbase | MIMAT0000524 | mmu-let-7e-5p | Nup214 | 227720 | ENSMUSG00000001855 | HITS-CLIP | Functional MTI (Weak) | 21258322 | validated |
| mirtarbase | MIMAT0000524 | mmu-let-7e-5p | Gnl3l | 237107 | ENSMUSG00000025266 | HITS-CLIP | Functional MTI (Weak) | 25083871 | validated |
| mirtarbase | MIMAT0000524 | mmu-let-7e-5p | Gnl3l | 237107 | ENSMUSG00000025266 | HITS-CLIP | Functional MTI (Weak) | 21258322 | validated |
| mirtarbase | MIMAT0000524 | mmu-let-7e-5p | Tnfrsf26 | 244237 | ENSMUSG00000045362 | HITS-CLIP | Functional MTI (Weak) | 21258322 | validated |
| mirtarbase | MIMAT0000524 | mmu-let-7e-5p | Col27a1 | 373864 | ENSMUSG00000045672 | qRT-PCR//Western blot | Functional MTI | 26272747 | validated |
| mirtarbase | MIMAT0000524 | mmu-let-7e-5p | Trim71 | 636931 | | Flow//GFP reporter assay//Immunocytochemistry//Immunohistochemistry//Immunoprecipitaion//In situ hybridization//Northern blot//qRT-PCR//Western blot | Functional MTI | 19898466 | validated |
| mirtarbase | MIMAT0000524 | mmu-let-7e-5p | Mup11 | 100039028 | ENSMUSG00000073834 | HITS-CLIP | Functional MTI (Weak) | 23597149 | validated |
| mirtarbase | MIMAT0000524 | mmu-let-7e-5p | Mup17 | 100039206 | ENSMUSG00000096688 | HITS-CLIP | Functional MTI (Weak) | 23597149 | validated |
| tarbase | MIMAT0000524 | mmu-let-7e-5p | Xpo6 | 74204 | ENSMUSG00000000131 | Degradome sequencing//Degradome sequencing | positive | | validated |
| tarbase | MIMAT0000524 | mmu-let-7e-5p | Ccnd2 | 12444 | ENSMUSG00000000184 | Degradome sequencing//Degradome sequencing | positive | | validated |
| tarbase | MIMAT0000524 | mmu-let-7e-5p | Trim25 | 217069 | ENSMUSG00000000275 | Degradome sequencing | positive | | validated |
| tarbase | MIMAT0000524 | mmu-let-7e-5p | Mnt | 17428 | ENSMUSG00000000282 | Degradome sequencing | positive | | validated |
| tarbase | MIMAT0000524 | mmu-let-7e-5p | Itgb2 | 16414 | ENSMUSG00000000290 | Degradome sequencing | positive | | validated |
| tarbase | MIMAT0000524 | mmu-let-7e-5p | Tmprss2 | 50528 | ENSMUSG00000000385 | Degradome sequencing//Degradome sequencing | positive | | validated |
| tarbase | MIMAT0000524 | mmu-let-7e-5p | Acvr1b | 11479 | ENSMUSG00000000532 | Degradome sequencing | positive | | validated |
| tarbase | MIMAT0000524 | mmu-let-7e-5p | Cd52 | 23833 | ENSMUSG00000000682 | Degradome sequencing//Degradome sequencing | positive | | validated |
| tarbase | MIMAT0000524 | mmu-let-7e-5p | Usp32 | 237898 | ENSMUSG00000000804 | Degradome sequencing | positive | | validated |
| tarbase | MIMAT0000524 | mmu-let-7e-5p | Mxd1 | 17119 | ENSMUSG00000001156 | Degradome sequencing | positive | | validated |
| tarbase | MIMAT0000524 | mmu-let-7e-5p | Calm1 | 12313 | ENSMUSG00000001175 | Degradome sequencing//Degradome sequencing | positive | | validated |
| tarbase | MIMAT0000524 | mmu-let-7e-5p | Agpat3 | 28169 | ENSMUSG00000001211 | Degradome sequencing | positive | | validated |
| tarbase | MIMAT0000524 | mmu-let-7e-5p | Sp1 | 20683 | ENSMUSG00000001280 | Degradome sequencing | positive | | validated |
| tarbase | MIMAT0000524 | mmu-let-7e-5p | Ell2 | | ENSMUSG00000001542 | Degradome sequencing | positive | | validated |
| tarbase | MIMAT0000524 | mmu-let-7e-5p | Ergic1 | 67458 | ENSMUSG00000001576 | Degradome sequencing | positive | | validated |
| tarbase | MIMAT0000524 | mmu-let-7e-5p | Gstt1 | 14871 | ENSMUSG00000001663 | Degradome sequencing//Degradome sequencing | positive | | validated |
| tarbase | MIMAT0000524 | mmu-let-7e-5p | Gramd3 | 107022 | ENSMUSG00000001700 | Degradome sequencing | positive | | validated |
| tarbase | MIMAT0000524 | mmu-let-7e-5p | Chordc1 | | ENSMUSG00000001774 | Degradome sequencing//Degradome sequencing | positive | | validated |
| tarbase | MIMAT0000524 | mmu-let-7e-5p | Kmt2a | 214162 | ENSMUSG00000002028 | Degradome sequencing | positive | | validated |
| tarbase | MIMAT0000524 | mmu-let-7e-5p | Celf2 | 14007 | ENSMUSG00000002107 | Degradome sequencing | positive | | validated |
| tarbase | MIMAT0000524 | mmu-let-7e-5p | Def6 | 23853 | ENSMUSG00000002257 | Degradome sequencing | positive | | validated |

| | | | | | | | | | |
|---|---|---|---|---|---|---|---|---|---|
| tarbase | MIMAT0000524 | mmu-let-7e-5p | Angptl4 | 57875 | ENSMUSG00000002289 | Degradome sequencing | positive | | validated |
| tarbase | MIMAT0000524 | mmu-let-7e-5p | Med6 | 69792 | ENSMUSG00000002679 | Degradome sequencing//Degradome sequencing | positive | | validated |
| tarbase | MIMAT0000524 | mmu-let-7e-5p | Baz1b | 22385 | ENSMUSG00000002748 | Degradome sequencing | positive | | validated |
| tarbase | MIMAT0000524 | mmu-let-7e-5p | Flii | 14248 | ENSMUSG00000002812 | Degradome sequencing//Degradome sequencing | positive | | validated |
| tarbase | MIMAT0000524 | mmu-let-7e-5p | Nab1 | 17936 | ENSMUSG00000002881 | Degradome sequencing | positive | | validated |
| tarbase | MIMAT0000524 | mmu-let-7e-5p | Hbp1 | 73389 | ENSMUSG00000002996 | Degradome sequencing | positive | | validated |
| tarbase | MIMAT0000524 | mmu-let-7e-5p | Cdk12 | 69131 | ENSMUSG00000003119 | Degradome sequencing | positive | | validated |
| tarbase | MIMAT0000524 | mmu-let-7e-5p | Slc2a3 | 20527 | ENSMUSG00000003153 | Degradome sequencing//Degradome sequencing | positive | | validated |
| tarbase | MIMAT0000524 | mmu-let-7e-5p | Dgcr2 | 13356 | ENSMUSG00000003166 | Degradome sequencing | positive | | validated |
| tarbase | MIMAT0000524 | mmu-let-7e-5p | Sh3gl1 | 20405 | ENSMUSG00000003200 | Degradome sequencing | positive | | validated |
| tarbase | MIMAT0000524 | mmu-let-7e-5p | Prkcsh | 19089 | ENSMUSG00000003402 | Degradome sequencing | positive | | validated |
| tarbase | MIMAT0000524 | mmu-let-7e-5p | Dusp3 | 72349 | ENSMUSG00000003518 | Degradome sequencing//Degradome sequencing | positive | | validated |
| tarbase | MIMAT0000524 | mmu-let-7e-5p | Prodh | 19125 | ENSMUSG00000003526 | Degradome sequencing | positive | | validated |
| tarbase | MIMAT0000524 | mmu-let-7e-5p | Aven | | ENSMUSG00000003604 | Degradome sequencing//Degradome sequencing | positive | | validated |
| tarbase | MIMAT0000524 | mmu-let-7e-5p | Snrnp200 | 320632 | ENSMUSG00000003660 | Degradome sequencing//Degradome sequencing//Degradome sequencing | positive | | validated |
| tarbase | MIMAT0000524 | mmu-let-7e-5p | Nfat5 | 54446 | ENSMUSG00000003847 | Degradome sequencing//Degradome sequencing | positive | | validated |
| tarbase | MIMAT0000524 | mmu-let-7e-5p | Stat3 | 20848 | ENSMUSG00000004040 | Degradome sequencing//Degradome sequencing | positive | | validated |
| tarbase | MIMAT0000524 | mmu-let-7e-5p | Phb2 | 12034 | ENSMUSG00000004264 | Degradome sequencing | positive | | validated |
| tarbase | MIMAT0000524 | mmu-let-7e-5p | Lpcat3 | | ENSMUSG00000004270 | Degradome sequencing//Degradome sequencing | positive | | validated |
| tarbase | MIMAT0000524 | mmu-let-7e-5p | Clcn3 | 12725 | ENSMUSG00000004319 | Degradome sequencing | positive | | validated |
| tarbase | MIMAT0000524 | mmu-let-7e-5p | Utp20 | 70683 | ENSMUSG00000004356 | Degradome sequencing | positive | | validated |
| tarbase | MIMAT0000524 | mmu-let-7e-5p | Coro1c | 23790 | ENSMUSG00000004530 | Degradome sequencing//Degradome sequencing//Degradome sequencing | positive | | validated |
| tarbase | MIMAT0000524 | mmu-let-7e-5p | Ndrg2 | 29811 | ENSMUSG00000004558 | Degradome sequencing//Degradome sequencing | positive | | validated |
| tarbase | MIMAT0000524 | mmu-let-7e-5p | Pkn2 | 109333 | ENSMUSG00000004591 | Degradome sequencing//Degradome sequencing | positive | | validated |
| tarbase | MIMAT0000524 | mmu-let-7e-5p | Arid3b | 56380 | ENSMUSG00000004661 | Degradome sequencing//Degradome sequencing | positive | | validated |
| tarbase | MIMAT0000524 | mmu-let-7e-5p | Myo9b | 17925 | ENSMUSG00000004677 | Degradome sequencing | positive | | validated |
| tarbase | MIMAT0000524 | mmu-let-7e-5p | Ulk2 | 29869 | ENSMUSG00000004798 | Degradome sequencing//Degradome sequencing//Degradome sequencing | positive | | validated |
| tarbase | MIMAT0000524 | mmu-let-7e-5p | Lbr | 98386 | ENSMUSG00000004880 | Degradome sequencing//Degradome sequencing | positive | | validated |
| tarbase | MIMAT0000524 | mmu-let-7e-5p | Pex5 | 19305 | ENSMUSG00000005069 | Degradome sequencing | positive | | validated |
| tarbase | MIMAT0000524 | mmu-let-7e-5p | Wdr1 | 22388 | ENSMUSG00000005103 | Degradome sequencing//Degradome sequencing | positive | | validated |
| tarbase | MIMAT0000524 | mmu-let-7e-5p | Polr2a | 20020 | ENSMUSG00000005198 | Degradome sequencing | positive | | validated |
| tarbase | MIMAT0000524 | mmu-let-7e-5p | Prlr | 19116 | ENSMUSG00000005268 | Degradome sequencing//Degradome sequencing | positive | | validated |

| | | | | | | | | | |
|---|---|---|---|---|---|---|---|---|---|
| tarbase | MIMAT0000524 | mmu-let-7e-5p | Txn2 | 56551 | ENSMUSG00000005354 | Degradome sequencing//Degradome sequencing | positive | | validated |
| tarbase | MIMAT0000524 | mmu-let-7e-5p | Crbn | | ENSMUSG00000005362 | Degradome sequencing//Degradome sequencing | positive | | validated |
| tarbase | MIMAT0000524 | mmu-let-7e-5p | Celf1 | 13046 | ENSMUSG00000005506 | Degradome sequencing | positive | | validated |
| tarbase | MIMAT0000524 | mmu-let-7e-5p | Por | 18984 | ENSMUSG00000005514 | Degradome sequencing//Degradome sequencing | positive | | validated |
| tarbase | MIMAT0000524 | mmu-let-7e-5p | Insr | 16337 | ENSMUSG00000005534 | Degradome sequencing//Degradome sequencing | positive | | validated |
| tarbase | MIMAT0000524 | mmu-let-7e-5p | Eif4g2 | 13690 | ENSMUSG00000005610 | Degradome sequencing//Degradome sequencing | positive | | validated |
| tarbase | MIMAT0000524 | mmu-let-7e-5p | Pcyt1a | 13026 | ENSMUSG00000005615 | Degradome sequencing | positive | | validated |
| tarbase | MIMAT0000524 | mmu-let-7e-5p | Cs | 12974 | ENSMUSG00000005683 | Degradome sequencing | positive | | validated |
| tarbase | MIMAT0000524 | mmu-let-7e-5p | Slc30a4 | 22785 | ENSMUSG00000005802 | Degradome sequencing | positive | | validated |
| tarbase | MIMAT0000524 | mmu-let-7e-5p | Metap1 | 75624 | ENSMUSG00000005813 | Degradome sequencing//Degradome sequencing//Degradome sequencing | positive | | validated |
| tarbase | MIMAT0000524 | mmu-let-7e-5p | Ncoa2 | 17978 | ENSMUSG00000005886 | Degradome sequencing//Degradome sequencing | positive | | validated |
| tarbase | MIMAT0000524 | mmu-let-7e-5p | Odr4 | 226499 | ENSMUSG00000006010 | Degradome sequencing | positive | | validated |
| tarbase | MIMAT0000524 | mmu-let-7e-5p | Kmt2b | 75410 | ENSMUSG00000006307 | Degradome sequencing | positive | | validated |
| tarbase | MIMAT0000524 | mmu-let-7e-5p | Gcat | 26912 | ENSMUSG00000006378 | Degradome sequencing | positive | | validated |
| tarbase | MIMAT0000524 | mmu-let-7e-5p | Mark3 | 17169 | ENSMUSG00000007411 | Degradome sequencing | positive | | validated |
| tarbase | MIMAT0000524 | mmu-let-7e-5p | Tgfbr1 | 21812 | ENSMUSG00000007613 | Degradome sequencing//Degradome sequencing//Degradome sequencing//Degradome sequencing//Degradome sequencing | positive | | validated |
| tarbase | MIMAT0000524 | mmu-let-7e-5p | Zmiz1 | 328365 | ENSMUSG00000007817 | Degradome sequencing | positive | | validated |
| tarbase | MIMAT0000524 | mmu-let-7e-5p | Fgfrl1 | 116701 | ENSMUSG00000008090 | Degradome sequencing | positive | | validated |
| tarbase | MIMAT0000524 | mmu-let-7e-5p | Prpf31 | 68988 | ENSMUSG00000008373 | Degradome sequencing | positive | | validated |
| tarbase | MIMAT0000524 | mmu-let-7e-5p | Sertad1 | 55942 | ENSMUSG00000008384 | Degradome sequencing//Degradome sequencing//Degradome sequencing//Degradome sequencing | positive | | validated |
| tarbase | MIMAT0000524 | mmu-let-7e-5p | Herpud2 | 80517 | ENSMUSG00000008429 | Degradome sequencing | positive | | validated |
| tarbase | MIMAT0000524 | mmu-let-7e-5p | Pou2f2 | 18987 | ENSMUSG00000008496 | Degradome sequencing//Degradome sequencing | positive | | validated |
| tarbase | MIMAT0000524 | mmu-let-7e-5p | Slc16a12 | 240638 | ENSMUSG00000009378 | Degradome sequencing | positive | | validated |
| tarbase | MIMAT0000524 | mmu-let-7e-5p | Tnpo1 | 238799 | ENSMUSG00000009470 | Degradome sequencing | positive | | validated |
| tarbase | MIMAT0000524 | mmu-let-7e-5p | Cbx5 | 12419 | ENSMUSG00000009575 | Degradome sequencing//Degradome sequencing//Degradome sequencing//Degradome sequencing//Degradome sequencing | positive | | validated |
| tarbase | MIMAT0000524 | mmu-let-7e-5p | Pla2g12b | 69836 | ENSMUSG00000009646 | Degradome sequencing | positive | | validated |
| tarbase | MIMAT0000524 | mmu-let-7e-5p | Slc38a3 | 76257 | ENSMUSG00000010064 | Degradome sequencing | positive | | validated |
| tarbase | MIMAT0000524 | mmu-let-7e-5p | Raver1 | 71766 | ENSMUSG00000010205 | Degradome sequencing | positive | | validated |
| tarbase | MIMAT0000524 | mmu-let-7e-5p | Kif11 | 16551 | ENSMUSG00000012443 | Degradome sequencing//Degradome sequencing | positive | | validated |
| tarbase | MIMAT0000524 | mmu-let-7e-5p | Atp6v0d1 | 11972 | ENSMUSG00000013160 | Degradome sequencing | positive | | validated |
| tarbase | MIMAT0000524 | mmu-let-7e-5p | Tmem17[illegible] | 72392 | ENSMUSG00000013495 | Degradome sequencing | positive | | validated |

| | | | | | | | | | |
|---|---|---|---|---|---|---|---|---|---|
| tarbase | MIMAT0000524 | mmu-let-7e-5p | Dedd | 21945 | ENSMUSG00000013973 | Degradome sequencing//Degradome sequencing//Degradome sequencing//Degradome sequencing//Degradome sequencing | positive | | validated |
| tarbase | MIMAT0000524 | mmu-let-7e-5p | Map3k4 | 26407 | ENSMUSG00000014426 | Degradome sequencing//Degradome sequencing//Degradome sequencing | positive | | validated |
| tarbase | MIMAT0000524 | mmu-let-7e-5p | Dnajb9 | | ENSMUSG00000014905 | Degradome sequencing | positive | | validated |
| tarbase | MIMAT0000524 | mmu-let-7e-5p | Slc25a13 | 50799 | ENSMUSG00000015112 | Degradome sequencing//Degradome sequencing//Degradome sequencing//Degradome sequencing//Degradome sequencing//Degradome sequencing | positive | | validated |
| tarbase | MIMAT0000524 | mmu-let-7e-5p | Slamf1 | 27218 | ENSMUSG00000015316 | Degradome sequencing//Degradome sequencing | positive | | validated |
| tarbase | MIMAT0000524 | mmu-let-7e-5p | Zdhhc12 | 66220 | ENSMUSG00000015335 | Degradome sequencing//Degradome sequencing | positive | | validated |
| tarbase | MIMAT0000524 | mmu-let-7e-5p | Cacfd1 | 381356 | ENSMUSG00000015488 | Degradome sequencing//Degradome sequencing | positive | | validated |
| tarbase | MIMAT0000524 | mmu-let-7e-5p | Arnt | 11863 | ENSMUSG00000015522 | Degradome sequencing | positive | | validated |
| tarbase | MIMAT0000524 | mmu-let-7e-5p | Setdb1 | 84505 | ENSMUSG00000015697 | Degradome sequencing//Degradome sequencing | positive | | validated |
| tarbase | MIMAT0000524 | mmu-let-7e-5p | Plekho1 | 67220 | ENSMUSG00000015745 | Degradome sequencing | positive | | validated |
| tarbase | MIMAT0000524 | mmu-let-7e-5p | Anp32e | 66471 | ENSMUSG00000015749 | Degradome sequencing//Degradome sequencing//Degradome sequencing//Degradome sequencing//Degradome sequencing | positive | | validated |
| tarbase | MIMAT0000524 | mmu-let-7e-5p | Tab2 | 68652 | ENSMUSG00000015755 | Degradome sequencing//Degradome sequencing | positive | | validated |
| tarbase | MIMAT0000524 | mmu-let-7e-5p | Chdh | 218865 | ENSMUSG00000015970 | Degradome sequencing//Degradome sequencing//Degradome sequencing//Degradome sequencing | positive | | validated |
| tarbase | MIMAT0000524 | mmu-let-7e-5p | Fli1 | 14247 | ENSMUSG00000016087 | Degradome sequencing//Degradome sequencing | positive | | validated |
| tarbase | MIMAT0000524 | mmu-let-7e-5p | Hsd11b1 | 15483 | ENSMUSG00000016194 | Degradome sequencing | positive | | validated |
| tarbase | MIMAT0000524 | mmu-let-7e-5p | Cd274 | 60533 | ENSMUSG00000016496 | Degradome sequencing//Degradome sequencing | positive | | validated |
| tarbase | MIMAT0000524 | mmu-let-7e-5p | Lamp2 | 16784 | ENSMUSG00000016534 | Degradome sequencing | positive | | validated |
| tarbase | MIMAT0000524 | mmu-let-7e-5p | Atxn10 | | ENSMUSG00000016541 | Degradome sequencing//Degradome sequencing | positive | | validated |
| tarbase | MIMAT0000524 | mmu-let-7e-5p | Eif3d | 55944 | ENSMUSG00000016554 | Degradome sequencing | positive | | validated |
| tarbase | MIMAT0000524 | mmu-let-7e-5p | H3f3b | 15081 | ENSMUSG00000016559 | Degradome sequencing | positive | | validated |
| tarbase | MIMAT0000524 | mmu-let-7e-5p | Cmah | 12763 | ENSMUSG00000016756 | Degradome sequencing//Degradome sequencing | positive | | validated |
| tarbase | MIMAT0000524 | mmu-let-7e-5p | Plcg1 | 18803 | ENSMUSG00000016933 | Degradome sequencing | positive | | validated |
| tarbase | MIMAT0000524 | mmu-let-7e-5p | Taok1 | 216965 | ENSMUSG00000017291 | Degradome sequencing//Degradome sequencing | positive | | validated |
| tarbase | MIMAT0000524 | mmu-let-7e-5p | Nlk | 18099 | ENSMUSG00000017376 | Degradome sequencing//Degradome sequencing//Degradome sequencing | positive | | validated |
| tarbase | MIMAT0000524 | mmu-let-7e-5p | Aldoc | 11676 | ENSMUSG00000017390 | Degradome sequencing | positive | | validated |

| | | | | | | | | | |
|---|---|---|---|---|---|---|---|---|---|
| tarbase | MIMAT0000524 | mmu-let-7e-5p | Arl5b | 75869 | ENSMUSG00000017418 | Degradome sequencing//Degradome sequencing | positive | | validated |
| tarbase | MIMAT0000524 | mmu-let-7e-5p | Suz12 | 52615 | ENSMUSG00000017548 | Degradome sequencing | positive | | validated |
| tarbase | MIMAT0000524 | mmu-let-7e-5p | Serinc3 | 26943 | ENSMUSG00000017707 | Degradome sequencing | positive | | validated |
| tarbase | MIMAT0000524 | mmu-let-7e-5p | Mmp9 | 17395 | ENSMUSG00000017737 | Degradome sequencing//Degradome sequencing//Degradome sequencing | positive | | validated |
| tarbase | MIMAT0000524 | mmu-let-7e-5p | Ctsa | 19025 | ENSMUSG00000017760 | Degradome sequencing//Degradome sequencing | positive | | validated |
| tarbase | MIMAT0000524 | mmu-let-7e-5p | Crk | 12928 | ENSMUSG00000017776 | Degradome sequencing//Degradome sequencing | positive | | validated |
| tarbase | MIMAT0000524 | mmu-let-7e-5p | Cyth3 | 19159 | ENSMUSG00000018001 | Degradome sequencing | positive | | validated |
| tarbase | MIMAT0000524 | mmu-let-7e-5p | Ro60 | 20822 | ENSMUSG00000018199 | Degradome sequencing | positive | | validated |
| tarbase | MIMAT0000524 | mmu-let-7e-5p | Il12rb2 | 16162 | ENSMUSG00000018341 | Degradome sequencing//Degradome sequencing | positive | | validated |
| tarbase | MIMAT0000524 | mmu-let-7e-5p | Srsf1 | 110809 | ENSMUSG00000018379 | Degradome sequencing | positive | | validated |
| tarbase | MIMAT0000524 | mmu-let-7e-5p | Akap1 | 11640 | ENSMUSG00000018428 | Degradome sequencing//Degradome sequencing | positive | | validated |
| tarbase | MIMAT0000524 | mmu-let-7e-5p | Appbp2 | | ENSMUSG00000018481 | Degradome sequencing//Degradome sequencing | positive | | validated |
| tarbase | MIMAT0000524 | mmu-let-7e-5p | Ncor1 | 20185 | ENSMUSG00000018501 | Degradome sequencing//Degradome sequencing | positive | | validated |
| tarbase | MIMAT0000524 | mmu-let-7e-5p | G3bp1 | | ENSMUSG00000018583 | Degradome sequencing | positive | | validated |
| tarbase | MIMAT0000524 | mmu-let-7e-5p | Tbx3 | 21386 | ENSMUSG00000018604 | Degradome sequencing | positive | | validated |
| tarbase | MIMAT0000524 | mmu-let-7e-5p | Pnpo | 103711 | ENSMUSG00000018659 | Degradome sequencing//Degradome sequencing | positive | | validated |
| tarbase | MIMAT0000524 | mmu-let-7e-5p | Dync1h1 | 13424 | ENSMUSG00000018707 | Degradome sequencing//Degradome sequencing | positive | | validated |
| tarbase | MIMAT0000524 | mmu-let-7e-5p | Ywhah | 22629 | ENSMUSG00000018965 | Degradome sequencing//Degradome sequencing//Degradome sequencing | positive | | validated |
| tarbase | MIMAT0000524 | mmu-let-7e-5p | Nars2 | 244141 | ENSMUSG00000018995 | Degradome sequencing | positive | | validated |
| tarbase | MIMAT0000524 | mmu-let-7e-5p | Slc35b4 | 58246 | ENSMUSG00000018999 | Degradome sequencing | positive | | validated |
| tarbase | MIMAT0000524 | mmu-let-7e-5p | H13 | 14950 | ENSMUSG00000019188 | Degradome sequencing | positive | | validated |
| tarbase | MIMAT0000524 | mmu-let-7e-5p | Gyg | 27357 | ENSMUSG00000019528 | Degradome sequencing | positive | | validated |
| tarbase | MIMAT0000524 | mmu-let-7e-5p | Arid3a | 13496 | ENSMUSG00000019564 | Degradome sequencing//Degradome sequencing | positive | | validated |
| tarbase | MIMAT0000524 | mmu-let-7e-5p | Cd164 | 53599 | ENSMUSG00000019818 | Degradome sequencing//Degradome sequencing | positive | | validated |
| tarbase | MIMAT0000524 | mmu-let-7e-5p | Utrn | 22288 | ENSMUSG00000019820 | Degradome sequencing//Degradome sequencing//Degradome sequencing//Degradome sequencing//Degradome sequencing | positive | | validated |
| tarbase | MIMAT0000524 | mmu-let-7e-5p | Slc16a10 | 72472 | ENSMUSG00000019838 | Degradome sequencing | positive | | validated |
| tarbase | MIMAT0000524 | mmu-let-7e-5p | Prep | | ENSMUSG00000019849 | Degradome sequencing//Degradome sequencing | positive | | validated |
| tarbase | MIMAT0000524 | mmu-let-7e-5p | Tnfaip3 | 21929 | ENSMUSG00000019850 | Degradome sequencing//Degradome sequencing | positive | | validated |
| tarbase | MIMAT0000524 | mmu-let-7e-5p | Rtn4ip1 | 170728 | ENSMUSG00000019864 | Degradome sequencing | positive | | validated |
| tarbase | MIMAT0000524 | mmu-let-7e-5p | Rhobtb1 | 69288 | ENSMUSG00000019944 | Degradome sequencing//Degradome sequencing | positive | | validated |
| tarbase | MIMAT0000524 | mmu-let-7e-5p | Uhrf1bp1 | 75089 | ENSMUSG00000019951 | Degradome sequencing | positive | | validated |

| | | | | | | | | | |
|---|---|---|---|---|---|---|---|---|---|
| tarbase | MIMAT0000524 | mmu-let-7e-5p | Dusp6 | 67603 | ENSMUSG00000019960 | Degradome sequencing//Degradome sequencing | positive | | validated |
| tarbase | MIMAT0000524 | mmu-let-7e-5p | Psen1 | 19164 | ENSMUSG00000019969 | Degradome sequencing//Degradome sequencing//Degradome sequencing | positive | | validated |
| tarbase | MIMAT0000524 | mmu-let-7e-5p | Apaf1 | 11783 | ENSMUSG00000019979 | Degradome sequencing | positive | | validated |
| tarbase | MIMAT0000524 | mmu-let-7e-5p | Fgd6 | 13998 | ENSMUSG00000020021 | Degradome sequencing | positive | | validated |
| tarbase | MIMAT0000524 | mmu-let-7e-5p | Tmcc3 | 319880 | ENSMUSG00000020023 | Degradome sequencing | positive | | validated |
| tarbase | MIMAT0000524 | mmu-let-7e-5p | Rfx4 | 71137 | ENSMUSG00000020037 | Degradome sequencing | positive | | validated |
| tarbase | MIMAT0000524 | mmu-let-7e-5p | Igf1 | 16000 | ENSMUSG00000020053 | Degradome sequencing//Degradome sequencing//Degradome sequencing//Degradome sequencing | positive | | validated |
| tarbase | MIMAT0000524 | mmu-let-7e-5p | Rufy2 | 70432 | ENSMUSG00000020070 | Degradome sequencing | positive | | validated |
| tarbase | MIMAT0000524 | mmu-let-7e-5p | Eif4ebp2 | 13688 | ENSMUSG00000020091 | Degradome sequencing//Degradome sequencing | positive | | validated |
| tarbase | MIMAT0000524 | mmu-let-7e-5p | Lrig3 | 320398 | ENSMUSG00000020105 | Degradome sequencing//Degradome sequencing | positive | | validated |
| tarbase | MIMAT0000524 | mmu-let-7e-5p | Tbk1 | 56480 | ENSMUSG00000020115 | Degradome sequencing//Degradome sequencing | positive | | validated |
| tarbase | MIMAT0000524 | mmu-let-7e-5p | Egfr | 13649 | ENSMUSG00000020122 | Degradome sequencing//Degradome sequencing | positive | | validated |
| tarbase | MIMAT0000524 | mmu-let-7e-5p | Vps54 | 245944 | ENSMUSG00000020128 | Degradome sequencing//Degradome sequencing//Degradome sequencing//Degradome sequencing//Degradome sequencing | positive | | validated |
| tarbase | MIMAT0000524 | mmu-let-7e-5p | Cpm | 70574 | ENSMUSG00000020183 | Degradome sequencing | positive | | validated |
| tarbase | MIMAT0000524 | mmu-let-7e-5p | Mdm2 | 17246 | ENSMUSG00000020184 | Degradome sequencing//Degradome sequencing | positive | | validated |
| tarbase | MIMAT0000524 | mmu-let-7e-5p | Mknk2 | 17347 | ENSMUSG00000020190 | Degradome sequencing//Degradome sequencing | positive | | validated |
| tarbase | MIMAT0000524 | mmu-let-7e-5p | Ncln | 103425 | ENSMUSG00000020238 | Degradome sequencing | positive | | validated |
| tarbase | MIMAT0000524 | mmu-let-7e-5p | Hcfc2 | 67933 | ENSMUSG00000020246 | Degradome sequencing//Degradome sequencing | positive | | validated |
| tarbase | MIMAT0000524 | mmu-let-7e-5p | Xpo1 | 103573 | ENSMUSG00000020290 | Degradome sequencing//Degradome sequencing | positive | | validated |
| tarbase | MIMAT0000524 | mmu-let-7e-5p | Cpeb4 | 67579 | ENSMUSG00000020300 | Degradome sequencing//Degradome sequencing//Degradome sequencing | positive | | validated |
| tarbase | MIMAT0000524 | mmu-let-7e-5p | Sptbn1 | 20742 | ENSMUSG00000020315 | Degradome sequencing//Degradome sequencing//Degradome sequencing | positive | | validated |
| tarbase | MIMAT0000524 | mmu-let-7e-5p | Il13 | 16163 | ENSMUSG00000020383 | Degradome sequencing | positive | | validated |
| tarbase | MIMAT0000524 | mmu-let-7e-5p | Ube2b | 22210 | ENSMUSG00000020390 | Degradome sequencing | positive | | validated |
| tarbase | MIMAT0000524 | mmu-let-7e-5p | Btg2 | | ENSMUSG00000020423 | Degradome sequencing//Degradome sequencing//Degradome sequencing//Degradome sequencing | positive | | validated |
| tarbase | MIMAT0000524 | mmu-let-7e-5p | Ppp4r3b | 104570 | ENSMUSG00000020463 | Degradome sequencing//Degradome sequencing | positive | | validated |
| tarbase | MIMAT0000524 | mmu-let-7e-5p | Rnft1 | 76892 | ENSMUSG00000020521 | Degradome sequencing//Degradome sequencing//Degradome sequencing | positive | | validated |
| tarbase | MIMAT0000524 | mmu-let-7e-5p | Fam49a | 76820 | ENSMUSG00000020589 | Degradome sequencing//Degradome sequencing//Degradome sequencing | positive | | validated |

| | | | | | | | | | |
|---|---|---|---|---|---|---|---|---|---|
| tarbase | MIMAT0000524 | mmu-let-7e-5p | Lpin1 | 14245 | ENSMUSG00000020593 | Degradome sequencing//Degradome sequencing | positive | | validated |
| tarbase | MIMAT0000524 | mmu-let-7e-5p | Trib2 | 217410 | ENSMUSG00000020601 | Degradome sequencing | positive | | validated |
| tarbase | MIMAT0000524 | mmu-let-7e-5p | Apob | 238055 | ENSMUSG00000020609 | Degradome sequencing | positive | | validated |
| tarbase | MIMAT0000524 | mmu-let-7e-5p | Itsn2 | 20403 | ENSMUSG00000020640 | Degradome sequencing//Degradome sequencing//Degradome sequencing//Degradome sequencing | positive | | validated |
| tarbase | MIMAT0000524 | mmu-let-7e-5p | Bcap29 | 12033 | ENSMUSG00000020650 | Degradome sequencing | positive | | validated |
| tarbase | MIMAT0000524 | mmu-let-7e-5p | Dnmt3a | 13435 | ENSMUSG00000020661 | Degradome sequencing//Degradome sequencing | positive | | validated |
| tarbase | MIMAT0000524 | mmu-let-7e-5p | Itgb3 | 16416 | ENSMUSG00000020689 | Degradome sequencing//Degradome sequencing | positive | | validated |
| tarbase | MIMAT0000524 | mmu-let-7e-5p | Rffl | 67338 | ENSMUSG00000020696 | Degradome sequencing//Degradome sequencing//Degradome sequencing | positive | | validated |
| tarbase | MIMAT0000524 | mmu-let-7e-5p | Map3k3 | 26406 | ENSMUSG00000020700 | Degradome sequencing//Degradome sequencing | positive | | validated |
| tarbase | MIMAT0000524 | mmu-let-7e-5p | Gga3 | 260302 | ENSMUSG00000020740 | Degradome sequencing | positive | | validated |
| tarbase | MIMAT0000524 | mmu-let-7e-5p | Ankfy1 | 11736 | ENSMUSG00000020790 | Degradome sequencing | positive | | validated |
| tarbase | MIMAT0000524 | mmu-let-7e-5p | Stat5b | 20851 | ENSMUSG00000020919 | Degradome sequencing | positive | | validated |
| tarbase | MIMAT0000524 | mmu-let-7e-5p | Tmem10 | 76547 | ENSMUSG00000020921 | Degradome sequencing | positive | | validated |
| tarbase | MIMAT0000524 | mmu-let-7e-5p | Map3k14 | 53859 | ENSMUSG00000020941 | Degradome sequencing | positive | | validated |
| tarbase | MIMAT0000524 | mmu-let-7e-5p | Sel1l | 20338 | ENSMUSG00000020964 | Degradome sequencing//Degradome sequencing | positive | | validated |
| tarbase | MIMAT0000524 | mmu-let-7e-5p | Pnn | 18949 | ENSMUSG00000020994 | Degradome sequencing | positive | | validated |
| tarbase | MIMAT0000524 | mmu-let-7e-5p | Hif1a | 15251 | ENSMUSG00000021109 | Degradome sequencing//Degradome sequencing | positive | | validated |
| tarbase | MIMAT0000524 | mmu-let-7e-5p | Pacs2 | 217893 | ENSMUSG00000021143 | Degradome sequencing | positive | | validated |
| tarbase | MIMAT0000524 | mmu-let-7e-5p | Wdr37 | 207615 | ENSMUSG00000021147 | Degradome sequencing//Degradome sequencing//Degradome sequencing | positive | | validated |
| tarbase | MIMAT0000524 | mmu-let-7e-5p | Esyt2 | 52635 | ENSMUSG00000021171 | Degradome sequencing | positive | | validated |
| tarbase | MIMAT0000524 | mmu-let-7e-5p | Pfkp | 56421 | ENSMUSG00000021196 | Degradome sequencing//Degradome sequencing | positive | | validated |
| tarbase | MIMAT0000524 | mmu-let-7e-5p | Entpd5 | 12499 | ENSMUSG00000021236 | Degradome sequencing//Degradome sequencing | positive | | validated |
| tarbase | MIMAT0000524 | mmu-let-7e-5p | Aldh6a1 | 104776 | ENSMUSG00000021238 | Degradome sequencing | positive | | validated |
| tarbase | MIMAT0000524 | mmu-let-7e-5p | Traf3 | 22031 | ENSMUSG00000021277 | Degradome sequencing | positive | | validated |
| tarbase | MIMAT0000524 | mmu-let-7e-5p | Gpr132 | 56696 | ENSMUSG00000021298 | Degradome sequencing//Degradome sequencing | positive | | validated |
| tarbase | MIMAT0000524 | mmu-let-7e-5p | Irf4 | 16364 | ENSMUSG00000021356 | Degradome sequencing//Degradome sequencing | positive | | validated |
| tarbase | MIMAT0000524 | mmu-let-7e-5p | Elovl2 | 54326 | ENSMUSG00000021364 | Degradome sequencing//Degradome sequencing//Degradome sequencing | positive | | validated |
| tarbase | MIMAT0000524 | mmu-let-7e-5p | Nup153 | 218210 | ENSMUSG00000021374 | Degradome sequencing | positive | | validated |
| tarbase | MIMAT0000524 | mmu-let-7e-5p | Sema4d | 20354 | ENSMUSG00000021451 | Degradome sequencing//Degradome sequencing | positive | | validated |
| tarbase | MIMAT0000524 | mmu-let-7e-5p | Ptch1 | 19206 | ENSMUSG00000021466 | Degradome sequencing | positive | | validated |
| tarbase | MIMAT0000524 | mmu-let-7e-5p | Ccnh | 66671 | ENSMUSG00000021548 | Degradome sequencing | positive | | validated |
| tarbase | MIMAT0000524 | mmu-let-7e-5p | Dapk1 | 69635 | ENSMUSG00000021559 | Degradome sequencing | positive | | validated |
| tarbase | MIMAT0000524 | mmu-let-7e-5p | Trip13 | | ENSMUSG00000021569 | Degradome sequencing//Degradome sequencing | positive | | validated |

| | | | | | | | | | |
|---|---|---|---|---|---|---|---|---|---|
| tarbase | MIMAT0000524 | mmu-let-7e-5p | Clptm1l | 218335 | ENSMUSG00000021610 | Degradome sequencing | positive | | validated |
| tarbase | MIMAT0000524 | mmu-let-7e-5p | Marveld2 | 218518 | ENSMUSG00000021636 | Degradome sequencing//Degradome sequencing | positive | | validated |
| tarbase | MIMAT0000524 | mmu-let-7e-5p | Gfm2 | 320806 | ENSMUSG00000021666 | Degradome sequencing | positive | | validated |
| tarbase | MIMAT0000524 | mmu-let-7e-5p | Iqgap2 | 544963 | ENSMUSG00000021676 | Degradome sequencing//Degradome sequencing//Degradome sequencing | positive | | validated |
| tarbase | MIMAT0000524 | mmu-let-7e-5p | Kif2a | 16563 | ENSMUSG00000021693 | Degradome sequencing//Degradome sequencing | positive | | validated |
| tarbase | MIMAT0000524 | mmu-let-7e-5p | Slc4a7 | 218756 | ENSMUSG00000021733 | Degradome sequencing//Degradome sequencing | positive | | validated |
| tarbase | MIMAT0000524 | mmu-let-7e-5p | Map3k1 | 26401 | ENSMUSG00000021754 | Degradome sequencing//Degradome sequencing//Degradome sequencing | positive | | validated |
| tarbase | MIMAT0000524 | mmu-let-7e-5p | Vcl | 22330 | ENSMUSG00000021823 | Degradome sequencing | positive | | validated |
| tarbase | MIMAT0000524 | mmu-let-7e-5p | Ap3m1 | 55946 | ENSMUSG00000021824 | Degradome sequencing | positive | | validated |
| tarbase | MIMAT0000524 | mmu-let-7e-5p | Mettl6 | 67011 | ENSMUSG00000021891 | Degradome sequencing//Degradome sequencing | positive | | validated |
| tarbase | MIMAT0000524 | mmu-let-7e-5p | Arhgef3 | 71704 | ENSMUSG00000021895 | Degradome sequencing | positive | | validated |
| tarbase | MIMAT0000524 | mmu-let-7e-5p | Ctsb | 13030 | ENSMUSG00000021939 | Degradome sequencing | positive | | validated |
| tarbase | MIMAT0000524 | mmu-let-7e-5p | Xpo4 | 57258 | ENSMUSG00000021952 | Degradome sequencing | positive | | validated |
| tarbase | MIMAT0000524 | mmu-let-7e-5p | Rb1 | 19645 | ENSMUSG00000022105 | Degradome sequencing | positive | | validated |
| tarbase | MIMAT0000524 | mmu-let-7e-5p | Lrp10 | | ENSMUSG00000022175 | Degradome sequencing | positive | | validated |
| tarbase | MIMAT0000524 | mmu-let-7e-5p | Golph3 | 66629 | ENSMUSG00000022200 | Degradome sequencing//Degradome sequencing | positive | | validated |
| tarbase | MIMAT0000524 | mmu-let-7e-5p | Dcaf11 | 28199 | ENSMUSG00000022214 | Degradome sequencing | positive | | validated |
| tarbase | MIMAT0000524 | mmu-let-7e-5p | Myc | 17869 | ENSMUSG00000022346 | Degradome sequencing//Degradome sequencing//Degradome sequencing | positive | | validated |
| tarbase | MIMAT0000524 | mmu-let-7e-5p | Fbxo32 | 67731 | ENSMUSG00000022358 | Degradome sequencing//Degradome sequencing//Degradome sequencing//Degradome sequencing//Degradome sequencing | positive | | validated |
| tarbase | MIMAT0000524 | mmu-let-7e-5p | Asap1 | 13196 | ENSMUSG00000022377 | Degradome sequencing | positive | | validated |
| tarbase | MIMAT0000524 | mmu-let-7e-5p | Mief1 | 239555 | ENSMUSG00000022412 | Degradome sequencing//Degradome sequencing | positive | | validated |
| tarbase | MIMAT0000524 | mmu-let-7e-5p | Myh9 | 17886 | ENSMUSG00000022443 | Degradome sequencing//Degradome sequencing//Degradome sequencing//Degradome sequencing | positive | | validated |
| tarbase | MIMAT0000524 | mmu-let-7e-5p | Slc38a2 | 67760 | ENSMUSG00000022462 | Degradome sequencing//Degradome sequencing | positive | | validated |
| tarbase | MIMAT0000524 | mmu-let-7e-5p | Txndc11 | 106200 | ENSMUSG00000022498 | Degradome sequencing | positive | | validated |
| tarbase | MIMAT0000524 | mmu-let-7e-5p | 1810013I | 69053 | ENSMUSG00000022507 | Degradome sequencing//Degradome sequencing | positive | | validated |
| tarbase | MIMAT0000524 | mmu-let-7e-5p | Crebbp | 12914 | ENSMUSG00000022521 | Degradome sequencing//Degradome sequencing//Degradome sequencing//Degradome sequencing//Degradome sequencing | positive | | validated |
| tarbase | MIMAT0000524 | mmu-let-7e-5p | Atp13a3 | 224088 | ENSMUSG00000022533 | Degradome sequencing//Degradome sequencing//Degradome sequencing | positive | | validated |
| tarbase | MIMAT0000524 | mmu-let-7e-5p | Hsf1 | 15499 | ENSMUSG00000022556 | Degradome sequencing | positive | | validated |

| | | | | | | | | | |
|---|---|---|---|---|---|---|---|---|---|
| tarbase | MIMAT0000524 | mmu-let-7e-5p | Plec | 18810 | ENSMUSG00000022565 | Degradome sequencing//Degradome sequencing | positive | | validated |
| tarbase | MIMAT0000524 | mmu-let-7e-5p | Cep97 | 74201 | ENSMUSG00000022604 | Degradome sequencing//Degradome sequencing | positive | | validated |
| tarbase | MIMAT0000524 | mmu-let-7e-5p | Alcam | | ENSMUSG00000022636 | Degradome sequencing | positive | | validated |
| tarbase | MIMAT0000524 | mmu-let-7e-5p | Cblb | 208650 | ENSMUSG00000022637 | Degradome sequencing | positive | | validated |
| tarbase | MIMAT0000524 | mmu-let-7e-5p | Cpox | 12892 | ENSMUSG00000022742 | Degradome sequencing | positive | | validated |
| tarbase | MIMAT0000524 | mmu-let-7e-5p | Senp5 | 320213 | ENSMUSG00000022772 | Degradome sequencing//Degradome sequencing//Degradome sequencing | positive | | validated |
| tarbase | MIMAT0000524 | mmu-let-7e-5p | Tfrc | 22042 | ENSMUSG00000022797 | Degradome sequencing//Degradome sequencing | positive | | validated |
| tarbase | MIMAT0000524 | mmu-let-7e-5p | Rabl3 | 67657 | ENSMUSG00000022827 | Degradome sequencing | positive | | validated |
| tarbase | MIMAT0000524 | mmu-let-7e-5p | Ehhadh | 74147 | ENSMUSG00000022853 | Degradome sequencing//Degradome sequencing//Degradome sequencing | positive | | validated |
| tarbase | MIMAT0000524 | mmu-let-7e-5p | Kpna1 | 16646 | ENSMUSG00000022905 | Degradome sequencing | positive | | validated |
| tarbase | MIMAT0000524 | mmu-let-7e-5p | Brwd1 | 93871 | ENSMUSG00000022914 | Degradome sequencing//Degradome sequencing//Degradome sequencing//Degradome sequencing//Degradome sequencing | positive | | validated |
| tarbase | MIMAT0000524 | mmu-let-7e-5p | Il10rb | 16155 | ENSMUSG00000022969 | Degradome sequencing | positive | | validated |
| tarbase | MIMAT0000524 | mmu-let-7e-5p | Lmbr1l | 74775 | ENSMUSG00000022999 | Degradome sequencing//Degradome sequencing | positive | | validated |
| tarbase | MIMAT0000524 | mmu-let-7e-5p | Akirin1 | | ENSMUSG00000023075 | Degradome sequencing//Degradome sequencing | positive | | validated |
| tarbase | MIMAT0000524 | mmu-let-7e-5p | Igf2r | 16004 | ENSMUSG00000023830 | Degradome sequencing//Degradome sequencing | positive | | validated |
| tarbase | MIMAT0000524 | mmu-let-7e-5p | Chd1 | 12648 | ENSMUSG00000023852 | Degradome sequencing//Degradome sequencing//Degradome sequencing//Degradome sequencing//Degradome sequencing | positive | | validated |
| tarbase | MIMAT0000524 | mmu-let-7e-5p | Cenpq | | ENSMUSG00000023919 | Degradome sequencing | positive | | validated |
| tarbase | MIMAT0000524 | mmu-let-7e-5p | Hsp90ab | 15516 | ENSMUSG00000023944 | Degradome sequencing//Degradome sequencing//Degradome sequencing//Degradome sequencing//Degradome sequencing//Degradome sequencing | positive | | validated |
| tarbase | MIMAT0000524 | mmu-let-7e-5p | Vegfa | 22339 | ENSMUSG00000023951 | Degradome sequencing | positive | | validated |
| tarbase | MIMAT0000524 | mmu-let-7e-5p | Ubr2 | 224826 | ENSMUSG00000023977 | Degradome sequencing//Degradome sequencing//Degradome sequencing | positive | | validated |
| tarbase | MIMAT0000524 | mmu-let-7e-5p | Glo1 | | ENSMUSG00000024026 | Degradome sequencing//Degradome sequencing | positive | | validated |
| tarbase | MIMAT0000524 | mmu-let-7e-5p | Wiz | 22404 | ENSMUSG00000024050 | Degradome sequencing | positive | | validated |
| tarbase | MIMAT0000524 | mmu-let-7e-5p | Lbh | 77889 | ENSMUSG00000024063 | Degradome sequencing//Degradome sequencing | positive | | validated |
| tarbase | MIMAT0000524 | mmu-let-7e-5p | Birc6 | 12211 | ENSMUSG00000024073 | Degradome sequencing | positive | | validated |
| tarbase | MIMAT0000524 | mmu-let-7e-5p | Strn | 268980 | ENSMUSG00000024077 | Degradome sequencing | positive | | validated |
| tarbase | MIMAT0000524 | mmu-let-7e-5p | Man2a1 | | ENSMUSG00000024085 | Degradome sequencing | positive | | validated |
| tarbase | MIMAT0000524 | mmu-let-7e-5p | Pdpk1 | 18607 | ENSMUSG00000024122 | Degradome sequencing | positive | | validated |

| | | | | | | | | | |
|---|---|---|---|---|---|---|---|---|---|
| tarbase | MIMAT0000524 | mmu-let-7e-5p | Abca3 | 27410 | ENSMUSG00000024130 | Degradome sequencing | positive | | validated |
| tarbase | MIMAT0000524 | mmu-let-7e-5p | C3 | 12266 | ENSMUSG00000024164 | Degradome sequencing//Degradome sequencing | positive | | validated |
| tarbase | MIMAT0000524 | mmu-let-7e-5p | Dusp1 | 19252 | ENSMUSG00000024190 | Degradome sequencing//Degradome sequencing | positive | | validated |
| tarbase | MIMAT0000524 | mmu-let-7e-5p | Cul2 | 71745 | ENSMUSG00000024231 | Degradome sequencing | positive | | validated |
| tarbase | MIMAT0000524 | mmu-let-7e-5p | Svil | 225115 | ENSMUSG00000024236 | Degradome sequencing//Degradome sequencing | positive | | validated |
| tarbase | MIMAT0000524 | mmu-let-7e-5p | Epc1 | 13831 | ENSMUSG00000024240 | Degradome sequencing//Degradome sequencing//Degradome sequencing | positive | | validated |
| tarbase | MIMAT0000524 | mmu-let-7e-5p | Sos1 | 20662 | ENSMUSG00000024241 | Degradome sequencing//Degradome sequencing | positive | | validated |
| tarbase | MIMAT0000524 | mmu-let-7e-5p | Map4k3 | 225028 | ENSMUSG00000024242 | Degradome sequencing | positive | | validated |
| tarbase | MIMAT0000524 | mmu-let-7e-5p | Abcg8 | 67470 | ENSMUSG00000024254 | Degradome sequencing | positive | | validated |
| tarbase | MIMAT0000524 | mmu-let-7e-5p | Mapre2 | 212307 | ENSMUSG00000024277 | Degradome sequencing | positive | | validated |
| tarbase | MIMAT0000524 | mmu-let-7e-5p | Wac | 225131 | ENSMUSG00000024283 | Degradome sequencing | positive | | validated |
| tarbase | MIMAT0000524 | mmu-let-7e-5p | Mib1 | 225164 | ENSMUSG00000024294 | Degradome sequencing | positive | | validated |
| tarbase | MIMAT0000524 | mmu-let-7e-5p | Tapbp | 21356 | ENSMUSG00000024308 | Degradome sequencing | positive | | validated |
| tarbase | MIMAT0000524 | mmu-let-7e-5p | Tmem17 | 72512 | ENSMUSG00000024349 | Degradome sequencing | positive | | validated |
| tarbase | MIMAT0000524 | mmu-let-7e-5p | C2 | 12263 | ENSMUSG00000024371 | Degradome sequencing | positive | | validated |
| tarbase | MIMAT0000524 | mmu-let-7e-5p | Map3k2 | 26405 | ENSMUSG00000024383 | Degradome sequencing | positive | | validated |
| tarbase | MIMAT0000524 | mmu-let-7e-5p | Bag6 | 224727 | ENSMUSG00000024392 | Degradome sequencing | positive | | validated |
| tarbase | MIMAT0000524 | mmu-let-7e-5p | Tnf | 21926 | ENSMUSG00000024401 | Degradome sequencing | positive | | validated |
| tarbase | MIMAT0000524 | mmu-let-7e-5p | Riok3 | 66878 | ENSMUSG00000024404 | Degradome sequencing//Degradome sequencing | positive | | validated |
| tarbase | MIMAT0000524 | mmu-let-7e-5p | Hdac3 | 15183 | ENSMUSG00000024454 | Degradome sequencing | positive | | validated |
| tarbase | MIMAT0000524 | mmu-let-7e-5p | Tcerg1 | 56070 | ENSMUSG00000024498 | Degradome sequencing//Degradome sequencing | positive | | validated |
| tarbase | MIMAT0000524 | mmu-let-7e-5p | Pmaip1 | 58801 | ENSMUSG00000024521 | Degradome sequencing//Degradome sequencing | positive | | validated |
| tarbase | MIMAT0000524 | mmu-let-7e-5p | Csnk1a1 | 93687 | ENSMUSG00000024576 | Degradome sequencing//Degradome sequencing | positive | | validated |
| tarbase | MIMAT0000524 | mmu-let-7e-5p | Lmnb1 | 16906 | ENSMUSG00000024590 | Degradome sequencing//Degradome sequencing | positive | | validated |
| tarbase | MIMAT0000524 | mmu-let-7e-5p | Fads3 | | ENSMUSG00000024664 | Degradome sequencing | positive | | validated |
| tarbase | MIMAT0000524 | mmu-let-7e-5p | Ms4a6b | 69774 | ENSMUSG00000024677 | Degradome sequencing | positive | | validated |
| tarbase | MIMAT0000524 | mmu-let-7e-5p | Ddb1 | 13194 | ENSMUSG00000024740 | Degradome sequencing//Degradome sequencing//Degradome sequencing | positive | | validated |
| tarbase | MIMAT0000524 | mmu-let-7e-5p | Cemip2 | 83921 | ENSMUSG00000024754 | Degradome sequencing//Degradome sequencing//Degradome sequencing//Degradome sequencing//Degradome sequencing | positive | | validated |
| tarbase | MIMAT0000524 | mmu-let-7e-5p | Rtn3 | 20168 | ENSMUSG00000024758 | Degradome sequencing//Degradome sequencing | positive | | validated |
| tarbase | MIMAT0000524 | mmu-let-7e-5p | Syvn1 | 74126 | ENSMUSG00000024807 | Degradome sequencing | positive | | validated |
| tarbase | MIMAT0000524 | mmu-let-7e-5p | Uhrf2 | 109113 | ENSMUSG00000024817 | Degradome sequencing//Degradome sequencing | positive | | validated |
| tarbase | MIMAT0000524 | mmu-let-7e-5p | Slc25a45 | 107375 | ENSMUSG00000024818 | Degradome sequencing//Degradome sequencing | positive | | validated |

| | | | | | | | | | |
|---|---|---|---|---|---|---|---|---|---|
| tarbase | MIMAT0000524 | mmu-let-7e-5p | Rps6kb2 | 58988 | ENSMUSG00000024830 | Degradome sequencing//Degradome sequencing//Degradome sequencing | positive | | validated |
| tarbase | MIMAT0000524 | mmu-let-7e-5p | Rab1b | 76308 | ENSMUSG00000024870 | Degradome sequencing//Degradome sequencing//Degradome sequencing//Degradome sequencing//Degradome sequencing | positive | | validated |
| tarbase | MIMAT0000524 | mmu-let-7e-5p | Scyl1 | 78891 | ENSMUSG00000024941 | Degradome sequencing | positive | | validated |
| tarbase | MIMAT0000524 | mmu-let-7e-5p | Cep55 | 74107 | ENSMUSG00000024989 | Degradome sequencing | positive | | validated |
| tarbase | MIMAT0000524 | mmu-let-7e-5p | Prdx3 | 11757 | ENSMUSG00000024997 | Degradome sequencing | positive | | validated |
| tarbase | MIMAT0000524 | mmu-let-7e-5p | Ccnj | 240665 | ENSMUSG00000025010 | Degradome sequencing//Degradome sequencing//Degradome sequencing | positive | | validated |
| tarbase | MIMAT0000524 | mmu-let-7e-5p | Msr1 | 20288 | ENSMUSG00000025044 | Degradome sequencing | positive | | validated |
| tarbase | MIMAT0000524 | mmu-let-7e-5p | Taf5 | 226182 | ENSMUSG00000025049 | Degradome sequencing//Degradome sequencing | positive | | validated |
| tarbase | MIMAT0000524 | mmu-let-7e-5p | Cenpx | 20892 | ENSMUSG00000025144 | Degradome sequencing | positive | | validated |
| tarbase | MIMAT0000524 | mmu-let-7e-5p | Slc16a3 | 80879 | ENSMUSG00000025161 | Degradome sequencing//Degradome sequencing | positive | | validated |
| tarbase | MIMAT0000524 | mmu-let-7e-5p | Loxl4 | | ENSMUSG00000025185 | Degradome sequencing | positive | | validated |
| tarbase | MIMAT0000524 | mmu-let-7e-5p | Chuk | 12675 | ENSMUSG00000025199 | Degradome sequencing | positive | | validated |
| tarbase | MIMAT0000524 | mmu-let-7e-5p | Scd2 | 20250 | ENSMUSG00000025203 | Degradome sequencing//Degradome sequencing | positive | | validated |
| tarbase | MIMAT0000524 | mmu-let-7e-5p | Sema4g | 26456 | ENSMUSG00000025207 | Degradome sequencing | positive | | validated |
| tarbase | MIMAT0000524 | mmu-let-7e-5p | Arih1 | 23806 | ENSMUSG00000025234 | Degradome sequencing//Degradome sequencing//Degradome sequencing//Degradome sequencing | positive | | validated |
| tarbase | MIMAT0000524 | mmu-let-7e-5p | Huwe1 | 59026 | ENSMUSG00000025261 | Degradome sequencing//Degradome sequencing//Degradome sequencing | positive | | validated |
| tarbase | MIMAT0000524 | mmu-let-7e-5p | Pfkfb1 | 18639 | ENSMUSG00000025271 | Degradome sequencing//Degradome sequencing | positive | | validated |
| tarbase | MIMAT0000524 | mmu-let-7e-5p | Dnajc14 | 74330 | ENSMUSG00000025354 | Degradome sequencing | positive | | validated |
| tarbase | MIMAT0000524 | mmu-let-7e-5p | Dgka | 13139 | ENSMUSG00000025357 | Degradome sequencing | positive | | validated |
| tarbase | MIMAT0000524 | mmu-let-7e-5p | Mbd6 | 110962 | ENSMUSG00000025409 | Degradome sequencing | positive | | validated |
| tarbase | MIMAT0000524 | mmu-let-7e-5p | Tmem192 | 73067 | ENSMUSG00000025521 | Degradome sequencing | positive | | validated |
| tarbase | MIMAT0000524 | mmu-let-7e-5p | Tnrc6c | 217351 | ENSMUSG00000025571 | Degradome sequencing | positive | | validated |
| tarbase | MIMAT0000524 | mmu-let-7e-5p | Mkln1 | 27418 | ENSMUSG00000025609 | Degradome sequencing | positive | | validated |
| tarbase | MIMAT0000524 | mmu-let-7e-5p | Bach1 | 12013 | ENSMUSG00000025612 | Degradome sequencing//Degradome sequencing//Degradome sequencing//Degradome sequencing | positive | | validated |
| tarbase | MIMAT0000524 | mmu-let-7e-5p | Shisa5 | 66940 | ENSMUSG00000025647 | Degradome sequencing | positive | | validated |
| tarbase | MIMAT0000524 | mmu-let-7e-5p | Pard3 | 93742 | ENSMUSG00000025812 | Degradome sequencing | positive | | validated |
| tarbase | MIMAT0000524 | mmu-let-7e-5p | Nrp2 | 18187 | ENSMUSG00000025969 | Degradome sequencing//Degradome sequencing//Degradome sequencing | positive | | validated |
| tarbase | MIMAT0000524 | mmu-let-7e-5p | Ctla4 | 12477 | ENSMUSG00000026011 | Degradome sequencing//Degradome sequencing | positive | | validated |
| tarbase | MIMAT0000524 | mmu-let-7e-5p | Cflar | 12633 | ENSMUSG00000026031 | Degradome sequencing//Degradome sequencing | positive | | validated |

| | | | | | | | | | |
|---|---|---|---|---|---|---|---|---|---|
| tarbase | MIMAT0000524 | mmu-let-7e-5p | Gls | 14660 | ENSMUSG00000026103 | Degradome sequencing//Degradome sequencing | positive | | validated |
| tarbase | MIMAT0000524 | mmu-let-7e-5p | Nabp1 | 109019 | ENSMUSG00000026107 | Degradome sequencing//Degradome sequencing | positive | | validated |
| tarbase | MIMAT0000524 | mmu-let-7e-5p | Sema4c | 20353 | ENSMUSG00000026121 | Degradome sequencing | positive | | validated |
| tarbase | MIMAT0000524 | mmu-let-7e-5p | Dst | 13518 | ENSMUSG00000026131 | Degradome sequencing//Degradome sequencing//Degradome sequencing | positive | | validated |
| tarbase | MIMAT0000524 | mmu-let-7e-5p | Fam135a | 68187 | ENSMUSG00000026153 | Degradome sequencing | positive | | validated |
| tarbase | MIMAT0000524 | mmu-let-7e-5p | Pnkd | 56695 | ENSMUSG00000026179 | Degradome sequencing | positive | | validated |
| tarbase | MIMAT0000524 | mmu-let-7e-5p | Tuba4a | 22145 | ENSMUSG00000026202 | Degradome sequencing//Degradome sequencing | positive | | validated |
| tarbase | MIMAT0000524 | mmu-let-7e-5p | Trip12 | 14897 | ENSMUSG00000026219 | Degradome sequencing | positive | | validated |
| tarbase | MIMAT0000524 | mmu-let-7e-5p | Itm2c | | ENSMUSG00000026223 | Degradome sequencing | positive | | validated |
| tarbase | MIMAT0000524 | mmu-let-7e-5p | Atg16l1 | 77040 | ENSMUSG00000026289 | Degradome sequencing//Degradome sequencing | positive | | validated |
| tarbase | MIMAT0000524 | mmu-let-7e-5p | Lrrfip1 | 16978 | ENSMUSG00000026305 | Degradome sequencing//Degradome sequencing | positive | | validated |
| tarbase | MIMAT0000524 | mmu-let-7e-5p | Ubxn4 | 67812 | ENSMUSG00000026353 | Degradome sequencing//Degradome sequencing | positive | | validated |
| tarbase | MIMAT0000524 | mmu-let-7e-5p | Ptpn4 | 19258 | ENSMUSG00000026384 | Degradome sequencing | positive | | validated |
| tarbase | MIMAT0000524 | mmu-let-7e-5p | Ptprc | 19264 | ENSMUSG00000026395 | Degradome sequencing//Degradome sequencing | positive | | validated |
| tarbase | MIMAT0000524 | mmu-let-7e-5p | Nr5a2 | 26424 | ENSMUSG00000026398 | Degradome sequencing | positive | | validated |
| tarbase | MIMAT0000524 | mmu-let-7e-5p | Pfkfb2 | 18640 | ENSMUSG00000026409 | Degradome sequencing//Degradome sequencing//Degradome sequencing | positive | | validated |
| tarbase | MIMAT0000524 | mmu-let-7e-5p | Srgap2 | 14270 | ENSMUSG00000026425 | Degradome sequencing | positive | | validated |
| tarbase | MIMAT0000524 | mmu-let-7e-5p | Tor1aip1 | 208263 | ENSMUSG00000026466 | Degradome sequencing//Degradome sequencing | positive | | validated |
| tarbase | MIMAT0000524 | mmu-let-7e-5p | Xpr1 | 19775 | ENSMUSG00000026469 | Degradome sequencing | positive | | validated |
| tarbase | MIMAT0000524 | mmu-let-7e-5p | Rgs16 | 19734 | ENSMUSG00000026475 | Degradome sequencing//Degradome sequencing | positive | | validated |
| tarbase | MIMAT0000524 | mmu-let-7e-5p | Ahctf1 | 226747 | ENSMUSG00000026491 | Degradome sequencing//Degradome sequencing//Degradome sequencing | positive | | validated |
| tarbase | MIMAT0000524 | mmu-let-7e-5p | Sec16b | 89867 | ENSMUSG00000026589 | Degradome sequencing | positive | | validated |
| tarbase | MIMAT0000524 | mmu-let-7e-5p | Abl2 | 11352 | ENSMUSG00000026596 | Degradome sequencing | positive | | validated |
| tarbase | MIMAT0000524 | mmu-let-7e-5p | Kctd3 | 226823 | ENSMUSG00000026608 | Degradome sequencing | positive | | validated |
| tarbase | MIMAT0000524 | mmu-let-7e-5p | Lpgat1 | 226856 | ENSMUSG00000026623 | Degradome sequencing//Degradome sequencing//Degradome sequencing | positive | | validated |
| tarbase | MIMAT0000524 | mmu-let-7e-5p | Plxna2 | 18845 | ENSMUSG00000026640 | Degradome sequencing | positive | | validated |
| tarbase | MIMAT0000524 | mmu-let-7e-5p | Uhmk1 | 16589 | ENSMUSG00000026667 | Degradome sequencing | positive | | validated |
| tarbase | MIMAT0000524 | mmu-let-7e-5p | Uap1 | 107652 | ENSMUSG00000026670 | Degradome sequencing | positive | | validated |
| tarbase | MIMAT0000524 | mmu-let-7e-5p | Pigc | 67292 | ENSMUSG00000026698 | Degradome sequencing | positive | | validated |
| tarbase | MIMAT0000524 | mmu-let-7e-5p | Rabgap1 | 29809 | ENSMUSG00000026721 | Degradome sequencing | positive | | validated |
| tarbase | MIMAT0000524 | mmu-let-7e-5p | Mllt10 | 17354 | ENSMUSG00000026743 | Degradome sequencing//Degradome sequencing | positive | | validated |
| tarbase | MIMAT0000524 | mmu-let-7e-5p | Eng | 13805 | ENSMUSG00000026814 | Degradome sequencing | positive | | validated |
| tarbase | MIMAT0000524 | mmu-let-7e-5p | Crat | 12908 | ENSMUSG00000026853 | Degradome sequencing | positive | | validated |
| tarbase | MIMAT0000524 | mmu-let-7e-5p | Gapvd1 | 66691 | ENSMUSG00000026867 | Degradome sequencing//Degradome sequencing//Degradome sequencing | positive | | validated |
| tarbase | MIMAT0000524 | mmu-let-7e-5p | Hc | 15139 | ENSMUSG00000026874 | Degradome sequencing | positive | | validated |

| | | | | | | | | | |
|---|---|---|---|---|---|---|---|---|---|
| tarbase | MIMAT0000524 | mmu-let-7e-5p | Brd3 | 67382 | ENSMUSG00000026918 | Degradome sequencing | positive | | validated |
| tarbase | MIMAT0000524 | mmu-let-7e-5p | Sec16a | 227648 | ENSMUSG00000026924 | Degradome sequencing//Degradome sequencing//Degradome sequencing//Degradome sequencing | positive | | validated |
| tarbase | MIMAT0000524 | mmu-let-7e-5p | Pmpca | 66865 | ENSMUSG00000026926 | Degradome sequencing | positive | | validated |
| tarbase | MIMAT0000524 | mmu-let-7e-5p | Abca2 | 11305 | ENSMUSG00000026944 | Degradome sequencing//Degradome sequencing//Degradome sequencing//Degradome sequencing | positive | | validated |
| tarbase | MIMAT0000524 | mmu-let-7e-5p | Rbms1 | 56878 | ENSMUSG00000026970 | Degradome sequencing//Degradome sequencing//Degradome sequencing//Degradome sequencing//Degradome sequencing//Degradome sequencing | positive | | validated |
| tarbase | MIMAT0000524 | mmu-let-7e-5p | Mar-07 | | ENSMUSG00000026977 | Degradome sequencing//Degradome sequencing//Degradome sequencing//Degradome sequencing//Degradome sequencing | positive | | validated |
| tarbase | MIMAT0000524 | mmu-let-7e-5p | Psd4 | 215632 | ENSMUSG00000026979 | Degradome sequencing | positive | | validated |
| tarbase | MIMAT0000524 | mmu-let-7e-5p | Abcb11 | 27413 | ENSMUSG00000027048 | Degradome sequencing | positive | | validated |
| tarbase | MIMAT0000524 | mmu-let-7e-5p | Clp1 | 98985 | ENSMUSG00000027079 | Degradome sequencing | positive | | validated |
| tarbase | MIMAT0000524 | mmu-let-7e-5p | Katnbl1 | 72425 | ENSMUSG00000027132 | Degradome sequencing | positive | | validated |
| tarbase | MIMAT0000524 | mmu-let-7e-5p | Nop10 | | ENSMUSG00000027133 | Degradome sequencing//Degradome sequencing | positive | | validated |
| tarbase | MIMAT0000524 | mmu-let-7e-5p | Hipk3 | 15259 | ENSMUSG00000027177 | Degradome sequencing//Degradome sequencing | positive | | validated |
| tarbase | MIMAT0000524 | mmu-let-7e-5p | Eif3j1 | | ENSMUSG00000027236 | Degradome sequencing//Degradome sequencing | negative | | validated |
| tarbase | MIMAT0000524 | mmu-let-7e-5p | Hao1 | 15112 | ENSMUSG00000027261 | Degradome sequencing | positive | | validated |
| tarbase | MIMAT0000524 | mmu-let-7e-5p | Snap23 | 20619 | ENSMUSG00000027287 | Degradome sequencing | positive | | validated |
| tarbase | MIMAT0000524 | mmu-let-7e-5p | Ubox5 | 140629 | ENSMUSG00000027300 | Degradome sequencing | positive | | validated |
| tarbase | MIMAT0000524 | mmu-let-7e-5p | 4930402I | 228602 | ENSMUSG00000027309 | Degradome sequencing//Degradome sequencing | positive | | validated |
| tarbase | MIMAT0000524 | mmu-let-7e-5p | Rpusd2 | | ENSMUSG00000027324 | Degradome sequencing//Degradome sequencing | positive | | validated |
| tarbase | MIMAT0000524 | mmu-let-7e-5p | Gpcpd1 | 74182 | ENSMUSG00000027346 | Degradome sequencing//Degradome sequencing//Degradome sequencing//Degradome sequencing | positive | | validated |
| tarbase | MIMAT0000524 | mmu-let-7e-5p | Rasgrp1 | 19419 | ENSMUSG00000027347 | Degradome sequencing//Degradome sequencing | positive | | validated |
| tarbase | MIMAT0000524 | mmu-let-7e-5p | Spred1 | 114715 | ENSMUSG00000027351 | Degradome sequencing | positive | | validated |
| tarbase | MIMAT0000524 | mmu-let-7e-5p | Stard7 | 99138 | ENSMUSG00000027367 | Degradome sequencing | positive | | validated |
| tarbase | MIMAT0000524 | mmu-let-7e-5p | Bcl2l11 | 12125 | ENSMUSG00000027381 | Degradome sequencing//Degradome sequencing | positive | | validated |
| tarbase | MIMAT0000524 | mmu-let-7e-5p | Slc20a1 | 20515 | ENSMUSG00000027397 | Degradome sequencing//Degradome sequencing//Degradome sequencing | positive | | validated |
| tarbase | MIMAT0000524 | mmu-let-7e-5p | Snx5 | 69178 | ENSMUSG00000027423 | Degradome sequencing//Degradome sequencing//Degradome sequencing//Degradome sequencing | positive | | validated |

| | | | | | | | | | |
|---|---|---|---|---|---|---|---|---|---|
| tarbase | MIMAT0000524 | mmu-let-7e-5p | Pag1 | 94212 | ENSMUSG00000027508 | Degradome sequencing//Degradome sequencing//Degradome sequencing//Degradome sequencing//Degradome sequencing | positive | | validated |
| tarbase | MIMAT0000524 | mmu-let-7e-5p | Gid8 | 76425 | ENSMUSG00000027573 | Degradome sequencing//Degradome sequencing//Degradome sequencing | positive | | validated |
| tarbase | MIMAT0000524 | mmu-let-7e-5p | Helz2 | 229003 | ENSMUSG00000027580 | Degradome sequencing//Degradome sequencing | positive | | validated |
| tarbase | MIMAT0000524 | mmu-let-7e-5p | Rpn2 | 20014 | ENSMUSG00000027642 | Degradome sequencing | positive | | validated |
| tarbase | MIMAT0000524 | mmu-let-7e-5p | Tti1 | 75425 | ENSMUSG00000027650 | Degradome sequencing | positive | | validated |
| tarbase | MIMAT0000524 | mmu-let-7e-5p | Skil | 20482 | ENSMUSG00000027660 | Degradome sequencing//Degradome sequencing | positive | | validated |
| tarbase | MIMAT0000524 | mmu-let-7e-5p | Pik3ca | 18706 | ENSMUSG00000027665 | Degradome sequencing | positive | | validated |
| tarbase | MIMAT0000524 | mmu-let-7e-5p | Ttc14 | 67120 | ENSMUSG00000027677 | Degradome sequencing//Degradome sequencing | positive | | validated |
| tarbase | MIMAT0000524 | mmu-let-7e-5p | Ncoa3 | 17979 | ENSMUSG00000027678 | Degradome sequencing | positive | | validated |
| tarbase | MIMAT0000524 | mmu-let-7e-5p | Sec62 | 69276 | ENSMUSG00000027706 | Degradome sequencing | positive | | validated |
| tarbase | MIMAT0000524 | mmu-let-7e-5p | Ccna2 | 12428 | ENSMUSG00000027715 | Degradome sequencing//Degradome sequencing//Degradome sequencing | positive | | validated |
| tarbase | MIMAT0000524 | mmu-let-7e-5p | Kpna4 | 16649 | ENSMUSG00000027782 | Degradome sequencing//Degradome sequencing | positive | | validated |
| tarbase | MIMAT0000524 | mmu-let-7e-5p | Tsc22d2 | 72033 | ENSMUSG00000027806 | Degradome sequencing//Degradome sequencing | positive | | validated |
| tarbase | MIMAT0000524 | mmu-let-7e-5p | Nras | 18176 | ENSMUSG00000027852 | Degradome sequencing//Degradome sequencing | positive | | validated |
| tarbase | MIMAT0000524 | mmu-let-7e-5p | Notch2 | 18129 | ENSMUSG00000027878 | Degradome sequencing//Degradome sequencing | positive | | validated |
| tarbase | MIMAT0000524 | mmu-let-7e-5p | Adar | 56417 | ENSMUSG00000027951 | Degradome sequencing//Degradome sequencing | positive | | validated |
| tarbase | MIMAT0000524 | mmu-let-7e-5p | Ash1l | 192195 | ENSMUSG00000028053 | Degradome sequencing//Degradome sequencing | positive | | validated |
| tarbase | MIMAT0000524 | mmu-let-7e-5p | Cd1d1 | 12479 | ENSMUSG00000028076 | Degradome sequencing | positive | | validated |
| tarbase | MIMAT0000524 | mmu-let-7e-5p | Lrba | 80877 | ENSMUSG00000028080 | Degradome sequencing | positive | | validated |
| tarbase | MIMAT0000524 | mmu-let-7e-5p | Rnf115 | 67845 | ENSMUSG00000028098 | Degradome sequencing//Degradome sequencing | positive | | validated |
| tarbase | MIMAT0000524 | mmu-let-7e-5p | Bcar3 | 29815 | ENSMUSG00000028121 | Degradome sequencing | positive | | validated |
| tarbase | MIMAT0000524 | mmu-let-7e-5p | Tmem56 | 99887 | ENSMUSG00000028132 | Degradome sequencing//Degradome sequencing | positive | | validated |
| tarbase | MIMAT0000524 | mmu-let-7e-5p | Tspan5 | 56224 | ENSMUSG00000028152 | Degradome sequencing | positive | | validated |
| tarbase | MIMAT0000524 | mmu-let-7e-5p | Rpf1 | 70285 | ENSMUSG00000028187 | Degradome sequencing | positive | | validated |
| tarbase | MIMAT0000524 | mmu-let-7e-5p | Trp53inp | 60599 | ENSMUSG00000028211 | Degradome sequencing//Degradome sequencing//Degradome sequencing | positive | | validated |
| tarbase | MIMAT0000524 | mmu-let-7e-5p | Ccne2 | 12448 | ENSMUSG00000028212 | Degradome sequencing | positive | | validated |
| tarbase | MIMAT0000524 | mmu-let-7e-5p | Tgs1 | 116940 | ENSMUSG00000028233 | Degradome sequencing//Degradome sequencing | positive | | validated |
| tarbase | MIMAT0000524 | mmu-let-7e-5p | Ndufaf4 | 68493 | ENSMUSG00000028261 | Degradome sequencing | positive | | validated |
| tarbase | MIMAT0000524 | mmu-let-7e-5p | Gbp2 | 14469 | ENSMUSG00000028270 | Degradome sequencing//Degradome sequencing | positive | | validated |
| tarbase | MIMAT0000524 | mmu-let-7e-5p | Tnfsf8 | 21949 | ENSMUSG00000028362 | Degradome sequencing | positive | | validated |

| | | | | | | | | | |
|---|---|---|---|---|---|---|---|---|---|
| tarbase | MIMAT0000524 | mmu-let-7e-5p | Ugcg | 22234 | ENSMUSG00000028381 | Degradome sequencing//Degradome sequencing//Degradome sequencing | positive | | validated |
| tarbase | MIMAT0000524 | mmu-let-7e-5p | Ptbp3 | 230257 | ENSMUSG00000028382 | Degradome sequencing | positive | | validated |
| tarbase | MIMAT0000524 | mmu-let-7e-5p | Ptprd | 19266 | ENSMUSG00000028399 | Degradome sequencing//Degradome sequencing//Degradome sequencing//Degradome sequencing | positive | | validated |
| tarbase | MIMAT0000524 | mmu-let-7e-5p | Zdhhc21 | 68268 | ENSMUSG00000028403 | Degradome sequencing | positive | | validated |
| tarbase | MIMAT0000524 | mmu-let-7e-5p | Rad23b | 19359 | ENSMUSG00000028426 | Degradome sequencing//Degradome sequencing//Degradome sequencing | positive | | validated |
| tarbase | MIMAT0000524 | mmu-let-7e-5p | Elp1 | 230233 | ENSMUSG00000028431 | Degradome sequencing//Degradome sequencing//Degradome sequencing//Degradome sequencing | positive | | validated |
| tarbase | MIMAT0000524 | mmu-let-7e-5p | Ubap2 | 68926 | ENSMUSG00000028433 | Degradome sequencing//Degradome sequencing//Degradome sequencing | positive | | validated |
| tarbase | MIMAT0000524 | mmu-let-7e-5p | Ubap1 | 67123 | ENSMUSG00000028437 | Degradome sequencing | positive | | validated |
| tarbase | MIMAT0000524 | mmu-let-7e-5p | Rgp1 | 242406 | ENSMUSG00000028468 | Degradome sequencing//Degradome sequencing//Degradome sequencing//Degradome sequencing | positive | | validated |
| tarbase | MIMAT0000524 | mmu-let-7e-5p | Usp24 | 329908 | ENSMUSG00000028514 | Degradome sequencing | positive | | validated |
| tarbase | MIMAT0000524 | mmu-let-7e-5p | Plpp3 | 67916 | ENSMUSG00000028517 | Degradome sequencing | positive | | validated |
| tarbase | MIMAT0000524 | mmu-let-7e-5p | Prkaa2 | 108079 | ENSMUSG00000028518 | Degradome sequencing | positive | | validated |
| tarbase | MIMAT0000524 | mmu-let-7e-5p | Ak4 | 11639 | ENSMUSG00000028527 | Degradome sequencing | positive | | validated |
| tarbase | MIMAT0000524 | mmu-let-7e-5p | Jak1 | 16451 | ENSMUSG00000028530 | Degradome sequencing//Degradome sequencing | positive | | validated |
| tarbase | MIMAT0000524 | mmu-let-7e-5p | Tnfrsf1b | | ENSMUSG00000028599 | Degradome sequencing//Degradome sequencing//Degradome sequencing | positive | | validated |
| tarbase | MIMAT0000524 | mmu-let-7e-5p | Macf1 | 11426 | ENSMUSG00000028649 | Degradome sequencing//Degradome sequencing//Degradome sequencing//Degradome sequencing//Degradome sequencing | positive | | validated |
| tarbase | MIMAT0000524 | mmu-let-7e-5p | Eloa | 27224 | ENSMUSG00000028668 | Degradome sequencing | positive | | validated |
| tarbase | MIMAT0000524 | mmu-let-7e-5p | Nsun4 | 72181 | ENSMUSG00000028706 | Degradome sequencing | positive | | validated |
| tarbase | MIMAT0000524 | mmu-let-7e-5p | Pqlc2 | 212555 | ENSMUSG00000028744 | Degradome sequencing//Degradome sequencing//Degradome sequencing | positive | | validated |
| tarbase | MIMAT0000524 | mmu-let-7e-5p | Ddost | 13200 | ENSMUSG00000028757 | Degradome sequencing//Degradome sequencing | positive | | validated |
| tarbase | MIMAT0000524 | mmu-let-7e-5p | Ak2 | 11637 | ENSMUSG00000028792 | Degradome sequencing//Degradome sequencing | positive | | validated |
| tarbase | MIMAT0000524 | mmu-let-7e-5p | Psmb2 | 26445 | ENSMUSG00000028837 | Degradome sequencing | positive | | validated |
| tarbase | MIMAT0000524 | mmu-let-7e-5p | Gpatch3 | 242691 | ENSMUSG00000028850 | Degradome sequencing | positive | | validated |
| tarbase | MIMAT0000524 | mmu-let-7e-5p | Eya3 | 14050 | ENSMUSG00000028886 | Degradome sequencing | positive | | validated |
| tarbase | MIMAT0000524 | mmu-let-7e-5p | Per3 | 18628 | ENSMUSG00000028957 | Degradome sequencing | positive | | validated |
| tarbase | MIMAT0000524 | mmu-let-7e-5p | Ube4b | 63958 | ENSMUSG00000028960 | Degradome sequencing//Degradome sequencing | positive | | validated |
| tarbase | MIMAT0000524 | mmu-let-7e-5p | Errfi1 | 74155 | ENSMUSG00000028967 | Degradome sequencing | positive | | validated |
| tarbase | MIMAT0000524 | mmu-let-7e-5p | Slc2a5 | 56485 | ENSMUSG00000028976 | Degradome sequencing | positive | | validated |
| tarbase | MIMAT0000524 | mmu-let-7e-5p | Mtor | 56717 | ENSMUSG00000028991 | Degradome sequencing | positive | | validated |
| tarbase | MIMAT0000524 | mmu-let-7e-5p | Kmt2e | 69188 | ENSMUSG00000029004 | Degradome sequencing//Degradome sequencing | positive | | validated |

| | | | | | | | | | |
|---|---|---|---|---|---|---|---|---|---|
| tarbase | MIMAT0000524 | mmu-let-7e-5p | Nelfa | 24116 | ENSMUSG00000029111 | Degradome sequencing//Degradome sequencing | positive | | validated |
| tarbase | MIMAT0000524 | mmu-let-7e-5p | | | ENSMUSG00000029144 | Degradome sequencing | positive | | validated |
| tarbase | MIMAT0000524 | mmu-let-7e-5p | Tmem165 | 21982 | ENSMUSG00000029234 | Degradome sequencing | positive | | validated |
| tarbase | MIMAT0000524 | mmu-let-7e-5p | Tgfbr3 | 21814 | ENSMUSG00000029287 | Degradome sequencing//Degradome sequencing//Degradome sequencing | positive | | validated |
| tarbase | MIMAT0000524 | mmu-let-7e-5p | Slc10a6 | 75750 | ENSMUSG00000029321 | Degradome sequencing | positive | | validated |
| tarbase | MIMAT0000524 | mmu-let-7e-5p | Abcb9 | 56325 | ENSMUSG00000029408 | Degradome sequencing | positive | | validated |
| tarbase | MIMAT0000524 | mmu-let-7e-5p | Scarb2 | 12492 | ENSMUSG00000029426 | Degradome sequencing//Degradome sequencing | positive | | validated |
| tarbase | MIMAT0000524 | mmu-let-7e-5p | Mapkapk | 17165 | ENSMUSG00000029454 | Degradome sequencing | positive | | validated |
| tarbase | MIMAT0000524 | mmu-let-7e-5p | Kdm2b | 30841 | ENSMUSG00000029475 | Degradome sequencing//Degradome sequencing | positive | | validated |
| tarbase | MIMAT0000524 | mmu-let-7e-5p | Ncor2 | 20602 | ENSMUSG00000029478 | Degradome sequencing | positive | | validated |
| tarbase | MIMAT0000524 | mmu-let-7e-5p | Ints1 | 68510 | ENSMUSG00000029547 | Degradome sequencing | positive | | validated |
| tarbase | MIMAT0000524 | mmu-let-7e-5p | Ddx54 | 71990 | ENSMUSG00000029599 | Degradome sequencing | positive | | validated |
| tarbase | MIMAT0000524 | mmu-let-7e-5p | Cdk8 | 264064 | ENSMUSG00000029635 | Degradome sequencing//Degradome sequencing | positive | | validated |
| tarbase | MIMAT0000524 | mmu-let-7e-5p | Polr1d | 20018 | ENSMUSG00000029642 | Degradome sequencing | positive | | validated |
| tarbase | MIMAT0000524 | mmu-let-7e-5p | Hsph1 | 15505 | ENSMUSG00000029657 | Degradome sequencing | positive | | validated |
| tarbase | MIMAT0000524 | mmu-let-7e-5p | Col1a2 | 12843 | ENSMUSG00000029661 | Degradome sequencing | positive | | validated |
| tarbase | MIMAT0000524 | mmu-let-7e-5p | Wasl | 73178 | ENSMUSG00000029684 | Degradome sequencing//Degradome sequencing | positive | | validated |
| tarbase | MIMAT0000524 | mmu-let-7e-5p | Lrwd1 | | ENSMUSG00000029703 | Degradome sequencing//Degradome sequencing//Degradome sequencing//Degradome sequencing | positive | | validated |
| tarbase | MIMAT0000524 | mmu-let-7e-5p | Gigyf1 | 57330 | ENSMUSG00000029714 | Degradome sequencing//Degradome sequencing | positive | | validated |
| tarbase | MIMAT0000524 | mmu-let-7e-5p | Cald1 | 109624 | ENSMUSG00000029761 | Degradome sequencing//Degradome sequencing//Degradome sequencing | positive | | validated |
| tarbase | MIMAT0000524 | mmu-let-7e-5p | Gars | 353172 | ENSMUSG00000029777 | Degradome sequencing//Degradome sequencing | positive | | validated |
| tarbase | MIMAT0000524 | mmu-let-7e-5p | Smarcad | 13990 | ENSMUSG00000029920 | Degradome sequencing | positive | | validated |
| tarbase | MIMAT0000524 | mmu-let-7e-5p | Pcyox1 | 66881 | ENSMUSG00000029998 | Degradome sequencing//Degradome sequencing | positive | | validated |
| tarbase | MIMAT0000524 | mmu-let-7e-5p | Fbxl14 | | ENSMUSG00000030019 | Degradome sequencing | positive | | validated |
| tarbase | MIMAT0000524 | mmu-let-7e-5p | Cnbp | 12785 | ENSMUSG00000030057 | Degradome sequencing//Degradome sequencing | positive | | validated |
| tarbase | MIMAT0000524 | mmu-let-7e-5p | Copg1 | 54161 | ENSMUSG00000030058 | Degradome sequencing | positive | | validated |
| tarbase | MIMAT0000524 | mmu-let-7e-5p | Foxp1 | 108655 | ENSMUSG00000030067 | Degradome sequencing//Degradome sequencing | positive | | validated |
| tarbase | MIMAT0000524 | mmu-let-7e-5p | Slc6a13 | 14412 | ENSMUSG00000030108 | Degradome sequencing | positive | | validated |
| tarbase | MIMAT0000524 | mmu-let-7e-5p | Plxnd1 | 67784 | ENSMUSG00000030123 | Degradome sequencing | positive | | validated |
| tarbase | MIMAT0000524 | mmu-let-7e-5p | Adipor2 | 68465 | ENSMUSG00000030168 | Degradome sequencing | positive | | validated |
| tarbase | MIMAT0000524 | mmu-let-7e-5p | Atf7ip | 54343 | ENSMUSG00000030213 | Degradome sequencing//Degradome sequencing//Degradome sequencing | positive | | validated |
| tarbase | MIMAT0000524 | mmu-let-7e-5p | Golt1b | 66964 | ENSMUSG00000030245 | Degradome sequencing//Degradome sequencing//Degradome sequencing | positive | | validated |
| tarbase | MIMAT0000524 | mmu-let-7e-5p | Etnk1 | 75320 | ENSMUSG00000030275 | Degradome sequencing//Degradome sequencing | positive | | validated |

| | | | | | | | | | |
|---|---|---|---|---|---|---|---|---|---|
| tarbase | MIMAT0000524 | mmu-let-7e-5p | Cmas | 12764 | ENSMUSG00000030282 | Degradome sequencing//Degradome sequencing//Degradome sequencing | positive | | validated |
| tarbase | MIMAT0000524 | mmu-let-7e-5p | Ergic2 | 67456 | ENSMUSG00000030304 | Degradome sequencing | positive | | validated |
| tarbase | MIMAT0000524 | mmu-let-7e-5p | Tnfrsf1a | 21937 | ENSMUSG00000030341 | Degradome sequencing | positive | | validated |
| tarbase | MIMAT0000524 | mmu-let-7e-5p | Vasp | 22323 | ENSMUSG00000030403 | Degradome sequencing | positive | | validated |
| tarbase | MIMAT0000524 | mmu-let-7e-5p | Nipa2 | 93790 | ENSMUSG00000030452 | Degradome sequencing | positive | | validated |
| tarbase | MIMAT0000524 | mmu-let-7e-5p | Psd3 | 234353 | ENSMUSG00000030465 | Degradome sequencing | positive | | validated |
| tarbase | MIMAT0000524 | mmu-let-7e-5p | Ctsc | | ENSMUSG00000030560 | Degradome sequencing | positive | | validated |
| tarbase | MIMAT0000524 | mmu-let-7e-5p | Pak4 | 70584 | ENSMUSG00000030602 | Degradome sequencing//Degradome sequencing | positive | | validated |
| tarbase | MIMAT0000524 | mmu-let-7e-5p | Cyp2r1 | 244209 | ENSMUSG00000030670 | Degradome sequencing | positive | | validated |
| tarbase | MIMAT0000524 | mmu-let-7e-5p | Ppp4c | 56420 | ENSMUSG00000030697 | Degradome sequencing//Degradome sequencing//Degradome sequencing | positive | | validated |
| tarbase | MIMAT0000524 | mmu-let-7e-5p | Lipt2 | | ENSMUSG00000030725 | Degradome sequencing//Degradome sequencing//Degradome sequencing | positive | | validated |
| tarbase | MIMAT0000524 | mmu-let-7e-5p | Pold3 | 67967 | ENSMUSG00000030726 | Degradome sequencing//Degradome sequencing//Degradome sequencing//Degradome sequencing | positive | | validated |
| tarbase | MIMAT0000524 | mmu-let-7e-5p | Slco2b1 | 101488 | ENSMUSG00000030737 | Degradome sequencing//Degradome sequencing | positive | | validated |
| tarbase | MIMAT0000524 | mmu-let-7e-5p | Spns1 | | ENSMUSG00000030741 | Degradome sequencing | positive | | validated |
| tarbase | MIMAT0000524 | mmu-let-7e-5p | Dgat2 | 67800 | ENSMUSG00000030747 | Degradome sequencing | positive | | validated |
| tarbase | MIMAT0000524 | mmu-let-7e-5p | Itgal | 16408 | ENSMUSG00000030830 | Degradome sequencing//Degradome sequencing | positive | | validated |
| tarbase | MIMAT0000524 | mmu-let-7e-5p | Abcc6 | 27421 | ENSMUSG00000030834 | Degradome sequencing | positive | | validated |
| tarbase | MIMAT0000524 | mmu-let-7e-5p | Plk1 | 18817 | ENSMUSG00000030867 | Degradome sequencing | positive | | validated |
| tarbase | MIMAT0000524 | mmu-let-7e-5p | Abraxas2 | 109359 | ENSMUSG00000030965 | Degradome sequencing | positive | | validated |
| tarbase | MIMAT0000524 | mmu-let-7e-5p | Acsm5 | 272428 | ENSMUSG00000030972 | Degradome sequencing | positive | | validated |
| tarbase | MIMAT0000524 | mmu-let-7e-5p | Pgap2 | 233575 | ENSMUSG00000030990 | Degradome sequencing//Degradome sequencing | positive | | validated |
| tarbase | MIMAT0000524 | mmu-let-7e-5p | Otud5 | 54644 | ENSMUSG00000031154 | Degradome sequencing//Degradome sequencing | positive | | validated |
| tarbase | MIMAT0000524 | mmu-let-7e-5p | Abcd1 | 11666 | ENSMUSG00000031378 | Degradome sequencing | positive | | validated |
| tarbase | MIMAT0000524 | mmu-let-7e-5p | Piga | 18700 | ENSMUSG00000031381 | Degradome sequencing//Degradome sequencing | positive | | validated |
| tarbase | MIMAT0000524 | mmu-let-7e-5p | F10 | 14058 | ENSMUSG00000031444 | Degradome sequencing | positive | | validated |
| tarbase | MIMAT0000524 | mmu-let-7e-5p | Adrb3 | 11556 | ENSMUSG00000031489 | Degradome sequencing//Degradome sequencing | positive | | validated |
| tarbase | MIMAT0000524 | mmu-let-7e-5p | Eif4ebp1 | 13685 | ENSMUSG00000031490 | Degradome sequencing//Degradome sequencing//Degradome sequencing | positive | | validated |
| tarbase | MIMAT0000524 | mmu-let-7e-5p | Col4a2 | 12827 | ENSMUSG00000031503 | Degradome sequencing//Degradome sequencing//Degradome sequencing//Degradome sequencing | positive | | validated |
| tarbase | MIMAT0000524 | mmu-let-7e-5p | Dusp4 | | ENSMUSG00000031530 | Degradome sequencing//Degradome sequencing | positive | | validated |
| tarbase | MIMAT0000524 | mmu-let-7e-5p | Kat6a | 244349 | ENSMUSG00000031540 | Degradome sequencing | positive | | validated |
| tarbase | MIMAT0000524 | mmu-let-7e-5p | Gpat4 | 102247 | ENSMUSG00000031545 | Degradome sequencing//Degradome sequencing//Degradome sequencing | positive | | validated |
| tarbase | MIMAT0000524 | mmu-let-7e-5p | Plpp5 | 71910 | ENSMUSG00000031570 | Degradome sequencing | positive | | validated |
| tarbase | MIMAT0000524 | mmu-let-7e-5p | Klhl2 | 77113 | ENSMUSG00000031605 | Degradome sequencing | positive | | validated |

| | | | | | | | | | |
|---|---|---|---|---|---|---|---|---|---|
| tarbase | MIMAT0000524 | mmu-let-7e-5p | Casp3 | 12367 | ENSMUSG00000031628 | Degradome sequencing//Degradome sequencing | positive | | validated |
| tarbase | MIMAT0000524 | mmu-let-7e-5p | Slc25a4 | 11739 | ENSMUSG00000031633 | Degradome sequencing | positive | | validated |
| tarbase | MIMAT0000524 | mmu-let-7e-5p | Sh3rf1 | 59009 | ENSMUSG00000031642 | Degradome sequencing | positive | | validated |
| tarbase | MIMAT0000524 | mmu-let-7e-5p | N4bp1 | 80750 | ENSMUSG00000031652 | Degradome sequencing | positive | | validated |
| tarbase | MIMAT0000524 | mmu-let-7e-5p | Sall1 | 58198 | ENSMUSG00000031665 | Degradome sequencing | positive | | validated |
| tarbase | MIMAT0000524 | mmu-let-7e-5p | Dnaja2 | 56445 | ENSMUSG00000031701 | Degradome sequencing | positive | | validated |
| tarbase | MIMAT0000524 | mmu-let-7e-5p | Gab1 | 14388 | ENSMUSG00000031714 | Degradome sequencing | positive | | validated |
| tarbase | MIMAT0000524 | mmu-let-7e-5p | Usp10 | 22224 | ENSMUSG00000031826 | Degradome sequencing//Degradome sequencing | positive | | validated |
| tarbase | MIMAT0000524 | mmu-let-7e-5p | Hsbp1 | 68196 | ENSMUSG00000031839 | Degradome sequencing//Degradome sequencing//Degradome sequencing | positive | | validated |
| tarbase | MIMAT0000524 | mmu-let-7e-5p | Hsd17b2 | 15486 | ENSMUSG00000031844 | Degradome sequencing//Degradome sequencing//Degradome sequencing//Degradome sequencing | positive | | validated |
| tarbase | MIMAT0000524 | mmu-let-7e-5p | Nfatc3 | 18021 | ENSMUSG00000031902 | Degradome sequencing//Degradome sequencing//Degradome sequencing//Degradome sequencing | positive | | validated |
| tarbase | MIMAT0000524 | mmu-let-7e-5p | Pla2g15 | 192654 | ENSMUSG00000031903 | Degradome sequencing | positive | | validated |
| tarbase | MIMAT0000524 | mmu-let-7e-5p | Ankrd49 | 56503 | ENSMUSG00000031931 | Degradome sequencing//Degradome sequencing | positive | | validated |
| tarbase | MIMAT0000524 | mmu-let-7e-5p | Kars | 85305 | ENSMUSG00000031948 | Degradome sequencing//Degradome sequencing | positive | | validated |
| tarbase | MIMAT0000524 | mmu-let-7e-5p | Aars | 234734 | ENSMUSG00000031960 | Degradome sequencing | positive | | validated |
| tarbase | MIMAT0000524 | mmu-let-7e-5p | Abcb10 | 56199 | ENSMUSG00000031974 | Degradome sequencing//Degradome sequencing | positive | | validated |
| tarbase | MIMAT0000524 | mmu-let-7e-5p | Urb2 | 382038 | ENSMUSG00000031976 | Degradome sequencing | positive | | validated |
| tarbase | MIMAT0000524 | mmu-let-7e-5p | Vps26b | | ENSMUSG00000031988 | Degradome sequencing//Degradome sequencing | positive | | validated |
| tarbase | MIMAT0000524 | mmu-let-7e-5p | Oaf | 102644 | ENSMUSG00000032014 | Degradome sequencing//Degradome sequencing//Degradome sequencing | positive | | validated |
| tarbase | MIMAT0000524 | mmu-let-7e-5p | Sc5d | 235293 | ENSMUSG00000032018 | Degradome sequencing | positive | | validated |
| tarbase | MIMAT0000524 | mmu-let-7e-5p | Crtam | | ENSMUSG00000032021 | Degradome sequencing | positive | | validated |
| tarbase | MIMAT0000524 | mmu-let-7e-5p | Acat1 | 110446 | ENSMUSG00000032047 | Degradome sequencing//Degradome sequencing | positive | | validated |
| tarbase | MIMAT0000524 | mmu-let-7e-5p | Rdx | 19684 | ENSMUSG00000032050 | Degradome sequencing//Degradome sequencing//Degradome sequencing//Degradome sequencing//Degradome sequencing | positive | | validated |
| tarbase | MIMAT0000524 | mmu-let-7e-5p | Apoa5 | 66113 | ENSMUSG00000032079 | Degradome sequencing//Degradome sequencing//Degradome sequencing | positive | | validated |
| tarbase | MIMAT0000524 | mmu-let-7e-5p | Cd3e | 12501 | ENSMUSG00000032093 | Degradome sequencing//Degradome sequencing | positive | | validated |
| tarbase | MIMAT0000524 | mmu-let-7e-5p | Slc37a4 | 14385 | ENSMUSG00000032114 | Degradome sequencing | positive | | validated |
| tarbase | MIMAT0000524 | mmu-let-7e-5p | Tyk2 | 54721 | ENSMUSG00000032175 | Degradome sequencing//Degradome sequencing | positive | | validated |
| tarbase | MIMAT0000524 | mmu-let-7e-5p | Smarca4 | 20586 | ENSMUSG00000032187 | Degradome sequencing//Degradome sequencing//Degradome sequencing | positive | | validated |
| tarbase | MIMAT0000524 | mmu-let-7e-5p | Ldlr | 16835 | ENSMUSG00000032193 | Degradome sequencing | positive | | validated |
| tarbase | MIMAT0000524 | mmu-let-7e-5p | Glce | 93683 | ENSMUSG00000032252 | Degradome sequencing | positive | | validated |

| | | | | | | | | | |
|---|---|---|---|---|---|---|---|---|---|
| tarbase | MIMAT0000524 | mmu-let-7e-5p | 1700017I | 74211 | ENSMUSG00000032300 | Degradome sequencing//Degradome sequencing//Degradome sequencing//Degradome sequencing | positive | | validated |
| tarbase | MIMAT0000524 | mmu-let-7e-5p | Tmem30a | 69981 | ENSMUSG00000032328 | Degradome sequencing//Degradome sequencing//Degradome sequencing//Degradome sequencing//Degradome sequencing | positive | | validated |
| tarbase | MIMAT0000524 | mmu-let-7e-5p | Ciao2a | 68250 | ENSMUSG00000032381 | Degradome sequencing//Degradome sequencing//Degradome sequencing | positive | | validated |
| tarbase | MIMAT0000524 | mmu-let-7e-5p | Parp16 | 214424 | ENSMUSG00000032392 | Degradome sequencing//Degradome sequencing | positive | | validated |
| tarbase | MIMAT0000524 | mmu-let-7e-5p | Pias1 | 56469 | ENSMUSG00000032405 | Degradome sequencing | positive | | validated |
| tarbase | MIMAT0000524 | mmu-let-7e-5p | Atr | 245000 | ENSMUSG00000032409 | Degradome sequencing//Degradome sequencing | positive | | validated |
| tarbase | MIMAT0000524 | mmu-let-7e-5p | Xrn1 | 24127 | ENSMUSG00000032410 | Degradome sequencing | positive | | validated |
| tarbase | MIMAT0000524 | mmu-let-7e-5p | Nt5e | | ENSMUSG00000032420 | Degradome sequencing | positive | | validated |
| tarbase | MIMAT0000524 | mmu-let-7e-5p | Mlh1 | 17350 | ENSMUSG00000032498 | Degradome sequencing | positive | | validated |
| tarbase | MIMAT0000524 | mmu-let-7e-5p | Trib1 | 211770 | ENSMUSG00000032501 | Degradome sequencing//Degradome sequencing//Degradome sequencing//Degradome sequencing//Degradome sequencing//Degradome sequencing | positive | | validated |
| tarbase | MIMAT0000524 | mmu-let-7e-5p | Wdr48 | 67561 | ENSMUSG00000032512 | Degradome sequencing | positive | | validated |
| tarbase | MIMAT0000524 | mmu-let-7e-5p | Vipr1 | 22354 | ENSMUSG00000032528 | Degradome sequencing//Degradome sequencing | positive | | validated |
| tarbase | MIMAT0000524 | mmu-let-7e-5p | Dnajc13 | | ENSMUSG00000032560 | Degradome sequencing | positive | | validated |
| tarbase | MIMAT0000524 | mmu-let-7e-5p | Qars | | ENSMUSG00000032604 | Degradome sequencing//Degradome sequencing//Degradome sequencing//Degradome sequencing | positive | | validated |
| tarbase | MIMAT0000524 | mmu-let-7e-5p | Atxn2l | 233871 | ENSMUSG00000032637 | Degradome sequencing//Degradome sequencing//Degradome sequencing//Degradome sequencing | positive | | validated |
| tarbase | MIMAT0000524 | mmu-let-7e-5p | Trib3 | 228775 | ENSMUSG00000032715 | Degradome sequencing | positive | | validated |
| tarbase | MIMAT0000524 | mmu-let-7e-5p | Arap1 | 69710 | ENSMUSG00000032812 | Degradome sequencing | positive | | validated |
| tarbase | MIMAT0000524 | mmu-let-7e-5p | Zswim6 | 67263 | ENSMUSG00000032846 | Degradome sequencing//Degradome sequencing | positive | | validated |
| tarbase | MIMAT0000524 | mmu-let-7e-5p | Cyb5r4 | 266690 | ENSMUSG00000032872 | Degradome sequencing//Degradome sequencing//Degradome sequencing | positive | | validated |
| tarbase | MIMAT0000524 | mmu-let-7e-5p | Mycbp2 | 105689 | ENSMUSG00000033004 | Degradome sequencing | positive | | validated |
| tarbase | MIMAT0000524 | mmu-let-7e-5p | Slc35d2 | 70484 | ENSMUSG00000033114 | Degradome sequencing//Degradome sequencing | positive | | validated |
| tarbase | MIMAT0000524 | mmu-let-7e-5p | Atg9a | 245860 | ENSMUSG00000033124 | Degradome sequencing//Degradome sequencing | positive | | validated |
| tarbase | MIMAT0000524 | mmu-let-7e-5p | Phldb2 | 208177 | ENSMUSG00000033149 | Degradome sequencing//Degradome sequencing//Degradome sequencing | positive | | validated |
| tarbase | MIMAT0000524 | mmu-let-7e-5p | Rac2 | 19354 | ENSMUSG00000033220 | Degradome sequencing//Degradome sequencing | positive | | validated |
| tarbase | MIMAT0000524 | mmu-let-7e-5p | Slc35a4 | 67843 | ENSMUSG00000033272 | Degradome sequencing//Degradome sequencing//Degradome sequencing | positive | | validated |
| tarbase | MIMAT0000524 | mmu-let-7e-5p | Ptprf | 19268 | ENSMUSG00000033295 | Degradome sequencing | positive | | validated |

| | | | | | | | | | |
|---|---|---|---|---|---|---|---|---|---|
| tarbase | MIMAT0000524 | mmu-let-7e-5p | Ctdp1 | 67655 | ENSMUSG00000033323 | Degradome sequencing | positive | | validated |
| tarbase | MIMAT0000524 | mmu-let-7e-5p | Dnm2 | 13430 | ENSMUSG00000033335 | Degradome sequencing | positive | | validated |
| tarbase | MIMAT0000524 | mmu-let-7e-5p | Map2k4 | 26398 | ENSMUSG00000033352 | Degradome sequencing | positive | | validated |
| tarbase | MIMAT0000524 | mmu-let-7e-5p | Agl | 77559 | ENSMUSG00000033400 | Degradome sequencing | positive | | validated |
| tarbase | MIMAT0000524 | mmu-let-7e-5p | Ctdspl2 | | ENSMUSG00000033411 | Degradome sequencing | positive | | validated |
| tarbase | MIMAT0000524 | mmu-let-7e-5p | Lpar6 | 67168 | ENSMUSG00000033446 | Degradome sequencing//Degradome sequencing//Degradome sequencing//Degradome sequencing//Degradome sequencing | positive | | validated |
| tarbase | MIMAT0000524 | mmu-let-7e-5p | Fam160b | 226252 | ENSMUSG00000033478 | Degradome sequencing//Degradome sequencing | positive | | validated |
| tarbase | MIMAT0000524 | mmu-let-7e-5p | Fndc3a | 319448 | ENSMUSG00000033487 | Degradome sequencing//Degradome sequencing//Degradome sequencing//Degradome sequencing | positive | | validated |
| tarbase | MIMAT0000524 | mmu-let-7e-5p | Larp4b | 217980 | ENSMUSG00000033499 | Degradome sequencing//Degradome sequencing//Degradome sequencing | positive | | validated |
| tarbase | MIMAT0000524 | mmu-let-7e-5p | Gtf2a2 | 235459 | ENSMUSG00000033543 | Degradome sequencing | positive | | validated |
| tarbase | MIMAT0000524 | mmu-let-7e-5p | Rbfox2 | 93686 | ENSMUSG00000033565 | Degradome sequencing | positive | | validated |
| tarbase | MIMAT0000524 | mmu-let-7e-5p | Pcgf3 | 69587 | ENSMUSG00000033623 | Degradome sequencing//Degradome sequencing//Degradome sequencing//Degradome sequencing//Degradome sequencing | positive | | validated |
| tarbase | MIMAT0000524 | mmu-let-7e-5p | Cep350 | 74081 | ENSMUSG00000033671 | Degradome sequencing | positive | | validated |
| tarbase | MIMAT0000524 | mmu-let-7e-5p | Ucp2 | 22228 | ENSMUSG00000033685 | Degradome sequencing//Degradome sequencing | positive | | validated |
| tarbase | MIMAT0000524 | mmu-let-7e-5p | Asb13 | | ENSMUSG00000033781 | Degradome sequencing | positive | | validated |
| tarbase | MIMAT0000524 | mmu-let-7e-5p | Atp7a | 11977 | ENSMUSG00000033792 | Degradome sequencing | positive | | validated |
| tarbase | MIMAT0000524 | mmu-let-7e-5p | Alg3 | 208624 | ENSMUSG00000033809 | Degradome sequencing | positive | | validated |
| tarbase | MIMAT0000524 | mmu-let-7e-5p | Lgals3bp | 19039 | ENSMUSG00000033880 | Degradome sequencing//Degradome sequencing//Degradome sequencing | positive | | validated |
| tarbase | MIMAT0000524 | mmu-let-7e-5p | Slc16a2 | 20502 | ENSMUSG00000033965 | Degradome sequencing | positive | | validated |
| tarbase | MIMAT0000524 | mmu-let-7e-5p | Rpap1 | 68925 | ENSMUSG00000034032 | Degradome sequencing | positive | | validated |
| tarbase | MIMAT0000524 | mmu-let-7e-5p | Phka1 | 18679 | ENSMUSG00000034055 | Degradome sequencing | positive | | validated |
| tarbase | MIMAT0000524 | mmu-let-7e-5p | Hdlbp | 110611 | ENSMUSG00000034088 | Degradome sequencing//Degradome sequencing | positive | | validated |
| tarbase | MIMAT0000524 | mmu-let-7e-5p | Pomt2 | 217734 | ENSMUSG00000034126 | Degradome sequencing | positive | | validated |
| tarbase | MIMAT0000524 | mmu-let-7e-5p | Lrrc58 | 320184 | ENSMUSG00000034158 | Degradome sequencing | positive | | validated |
| tarbase | MIMAT0000524 | mmu-let-7e-5p | Chmp7 | 105513 | ENSMUSG00000034190 | Degradome sequencing | positive | | validated |
| tarbase | MIMAT0000524 | mmu-let-7e-5p | Usp54 | 78787 | ENSMUSG00000034235 | Degradome sequencing | positive | | validated |
| tarbase | MIMAT0000524 | mmu-let-7e-5p | Setd5 | 72895 | ENSMUSG00000034269 | Degradome sequencing | positive | | validated |
| tarbase | MIMAT0000524 | mmu-let-7e-5p | Nek9 | 217718 | ENSMUSG00000034290 | Degradome sequencing | positive | | validated |
| tarbase | MIMAT0000524 | mmu-let-7e-5p | Cbl | 12402 | ENSMUSG00000034342 | Degradome sequencing | positive | | validated |
| tarbase | MIMAT0000524 | mmu-let-7e-5p | Smc4 | 70099 | ENSMUSG00000034349 | Degradome sequencing//Degradome sequencing | positive | | validated |
| tarbase | MIMAT0000524 | mmu-let-7e-5p | Wdr5b | | ENSMUSG00000034379 | Degradome sequencing | positive | | validated |
| tarbase | MIMAT0000524 | mmu-let-7e-5p | Trmt5 | 76357 | ENSMUSG00000034442 | Degradome sequencing | positive | | validated |
| tarbase | MIMAT0000524 | mmu-let-7e-5p | Eda2r | 245527 | ENSMUSG00000034457 | Degradome sequencing | positive | | validated |
| tarbase | MIMAT0000524 | mmu-let-7e-5p | Slc41a2 | 338365 | ENSMUSG00000034591 | Degradome sequencing//Degradome sequencing | positive | | validated |

| | | | | | | | | | |
|---|---|---|---|---|---|---|---|---|---|
| tarbase | MIMAT0000524 | mmu-let-7e-5p | Tut4 | 230594 | ENSMUSG00000034610 | Degradome sequencing//Degradome sequencing | positive | | validated |
| tarbase | MIMAT0000524 | mmu-let-7e-5p | Pik3ip1 | 216505 | ENSMUSG00000034614 | Degradome sequencing//Degradome sequencing | positive | | validated |
| tarbase | MIMAT0000524 | mmu-let-7e-5p | Zyg11b | 414872 | ENSMUSG00000034636 | Degradome sequencing | positive | | validated |
| tarbase | MIMAT0000524 | mmu-let-7e-5p | Bmp2k | 140780 | ENSMUSG00000034663 | Degradome sequencing | positive | | validated |
| tarbase | MIMAT0000524 | mmu-let-7e-5p | Xpot | 73192 | ENSMUSG00000034667 | Degradome sequencing//Degradome sequencing | positive | | validated |
| tarbase | MIMAT0000524 | mmu-let-7e-5p | Pbx2 | 18515 | ENSMUSG00000034673 | Degradome sequencing//Degradome sequencing//Degradome sequencing | positive | | validated |
| tarbase | MIMAT0000524 | mmu-let-7e-5p | Tmx4 | 52837 | ENSMUSG00000034723 | Degradome sequencing//Degradome sequencing//Degradome sequencing//Degradome sequencing | positive | | validated |
| tarbase | MIMAT0000524 | mmu-let-7e-5p | Cnot6l | 231464 | ENSMUSG00000034724 | Degradome sequencing//Degradome sequencing//Degradome sequencing | positive | | validated |
| tarbase | MIMAT0000524 | mmu-let-7e-5p | Map4k5 | 399510 | ENSMUSG00000034761 | Degradome sequencing//Degradome sequencing | positive | | validated |
| tarbase | MIMAT0000524 | mmu-let-7e-5p | Dusp5 | 240672 | ENSMUSG00000034765 | Degradome sequencing//Degradome sequencing | positive | | validated |
| tarbase | MIMAT0000524 | mmu-let-7e-5p | Tet3 | 194388 | ENSMUSG00000034832 | Degradome sequencing//Degradome sequencing//Degradome sequencing | positive | | validated |
| tarbase | MIMAT0000524 | mmu-let-7e-5p | Acot11 | 329910 | ENSMUSG00000034853 | Degradome sequencing | positive | | validated |
| tarbase | MIMAT0000524 | mmu-let-7e-5p | Rnf44 | 105239 | ENSMUSG00000034928 | Degradome sequencing//Degradome sequencing//Degradome sequencing | positive | | validated |
| tarbase | MIMAT0000524 | mmu-let-7e-5p | Dhx8 | 217207 | ENSMUSG00000034931 | Degradome sequencing//Degradome sequencing | positive | | validated |
| tarbase | MIMAT0000524 | mmu-let-7e-5p | Eef2 | 13629 | ENSMUSG00000034994 | Degradome sequencing | positive | | validated |
| tarbase | MIMAT0000524 | mmu-let-7e-5p | Baz1a | 217578 | ENSMUSG00000035021 | Degradome sequencing//Degradome sequencing//Degradome sequencing//Degradome sequencing//Degradome sequencing | positive | | validated |
| tarbase | MIMAT0000524 | mmu-let-7e-5p | Arhgap5 | 11855 | ENSMUSG00000035133 | Degradome sequencing | positive | | validated |
| tarbase | MIMAT0000524 | mmu-let-7e-5p | Ap2b1 | 71770 | ENSMUSG00000035152 | Degradome sequencing | positive | | validated |
| tarbase | MIMAT0000524 | mmu-let-7e-5p | Hectd1 | 207304 | ENSMUSG00000035247 | Degradome sequencing | positive | | validated |
| tarbase | MIMAT0000524 | mmu-let-7e-5p | Adrb1 | 11554 | ENSMUSG00000035283 | Degradome sequencing | positive | | validated |
| tarbase | MIMAT0000524 | mmu-let-7e-5p | Galm | 319625 | ENSMUSG00000035473 | Degradome sequencing//Degradome sequencing//Degradome sequencing | positive | | validated |
| tarbase | MIMAT0000524 | mmu-let-7e-5p | Rnf38 | 73469 | ENSMUSG00000035696 | Degradome sequencing//Degradome sequencing | positive | | validated |
| tarbase | MIMAT0000524 | mmu-let-7e-5p | Syt1 | 20979 | ENSMUSG00000035864 | Degradome sequencing | positive | | validated |
| tarbase | MIMAT0000524 | mmu-let-7e-5p | Hykk | 235386 | ENSMUSG00000035878 | Degradome sequencing | positive | | validated |
| tarbase | MIMAT0000524 | mmu-let-7e-5p | Uba6 | | ENSMUSG00000035898 | Degradome sequencing//Degradome sequencing | positive | | validated |
| tarbase | MIMAT0000524 | mmu-let-7e-5p | Fnip1 | 216742 | ENSMUSG00000035992 | Degradome sequencing//Degradome sequencing//Degradome sequencing//Degradome sequencing | positive | | validated |
| tarbase | MIMAT0000524 | mmu-let-7e-5p | Ripor2 | 193385 | ENSMUSG00000036006 | Degradome sequencing | positive | | validated |
| tarbase | MIMAT0000524 | mmu-let-7e-5p | Smug1 | 71726 | ENSMUSG00000036061 | Degradome sequencing//Degradome sequencing//Degradome sequencing | positive | | validated |

| | | | | | | | | | |
|---|---|---|---|---|---|---|---|---|---|
| tarbase | MIMAT0000524 | mmu-let-7e-5p | Sigmar1 | 18391 | ENSMUSG00000036078 | Degradome sequencing//Degradome sequencing | positive | | validated |
| tarbase | MIMAT0000524 | mmu-let-7e-5p | Slc17a3 | 105355 | ENSMUSG00000036083 | Degradome sequencing | positive | | validated |
| tarbase | MIMAT0000524 | mmu-let-7e-5p | Arl5a | 75423 | ENSMUSG00000036093 | Degradome sequencing//Degradome sequencing | positive | | validated |
| tarbase | MIMAT0000524 | mmu-let-7e-5p | Slf2 | 226151 | ENSMUSG00000036097 | Degradome sequencing//Degradome sequencing//Degradome sequencing//Degradome sequencing | positive | | validated |
| tarbase | MIMAT0000524 | mmu-let-7e-5p | Vezt | 215008 | ENSMUSG00000036099 | Degradome sequencing | positive | | validated |
| tarbase | MIMAT0000524 | mmu-let-7e-5p | Gxylt1 | 223827 | ENSMUSG00000036197 | Degradome sequencing//Degradome sequencing | positive | | validated |
| tarbase | MIMAT0000524 | mmu-let-7e-5p | Naa30 | 70646 | ENSMUSG00000036282 | Degradome sequencing | positive | | validated |
| tarbase | MIMAT0000524 | mmu-let-7e-5p | Gramd1c | 207798 | ENSMUSG00000036292 | Degradome sequencing//Degradome sequencing//Degradome sequencing//Degradome sequencing | positive | | validated |
| tarbase | MIMAT0000524 | mmu-let-7e-5p | Hif1an | 319594 | ENSMUSG00000036450 | Degradome sequencing//Degradome sequencing//Degradome sequencing//Degradome sequencing | positive | | validated |
| tarbase | MIMAT0000524 | mmu-let-7e-5p | Cnot1 | 234594 | ENSMUSG00000036550 | Degradome sequencing | positive | | validated |
| tarbase | MIMAT0000524 | mmu-let-7e-5p | Plxnb2 | 140570 | ENSMUSG00000036606 | Degradome sequencing | positive | | validated |
| tarbase | MIMAT0000524 | mmu-let-7e-5p | Clcn7 | 26373 | ENSMUSG00000036636 | Degradome sequencing | positive | | validated |
| tarbase | MIMAT0000524 | mmu-let-7e-5p | Ago2 | | ENSMUSG00000036698 | Degradome sequencing//Degradome sequencing//Degradome sequencing | positive | | validated |
| tarbase | MIMAT0000524 | mmu-let-7e-5p | Oxsr1 | 108737 | ENSMUSG00000036737 | Degradome sequencing//Degradome sequencing//Degradome sequencing | positive | | validated |
| tarbase | MIMAT0000524 | mmu-let-7e-5p | Jmjd4 | 194952 | ENSMUSG00000036819 | Degradome sequencing | positive | | validated |
| tarbase | MIMAT0000524 | mmu-let-7e-5p | Ehmt1 | 77683 | ENSMUSG00000036893 | Degradome sequencing | positive | | validated |
| tarbase | MIMAT0000524 | mmu-let-7e-5p | C1qb | 12260 | ENSMUSG00000036905 | Degradome sequencing | positive | | validated |
| tarbase | MIMAT0000524 | mmu-let-7e-5p | Aifm1 | 26926 | ENSMUSG00000036932 | Degradome sequencing//Degradome sequencing | positive | | validated |
| tarbase | MIMAT0000524 | mmu-let-7e-5p | Rab8b | 235442 | ENSMUSG00000036943 | Degradome sequencing//Degradome sequencing | positive | | validated |
| tarbase | MIMAT0000524 | mmu-let-7e-5p | Sh3glb1 | 54673 | ENSMUSG00000037062 | Degradome sequencing | positive | | validated |
| tarbase | MIMAT0000524 | mmu-let-7e-5p | Scd1 | 20249 | ENSMUSG00000037071 | Degradome sequencing//Degradome sequencing | positive | | validated |
| tarbase | MIMAT0000524 | mmu-let-7e-5p | Slc35b2 | 73836 | ENSMUSG00000037089 | Degradome sequencing | positive | | validated |
| tarbase | MIMAT0000524 | mmu-let-7e-5p | Ralgapa2 | 241694 | ENSMUSG00000037110 | Degradome sequencing//Degradome sequencing | positive | | validated |
| tarbase | MIMAT0000524 | mmu-let-7e-5p | E330009 | 243780 | ENSMUSG00000037172 | Degradome sequencing | positive | | validated |
| tarbase | MIMAT0000524 | mmu-let-7e-5p | Spry1 | 24063 | ENSMUSG00000037211 | Degradome sequencing | positive | | validated |
| tarbase | MIMAT0000524 | mmu-let-7e-5p | Mospd3 | 68929 | ENSMUSG00000037221 | Degradome sequencing | positive | | validated |
| tarbase | MIMAT0000524 | mmu-let-7e-5p | Fgf2 | 14173 | ENSMUSG00000037225 | Degradome sequencing//Degradome sequencing | positive | | validated |
| tarbase | MIMAT0000524 | mmu-let-7e-5p | Mxd4 | | ENSMUSG00000037235 | Degradome sequencing//Degradome sequencing | positive | | validated |
| tarbase | MIMAT0000524 | mmu-let-7e-5p | Matr3 | | ENSMUSG00000037236 | Degradome sequencing//Degradome sequencing//Degradome sequencing | positive | | validated |
| tarbase | MIMAT0000524 | mmu-let-7e-5p | Clic4 | 29876 | ENSMUSG00000037242 | Degradome sequencing | positive | | validated |
| tarbase | MIMAT0000524 | mmu-let-7e-5p | Larp1 | 73158 | ENSMUSG00000037331 | Degradome sequencing//Degradome sequencing | positive | | validated |

| | | | | | | | | | |
|---|---|---|---|---|---|---|---|---|---|
| tarbase | MIMAT0000524 | mmu-let-7e-5p | Paqr7 | 71904 | ENSMUSG00000037348 | Degradome sequencing | positive | | validated |
| tarbase | MIMAT0000524 | mmu-let-7e-5p | Letmd1 | 68614 | ENSMUSG00000037353 | Degradome sequencing//Degradome sequencing | positive | | validated |
| tarbase | MIMAT0000524 | mmu-let-7e-5p | Kdm6a | 22289 | ENSMUSG00000037369 | Degradome sequencing//Degradome sequencing//Degradome sequencing | positive | | validated |
| tarbase | MIMAT0000524 | mmu-let-7e-5p | Tbc1d2b | 67016 | ENSMUSG00000037410 | Degradome sequencing | positive | | validated |
| tarbase | MIMAT0000524 | mmu-let-7e-5p | Ranbp10 | 74334 | ENSMUSG00000037415 | Degradome sequencing | positive | | validated |
| tarbase | MIMAT0000524 | mmu-let-7e-5p | Klf10 | 21847 | ENSMUSG00000037465 | Degradome sequencing | positive | | validated |
| tarbase | MIMAT0000524 | mmu-let-7e-5p | Ppfia1 | 233977 | ENSMUSG00000037519 | Degradome sequencing//Degradome sequencing//Degradome sequencing//Degradome sequencing | positive | | validated |
| tarbase | MIMAT0000524 | mmu-let-7e-5p | Rfx7 | 319758 | ENSMUSG00000037674 | Degradome sequencing//Degradome sequencing | positive | | validated |
| tarbase | MIMAT0000524 | mmu-let-7e-5p | Inf2 | 70435 | ENSMUSG00000037679 | Degradome sequencing | positive | | validated |
| tarbase | MIMAT0000524 | mmu-let-7e-5p | Ddhd1 | 114874 | ENSMUSG00000037697 | Degradome sequencing//Degradome sequencing | positive | | validated |
| tarbase | MIMAT0000524 | mmu-let-7e-5p | Lzts3 | 241638 | ENSMUSG00000037703 | Degradome sequencing | positive | | validated |
| tarbase | MIMAT0000524 | mmu-let-7e-5p | Tmem33 | 67878 | ENSMUSG00000037720 | Degradome sequencing//Degradome sequencing | positive | | validated |
| tarbase | MIMAT0000524 | mmu-let-7e-5p | Fam222b | 216971 | ENSMUSG00000037750 | Degradome sequencing | positive | | validated |
| tarbase | MIMAT0000524 | mmu-let-7e-5p | Jmjd1c | 108829 | ENSMUSG00000037876 | Degradome sequencing//Degradome sequencing//Degradome sequencing//Degradome sequencing | positive | | validated |
| tarbase | MIMAT0000524 | mmu-let-7e-5p | Ccr7 | 12775 | ENSMUSG00000037944 | Degradome sequencing//Degradome sequencing | positive | | validated |
| tarbase | MIMAT0000524 | mmu-let-7e-5p | Cramp1l | 57354 | ENSMUSG00000038002 | Degradome sequencing//Degradome sequencing//Degradome sequencing | positive | | validated |
| tarbase | MIMAT0000524 | mmu-let-7e-5p | Socs1 | 12703 | ENSMUSG00000038037 | Degradome sequencing | positive | | validated |
| tarbase | MIMAT0000524 | mmu-let-7e-5p | Haus6 | 230376 | ENSMUSG00000038047 | Degradome sequencing | positive | | validated |
| tarbase | MIMAT0000524 | mmu-let-7e-5p | Kmt2c | 231051 | ENSMUSG00000038056 | Degradome sequencing | positive | | validated |
| tarbase | MIMAT0000524 | mmu-let-7e-5p | Cdkn2aip | 70925 | ENSMUSG00000038069 | Degradome sequencing | positive | | validated |
| tarbase | MIMAT0000524 | mmu-let-7e-5p | Trappc11 | 320714 | ENSMUSG00000038102 | Degradome sequencing | positive | | validated |
| tarbase | MIMAT0000524 | mmu-let-7e-5p | Ccdc50 | 67501 | ENSMUSG00000038127 | Degradome sequencing | positive | | validated |
| tarbase | MIMAT0000524 | mmu-let-7e-5p | Prdm1 | 12142 | ENSMUSG00000038151 | Degradome sequencing//Degradome sequencing | positive | | validated |
| tarbase | MIMAT0000524 | mmu-let-7e-5p | Ttc39b | 69863 | ENSMUSG00000038172 | Degradome sequencing | positive | | validated |
| tarbase | MIMAT0000524 | mmu-let-7e-5p | Slc43a2 | 215113 | ENSMUSG00000038178 | Degradome sequencing | positive | | validated |
| tarbase | MIMAT0000524 | mmu-let-7e-5p | Fbxo8 | 50753 | ENSMUSG00000038206 | Degradome sequencing//Degradome sequencing | positive | | validated |
| tarbase | MIMAT0000524 | mmu-let-7e-5p | Tapbpl | | ENSMUSG00000038213 | Degradome sequencing | positive | | validated |
| tarbase | MIMAT0000524 | mmu-let-7e-5p | Serpinf2 | 18816 | ENSMUSG00000038224 | Degradome sequencing | positive | | validated |
| tarbase | MIMAT0000524 | mmu-let-7e-5p | | | ENSMUSG00000038244 | Degradome sequencing | positive | | validated |
| tarbase | MIMAT0000524 | mmu-let-7e-5p | Usp38 | 74841 | ENSMUSG00000038250 | Degradome sequencing | positive | | validated |
| tarbase | MIMAT0000524 | mmu-let-7e-5p | Slc22a23 | 73102 | ENSMUSG00000038267 | Degradome sequencing//Degradome sequencing//Degradome sequencing | positive | | validated |
| tarbase | MIMAT0000524 | mmu-let-7e-5p | Ostm1 | 14628 | ENSMUSG00000038280 | Degradome sequencing | positive | | validated |
| tarbase | MIMAT0000524 | mmu-let-7e-5p | Snx25 | 102141 | ENSMUSG00000038291 | Degradome sequencing | positive | | validated |
| tarbase | MIMAT0000524 | mmu-let-7e-5p | Pdzk1 | 59020 | ENSMUSG00000038298 | Degradome sequencing | positive | | validated |

| | | | | | | | | | |
|---|---|---|---|---|---|---|---|---|---|
| tarbase | MIMAT0000524 | mmu-let-7e-5p | Sesn1 | 140742 | ENSMUSG00000038332 | Degradome sequencing//Degradome sequencing//Degradome sequencing//Degradome sequencing//Degradome sequencing | positive | | validated |
| tarbase | MIMAT0000524 | mmu-let-7e-5p | Mlxip | 208104 | ENSMUSG00000038342 | Degradome sequencing | positive | | validated |
| tarbase | MIMAT0000524 | mmu-let-7e-5p | Ncoa6 | 56406 | ENSMUSG00000038369 | Degradome sequencing//Degradome sequencing | positive | | validated |
| tarbase | MIMAT0000524 | mmu-let-7e-5p | Egr1 | | ENSMUSG00000038418 | Degradome sequencing | positive | | validated |
| tarbase | MIMAT0000524 | mmu-let-7e-5p | Cdc40 | 71713 | ENSMUSG00000038446 | Degradome sequencing//Degradome sequencing | positive | | validated |
| tarbase | MIMAT0000524 | mmu-let-7e-5p | Parp12 | 243771 | ENSMUSG00000038507 | Degradome sequencing | positive | | validated |
| tarbase | MIMAT0000524 | mmu-let-7e-5p | Jarid2 | 16468 | ENSMUSG00000038518 | Degradome sequencing//Degradome sequencing | positive | | validated |
| tarbase | MIMAT0000524 | mmu-let-7e-5p | Rgs4 | 19736 | ENSMUSG00000038530 | Degradome sequencing | positive | | validated |
| tarbase | MIMAT0000524 | mmu-let-7e-5p | Zfp280d | 235469 | ENSMUSG00000038535 | Degradome sequencing//Degradome sequencing | positive | | validated |
| tarbase | MIMAT0000524 | mmu-let-7e-5p | Ubn2 | 320538 | ENSMUSG00000038538 | Degradome sequencing//Degradome sequencing | positive | | validated |
| tarbase | MIMAT0000524 | mmu-let-7e-5p | Pptc7 | 320717 | ENSMUSG00000038582 | Degradome sequencing//Degradome sequencing | positive | | validated |
| tarbase | MIMAT0000524 | mmu-let-7e-5p | Dock10 | 210293 | ENSMUSG00000038608 | Degradome sequencing | positive | | validated |
| tarbase | MIMAT0000524 | mmu-let-7e-5p | Ric1 | | ENSMUSG00000038658 | Degradome sequencing | positive | | validated |
| tarbase | MIMAT0000524 | mmu-let-7e-5p | Herc1 | 235439 | ENSMUSG00000038664 | Degradome sequencing//Degradome sequencing | positive | | validated |
| tarbase | MIMAT0000524 | mmu-let-7e-5p | Trps1 | 83925 | ENSMUSG00000038679 | Degradome sequencing | positive | | validated |
| tarbase | MIMAT0000524 | mmu-let-7e-5p | Mapkap1 | 227743 | ENSMUSG00000038696 | Degradome sequencing | positive | | validated |
| tarbase | MIMAT0000524 | mmu-let-7e-5p | Taf5l | 102162 | ENSMUSG00000038697 | Degradome sequencing//Degradome sequencing//Degradome sequencing | positive | | validated |
| tarbase | MIMAT0000524 | mmu-let-7e-5p | Dsel | 319901 | ENSMUSG00000038702 | Degradome sequencing | positive | | validated |
| tarbase | MIMAT0000524 | mmu-let-7e-5p | Golga4 | 54214 | ENSMUSG00000038708 | Degradome sequencing//Degradome sequencing | positive | | validated |
| tarbase | MIMAT0000524 | mmu-let-7e-5p | Wdr26 | 226757 | ENSMUSG00000038733 | Degradome sequencing//Degradome sequencing | positive | | validated |
| tarbase | MIMAT0000524 | mmu-let-7e-5p | Ptpn3 | 545622 | ENSMUSG00000038764 | Degradome sequencing | positive | | validated |
| tarbase | MIMAT0000524 | mmu-let-7e-5p | Gabpb2 | 213054 | ENSMUSG00000038766 | Degradome sequencing | positive | | validated |
| tarbase | MIMAT0000524 | mmu-let-7e-5p | Itpkb | 320404 | ENSMUSG00000038855 | Degradome sequencing//Degradome sequencing | positive | | validated |
| tarbase | MIMAT0000524 | mmu-let-7e-5p | Zfhx3 | 11906 | ENSMUSG00000038872 | Degradome sequencing | positive | | validated |
| tarbase | MIMAT0000524 | mmu-let-7e-5p | Irs2 | 384783 | ENSMUSG00000038894 | Degradome sequencing | positive | | validated |
| tarbase | MIMAT0000524 | mmu-let-7e-5p | Prc1 | 233406 | ENSMUSG00000038943 | Degradome sequencing | positive | | validated |
| tarbase | MIMAT0000524 | mmu-let-7e-5p | Pdk2 | 18604 | ENSMUSG00000038967 | Degradome sequencing//Degradome sequencing | positive | | validated |
| tarbase | MIMAT0000524 | mmu-let-7e-5p | Cables2 | 252966 | ENSMUSG00000038990 | Degradome sequencing | positive | | validated |
| tarbase | MIMAT0000524 | mmu-let-7e-5p | Rreb1 | 68750 | ENSMUSG00000039087 | Degradome sequencing//Degradome sequencing//Degradome sequencing | positive | | validated |
| tarbase | MIMAT0000524 | mmu-let-7e-5p | Nrn1 | 68404 | ENSMUSG00000039114 | Degradome sequencing | positive | | validated |
| tarbase | MIMAT0000524 | mmu-let-7e-5p | Akna | 100182 | ENSMUSG00000039158 | Degradome sequencing//Degradome sequencing | positive | | validated |
| tarbase | MIMAT0000524 | mmu-let-7e-5p | Dap | 223453 | ENSMUSG00000039168 | Degradome sequencing//Degradome sequencing | positive | | validated |

| | | | | | | | | | |
|---|---|---|---|---|---|---|---|---|---|
| tarbase | MIMAT0000524 | mmu-let-7e-5p | Daglb | 231871 | ENSMUSG00000039206 | Degradome sequencing | positive | | validated |
| tarbase | MIMAT0000524 | mmu-let-7e-5p | Gpatch2 | 67769 | ENSMUSG00000039210 | Degradome sequencing | positive | | validated |
| tarbase | MIMAT0000524 | mmu-let-7e-5p | Srrm2 | 75956 | ENSMUSG00000039218 | Degradome sequencing//Degradome sequencing | positive | | validated |
| tarbase | MIMAT0000524 | mmu-let-7e-5p | Rftn1 | 76438 | ENSMUSG00000039316 | Degradome sequencing//Degradome sequencing | positive | | validated |
| tarbase | MIMAT0000524 | mmu-let-7e-5p | Sec24c | 218811 | ENSMUSG00000039367 | Degradome sequencing//Degradome sequencing | positive | | validated |
| tarbase | MIMAT0000524 | mmu-let-7e-5p | Mtmr12 | 268783 | ENSMUSG00000039458 | Degradome sequencing | positive | | validated |
| tarbase | MIMAT0000524 | mmu-let-7e-5p | Mocos | 68591 | ENSMUSG00000039616 | Degradome sequencing//Degradome sequencing//Degradome sequencing//Degradome sequencing | positive | | validated |
| tarbase | MIMAT0000524 | mmu-let-7e-5p | Hnrnpu | | ENSMUSG00000039630 | Degradome sequencing//Degradome sequencing | positive | | validated |
| tarbase | MIMAT0000524 | mmu-let-7e-5p | Mrpl12 | | ENSMUSG00000039640 | Degradome sequencing//Degradome sequencing//Degradome sequencing | positive | | validated |
| tarbase | MIMAT0000524 | mmu-let-7e-5p | Lap3 | 66988 | ENSMUSG00000039682 | Degradome sequencing | positive | | validated |
| tarbase | MIMAT0000524 | mmu-let-7e-5p | Nploc4 | 217365 | ENSMUSG00000039703 | Degradome sequencing | positive | | validated |
| tarbase | MIMAT0000524 | mmu-let-7e-5p | Exo1 | 26909 | ENSMUSG00000039748 | Degradome sequencing | positive | | validated |
| tarbase | MIMAT0000524 | mmu-let-7e-5p | Alkbh4 | 72041 | ENSMUSG00000039754 | Degradome sequencing//Degradome sequencing//Degradome sequencing//Degradome sequencing | positive | | validated |
| tarbase | MIMAT0000524 | mmu-let-7e-5p | Rhbdd2 | 215160 | ENSMUSG00000039917 | Degradome sequencing | positive | | validated |
| tarbase | MIMAT0000524 | mmu-let-7e-5p | Hip1 | 215114 | ENSMUSG00000039959 | Degradome sequencing | positive | | validated |
| tarbase | MIMAT0000524 | mmu-let-7e-5p | Saa4 | 20211 | ENSMUSG00000040017 | Degradome sequencing | positive | | validated |
| tarbase | MIMAT0000524 | mmu-let-7e-5p | Rab11fip | 74998 | ENSMUSG00000040022 | Degradome sequencing | positive | | validated |
| tarbase | MIMAT0000524 | mmu-let-7e-5p | Elavl1 | 15568 | ENSMUSG00000040028 | Degradome sequencing | positive | | validated |
| tarbase | MIMAT0000524 | mmu-let-7e-5p | Baz2a | 116848 | ENSMUSG00000040054 | Degradome sequencing | positive | | validated |
| tarbase | MIMAT0000524 | mmu-let-7e-5p | Gramd1b | 235283 | ENSMUSG00000040111 | Degradome sequencing//Degradome sequencing | positive | | validated |
| tarbase | MIMAT0000524 | mmu-let-7e-5p | | | ENSMUSG00000040195 | Degradome sequencing | positive | | validated |
| tarbase | MIMAT0000524 | mmu-let-7e-5p | Prrc2c | 226562 | ENSMUSG00000040225 | Degradome sequencing//Degradome sequencing//Degradome sequencing | positive | | validated |
| tarbase | MIMAT0000524 | mmu-let-7e-5p | Trappc5 | | ENSMUSG00000040236 | Degradome sequencing | positive | | validated |
| tarbase | MIMAT0000524 | mmu-let-7e-5p | Lrp1 | 16971 | ENSMUSG00000040249 | Degradome sequencing//Degradome sequencing | positive | | validated |
| tarbase | MIMAT0000524 | mmu-let-7e-5p | Bach2 | 12014 | ENSMUSG00000040270 | Degradome sequencing | positive | | validated |
| tarbase | MIMAT0000524 | mmu-let-7e-5p | BC05204 | 399568 | ENSMUSG00000040282 | Degradome sequencing//Degradome sequencing | positive | | validated |
| tarbase | MIMAT0000524 | mmu-let-7e-5p | Ddx58 | | ENSMUSG00000040296 | Degradome sequencing | positive | | validated |
| tarbase | MIMAT0000524 | mmu-let-7e-5p | Suco | 226551 | ENSMUSG00000040297 | Degradome sequencing//Degradome sequencing | positive | | validated |
| tarbase | MIMAT0000524 | mmu-let-7e-5p | Dcaf1 | 321006 | ENSMUSG00000040325 | Degradome sequencing | positive | | validated |
| tarbase | MIMAT0000524 | mmu-let-7e-5p | Trim41 | 211007 | ENSMUSG00000040365 | Degradome sequencing | positive | | validated |
| tarbase | MIMAT0000524 | mmu-let-7e-5p | Rc3h1 | 381305 | ENSMUSG00000040423 | Degradome sequencing | positive | | validated |
| tarbase | MIMAT0000524 | mmu-let-7e-5p | Os9 | 216440 | ENSMUSG00000040462 | Degradome sequencing | positive | | validated |
| tarbase | MIMAT0000524 | mmu-let-7e-5p | Mar-09 | | ENSMUSG00000040502 | Degradome sequencing | positive | | validated |
| tarbase | MIMAT0000524 | mmu-let-7e-5p | Pvr | 52118 | ENSMUSG00000040511 | Degradome sequencing//Degradome sequencing | positive | | validated |

| | | | | | | | | | |
|---|---|---|---|---|---|---|---|---|---|
| tarbase | MIMAT0000524 | mmu-let-7e-5p | Tex2 | 21763 | ENSMUSG00000040548 | Degradome sequencing//Degradome sequencing | positive | | validated |
| tarbase | MIMAT0000524 | mmu-let-7e-5p | Ppip5k2 | 227399 | ENSMUSG00000040648 | Degradome sequencing | positive | | validated |
| tarbase | MIMAT0000524 | mmu-let-7e-5p | Efhd2 | 27984 | ENSMUSG00000040659 | Degradome sequencing//Degradome sequencing | positive | | validated |
| tarbase | MIMAT0000524 | mmu-let-7e-5p | Rad54l2 | 81000 | ENSMUSG00000040661 | Degradome sequencing | positive | | validated |
| tarbase | MIMAT0000524 | mmu-let-7e-5p | Nup88 | 19069 | ENSMUSG00000040667 | Degradome sequencing//Degradome sequencing//Degradome sequencing | positive | | validated |
| tarbase | MIMAT0000524 | mmu-let-7e-5p | Kremen2 | 73016 | ENSMUSG00000040680 | Degradome sequencing//Degradome sequencing | positive | | validated |
| tarbase | MIMAT0000524 | mmu-let-7e-5p | Limd2 | 67803 | ENSMUSG00000040699 | Degradome sequencing | positive | | validated |
| tarbase | MIMAT0000524 | mmu-let-7e-5p | Rnf167 | 70510 | ENSMUSG00000040746 | Degradome sequencing//Degradome sequencing | positive | | validated |
| tarbase | MIMAT0000524 | mmu-let-7e-5p | Psme4 | 103554 | ENSMUSG00000040850 | Degradome sequencing//Degradome sequencing | positive | | validated |
| tarbase | MIMAT0000524 | mmu-let-7e-5p | Ino80d | 227195 | ENSMUSG00000040865 | Degradome sequencing//Degradome sequencing | positive | | validated |
| tarbase | MIMAT0000524 | mmu-let-7e-5p | S100pbp | 74648 | ENSMUSG00000040928 | Degradome sequencing//Degradome sequencing//Degradome sequencing//Degradome sequencing | positive | | validated |
| tarbase | MIMAT0000524 | mmu-let-7e-5p | Tet2 | 214133 | ENSMUSG00000040943 | Degradome sequencing//Degradome sequencing | positive | | validated |
| tarbase | MIMAT0000524 | mmu-let-7e-5p | Chd7 | 320790 | ENSMUSG00000041235 | Degradome sequencing//Degradome sequencing//Degradome sequencing//Degradome sequencing | positive | | validated |
| tarbase | MIMAT0000524 | mmu-let-7e-5p | Wapl | 218914 | ENSMUSG00000041408 | Degradome sequencing//Degradome sequencing//Degradome sequencing | positive | | validated |
| tarbase | MIMAT0000524 | mmu-let-7e-5p | Dicer1 | 192119 | ENSMUSG00000041415 | Degradome sequencing//Degradome sequencing//Degradome sequencing//Degradome sequencing//Degradome sequencing | positive | | validated |
| tarbase | MIMAT0000524 | mmu-let-7e-5p | Pik3r1 | 18708 | ENSMUSG00000041417 | Degradome sequencing//Degradome sequencing//Degradome sequencing//Degradome sequencing | positive | | validated |
| tarbase | MIMAT0000524 | mmu-let-7e-5p | Zfp281 | 226442 | ENSMUSG00000041483 | Degradome sequencing//Degradome sequencing//Degradome sequencing | positive | | validated |
| tarbase | MIMAT0000524 | mmu-let-7e-5p | Kif14 | 381293 | ENSMUSG00000041498 | Degradome sequencing | positive | | validated |
| tarbase | MIMAT0000524 | mmu-let-7e-5p | Rnf123 | 84585 | ENSMUSG00000041528 | Degradome sequencing | positive | | validated |
| tarbase | MIMAT0000524 | mmu-let-7e-5p | Ago1 | 236511 | ENSMUSG00000041530 | Degradome sequencing//Degradome sequencing | positive | | validated |
| tarbase | MIMAT0000524 | mmu-let-7e-5p | Sox5 | 20678 | ENSMUSG00000041540 | Degradome sequencing//Degradome sequencing//Degradome sequencing | positive | | validated |

| | | | | | | | | | |
|---|---|---|---|---|---|---|---|---|---|
| tarbase | MIMAT0000524 | mmu-let-7e-5p | Kif21b | 16565 | ENSMUSG00000041642 | Degradome sequencing//Degradome sequencing | positive | | validated |
| tarbase | MIMAT0000524 | mmu-let-7e-5p | Pcca | 110821 | ENSMUSG00000041650 | Degradome sequencing | positive | | validated |
| tarbase | MIMAT0000524 | mmu-let-7e-5p | Fcho2 | 218503 | ENSMUSG00000041685 | Degradome sequencing//Degradome sequencing//Degradome sequencing | positive | | validated |
| tarbase | MIMAT0000524 | mmu-let-7e-5p | Gpr155 | 68526 | ENSMUSG00000041762 | Degradome sequencing//Degradome sequencing | positive | | validated |
| tarbase | MIMAT0000524 | mmu-let-7e-5p | Tpp2 | 22019 | ENSMUSG00000041763 | Degradome sequencing//Degradome sequencing//Degradome sequencing//Degradome sequencing//Degradome sequencing | positive | | validated |
| tarbase | MIMAT0000524 | mmu-let-7e-5p | Slc16a6 | 104681 | ENSMUSG00000041920 | Degradome sequencing | positive | | validated |
| tarbase | MIMAT0000524 | mmu-let-7e-5p | Mvk | 17855 | ENSMUSG00000041939 | Degradome sequencing | positive | | validated |
| tarbase | MIMAT0000524 | mmu-let-7e-5p | Zbed3 | 72114 | ENSMUSG00000041995 | Degradome sequencing | positive | | validated |
| tarbase | MIMAT0000524 | mmu-let-7e-5p | Tlk1 | 228012 | ENSMUSG00000041997 | Degradome sequencing//Degradome sequencing | positive | | validated |
| tarbase | MIMAT0000524 | mmu-let-7e-5p | Cox10 | 70383 | ENSMUSG00000042148 | Degradome sequencing | positive | | validated |
| tarbase | MIMAT0000524 | mmu-let-7e-5p | Fbxo38 | 107035 | ENSMUSG00000042211 | Degradome sequencing | positive | | validated |
| tarbase | MIMAT0000524 | mmu-let-7e-5p | Ccr8 | 12776 | ENSMUSG00000042262 | Degradome sequencing//Degradome sequencing | positive | | validated |
| tarbase | MIMAT0000524 | mmu-let-7e-5p | S100a13 | 20196 | ENSMUSG00000042312 | Degradome sequencing//Degradome sequencing | positive | | validated |
| tarbase | MIMAT0000524 | mmu-let-7e-5p | Snx18 | 170625 | ENSMUSG00000042364 | Degradome sequencing | positive | | validated |
| tarbase | MIMAT0000524 | mmu-let-7e-5p | Abcb4 | 18670 | ENSMUSG00000042476 | Degradome sequencing | positive | | validated |
| tarbase | MIMAT0000524 | mmu-let-7e-5p | Elmsan1 | 238317 | ENSMUSG00000042507 | Degradome sequencing | positive | | validated |
| tarbase | MIMAT0000524 | mmu-let-7e-5p | Asxl1 | 228790 | ENSMUSG00000042548 | Degradome sequencing | positive | | validated |
| tarbase | MIMAT0000524 | mmu-let-7e-5p | Mier2 | 70427 | ENSMUSG00000042570 | Degradome sequencing | positive | | validated |
| tarbase | MIMAT0000524 | mmu-let-7e-5p | Ube2q1 | 70093 | ENSMUSG00000042572 | Degradome sequencing | positive | | validated |
| tarbase | MIMAT0000524 | mmu-let-7e-5p | Sh2b3 | 16923 | ENSMUSG00000042594 | Degradome sequencing//Degradome sequencing//Degradome sequencing | positive | | validated |
| tarbase | MIMAT0000524 | mmu-let-7e-5p | Stk40 | 74178 | ENSMUSG00000042608 | Degradome sequencing | positive | | validated |
| tarbase | MIMAT0000524 | mmu-let-7e-5p | Arrdc4 | 66412 | ENSMUSG00000042659 | Degradome sequencing | positive | | validated |
| tarbase | MIMAT0000524 | mmu-let-7e-5p | Mapk6 | 50772 | ENSMUSG00000042688 | Degradome sequencing//Degradome sequencing//Degradome sequencing//Degradome sequencing | positive | | validated |
| tarbase | MIMAT0000524 | mmu-let-7e-5p | Bmt2 | 101148 | ENSMUSG00000042742 | Degradome sequencing//Degradome sequencing | positive | | validated |
| tarbase | MIMAT0000524 | mmu-let-7e-5p | Smg7 | 226517 | ENSMUSG00000042772 | Degradome sequencing | positive | | validated |
| tarbase | MIMAT0000524 | mmu-let-7e-5p | Klhl6 | 239743 | ENSMUSG00000043008 | Degradome sequencing//Degradome sequencing//Degradome sequencing | positive | | validated |
| tarbase | MIMAT0000524 | mmu-let-7e-5p | Onecut1 | 15379 | ENSMUSG00000043013 | Degradome sequencing | positive | | validated |
| tarbase | MIMAT0000524 | mmu-let-7e-5p | Edem3 | 66967 | ENSMUSG00000043019 | Degradome sequencing | positive | | validated |
| tarbase | MIMAT0000524 | mmu-let-7e-5p | Mob1a | 232157 | ENSMUSG00000043131 | Degradome sequencing//Degradome sequencing | positive | | validated |
| tarbase | MIMAT0000524 | mmu-let-7e-5p | Setx | 269254 | ENSMUSG00000043535 | Degradome sequencing | positive | | validated |
| tarbase | MIMAT0000524 | mmu-let-7e-5p | Ptpn11 | 19247 | ENSMUSG00000043733 | Degradome sequencing | positive | | validated |
| tarbase | MIMAT0000524 | mmu-let-7e-5p | Taf10 | 24075 | ENSMUSG00000043866 | Degradome sequencing | positive | | validated |
| tarbase | MIMAT0000524 | mmu-let-7e-5p | S1pr2 | 14739 | ENSMUSG00000043895 | Degradome sequencing | positive | | validated |
| tarbase | MIMAT0000524 | mmu-let-7e-5p | Mgat2 | 217664 | ENSMUSG00000043998 | Degradome sequencing | positive | | validated |

| | | | | | | | | | |
|---|---|---|---|---|---|---|---|---|---|
| tarbase | MIMAT0000524 | mmu-let-7e-5p | Rsbn1 | 229675 | ENSMUSG00000044098 | Degradome sequencing//Degradome sequencing//Degradome sequencing | positive | | validated |
| tarbase | MIMAT0000524 | mmu-let-7e-5p | Foxo1 | 56458 | ENSMUSG00000044167 | Degradome sequencing | positive | | validated |
| tarbase | MIMAT0000524 | mmu-let-7e-5p | Crb3 | 224912 | ENSMUSG00000044279 | Degradome sequencing | positive | | validated |
| tarbase | MIMAT0000524 | mmu-let-7e-5p | Tent5c | 74645 | ENSMUSG00000044468 | Degradome sequencing | positive | | validated |
| tarbase | MIMAT0000524 | mmu-let-7e-5p | Zfand3 | 21769 | ENSMUSG00000044477 | Degradome sequencing | positive | | validated |
| tarbase | MIMAT0000524 | mmu-let-7e-5p | 25100390 | 77034 | ENSMUSG00000044496 | Degradome sequencing//Degradome sequencing | positive | | validated |
| tarbase | MIMAT0000524 | mmu-let-7e-5p | Zbtb39 | 320080 | ENSMUSG00000044617 | Degradome sequencing | positive | | validated |
| tarbase | MIMAT0000524 | mmu-let-7e-5p | | | ENSMUSG00000044783 | Degradome sequencing | positive | | validated |
| tarbase | MIMAT0000524 | mmu-let-7e-5p | Zfp36 | | ENSMUSG00000044786 | Degradome sequencing//Degradome sequencing//Degradome sequencing//Degradome sequencing | positive | | validated |
| tarbase | MIMAT0000524 | mmu-let-7e-5p | Setd2 | 235626 | ENSMUSG00000044791 | Degradome sequencing//Degradome sequencing | positive | | validated |
| tarbase | MIMAT0000524 | mmu-let-7e-5p | Sft2d3 | 67158 | ENSMUSG00000044982 | Degradome sequencing//Degradome sequencing | positive | | validated |
| tarbase | MIMAT0000524 | mmu-let-7e-5p | Kmt5b | 225888 | ENSMUSG00000045098 | Degradome sequencing | positive | | validated |
| tarbase | MIMAT0000524 | mmu-let-7e-5p | Rtn4rl1 | 237847 | ENSMUSG00000045287 | Degradome sequencing | positive | | validated |
| tarbase | MIMAT0000524 | mmu-let-7e-5p | Tnfrsf26 | 244237 | ENSMUSG00000045362 | Degradome sequencing | positive | | validated |
| tarbase | MIMAT0000524 | mmu-let-7e-5p | Wdr81 | 192652 | ENSMUSG00000045374 | Degradome sequencing | positive | | validated |
| tarbase | MIMAT0000524 | mmu-let-7e-5p | Akr1e1 | 56043 | ENSMUSG00000045410 | Degradome sequencing | positive | | validated |
| tarbase | MIMAT0000524 | mmu-let-7e-5p | Dipk2a | 68861 | ENSMUSG00000045414 | Degradome sequencing//Degradome sequencing | positive | | validated |
| tarbase | MIMAT0000524 | mmu-let-7e-5p | Tmem60 | | ENSMUSG00000045435 | Degradome sequencing//Degradome sequencing | positive | | validated |
| tarbase | MIMAT0000524 | mmu-let-7e-5p | Trrap | 100683 | ENSMUSG00000045482 | Degradome sequencing | positive | | validated |
| tarbase | MIMAT0000524 | mmu-let-7e-5p | Penk | 18619 | ENSMUSG00000045573 | Degradome sequencing//Degradome sequencing | positive | | validated |
| tarbase | MIMAT0000524 | mmu-let-7e-5p | Adrb2 | 11555 | ENSMUSG00000045730 | Degradome sequencing | positive | | validated |
| tarbase | MIMAT0000524 | mmu-let-7e-5p | Slc16a5 | 217316 | ENSMUSG00000045775 | Degradome sequencing | positive | | validated |
| tarbase | MIMAT0000524 | mmu-let-7e-5p | Eif4g1 | 208643 | ENSMUSG00000045983 | Degradome sequencing | positive | | validated |
| tarbase | MIMAT0000524 | mmu-let-7e-5p | Onecut2 | | ENSMUSG00000045991 | Degradome sequencing | positive | | validated |
| tarbase | MIMAT0000524 | mmu-let-7e-5p | Pofut1 | 140484 | ENSMUSG00000046020 | Degradome sequencing//Degradome sequencing//Degradome sequencing | positive | | validated |
| tarbase | MIMAT0000524 | mmu-let-7e-5p | Ppp1r15b | 108954 | ENSMUSG00000046062 | Degradome sequencing//Degradome sequencing//Degradome sequencing | positive | | validated |
| tarbase | MIMAT0000524 | mmu-let-7e-5p | Igfals | | ENSMUSG00000046070 | Degradome sequencing//Degradome sequencing//Degradome sequencing | positive | | validated |
| tarbase | MIMAT0000524 | mmu-let-7e-5p | Patl1 | 225929 | ENSMUSG00000046139 | Degradome sequencing//Degradome sequencing//Degradome sequencing//Degradome sequencing | positive | | validated |
| tarbase | MIMAT0000524 | mmu-let-7e-5p | Yod1 | | ENSMUSG00000046404 | Degradome sequencing//Degradome sequencing | positive | | validated |
| tarbase | MIMAT0000524 | mmu-let-7e-5p | Cxxc5 | 67393 | ENSMUSG00000046668 | Degradome sequencing//Degradome sequencing//Degradome sequencing//Degradome sequencing | positive | | validated |
| tarbase | MIMAT0000524 | mmu-let-7e-5p | Cldn12 | 64945 | ENSMUSG00000046798 | Degradome sequencing//Degradome sequencing | positive | | validated |

| | | | | | | | | | |
|---|---|---|---|---|---|---|---|---|---|
| tarbase | MIMAT0000524 | mmu-let-7e-5p | Zfp740 | 68744 | ENSMUSG00000046897 | Degradome sequencing | positive | | validated |
| tarbase | MIMAT0000524 | mmu-let-7e-5p | Zfp654 | 72020 | ENSMUSG00000047141 | Degradome sequencing//Degradome sequencing//Degradome sequencing | positive | | validated |
| tarbase | MIMAT0000524 | mmu-let-7e-5p | Ythdf3 | 229096 | ENSMUSG00000047213 | Degradome sequencing//Degradome sequencing//Degradome sequencing | positive | | validated |
| tarbase | MIMAT0000524 | mmu-let-7e-5p | Gphn | 268566 | ENSMUSG00000047454 | Degradome sequencing | positive | | validated |
| tarbase | MIMAT0000524 | mmu-let-7e-5p | Dynlrb1 | 67068 | ENSMUSG00000047459 | Degradome sequencing | positive | | validated |
| tarbase | MIMAT0000524 | mmu-let-7e-5p | Zc3hav1l | 209032 | ENSMUSG00000047749 | Degradome sequencing | positive | | validated |
| tarbase | MIMAT0000524 | mmu-let-7e-5p | Gjb1 | 14618 | ENSMUSG00000047797 | Degradome sequencing//Degradome sequencing | positive | | validated |
| tarbase | MIMAT0000524 | mmu-let-7e-5p | Zfp473 | 243963 | ENSMUSG00000048012 | Degradome sequencing//Degradome sequencing | positive | | validated |
| tarbase | MIMAT0000524 | mmu-let-7e-5p | Kmt2d | 381022 | ENSMUSG00000048154 | Degradome sequencing | positive | | validated |
| tarbase | MIMAT0000524 | mmu-let-7e-5p | Asb8 | 78541 | ENSMUSG00000048175 | Degradome sequencing | positive | | validated |
| tarbase | MIMAT0000524 | mmu-let-7e-5p | Pskh1 | 244631 | ENSMUSG00000048310 | Degradome sequencing//Degradome sequencing | positive | | validated |
| tarbase | MIMAT0000524 | mmu-let-7e-5p | Zfp407 | 240476 | ENSMUSG00000048410 | Degradome sequencing//Degradome sequencing//Degradome sequencing//Degradome sequencing | positive | | validated |
| tarbase | MIMAT0000524 | mmu-let-7e-5p | Nrip1 | 268903 | ENSMUSG00000048490 | Degradome sequencing//Degradome sequencing//Degradome sequencing | positive | | validated |
| tarbase | MIMAT0000524 | mmu-let-7e-5p | Lemd3 | 380664 | ENSMUSG00000048661 | Degradome sequencing | positive | | validated |
| tarbase | MIMAT0000524 | mmu-let-7e-5p | Rhno1 | 72440 | ENSMUSG00000048668 | Degradome sequencing | positive | | validated |
| tarbase | MIMAT0000524 | mmu-let-7e-5p | Ggnbp1 | 70772 | ENSMUSG00000048731 | Degradome sequencing | positive | | validated |
| tarbase | MIMAT0000524 | mmu-let-7e-5p | Phf3 | 213109 | ENSMUSG00000048874 | Degradome sequencing//Degradome sequencing | positive | | validated |
| tarbase | MIMAT0000524 | mmu-let-7e-5p | Sephs2 | 20768 | ENSMUSG00000049091 | Degradome sequencing//Degradome sequencing | positive | | validated |
| tarbase | MIMAT0000524 | mmu-let-7e-5p | Zfp644 | 52397 | ENSMUSG00000049606 | Degradome sequencing//Degradome sequencing//Degradome sequencing | positive | | validated |
| tarbase | MIMAT0000524 | mmu-let-7e-5p | Zbtb5 | 230119 | ENSMUSG00000049657 | Degradome sequencing//Degradome sequencing | positive | | validated |
| tarbase | MIMAT0000524 | mmu-let-7e-5p | Zfp280b | 64453 | ENSMUSG00000049764 | Degradome sequencing//Degradome sequencing//Degradome sequencing | positive | | validated |
| tarbase | MIMAT0000524 | mmu-let-7e-5p | Suox | 211389 | ENSMUSG00000049858 | Degradome sequencing | positive | | validated |
| tarbase | MIMAT0000524 | mmu-let-7e-5p | Slc35c1 | 228368 | ENSMUSG00000049922 | Degradome sequencing | positive | | validated |
| tarbase | MIMAT0000524 | mmu-let-7e-5p | H2ax | 15270 | ENSMUSG00000049932 | Degradome sequencing//Degradome sequencing//Degradome sequencing | positive | | validated |
| tarbase | MIMAT0000524 | mmu-let-7e-5p | Grem2 | 23893 | ENSMUSG00000050069 | Degradome sequencing | positive | | validated |
| tarbase | MIMAT0000524 | mmu-let-7e-5p | 1600012I | 67912 | ENSMUSG00000050088 | Degradome sequencing | positive | | validated |
| tarbase | MIMAT0000524 | mmu-let-7e-5p | Pigm | 67556 | ENSMUSG00000050229 | Degradome sequencing//Degradome sequencing//Degradome sequencing | positive | | validated |
| tarbase | MIMAT0000524 | mmu-let-7e-5p | Hic2 | 58180 | ENSMUSG00000050240 | Degradome sequencing | positive | | validated |
| tarbase | MIMAT0000524 | mmu-let-7e-5p | Rictor | 78757 | ENSMUSG00000050310 | Degradome sequencing//Degradome sequencing | positive | | validated |
| tarbase | MIMAT0000524 | mmu-let-7e-5p | Tor1aip2 | 240832 | ENSMUSG00000050565 | Degradome sequencing//Degradome sequencing//Degradome sequencing//Degradome sequencing | positive | | validated |

| | | | | | | | | | |
|---|---|---|---|---|---|---|---|---|---|
| tarbase | MIMAT0000524 | mmu-let-7e-5p | Maml1 | 103806 | ENSMUSG00000050567 | Degradome sequencing//Degradome sequencing | positive | | validated |
| tarbase | MIMAT0000524 | mmu-let-7e-5p | Tmem37 | | ENSMUSG00000050777 | Degradome sequencing//Degradome sequencing//Degradome sequencing | positive | | validated |
| tarbase | MIMAT0000524 | mmu-let-7e-5p | Tmem12[illegible] | 71929 | ENSMUSG00000050912 | Degradome sequencing | positive | | validated |
| tarbase | MIMAT0000524 | mmu-let-7e-5p | Ankrd37 | 654824 | ENSMUSG00000050914 | Degradome sequencing | positive | | validated |
| tarbase | MIMAT0000524 | mmu-let-7e-5p | Gprc5c | 70355 | ENSMUSG00000051043 | Degradome sequencing | positive | | validated |
| tarbase | MIMAT0000524 | mmu-let-7e-5p | Adnp | 11538 | ENSMUSG00000051149 | Degradome sequencing | positive | | validated |
| tarbase | MIMAT0000524 | mmu-let-7e-5p | Eml5 | 319670 | ENSMUSG00000051166 | Degradome sequencing | positive | | validated |
| tarbase | MIMAT0000524 | mmu-let-7e-5p | Zfp524 | 66056 | ENSMUSG00000051184 | Degradome sequencing | positive | | validated |
| tarbase | MIMAT0000524 | mmu-let-7e-5p | Fam174a | 67698 | ENSMUSG00000051185 | Degradome sequencing | positive | | validated |
| tarbase | MIMAT0000524 | mmu-let-7e-5p | Rnf7 | 19823 | ENSMUSG00000051234 | Degradome sequencing//Degradome sequencing | positive | | validated |
| tarbase | MIMAT0000524 | mmu-let-7e-5p | Usp42 | 76800 | ENSMUSG00000051306 | Degradome sequencing | positive | | validated |
| tarbase | MIMAT0000524 | mmu-let-7e-5p | Plagl2 | 54711 | ENSMUSG00000051413 | Degradome sequencing//Degradome sequencing//Degradome sequencing//Degradome sequencing | positive | | validated |
| tarbase | MIMAT0000524 | mmu-let-7e-5p | Mical3 | 194401 | ENSMUSG00000051586 | Degradome sequencing | positive | | validated |
| tarbase | MIMAT0000524 | mmu-let-7e-5p | B3gnt2 | | ENSMUSG00000051650 | Degradome sequencing//Degradome sequencing | positive | | validated |
| tarbase | MIMAT0000524 | mmu-let-7e-5p | Trim32 | 69807 | ENSMUSG00000051675 | Degradome sequencing | positive | | validated |
| tarbase | MIMAT0000524 | mmu-let-7e-5p | Zfp217 | 228913 | ENSMUSG00000052056 | Degradome sequencing//Degradome sequencing | positive | | validated |
| tarbase | MIMAT0000524 | mmu-let-7e-5p | Dock8 | 76088 | ENSMUSG00000052085 | Degradome sequencing//Degradome sequencing | positive | | validated |
| tarbase | MIMAT0000524 | mmu-let-7e-5p | Zfp622 | 52521 | ENSMUSG00000052253 | Degradome sequencing//Degradome sequencing | positive | | validated |
| tarbase | MIMAT0000524 | mmu-let-7e-5p | Ankrd44 | 329154 | ENSMUSG00000052331 | Degradome sequencing//Degradome sequencing | positive | | validated |
| tarbase | MIMAT0000524 | mmu-let-7e-5p | Nrros | 224109 | ENSMUSG00000052384 | Degradome sequencing//Degradome sequencing | positive | | validated |
| tarbase | MIMAT0000524 | mmu-let-7e-5p | Ezr | 22350 | ENSMUSG00000052397 | Degradome sequencing//Degradome sequencing | positive | | validated |
| tarbase | MIMAT0000524 | mmu-let-7e-5p | Nav2 | 78286 | ENSMUSG00000052512 | Degradome sequencing//Degradome sequencing//Degradome sequencing | positive | | validated |
| tarbase | MIMAT0000524 | mmu-let-7e-5p | Tnrc6a | 233833 | ENSMUSG00000052707 | Degradome sequencing | positive | | validated |
| tarbase | MIMAT0000524 | mmu-let-7e-5p | Swt1 | 66875 | ENSMUSG00000052748 | Degradome sequencing//Degradome sequencing | positive | | validated |
| tarbase | MIMAT0000524 | mmu-let-7e-5p | Junb | 16477 | ENSMUSG00000052837 | Degradome sequencing//Degradome sequencing//Degradome sequencing | positive | | validated |
| tarbase | MIMAT0000524 | mmu-let-7e-5p | Rnf26 | | ENSMUSG00000053128 | Degradome sequencing | positive | | validated |
| tarbase | MIMAT0000524 | mmu-let-7e-5p | Trmt1l | 98685 | ENSMUSG00000053286 | Degradome sequencing | positive | | validated |
| tarbase | MIMAT0000524 | mmu-let-7e-5p | Kdm3a | 104263 | ENSMUSG00000053470 | Degradome sequencing | positive | | validated |
| tarbase | MIMAT0000524 | mmu-let-7e-5p | Nrd1 | 230598 | ENSMUSG00000053510 | Degradome sequencing | positive | | validated |
| tarbase | MIMAT0000524 | mmu-let-7e-5p | Sh3pxd2[illegible] | 14218 | ENSMUSG00000053617 | Degradome sequencing | positive | | validated |
| tarbase | MIMAT0000524 | mmu-let-7e-5p | Dusp7 | 235584 | ENSMUSG00000053716 | Degradome sequencing | positive | | validated |
| tarbase | MIMAT0000524 | mmu-let-7e-5p | Ndst1 | 15531 | ENSMUSG00000054008 | Degradome sequencing | positive | | validated |

| | | | | | | | | | |
|---|---|---|---|---|---|---|---|---|---|
| tarbase | MIMAT0000524 | mmu-let-7e-5p | Ercc6 | 319955 | ENSMUSG00000054051 | Degradome sequencing//Degradome sequencing//Degradome sequencing//Degradome sequencing//Degradome sequencing | positive | | validated |
| tarbase | MIMAT0000524 | mmu-let-7e-5p | Lifr | 16880 | ENSMUSG00000054263 | Degradome sequencing | positive | | validated |
| tarbase | MIMAT0000524 | mmu-let-7e-5p | Mdm4 | 17248 | ENSMUSG00000054387 | Degradome sequencing//Degradome sequencing//Degradome sequencing | positive | | validated |
| tarbase | MIMAT0000524 | mmu-let-7e-5p | Spcs3 | 76687 | ENSMUSG00000054408 | Degradome sequencing//Degradome sequencing//Degradome sequencing//Degradome sequencing | positive | | validated |
| tarbase | MIMAT0000524 | mmu-let-7e-5p | Mettl7a1 | 70152 | ENSMUSG00000054619 | Degradome sequencing | positive | | validated |
| tarbase | MIMAT0000524 | mmu-let-7e-5p | 1600014( | 72244 | ENSMUSG00000054676 | Degradome sequencing//Degradome sequencing//Degradome sequencing | positive | | validated |
| tarbase | MIMAT0000524 | mmu-let-7e-5p | Nsd3 | 234135 | ENSMUSG00000054823 | Degradome sequencing//Degradome sequencing | positive | | validated |
| tarbase | MIMAT0000524 | mmu-let-7e-5p | Pcnx3 | 104401 | ENSMUSG00000054874 | Degradome sequencing | positive | | validated |
| tarbase | MIMAT0000524 | mmu-let-7e-5p | Zzef1 | 195018 | ENSMUSG00000055670 | Degradome sequencing//Degradome sequencing | positive | | validated |
| tarbase | MIMAT0000524 | mmu-let-7e-5p | Ghr | 14600 | ENSMUSG00000055737 | Degradome sequencing//Degradome sequencing//Degradome sequencing//Degradome sequencing | positive | | validated |
| tarbase | MIMAT0000524 | mmu-let-7e-5p | Mia3 | | ENSMUSG00000056050 | Degradome sequencing//Degradome sequencing//Degradome sequencing | positive | | validated |
| tarbase | MIMAT0000524 | mmu-let-7e-5p | Dennd1b | 329260 | ENSMUSG00000056268 | Degradome sequencing | positive | | validated |
| tarbase | MIMAT0000524 | mmu-let-7e-5p | Lig1 | 16881 | ENSMUSG00000056394 | Degradome sequencing | positive | | validated |
| tarbase | MIMAT0000524 | mmu-let-7e-5p | Chd9 | 109151 | ENSMUSG00000056608 | Degradome sequencing//Degradome sequencing//Degradome sequencing//Degradome sequencing | positive | | validated |
| tarbase | MIMAT0000524 | mmu-let-7e-5p | Nbeal2 | 235627 | ENSMUSG00000056724 | Degradome sequencing | positive | | validated |
| tarbase | MIMAT0000524 | mmu-let-7e-5p | Bcl2 | 12043 | ENSMUSG00000057329 | Degradome sequencing | positive | | validated |
| tarbase | MIMAT0000524 | mmu-let-7e-5p | Cep170 | 545389 | ENSMUSG00000057335 | Degradome sequencing//Degradome sequencing | positive | | validated |
| tarbase | MIMAT0000524 | mmu-let-7e-5p | Nsd2 | 107823 | ENSMUSG00000057406 | Degradome sequencing | positive | | validated |
| tarbase | MIMAT0000524 | mmu-let-7e-5p | E2f6 | 50496 | ENSMUSG00000057469 | Degradome sequencing//Degradome sequencing | positive | | validated |
| tarbase | MIMAT0000524 | mmu-let-7e-5p | Sptan1 | 20740 | ENSMUSG00000057738 | Degradome sequencing | positive | | validated |
| tarbase | MIMAT0000524 | mmu-let-7e-5p | Abat | 268860 | ENSMUSG00000057880 | Degradome sequencing | positive | | validated |
| tarbase | MIMAT0000524 | mmu-let-7e-5p | Arhgap35 | 232906 | ENSMUSG00000058230 | Degradome sequencing | positive | | validated |
| tarbase | MIMAT0000524 | mmu-let-7e-5p | Rrp1b | 72462 | ENSMUSG00000058392 | Degradome sequencing | positive | | validated |
| tarbase | MIMAT0000524 | mmu-let-7e-5p | Dhcr7 | 13360 | ENSMUSG00000058454 | Degradome sequencing | positive | | validated |
| tarbase | MIMAT0000524 | mmu-let-7e-5p | 0610030I | 68364 | ENSMUSG00000058706 | Degradome sequencing//Degradome sequencing | positive | | validated |
| tarbase | MIMAT0000524 | mmu-let-7e-5p | Egln2 | 112406 | ENSMUSG00000058709 | Degradome sequencing//Degradome sequencing//Degradome sequencing | positive | | validated |
| tarbase | MIMAT0000524 | mmu-let-7e-5p | Usp47 | 74996 | ENSMUSG00000059263 | Degradome sequencing//Degradome sequencing//Degradome sequencing | positive | | validated |
| tarbase | MIMAT0000524 | mmu-let-7e-5p | Tonsl | 72749 | ENSMUSG00000059323 | Degradome sequencing | positive | | validated |
| tarbase | MIMAT0000524 | mmu-let-7e-5p | Gckr | 231103 | ENSMUSG00000059434 | Degradome sequencing | positive | | validated |
| tarbase | MIMAT0000524 | mmu-let-7e-5p | Max | 17187 | ENSMUSG00000059436 | Degradome sequencing//Degradome sequencing | positive | | validated |

| | | | | | | | | | |
|---|---|---|---|---|---|---|---|---|---|
| tarbase | MIMAT0000524 | mmu-let-7e-5p | Kbtbd2 | 210973 | ENSMUSG00000059486 | Degradome sequencing//Degradome sequencing | positive | | validated |
| tarbase | MIMAT0000524 | mmu-let-7e-5p | Arhgef12 | 69632 | ENSMUSG00000059495 | Degradome sequencing | positive | | validated |
| tarbase | MIMAT0000524 | mmu-let-7e-5p | Zfp266 | 77519 | ENSMUSG00000060510 | Degradome sequencing | positive | | validated |
| tarbase | MIMAT0000524 | mmu-let-7e-5p | Fnip2 | 329679 | ENSMUSG00000061175 | Degradome sequencing | positive | | validated |
| tarbase | MIMAT0000524 | mmu-let-7e-5p | Cxcl12 | 20315 | ENSMUSG00000061353 | Degradome sequencing | positive | | validated |
| tarbase | MIMAT0000524 | mmu-let-7e-5p | Hipk2 | 15258 | ENSMUSG00000061436 | Degradome sequencing//Degradome sequencing//Degradome sequencing | positive | | validated |
| tarbase | MIMAT0000524 | mmu-let-7e-5p | Dot1l | 208266 | ENSMUSG00000061589 | Degradome sequencing | positive | | validated |
| tarbase | MIMAT0000524 | mmu-let-7e-5p | Armt1 | 73419 | ENSMUSG00000061759 | Degradome sequencing//Degradome sequencing | positive | | validated |
| tarbase | MIMAT0000524 | mmu-let-7e-5p | Hsd3b3 | 15494 | ENSMUSG00000062410 | Degradome sequencing//Degradome sequencing | positive | | validated |
| tarbase | MIMAT0000524 | mmu-let-7e-5p | Tlr12 | 384059 | ENSMUSG00000062545 | Degradome sequencing | positive | | validated |
| tarbase | MIMAT0000524 | mmu-let-7e-5p | Phactr2 | 215789 | ENSMUSG00000062866 | Degradome sequencing//Degradome sequencing | positive | | validated |
| tarbase | MIMAT0000524 | mmu-let-7e-5p | Klhl24 | 75785 | ENSMUSG00000062901 | Degradome sequencing | positive | | validated |
| tarbase | MIMAT0000524 | mmu-let-7e-5p | Mtap | 66902 | ENSMUSG00000062937 | Degradome sequencing//Degradome sequencing//Degradome sequencing//Degradome sequencing | positive | | validated |
| tarbase | MIMAT0000524 | mmu-let-7e-5p | Kdr | 16542 | ENSMUSG00000062960 | Degradome sequencing//Degradome sequencing//Degradome sequencing//Degradome sequencing | positive | | validated |
| tarbase | MIMAT0000524 | mmu-let-7e-5p | Kif1b | 16561 | ENSMUSG00000063077 | Degradome sequencing//Degradome sequencing//Degradome sequencing//Degradome sequencing | positive | | validated |
| tarbase | MIMAT0000524 | mmu-let-7e-5p | Zfp26 | 22688 | ENSMUSG00000063108 | Degradome sequencing//Degradome sequencing | positive | | validated |
| tarbase | MIMAT0000524 | mmu-let-7e-5p | Mapk1 | 26413 | ENSMUSG00000063358 | Degradome sequencing//Degradome sequencing | positive | | validated |
| tarbase | MIMAT0000524 | mmu-let-7e-5p | Rnf217 | | ENSMUSG00000063760 | Degradome sequencing | positive | | validated |
| tarbase | MIMAT0000524 | mmu-let-7e-5p | Uqcrh | 66576 | ENSMUSG00000063882 | Degradome sequencing | positive | | validated |
| tarbase | MIMAT0000524 | mmu-let-7e-5p | Crem | 12916 | ENSMUSG00000063889 | Degradome sequencing//Degradome sequencing | positive | | validated |
| tarbase | MIMAT0000524 | mmu-let-7e-5p | Nupl1 | | ENSMUSG00000063895 | Degradome sequencing//Degradome sequencing | positive | | validated |
| tarbase | MIMAT0000524 | mmu-let-7e-5p | Med14 | 26896 | ENSMUSG00000064127 | Degradome sequencing//Degradome sequencing | positive | | validated |
| tarbase | MIMAT0000524 | mmu-let-7e-5p | Rab3ip | 216363 | ENSMUSG00000064181 | Degradome sequencing | positive | | validated |
| tarbase | MIMAT0000524 | mmu-let-7e-5p | Cpped1 | 223978 | ENSMUSG00000065979 | Degradome sequencing | positive | | validated |
| tarbase | MIMAT0000524 | mmu-let-7e-5p | Cyp4a10 | 13117 | ENSMUSG00000066072 | Degradome sequencing | positive | | validated |
| tarbase | MIMAT0000524 | mmu-let-7e-5p | Slc31a1 | 20529 | ENSMUSG00000066150 | Degradome sequencing//Degradome sequencing | positive | | validated |
| tarbase | MIMAT0000524 | mmu-let-7e-5p | Slc31a2 | 20530 | ENSMUSG00000066152 | Degradome sequencing//Degradome sequencing | positive | | validated |
| tarbase | MIMAT0000524 | mmu-let-7e-5p | Rtp3 | 235636 | ENSMUSG00000066319 | Degradome sequencing//Degradome sequencing//Degradome sequencing | positive | | validated |
| tarbase | MIMAT0000524 | mmu-let-7e-5p | Zfyve26 | 211978 | ENSMUSG00000066440 | Degradome sequencing//Degradome sequencing//Degradome sequencing//Degradome sequencing | positive | | validated |

| | | | | | | | | | |
|---|---|---|---|---|---|---|---|---|---|
| tarbase | MIMAT0000524 | mmu-let-7e-5p | 4931406I | 233103 | ENSMUSG00000066571 | Degradome sequencing | positive | | validated |
| tarbase | MIMAT0000524 | mmu-let-7e-5p | Sptbn2 | 20743 | ENSMUSG00000067889 | Degradome sequencing//Degradome sequencing | positive | | validated |
| tarbase | MIMAT0000524 | mmu-let-7e-5p | Afdn | 17356 | ENSMUSG00000068036 | Degradome sequencing | positive | | validated |
| tarbase | MIMAT0000524 | mmu-let-7e-5p | Il2rb | 16185 | ENSMUSG00000068227 | Degradome sequencing//Degradome sequencing | positive | | validated |
| tarbase | MIMAT0000524 | mmu-let-7e-5p | Mgam | 232714 | ENSMUSG00000068587 | Degradome sequencing//Degradome sequencing | positive | | validated |
| tarbase | MIMAT0000524 | mmu-let-7e-5p | Cry2 | 12953 | ENSMUSG00000068742 | Degradome sequencing//Degradome sequencing//Degradome sequencing//Degradome sequencing//Degradome sequencing//Degradome sequencing | positive | | validated |
| tarbase | MIMAT0000524 | mmu-let-7e-5p | Sort1 | 20661 | ENSMUSG00000068747 | Degradome sequencing | positive | | validated |
| tarbase | MIMAT0000524 | mmu-let-7e-5p | Tmem19 | 67226 | ENSMUSG00000069520 | Degradome sequencing | positive | | validated |
| tarbase | MIMAT0000524 | mmu-let-7e-5p | Scyl2 | 213326 | ENSMUSG00000069539 | Degradome sequencing//Degradome sequencing | positive | | validated |
| tarbase | MIMAT0000524 | mmu-let-7e-5p | Arid1b | 239985 | ENSMUSG00000069729 | Degradome sequencing | positive | | validated |
| tarbase | MIMAT0000524 | mmu-let-7e-5p | Msi2 | 76626 | ENSMUSG00000069769 | Degradome sequencing | positive | | validated |
| tarbase | MIMAT0000524 | mmu-let-7e-5p | Ahnak | 66395 | ENSMUSG00000069833 | Degradome sequencing | positive | | validated |
| tarbase | MIMAT0000524 | mmu-let-7e-5p | Rnf213 | 672511 | ENSMUSG00000070327 | Degradome sequencing//Degradome sequencing//Degradome sequencing | positive | | validated |
| tarbase | MIMAT0000524 | mmu-let-7e-5p | Rbm47 | 245945 | ENSMUSG00000070780 | Degradome sequencing//Degradome sequencing//Degradome sequencing | positive | | validated |
| tarbase | MIMAT0000524 | mmu-let-7e-5p | Rraga | 68441 | ENSMUSG00000070934 | Degradome sequencing//Degradome sequencing | positive | | validated |
| tarbase | MIMAT0000524 | mmu-let-7e-5p | Acnat1 | 230161 | ENSMUSG00000070985 | Degradome sequencing | positive | | validated |
| tarbase | MIMAT0000524 | mmu-let-7e-5p | Smim15 | 75616 | ENSMUSG00000071180 | Degradome sequencing//Degradome sequencing | positive | | validated |
| tarbase | MIMAT0000524 | mmu-let-7e-5p | Ccnf | 12449 | ENSMUSG00000072082 | Degradome sequencing | positive | | validated |
| tarbase | MIMAT0000524 | mmu-let-7e-5p | Smim10l1 | | ENSMUSG00000072704 | Degradome sequencing | positive | | validated |
| tarbase | MIMAT0000524 | mmu-let-7e-5p | Nfxl1 | 100978 | ENSMUSG00000072889 | Degradome sequencing | positive | | validated |
| tarbase | MIMAT0000524 | mmu-let-7e-5p | Nbeal1 | 269198 | ENSMUSG00000073664 | Degradome sequencing | positive | | validated |
| tarbase | MIMAT0000524 | mmu-let-7e-5p | Mup11 | 100039028 | ENSMUSG00000073834 | Degradome sequencing//Degradome sequencing//Degradome sequencing | positive | | validated |
| tarbase | MIMAT0000524 | mmu-let-7e-5p | Nlrc5 | 434341 | ENSMUSG00000074151 | Degradome sequencing | positive | | validated |
| tarbase | MIMAT0000524 | mmu-let-7e-5p | Zfp568 | 243905 | ENSMUSG00000074221 | Degradome sequencing//Degradome sequencing//Degradome sequencing//Degradome sequencing//Degradome sequencing | positive | | validated |
| tarbase | MIMAT0000524 | mmu-let-7e-5p | Csnk2a1 | 12995 | ENSMUSG00000074698 | Degradome sequencing | positive | | validated |
| tarbase | MIMAT0000524 | mmu-let-7e-5p | Atxn7l3b | 382423 | ENSMUSG00000074748 | Degradome sequencing//Degradome sequencing | positive | | validated |
| tarbase | MIMAT0000524 | mmu-let-7e-5p | Qser1 | 99003 | ENSMUSG00000074994 | Degradome sequencing | positive | | validated |
| tarbase | MIMAT0000524 | mmu-let-7e-5p | Heg1 | 77446 | ENSMUSG00000075254 | Degradome sequencing//Degradome sequencing | positive | | validated |
| tarbase | MIMAT0000524 | mmu-let-7e-5p | Fign | 60344 | ENSMUSG00000075324 | Degradome sequencing | positive | | validated |
| tarbase | MIMAT0000524 | mmu-let-7e-5p | Rc3h2 | 319817 | ENSMUSG00000075376 | Degradome sequencing//Degradome sequencing | positive | | validated |
| tarbase | MIMAT0000524 | mmu-let-7e-5p | Zfp652 | | ENSMUSG00000075595 | Degradome sequencing | positive | | validated |

| | | | | | | | | | |
|---|---|---|---|---|---|---|---|---|---|
| tarbase | MIMAT0000524 | mmu-let-7e-5p | Klhdc7a | | ENSMUSG00000078234 | Degradome sequencing | positive | | validated |
| tarbase | MIMAT0000524 | mmu-let-7e-5p | Ddi2 | 68817 | ENSMUSG00000078515 | Degradome sequencing | positive | | validated |
| tarbase | MIMAT0000524 | mmu-let-7e-5p | Slfn1 | 20555 | ENSMUSG00000078763 | Degradome sequencing | positive | | validated |
| tarbase | MIMAT0000524 | mmu-let-7e-5p | Ifi47 | 15953 | ENSMUSG00000078920 | Degradome sequencing | positive | | validated |
| tarbase | MIMAT0000524 | mmu-let-7e-5p | Kdelr2 | 66913 | ENSMUSG00000079111 | Degradome sequencing | positive | | validated |
| tarbase | MIMAT0000524 | mmu-let-7e-5p | Zfp664 | 269704 | ENSMUSG00000079215 | Degradome sequencing//Degradome sequencing//Degradome sequencing | positive | | validated |
| tarbase | MIMAT0000524 | mmu-let-7e-5p | Tmppe | | ENSMUSG00000079260 | Degradome sequencing//Degradome sequencing//Degradome sequencing//Degradome sequencing | positive | | validated |
| tarbase | MIMAT0000524 | mmu-let-7e-5p | Plcxd2 | 433022 | ENSMUSG00000087141 | Degradome sequencing//Degradome sequencing//Degradome sequencing | positive | | validated |
| tarbase | MIMAT0000524 | mmu-let-7e-5p | Galnt2 | 108148 | ENSMUSG00000089704 | Degradome sequencing | positive | | validated |
| tarbase | MIMAT0000524 | mmu-let-7e-5p | Shkbp1 | 192192 | ENSMUSG00000089832 | Degradome sequencing | positive | | validated |
| tarbase | MIMAT0000524 | mmu-let-7e-5p | Acad11 | 102632 | ENSMUSG00000090150 | Degradome sequencing//Degradome sequencing | positive | | validated |
| tarbase | MIMAT0000524 | mmu-let-7e-5p | 49305230 | 67647 | ENSMUSG00000090394 | Degradome sequencing | positive | | validated |
| tarbase | MIMAT0000524 | mmu-let-7e-5p | Ccdc71l | 72123 | ENSMUSG00000090946 | Degradome sequencing//Degradome sequencing//Degradome sequencing//Degradome sequencing//Degradome sequencing | positive | | validated |
| tarbase | MIMAT0000524 | mmu-let-7e-5p | Lrch4 | 231798 | ENSMUSG00000093445 | Degradome sequencing | positive | | validated |
| tarbase | MIMAT0000524 | mmu-let-7e-5p | Hmgcs1 | 208715 | ENSMUSG00000093930 | Degradome sequencing//Degradome sequencing//Degradome sequencing | positive | | validated |
| tarbase | MIMAT0000524 | mmu-let-7e-5p | Rnasek | 52898 | ENSMUSG00000093989 | Degradome sequencing//Degradome sequencing | positive | | validated |
| tarbase | MIMAT0000524 | mmu-let-7e-5p | Gm38394 | | ENSMUSG00000094410 | Degradome sequencing//Degradome sequencing//Degradome sequencing//Degradome sequencing | positive | | validated |
| tarbase | MIMAT0000524 | mmu-let-7e-5p | Fignl2 | 668225 | ENSMUSG00000095440 | Degradome sequencing//Degradome sequencing//Degradome sequencing | positive | | validated |
| tarbase | MIMAT0000524 | mmu-let-7e-5p | Syne1 | 64009 | ENSMUSG00000096054 | Degradome sequencing | positive | | validated |
| tarbase | MIMAT0000524 | mmu-let-7e-5p | Cmtm4 | 97487 | ENSMUSG00000096188 | Degradome sequencing | positive | | validated |
| tarbase | MIMAT0000524 | mmu-let-7e-5p | | | ENSMUSG00000096688 | Degradome sequencing//Degradome sequencing//Degradome sequencing//Degradome sequencing | positive | | validated |
| tarbase | MIMAT0000524 | mmu-let-7e-5p | | | ENSMUSG00000099515 | Degradome sequencing | positive | | validated |

| Downregulated in Aged Injured compared to Aged Control | | | | | | | | | |
|---|---|---|---|---|---|---|---|---|---|
| miR-328-3p targets | | | | | | | | | |
| **database** | **mature_mirna_acc** | **mature_mirna_id** | **target_symbol** | **target_entrez** | **target_ensembl** | **experiment** | **support_type** | **pubmed_id** | **type** |
| mirecords | MIMAT0000565 | mmu-miR-328-3p | Bace1 | 23821 | ENSMUSG00000032086 | Western blot//Luciferase activity assay | | 18986979 | validated |
| mirtarbase | MIMAT0000565 | mmu-miR-328-3p | Dcc | 13176 | ENSMUSG00000060534 | HITS-CLIP | Functional MTI (Weak) | 23597149 | validated |
| mirtarbase | MIMAT0000565 | mmu-miR-328-3p | Dcc | 13176 | ENSMUSG00000060534 | HITS-CLIP | Functional MTI (Weak) | 21258322 | validated |
| mirtarbase | MIMAT0000565 | mmu-miR-328-3p | Grk4 | 14772 | ENSMUSG00000052783 | HITS-CLIP | Functional MTI (Weak) | 21258322 | validated |
| mirtarbase | MIMAT0000565 | mmu-miR-328-3p | H2-Q4 | 15015 | ENSMUSG00000035929 | HITS-CLIP | Functional MTI (Weak) | 21258322 | validated |
| mirtarbase | MIMAT0000565 | mmu-miR-328-3p | Maoa | 17161 | ENSMUSG00000025037 | HITS-CLIP | Functional MTI (Weak) | 21258322 | validated |
| mirtarbase | MIMAT0000565 | mmu-miR-328-3p | Map4 | 17758 | ENSMUSG00000032479 | HITS-CLIP | Functional MTI (Weak) | 25083871 | validated |
| mirtarbase | MIMAT0000565 | mmu-miR-328-3p | Rad51d | 19364 | | HITS-CLIP | Functional MTI (Weak) | 21258322 | validated |
| mirtarbase | MIMAT0000565 | mmu-miR-328-3p | Bace1 | 23821 | ENSMUSG00000032086 | Luciferase reporter assay//Northern blot//EMSA | Functional MTI | 18986979 | validated |
| mirtarbase | MIMAT0000565 | mmu-miR-328-3p | Bace2 | 56175 | ENSMUSG00000040605 | Luciferase reporter assay | Non-Functional MTI | 18986979 | validated |
| mirtarbase | MIMAT0000565 | mmu-miR-328-3p | Rint1 | 72772 | ENSMUSG00000028999 | HITS-CLIP | Functional MTI (Weak) | 21258322 | validated |
| mirtarbase | MIMAT0000565 | mmu-miR-328-3p | Ms4a6c | 73656 | ENSMUSG00000079419 | HITS-CLIP | Functional MTI (Weak) | 21258322 | validated |
| mirtarbase | MIMAT0000565 | mmu-miR-328-3p | Dhx8 | 217207 | ENSMUSG00000034931 | HITS-CLIP | Functional MTI (Weak) | 21258322 | validated |
| mirtarbase | MIMAT0000565 | mmu-miR-328-3p | Tbc1d8b | 245638 | ENSMUSG00000042473 | HITS-CLIP | Functional MTI (Weak) | 21258322 | validated |
| mirtarbase | MIMAT0000565 | mmu-miR-328-3p | Nkain3 | 269513 | ENSMUSG00000055761 | HITS-CLIP | Functional MTI (Weak) | 21258322 | validated |
| mirtarbase | MIMAT0000565 | mmu-miR-328-3p | Lig4 | 319583 | ENSMUSG00000049717 | HITS-CLIP | Functional MTI (Weak) | 23142080 | validated |
| mirtarbase | MIMAT0000565 | mmu-miR-328-3p | Aim2 | 383619 | ENSMUSG00000037860 | HITS-CLIP | Functional MTI (Weak) | 21258322 | validated |
| mirtarbase | MIMAT0000565 | mmu-miR-328-3p | Zfp970 | 628308 | | HITS-CLIP | Functional MTI (Weak) | 21258322 | validated |
| mirtarbase | MIMAT0000565 | mmu-miR-328-3p | Gm7609 | 665378 | ENSMUSG00000079457 | HITS-CLIP | Functional MTI (Weak) | 21258322 | validated |
| tarbase | MIMAT0000565 | mmu-miR-328-3p | Klf6 | 23849 | ENSMUSG00000000078 | Degradome sequencing | positive | | validated |
| tarbase | MIMAT0000565 | mmu-miR-328-3p | Comt | 12846 | ENSMUSG00000000326 | Degradome sequencing | positive | | validated |
| tarbase | MIMAT0000565 | mmu-miR-328-3p | Itga5 | 16402 | ENSMUSG00000000555 | Degradome sequencing | positive | | validated |
| tarbase | MIMAT0000565 | mmu-miR-328-3p | Zfp512b | 269401 | ENSMUSG00000000823 | Degradome sequencing | positive | | validated |
| tarbase | MIMAT0000565 | mmu-miR-328-3p | Cct3 | 12462 | ENSMUSG00000001416 | Degradome sequencing | positive | | validated |
| tarbase | MIMAT0000565 | mmu-miR-328-3p | Snd1 | 56463 | ENSMUSG00000001424 | Degradome sequencing | positive | | validated |
| tarbase | MIMAT0000565 | mmu-miR-328-3p | Ell2 | | ENSMUSG00000001542 | Degradome sequencing | positive | | validated |

| | | | | | | | | | |
|---|---|---|---|---|---|---|---|---|---|
| tarbase | MIMAT0000565 | mmu-miR-328-3p | Ugp2 | 216558 | ENSMUSG00000001891 | Degradome sequencing//Degradome sequencing | positive | | validated |
| tarbase | MIMAT0000565 | mmu-miR-328-3p | Supt6 | 20926 | ENSMUSG00000002052 | Degradome sequencing | positive | | validated |
| tarbase | MIMAT0000565 | mmu-miR-328-3p | Supt5 | 20924 | ENSMUSG00000003435 | Degradome sequencing | positive | | validated |
| tarbase | MIMAT0000565 | mmu-miR-328-3p | Inmt | 21743 | ENSMUSG00000003477 | Degradome sequencing | positive | | validated |
| tarbase | MIMAT0000565 | mmu-miR-328-3p | Calr | | ENSMUSG00000003814 | Degradome sequencing | positive | | validated |
| tarbase | MIMAT0000565 | mmu-miR-328-3p | Stat3 | 20848 | ENSMUSG00000004040 | Degradome sequencing//Degradome sequencing | positive | | validated |
| tarbase | MIMAT0000565 | mmu-miR-328-3p | Psap | 19156 | ENSMUSG00000004207 | Degradome sequencing | positive | | validated |
| tarbase | MIMAT0000565 | mmu-miR-328-3p | Mcm5 | 17218 | ENSMUSG00000005410 | Degradome sequencing | positive | | validated |
| tarbase | MIMAT0000565 | mmu-miR-328-3p | Metap1 | 75624 | ENSMUSG00000005813 | Degradome sequencing | positive | | validated |
| tarbase | MIMAT0000565 | mmu-miR-328-3p | Rbm14 | 56275 | ENSMUSG00000006456 | Degradome sequencing | positive | | validated |
| tarbase | MIMAT0000565 | mmu-miR-328-3p | Hap1 | 15114 | ENSMUSG00000006930 | Degradome sequencing | positive | | validated |
| tarbase | MIMAT0000565 | mmu-miR-328-3p | Pole | 18973 | ENSMUSG00000007080 | Degradome sequencing | positive | | validated |
| tarbase | MIMAT0000565 | mmu-miR-328-3p | 2610507B11Rik | 72503 | ENSMUSG00000010277 | Degradome sequencing//Degradome sequencing//Degradome sequencing//Degradome sequencing | positive | | validated |
| tarbase | MIMAT0000565 | mmu-miR-328-3p | Faf1 | 14084 | ENSMUSG00000010517 | Degradome sequencing | positive | | validated |
| tarbase | MIMAT0000565 | mmu-miR-328-3p | Fads1 | 76267 | ENSMUSG00000010663 | Degradome sequencing | positive | | validated |
| tarbase | MIMAT0000565 | mmu-miR-328-3p | Yes1 | 22612 | ENSMUSG00000014932 | Degradome sequencing | positive | | validated |
| tarbase | MIMAT0000565 | mmu-miR-328-3p | C8g | 69379 | ENSMUSG00000015083 | Degradome sequencing//Degradome sequencing//Degradome sequencing//Degradome sequencing | positive | | validated |
| tarbase | MIMAT0000565 | mmu-miR-328-3p | Ctsz | 64138 | ENSMUSG00000016256 | Degradome sequencing | positive | | validated |
| tarbase | MIMAT0000565 | mmu-miR-328-3p | Hnf4a | 15378 | ENSMUSG00000017950 | Degradome sequencing | positive | | validated |
| tarbase | MIMAT0000565 | mmu-miR-328-3p | Med1 | 19014 | ENSMUSG00000018160 | Degradome sequencing | positive | | validated |
| tarbase | MIMAT0000565 | mmu-miR-328-3p | Stard3 | 59045 | ENSMUSG00000018167 | Degradome sequencing | positive | | validated |
| tarbase | MIMAT0000565 | mmu-miR-328-3p | Pip4k2b | 108083 | ENSMUSG00000018547 | Degradome sequencing | positive | | validated |
| tarbase | MIMAT0000565 | mmu-miR-328-3p | Phf23 | 78246 | ENSMUSG00000018572 | Degradome sequencing | positive | | validated |
| tarbase | MIMAT0000565 | mmu-miR-328-3p | Sparc | 20692 | ENSMUSG00000018593 | Degradome sequencing | positive | | validated |
| tarbase | MIMAT0000565 | mmu-miR-328-3p | Nudt4 | 71207 | ENSMUSG00000020029 | Degradome sequencing | positive | | validated |
| tarbase | MIMAT0000565 | mmu-miR-328-3p | Rfx4 | 71137 | ENSMUSG00000020037 | Degradome sequencing | positive | | validated |
| tarbase | MIMAT0000565 | mmu-miR-328-3p | Ncln | 103425 | ENSMUSG00000020238 | Degradome sequencing | positive | | validated |
| tarbase | MIMAT0000565 | mmu-miR-328-3p | Mgat1 | 17308 | ENSMUSG00000020346 | Degradome sequencing//Degradome sequencing//Degradome sequencing//Degradome sequencing | positive | | validated |
| tarbase | MIMAT0000565 | mmu-miR-328-3p | Canx | 12330 | ENSMUSG00000020368 | Degradome sequencing | positive | | validated |
| tarbase | MIMAT0000565 | mmu-miR-328-3p | Rack1 | | ENSMUSG00000020372 | Degradome sequencing | positive | | validated |
| tarbase | MIMAT0000565 | mmu-miR-328-3p | Med7 | 66213 | ENSMUSG00000020397 | Degradome sequencing | positive | | validated |
| tarbase | MIMAT0000565 | mmu-miR-328-3p | Cnot8 | 69125 | ENSMUSG00000020515 | Degradome sequencing | positive | | validated |

| | | | | | | | | | |
|---|---|---|---|---|---|---|---|---|---|
| tarbase | MIMAT0000565 | mmu-miR-328-3p | Zfp36l1 | 12192 | ENSMUSG00000021127 | Degradome sequencing//Degradome sequencing | positive | | validated |
| tarbase | MIMAT0000565 | mmu-miR-328-3p | Susd6 | 217684 | ENSMUSG00000021133 | Degradome sequencing | positive | | validated |
| tarbase | MIMAT0000565 | mmu-miR-328-3p | Entpd5 | 12499 | ENSMUSG00000021236 | Degradome sequencing | positive | | validated |
| tarbase | MIMAT0000565 | mmu-miR-328-3p | Nsd1 | 18193 | ENSMUSG00000021488 | Degradome sequencing | positive | | validated |
| tarbase | MIMAT0000565 | mmu-miR-328-3p | F12 | 58992 | ENSMUSG00000021492 | Degradome sequencing | positive | | validated |
| tarbase | MIMAT0000565 | mmu-miR-328-3p | Hmgcr | 15357 | ENSMUSG00000021670 | Degradome sequencing | positive | | validated |
| tarbase | MIMAT0000565 | mmu-miR-328-3p | Lrp10 | | ENSMUSG00000022175 | Degradome sequencing//Degradome sequencing//Degradome sequencing | positive | | validated |
| tarbase | MIMAT0000565 | mmu-miR-328-3p | Dhrs4 | 28200 | ENSMUSG00000022210 | Degradome sequencing | positive | | validated |
| tarbase | MIMAT0000565 | mmu-miR-328-3p | Mief1 | 239555 | ENSMUSG00000022412 | Degradome sequencing | positive | | validated |
| tarbase | MIMAT0000565 | mmu-miR-328-3p | Myh9 | 17886 | ENSMUSG00000022443 | Degradome sequencing//Degradome sequencing | positive | | validated |
| tarbase | MIMAT0000565 | mmu-miR-328-3p | Srebf2 | 20788 | ENSMUSG00000022463 | Degradome sequencing | positive | | validated |
| tarbase | MIMAT0000565 | mmu-miR-328-3p | Gpt | 76282 | ENSMUSG00000022546 | Degradome sequencing | positive | | validated |
| tarbase | MIMAT0000565 | mmu-miR-328-3p | Cyc1 | 66445 | ENSMUSG00000022551 | Degradome sequencing | positive | | validated |
| tarbase | MIMAT0000565 | mmu-miR-328-3p | Maf1 | 68877 | ENSMUSG00000022553 | Degradome sequencing//Degradome sequencing | positive | | validated |
| tarbase | MIMAT0000565 | mmu-miR-328-3p | Plec | 18810 | ENSMUSG00000022565 | Degradome sequencing | positive | | validated |
| tarbase | MIMAT0000565 | mmu-miR-328-3p | Tymp | | ENSMUSG00000022615 | Degradome sequencing | positive | | validated |
| tarbase | MIMAT0000565 | mmu-miR-328-3p | Serpind1 | 15160 | ENSMUSG00000022766 | Degradome sequencing//Degradome sequencing | positive | | validated |
| tarbase | MIMAT0000565 | mmu-miR-328-3p | Mylk | 107589 | ENSMUSG00000022836 | Degradome sequencing | positive | | validated |
| tarbase | MIMAT0000565 | mmu-miR-328-3p | Ets2 | 23872 | ENSMUSG00000022895 | Degradome sequencing | positive | | validated |
| tarbase | MIMAT0000565 | mmu-miR-328-3p | Rcan1 | 54720 | ENSMUSG00000022951 | Degradome sequencing | positive | | validated |
| tarbase | MIMAT0000565 | mmu-miR-328-3p | Gart | 14450 | ENSMUSG00000022962 | Degradome sequencing | positive | | validated |
| tarbase | MIMAT0000565 | mmu-miR-328-3p | Gpd1 | 14555 | ENSMUSG00000023019 | Degradome sequencing | positive | | validated |
| tarbase | MIMAT0000565 | mmu-miR-328-3p | Nagpa | 27426 | ENSMUSG00000023143 | Degradome sequencing//Degradome sequencing | positive | | validated |
| tarbase | MIMAT0000565 | mmu-miR-328-3p | Cpn2 | 71756 | ENSMUSG00000023176 | Degradome sequencing//Degradome sequencing | positive | | validated |
| tarbase | MIMAT0000565 | mmu-miR-328-3p | Tpi1 | 21991 | ENSMUSG00000023456 | Degradome sequencing//Degradome sequencing | positive | | validated |
| tarbase | MIMAT0000565 | mmu-miR-328-3p | Cyp4f14 | 64385 | ENSMUSG00000024292 | Degradome sequencing | positive | | validated |
| tarbase | MIMAT0000565 | mmu-miR-328-3p | Slc39a7 | 14977 | ENSMUSG00000024327 | Degradome sequencing | positive | | validated |
| tarbase | MIMAT0000565 | mmu-miR-328-3p | Lmnb1 | 16906 | ENSMUSG00000024590 | Degradome sequencing | positive | | validated |
| tarbase | MIMAT0000565 | mmu-miR-328-3p | Hmgxb3 | 106894 | ENSMUSG00000024622 | Degradome sequencing | positive | | validated |
| tarbase | MIMAT0000565 | mmu-miR-328-3p | Tle4 | 21888 | ENSMUSG00000024642 | Degradome sequencing | positive | | validated |
| tarbase | MIMAT0000565 | mmu-miR-328-3p | Cdc42bpg | 240505 | ENSMUSG00000024769 | Degradome sequencing | positive | | validated |
| tarbase | MIMAT0000565 | mmu-miR-328-3p | Habp2 | 226243 | ENSMUSG00000025075 | Degradome sequencing | positive | | validated |
| tarbase | MIMAT0000565 | mmu-miR-328-3p | P4hb | 18453 | ENSMUSG00000025130 | Degradome sequencing | positive | | validated |

| | | | | | | | | | |
|---|---|---|---|---|---|---|---|---|---|
| tarbase | MIMAT0000565 | mmu-miR-328-3p | Actr1a | 54130 | ENSMUSG00000025228 | Degradome sequencing | positive | | validated |
| tarbase | MIMAT0000565 | mmu-miR-328-3p | Hsd17b10 | 15108 | ENSMUSG00000025260 | Degradome sequencing | positive | | validated |
| tarbase | MIMAT0000565 | mmu-miR-328-3p | Itgb1 | 16412 | ENSMUSG00000025809 | Degradome sequencing | positive | | validated |
| tarbase | MIMAT0000565 | mmu-miR-328-3p | Glul | 14645 | ENSMUSG00000026473 | Degradome sequencing//Degradome sequencing//Degradome sequencing | positive | | validated |
| tarbase | MIMAT0000565 | mmu-miR-328-3p | Coq8a | 67426 | ENSMUSG00000026489 | Degradome sequencing | positive | | validated |
| tarbase | MIMAT0000565 | mmu-miR-328-3p | Dcaf8 | 98193 | ENSMUSG00000026554 | Degradome sequencing | positive | | validated |
| tarbase | MIMAT0000565 | mmu-miR-328-3p | F5 | 14067 | ENSMUSG00000026579 | Degradome sequencing | positive | | validated |
| tarbase | MIMAT0000565 | mmu-miR-328-3p | Atf2 | 11909 | ENSMUSG00000027104 | Degradome sequencing | positive | | validated |
| tarbase | MIMAT0000565 | mmu-miR-328-3p | Hsd17b12 | 56348 | ENSMUSG00000027195 | Degradome sequencing | positive | | validated |
| tarbase | MIMAT0000565 | mmu-miR-328-3p | Ivd | 56357 | ENSMUSG00000027332 | Degradome sequencing | positive | | validated |
| tarbase | MIMAT0000565 | mmu-miR-328-3p | Pck1 | 18534 | ENSMUSG00000027513 | Degradome sequencing | positive | | validated |
| tarbase | MIMAT0000565 | mmu-miR-328-3p | Hmgcs2 | 15360 | ENSMUSG00000027875 | Degradome sequencing//Degradome sequencing//Degradome sequencing | positive | | validated |
| tarbase | MIMAT0000565 | mmu-miR-328-3p | Trp53inp1 | 60599 | ENSMUSG00000028211 | Degradome sequencing | positive | | validated |
| tarbase | MIMAT0000565 | mmu-miR-328-3p | Nbn | 27354 | ENSMUSG00000028224 | Degradome sequencing | positive | | validated |
| tarbase | MIMAT0000565 | mmu-miR-328-3p | Stra6l | 74152 | ENSMUSG00000028327 | Degradome sequencing | positive | | validated |
| tarbase | MIMAT0000565 | mmu-miR-328-3p | Nr4a3 | 18124 | ENSMUSG00000028341 | Degradome sequencing | positive | | validated |
| tarbase | MIMAT0000565 | mmu-miR-328-3p | Nol6 | 230082 | ENSMUSG00000028430 | Degradome sequencing | positive | | validated |
| tarbase | MIMAT0000565 | mmu-miR-328-3p | Ubap2 | 68926 | ENSMUSG00000028433 | Degradome sequencing | positive | | validated |
| tarbase | MIMAT0000565 | mmu-miR-328-3p | Tln1 | 21894 | ENSMUSG00000028465 | Degradome sequencing | positive | | validated |
| tarbase | MIMAT0000565 | mmu-miR-328-3p | Macf1 | 11426 | ENSMUSG00000028649 | Degradome sequencing | positive | | validated |
| tarbase | MIMAT0000565 | mmu-miR-328-3p | Capzb | 12345 | ENSMUSG00000028745 | Degradome sequencing//Degradome sequencing//Degradome sequencing | positive | | validated |
| tarbase | MIMAT0000565 | mmu-miR-328-3p | Tinagl1 | 94242 | ENSMUSG00000028776 | Degradome sequencing | positive | | validated |
| tarbase | MIMAT0000565 | mmu-miR-328-3p | Sfpq | 71514 | ENSMUSG00000028820 | Degradome sequencing | positive | | validated |
| tarbase | MIMAT0000565 | mmu-miR-328-3p | H6pd | 100198 | ENSMUSG00000028980 | Degradome sequencing//Degradome sequencing | positive | | validated |
| tarbase | MIMAT0000565 | mmu-miR-328-3p | Kmt2e | 69188 | ENSMUSG00000029004 | Degradome sequencing | positive | | validated |
| tarbase | MIMAT0000565 | mmu-miR-328-3p | Ugdh | 22235 | ENSMUSG00000029201 | Degradome sequencing//Degradome sequencing | positive | | validated |
| tarbase | MIMAT0000565 | mmu-miR-328-3p | Pds5a | 71521 | ENSMUSG00000029202 | Degradome sequencing | positive | | validated |
| tarbase | MIMAT0000565 | mmu-miR-328-3p | Scarb2 | 12492 | ENSMUSG00000029426 | Degradome sequencing | positive | | validated |
| tarbase | MIMAT0000565 | mmu-miR-328-3p | Aldh2 | 11669 | ENSMUSG00000029455 | Degradome sequencing//Degradome sequencing | positive | | validated |
| tarbase | MIMAT0000565 | mmu-miR-328-3p | Dhx37 | 208144 | ENSMUSG00000029480 | Degradome sequencing | positive | | validated |
| tarbase | MIMAT0000565 | mmu-miR-328-3p | Aacs | 78894 | ENSMUSG00000029482 | Degradome sequencing | positive | | validated |
| tarbase | MIMAT0000565 | mmu-miR-328-3p | Ints1 | 68510 | ENSMUSG00000029547 | Degradome sequencing | positive | | validated |
| tarbase | MIMAT0000565 | mmu-miR-328-3p | Arpc1a | 56443 | ENSMUSG00000029621 | Degradome sequencing | positive | | validated |
| tarbase | MIMAT0000565 | mmu-miR-328-3p | Gigyf1 | 57330 | ENSMUSG00000029714 | Degradome sequencing | positive | | validated |
| tarbase | MIMAT0000565 | mmu-miR-328-3p | Itpr1 | 16438 | ENSMUSG00000030102 | Degradome sequencing | positive | | validated |

| | | | | | | | | | |
|---|---|---|---|---|---|---|---|---|---|
| tarbase | MIMAT0000565 | mmu-miR-328-3p | Edem1 | 192193 | ENSMUSG00000030104 | Degradome sequencing//Degradome sequencing | positive | | validated |
| tarbase | MIMAT0000565 | mmu-miR-328-3p | Slc6a13 | 14412 | ENSMUSG00000030108 | Degradome sequencing | positive | | validated |
| tarbase | MIMAT0000565 | mmu-miR-328-3p | Gabarapl1 | 57436 | ENSMUSG00000030161 | Degradome sequencing | positive | | validated |
| tarbase | MIMAT0000565 | mmu-miR-328-3p | Aebp2 | 11569 | ENSMUSG00000030232 | Degradome sequencing | positive | | validated |
| tarbase | MIMAT0000565 | mmu-miR-328-3p | Furin | 18550 | ENSMUSG00000030530 | Degradome sequencing | positive | | validated |
| tarbase | MIMAT0000565 | mmu-miR-328-3p | Arl6ip1 | 54208 | ENSMUSG00000030654 | Degradome sequencing | positive | | validated |
| tarbase | MIMAT0000565 | mmu-miR-328-3p | Mrpl17 | 27397 | ENSMUSG00000030879 | Degradome sequencing | positive | | validated |
| tarbase | MIMAT0000565 | mmu-miR-328-3p | Arfip2 | 76932 | ENSMUSG00000030881 | Degradome sequencing//Degradome sequencing | positive | | validated |
| tarbase | MIMAT0000565 | mmu-miR-328-3p | Taf1 | 270627 | ENSMUSG00000031314 | Degradome sequencing | positive | | validated |
| tarbase | MIMAT0000565 | mmu-miR-328-3p | Slc25a15 | 18408 | ENSMUSG00000031482 | Degradome sequencing | positive | | validated |
| tarbase | MIMAT0000565 | mmu-miR-328-3p | Gpat4 | 102247 | ENSMUSG00000031545 | Degradome sequencing | positive | | validated |
| tarbase | MIMAT0000565 | mmu-miR-328-3p | Tnpo2 | 212999 | ENSMUSG00000031691 | Degradome sequencing//Degradome sequencing | positive | | validated |
| tarbase | MIMAT0000565 | mmu-miR-328-3p | Cfdp1 | | ENSMUSG00000031954 | Degradome sequencing | positive | | validated |
| tarbase | MIMAT0000565 | mmu-miR-328-3p | Aars | 234734 | ENSMUSG00000031960 | Degradome sequencing | positive | | validated |
| tarbase | MIMAT0000565 | mmu-miR-328-3p | Agt | 11606 | ENSMUSG00000031980 | Degradome sequencing | positive | | validated |
| tarbase | MIMAT0000565 | mmu-miR-328-3p | Apoa4 | 11808 | ENSMUSG00000032080 | Degradome sequencing | positive | | validated |
| tarbase | MIMAT0000565 | mmu-miR-328-3p | Neil1 | 72774 | ENSMUSG00000032298 | Degradome sequencing | positive | | validated |
| tarbase | MIMAT0000565 | mmu-miR-328-3p | Atr | 245000 | ENSMUSG00000032409 | Degradome sequencing | positive | | validated |
| tarbase | MIMAT0000565 | mmu-miR-328-3p | Gnai2 | 14678 | ENSMUSG00000032562 | Degradome sequencing | positive | | validated |
| tarbase | MIMAT0000565 | mmu-miR-328-3p | Slc16a1 | 20501 | ENSMUSG00000032902 | Degradome sequencing | positive | | validated |
| tarbase | MIMAT0000565 | mmu-miR-328-3p | Ptprf | 19268 | ENSMUSG00000033295 | Degradome sequencing | positive | | validated |
| tarbase | MIMAT0000565 | mmu-miR-328-3p | Tomm6 | | ENSMUSG00000033475 | Degradome sequencing//Degradome sequencing | positive | | validated |
| tarbase | MIMAT0000565 | mmu-miR-328-3p | Cct2 | 12461 | ENSMUSG00000034024 | Degradome sequencing | positive | | validated |
| tarbase | MIMAT0000565 | mmu-miR-328-3p | Lrrc58 | 320184 | ENSMUSG00000034158 | Degradome sequencing | positive | | validated |
| tarbase | MIMAT0000565 | mmu-miR-328-3p | Rnf43 | 207742 | ENSMUSG00000034177 | Degradome sequencing//Degradome sequencing | positive | | validated |
| tarbase | MIMAT0000565 | mmu-miR-328-3p | Mtmr3 | 74302 | ENSMUSG00000034354 | Degradome sequencing | positive | | validated |
| tarbase | MIMAT0000565 | mmu-miR-328-3p | Gulo | 268756 | ENSMUSG00000034450 | Degradome sequencing | positive | | validated |
| tarbase | MIMAT0000565 | mmu-miR-328-3p | Mon2 | 67074 | ENSMUSG00000034602 | Degradome sequencing | positive | | validated |
| tarbase | MIMAT0000565 | mmu-miR-328-3p | Colgalt1 | 234407 | ENSMUSG00000034807 | Degradome sequencing | positive | | validated |
| tarbase | MIMAT0000565 | mmu-miR-328-3p | Tet3 | 194388 | ENSMUSG00000034832 | Degradome sequencing | positive | | validated |
| tarbase | MIMAT0000565 | mmu-miR-328-3p | Rnf44 | 105239 | ENSMUSG00000034928 | Degradome sequencing | positive | | validated |
| tarbase | MIMAT0000565 | mmu-miR-328-3p | Eef2 | 13629 | ENSMUSG00000034994 | Degradome sequencing | positive | | validated |
| tarbase | MIMAT0000565 | mmu-miR-328-3p | Stab2 | 192188 | ENSMUSG00000035459 | Degradome sequencing | positive | | validated |
| tarbase | MIMAT0000565 | mmu-miR-328-3p | Zhx3 | 320799 | ENSMUSG00000035877 | Degradome sequencing | positive | | validated |
| tarbase | MIMAT0000565 | mmu-miR-328-3p | Hykk | 235386 | ENSMUSG00000035878 | Degradome sequencing | positive | | validated |
| tarbase | MIMAT0000565 | mmu-miR-328-3p | Zfp57 | 22715 | ENSMUSG00000036036 | Degradome sequencing | positive | | validated |
| tarbase | MIMAT0000565 | mmu-miR-328-3p | Sigmar1 | 18391 | ENSMUSG00000036078 | Degradome sequencing | positive | | validated |
| tarbase | MIMAT0000565 | mmu-miR-328-3p | Slc17a2 | 218103 | ENSMUSG00000036110 | Degradome sequencing | positive | | validated |
| tarbase | MIMAT0000565 | mmu-miR-328-3p | Fgf1 | 14164 | ENSMUSG00000036585 | Degradome sequencing | positive | | validated |

| | | | | | | | | | |
|---|---|---|---|---|---|---|---|---|---|
| tarbase | MIMAT0000565 | mmu-miR-328-3p | Sun1 | 77053 | ENSMUSG00000036817 | Degradome sequencing//Degradome sequencing | positive | | validated |
| tarbase | MIMAT0000565 | mmu-miR-328-3p | Lrrc20 | 216011 | ENSMUSG00000037151 | Degradome sequencing | positive | | validated |
| tarbase | MIMAT0000565 | mmu-miR-328-3p | Ctbp1 | 13016 | ENSMUSG00000037373 | Degradome sequencing | positive | | validated |
| tarbase | MIMAT0000565 | mmu-miR-328-3p | Ranbp10 | 74334 | ENSMUSG00000037415 | Degradome sequencing | positive | | validated |
| tarbase | MIMAT0000565 | mmu-miR-328-3p | Fam168b | 214469 | ENSMUSG00000037503 | Degradome sequencing | positive | | validated |
| tarbase | MIMAT0000565 | mmu-miR-328-3p | Crp | 12944 | ENSMUSG00000037942 | Degradome sequencing//Degradome sequencing | positive | | validated |
| tarbase | MIMAT0000565 | mmu-miR-328-3p | Sesn1 | 140742 | ENSMUSG00000038332 | Degradome sequencing | positive | | validated |
| tarbase | MIMAT0000565 | mmu-miR-328-3p | Txlng | 353170 | ENSMUSG00000038344 | Degradome sequencing | positive | | validated |
| tarbase | MIMAT0000565 | mmu-miR-328-3p | Foxq1 | | ENSMUSG00000038415 | Degradome sequencing | positive | | validated |
| tarbase | MIMAT0000565 | mmu-miR-328-3p | Mesd | 67943 | ENSMUSG00000038503 | Degradome sequencing | positive | | validated |
| tarbase | MIMAT0000565 | mmu-miR-328-3p | Jarid2 | 16468 | ENSMUSG00000038518 | Degradome sequencing | positive | | validated |
| tarbase | MIMAT0000565 | mmu-miR-328-3p | Irs2 | 384783 | ENSMUSG00000038894 | Degradome sequencing | positive | | validated |
| tarbase | MIMAT0000565 | mmu-miR-328-3p | Lsm14b | 241846 | ENSMUSG00000039108 | Degradome sequencing | positive | | validated |
| tarbase | MIMAT0000565 | mmu-miR-328-3p | Nrn1 | 68404 | ENSMUSG00000039114 | Degradome sequencing | positive | | validated |
| tarbase | MIMAT0000565 | mmu-miR-328-3p | Prrc2b | 227723 | ENSMUSG00000039262 | Degradome sequencing//Degradome sequencing | positive | | validated |
| tarbase | MIMAT0000565 | mmu-miR-328-3p | Ccdc32 | 269336 | ENSMUSG00000039983 | Degradome sequencing | positive | | validated |
| tarbase | MIMAT0000565 | mmu-miR-328-3p | Baz2a | 116848 | ENSMUSG00000040054 | Degradome sequencing | positive | | validated |
| tarbase | MIMAT0000565 | mmu-miR-328-3p | Thbs1 | 21825 | ENSMUSG00000040152 | Degradome sequencing | positive | | validated |
| tarbase | MIMAT0000565 | mmu-miR-328-3p | Pklr | 18770 | ENSMUSG00000041237 | Degradome sequencing | positive | | validated |
| tarbase | MIMAT0000565 | mmu-miR-328-3p | Pcf11 | 74737 | ENSMUSG00000041328 | Degradome sequencing | positive | | validated |
| tarbase | MIMAT0000565 | mmu-miR-328-3p | Lman1 | 70361 | ENSMUSG00000041891 | Degradome sequencing | positive | | validated |
| tarbase | MIMAT0000565 | mmu-miR-328-3p | Mvk | 17855 | ENSMUSG00000041939 | Degradome sequencing | positive | | validated |
| tarbase | MIMAT0000565 | mmu-miR-328-3p | Dmgdh | 74129 | ENSMUSG00000042102 | Degradome sequencing//Degradome sequencing | positive | | validated |
| tarbase | MIMAT0000565 | mmu-miR-328-3p | Cmklr1 | 14747 | ENSMUSG00000042190 | Degradome sequencing | positive | | validated |
| tarbase | MIMAT0000565 | mmu-miR-328-3p | Pelo | 105083 | ENSMUSG00000042275 | Degradome sequencing | positive | | validated |
| tarbase | MIMAT0000565 | mmu-miR-328-3p | Hsd3b7 | 101502 | ENSMUSG00000042289 | Degradome sequencing | positive | | validated |
| tarbase | MIMAT0000565 | mmu-miR-328-3p | Rbm7 | 67010 | ENSMUSG00000042396 | Degradome sequencing//Degradome sequencing | positive | | validated |
| tarbase | MIMAT0000565 | mmu-miR-328-3p | Atxn2 | 20239 | ENSMUSG00000042605 | Degradome sequencing | positive | | validated |
| tarbase | MIMAT0000565 | mmu-miR-328-3p | Dhx9 | 13211 | ENSMUSG00000042699 | Degradome sequencing//Degradome sequencing | positive | | validated |
| tarbase | MIMAT0000565 | mmu-miR-328-3p | Arf6 | | ENSMUSG00000044147 | Degradome sequencing | positive | | validated |
| tarbase | MIMAT0000565 | mmu-miR-328-3p | Gpr146 | 80290 | ENSMUSG00000044197 | Degradome sequencing | positive | | validated |
| tarbase | MIMAT0000565 | mmu-miR-328-3p | P2ry4 | 57385 | ENSMUSG00000044359 | Degradome sequencing | positive | | validated |
| tarbase | MIMAT0000565 | mmu-miR-328-3p | Tst | 22117 | ENSMUSG00000044986 | Degradome sequencing | positive | | validated |
| tarbase | MIMAT0000565 | mmu-miR-328-3p | Wdr81 | 192652 | ENSMUSG00000045374 | Degradome sequencing | positive | | validated |
| tarbase | MIMAT0000565 | mmu-miR-328-3p | Ccdc149 | 100503884 | ENSMUSG00000045790 | Degradome sequencing | positive | | validated |
| tarbase | MIMAT0000565 | mmu-miR-328-3p | Zfp36l2 | 12193 | ENSMUSG00000045817 | Degradome sequencing | positive | | validated |
| tarbase | MIMAT0000565 | mmu-miR-328-3p | Eif4g1 | 208643 | ENSMUSG00000045983 | Degradome sequencing | positive | | validated |
| tarbase | MIMAT0000565 | mmu-miR-328-3p | Olig1 | 50914 | ENSMUSG00000046160 | Degradome sequencing | positive | | validated |

| | | | | | | | | | |
|---|---|---|---|---|---|---|---|---|---|
| tarbase | MIMAT0000565 | mmu-miR-328-3p | E2f8 | 108961 | ENSMUSG00000046179 | Degradome sequencing | positive | | validated |
| tarbase | MIMAT0000565 | mmu-miR-328-3p | Zfp319 | | ENSMUSG00000046556 | Degradome sequencing | positive | | validated |
| tarbase | MIMAT0000565 | mmu-miR-328-3p | Gm5424 | | ENSMUSG00000046687 | Degradome sequencing | positive | | validated |
| tarbase | MIMAT0000565 | mmu-miR-328-3p | Rnf152 | 320311 | ENSMUSG00000047496 | Degradome sequencing | positive | | validated |
| tarbase | MIMAT0000565 | mmu-miR-328-3p | Apof | | ENSMUSG00000047631 | Degradome sequencing//Degradome sequencing | positive | | validated |
| tarbase | MIMAT0000565 | mmu-miR-328-3p | Wbp1l | 226178 | ENSMUSG00000047731 | Degradome sequencing | positive | | validated |
| tarbase | MIMAT0000565 | mmu-miR-328-3p | Angptl8 | 624219 | ENSMUSG00000047822 | Degradome sequencing | positive | | validated |
| tarbase | MIMAT0000565 | mmu-miR-328-3p | Mlec | 109154 | ENSMUSG00000048578 | Degradome sequencing | positive | | validated |
| tarbase | MIMAT0000565 | mmu-miR-328-3p | Foxo3 | 56484 | ENSMUSG00000048756 | Degradome sequencing | positive | | validated |
| tarbase | MIMAT0000565 | mmu-miR-328-3p | Agtr1a | 11607 | ENSMUSG00000049115 | Degradome sequencing | positive | | validated |
| tarbase | MIMAT0000565 | mmu-miR-328-3p | Retreg2 | 227298 | ENSMUSG00000049339 | Degradome sequencing | positive | | validated |
| tarbase | MIMAT0000565 | mmu-miR-328-3p | Tsku | 244152 | ENSMUSG00000049580 | Degradome sequencing | positive | | validated |
| tarbase | MIMAT0000565 | mmu-miR-328-3p | Armcx4 | 100503043 | ENSMUSG00000049804 | Degradome sequencing | positive | | validated |
| tarbase | MIMAT0000565 | mmu-miR-328-3p | Adra1b | 11548 | ENSMUSG00000050541 | Degradome sequencing | positive | | validated |
| tarbase | MIMAT0000565 | mmu-miR-328-3p | Gprc5c | 70355 | ENSMUSG00000051043 | Degradome sequencing | positive | | validated |
| tarbase | MIMAT0000565 | mmu-miR-328-3p | Pcdh1 | 75599 | ENSMUSG00000051375 | Degradome sequencing | positive | | validated |
| tarbase | MIMAT0000565 | mmu-miR-328-3p | Irf2bp2 | 270110 | ENSMUSG00000051495 | Degradome sequencing | positive | | validated |
| tarbase | MIMAT0000565 | mmu-miR-328-3p | Pcbp1 | 23983 | ENSMUSG00000051695 | Degradome sequencing | positive | | validated |
| tarbase | MIMAT0000565 | mmu-miR-328-3p | Rab4b | 19342 | ENSMUSG00000053291 | Degradome sequencing | positive | | validated |
| tarbase | MIMAT0000565 | mmu-miR-328-3p | Gon4l | 76022 | ENSMUSG00000054199 | Degradome sequencing | positive | | validated |
| tarbase | MIMAT0000565 | mmu-miR-328-3p | Fgfr3 | 14184 | ENSMUSG00000054252 | Degradome sequencing | positive | | validated |
| tarbase | MIMAT0000565 | mmu-miR-328-3p | Nfic | 18029 | ENSMUSG00000055053 | Degradome sequencing | positive | | validated |
| tarbase | MIMAT0000565 | mmu-miR-328-3p | Per2 | 18627 | ENSMUSG00000055866 | Degradome sequencing | positive | | validated |
| tarbase | MIMAT0000565 | mmu-miR-328-3p | Mia3 | | ENSMUSG00000056050 | Degradome sequencing | positive | | validated |
| tarbase | MIMAT0000565 | mmu-miR-328-3p | Ilrun | 224647 | ENSMUSG00000056692 | Degradome sequencing | positive | | validated |
| tarbase | MIMAT0000565 | mmu-miR-328-3p | Sptan1 | 20740 | ENSMUSG00000057738 | Degradome sequencing | positive | | validated |
| tarbase | MIMAT0000565 | mmu-miR-328-3p | Rnf169 | 108937 | ENSMUSG00000058761 | Degradome sequencing//Degradome sequencing | positive | | validated |
| tarbase | MIMAT0000565 | mmu-miR-328-3p | Vwa8 | 219189 | ENSMUSG00000058997 | Degradome sequencing | positive | | validated |
| tarbase | MIMAT0000565 | mmu-miR-328-3p | Hnrnpm | 76936 | ENSMUSG00000059208 | Degradome sequencing | positive | | validated |
| tarbase | MIMAT0000565 | mmu-miR-328-3p | Hopx | 74318 | ENSMUSG00000059325 | Degradome sequencing | positive | | validated |
| tarbase | MIMAT0000565 | mmu-miR-328-3p | Serpina6 | 12401 | ENSMUSG00000060807 | Degradome sequencing | positive | | validated |
| tarbase | MIMAT0000565 | mmu-miR-328-3p | Zcchc14 | 142682 | ENSMUSG00000061410 | Degradome sequencing | positive | | validated |
| tarbase | MIMAT0000565 | mmu-miR-328-3p | Rnf217 | | ENSMUSG00000063760 | Degradome sequencing | positive | | validated |
| tarbase | MIMAT0000565 | mmu-miR-328-3p | Bmpr2 | 12168 | ENSMUSG00000067336 | Degradome sequencing | positive | | validated |
| tarbase | MIMAT0000565 | mmu-miR-328-3p | Sptbn2 | 20743 | ENSMUSG00000067889 | Degradome sequencing//Degradome sequencing | positive | | validated |
| tarbase | MIMAT0000565 | mmu-miR-328-3p | Gm5096 | | ENSMUSG00000069324 | Degradome sequencing//Degradome sequencing | positive | | validated |
| tarbase | MIMAT0000565 | mmu-miR-328-3p | Ugt2b36 | | ENSMUSG00000070704 | Degradome sequencing | positive | | validated |
| tarbase | MIMAT0000565 | mmu-miR-328-3p | Prr16 | | ENSMUSG00000073565 | Degradome sequencing | positive | | validated |
| tarbase | MIMAT0000565 | mmu-miR-328-3p | Ttpa | 50500 | ENSMUSG00000073988 | Degradome sequencing | positive | | validated |
| tarbase | MIMAT0000565 | mmu-miR-328-3p | Osgin1 | 71839 | ENSMUSG00000074063 | Degradome sequencing//Degradome sequencing | positive | | validated |

| | | | | | | | | | |
|---|---|---|---|---|---|---|---|---|---|
| tarbase | MIMAT0000565 | mmu-miR-328-3p | Rpl13a | 22121 | ENSMUSG00000074129 | Degradome sequencing | positive | | validated |
| tarbase | MIMAT0000565 | mmu-miR-328-3p | Zfp568 | 243905 | ENSMUSG00000074221 | Degradome sequencing | positive | | validated |
| tarbase | MIMAT0000565 | mmu-miR-328-3p | Plxnc1 | 54712 | ENSMUSG00000074785 | Degradome sequencing//Degradome sequencing | positive | | validated |
| tarbase | MIMAT0000565 | mmu-miR-328-3p | Plcxd2 | 433022 | ENSMUSG00000087141 | Degradome sequencing | positive | | validated |
| tarbase | MIMAT0000565 | mmu-miR-328-3p | Bcl2l2 | 12050 | ENSMUSG00000089682 | Degradome sequencing | positive | | validated |
| tarbase | MIMAT0000565 | mmu-miR-328-3p | Rbm4 | 19653 | ENSMUSG00000094936 | Degradome sequencing | positive | | validated |

| Downregulated in Aged Injured compared to Aged Control | | | | | | | | | |
|---|---|---|---|---|---|---|---|---|---|
| miR-151-3p targets | | | | | | | | | |
| database | mature_mirna_acc | mature_mirna_id | target_symbol | target_entrez | target_ensembl | experiment | support_type | pubmed_id | type |
| mirtarbase | MIMAT0000161 | mmu-miR-151-3p | Clns1a | 12729 | | HITS-CLIP | Functional MTI (Weak) | 21258322 | validated |
| mirtarbase | MIMAT0000161 | mmu-miR-151-3p | Unc5b | 107449 | ENSMUSG00000020099 | HITS-CLIP | Functional MTI (Weak) | 23142080 | validated |
| mirtarbase | MIMAT0000161 | mmu-miR-151-3p | Rmi2 | 223970 | ENSMUSG00000037991 | HITS-CLIP | Functional MTI (Weak) | 25083871 | validated |
| mirtarbase | MIMAT0000161 | mmu-miR-151-3p | Lrrc4c | 241568 | ENSMUSG00000050587 | HITS-CLIP | Functional MTI (Weak) | 21258322 | validated |
| mirtarbase | MIMAT0000161 | mmu-miR-151-3p | Mup10 | 100039008 | ENSMUSG00000078680 | HITS-CLIP | Functional MTI (Weak) | 23597149 | validated |
| mirtarbase | MIMAT0000161 | mmu-miR-151-3p | Mup11 | 100039028 | ENSMUSG00000073834 | HITS-CLIP | Functional MTI (Weak) | 23597149 | validated |
| mirtarbase | MIMAT0000161 | mmu-miR-151-3p | Mup12 | 100039054 | ENSMUSG00000094793 | HITS-CLIP | Functional MTI (Weak) | 23597149 | validated |
| mirtarbase | MIMAT0000161 | mmu-miR-151-3p | Mup14 | 100039116 | ENSMUSG00000073830 | HITS-CLIP | Functional MTI (Weak) | 23597149 | validated |
| mirtarbase | MIMAT0000161 | mmu-miR-151-3p | Mup14 | 100039116 | ENSMUSG00000106700 | HITS-CLIP | Functional MTI (Weak) | 23597149 | validated |
| mirtarbase | MIMAT0000161 | mmu-miR-151-3p | Mup15 | 100039150 | ENSMUSG00000096674 | HITS-CLIP | Functional MTI (Weak) | 23597149 | validated |
| mirtarbase | MIMAT0000161 | mmu-miR-151-3p | Mup15 | 100039150 | ENSMUSG00000107311 | HITS-CLIP | Functional MTI (Weak) | 23597149 | validated |
| mirtarbase | MIMAT0000161 | mmu-miR-151-3p | Mup17 | 100039206 | ENSMUSG00000096688 | HITS-CLIP | Functional MTI (Weak) | 23597149 | validated |
| mirtarbase | MIMAT0000161 | mmu-miR-151-3p | Mup19 | 100189605 | ENSMUSG00000078673 | HITS-CLIP | Functional MTI (Weak) | 23597149 | validated |
| tarbase | MIMAT0000161 | mmu-miR-151-3p | Ell2 | | ENSMUSG00000001542 | Degradome sequencing | positive | | validated |
| tarbase | MIMAT0000161 | mmu-miR-151-3p | Ergic1 | 67458 | ENSMUSG00000001576 | Degradome sequencing | positive | | validated |
| tarbase | MIMAT0000161 | mmu-miR-151-3p | Trim28 | 21849 | ENSMUSG00000005566 | Degradome sequencing | positive | | validated |
| tarbase | MIMAT0000161 | mmu-miR-151-3p | Hnrnph1 | 59013 | ENSMUSG00000007850 | Degradome sequencing | positive | | validated |
| tarbase | MIMAT0000161 | mmu-miR-151-3p | Tnpo1 | 238799 | ENSMUSG00000009470 | Degradome sequencing | positive | | validated |
| tarbase | MIMAT0000161 | mmu-miR-151-3p | Cbx5 | 12419 | ENSMUSG00000009575 | Degradome sequencing | positive | | validated |
| tarbase | MIMAT0000161 | mmu-miR-151-3p | Slc38a3 | 76257 | ENSMUSG00000010064 | Degradome sequencing | positive | | validated |

| | | | | | | | | | |
|---|---|---|---|---|---|---|---|---|---|
| tarbase | MIMAT0000161 | mmu-miR-151-3p | Efr3a | 76740 | ENSMUSG00000015002 | Degradome sequencing | positive | | validated |
| tarbase | MIMAT0000161 | mmu-miR-151-3p | Tox4 | | ENSMUSG00000016831 | Degradome sequencing | positive | | validated |
| tarbase | MIMAT0000161 | mmu-miR-151-3p | Hnf4a | 15378 | ENSMUSG00000017950 | Degradome sequencing | positive | | validated |
| tarbase | MIMAT0000161 | mmu-miR-151-3p | Reep3 | 28193 | ENSMUSG00000019873 | Degradome sequencing | positive | | validated |
| tarbase | MIMAT0000161 | mmu-miR-151-3p | Ccar1 | 67500 | ENSMUSG00000020074 | Degradome sequencing | positive | | validated |
| tarbase | MIMAT0000161 | mmu-miR-151-3p | Apob | 238055 | ENSMUSG00000020609 | Degradome sequencing | positive | | validated |
| tarbase | MIMAT0000161 | mmu-miR-151-3p | Gna13 | 14674 | ENSMUSG00000020611 | Degradome sequencing | positive | | validated |
| tarbase | MIMAT0000161 | mmu-miR-151-3p | Ddx5 | 13207 | ENSMUSG00000020719 | Degradome sequencing//Degradome sequencing | positive | | validated |
| tarbase | MIMAT0000161 | mmu-miR-151-3p | Spag9 | 70834 | ENSMUSG00000020859 | Degradome sequencing | positive | | validated |
| tarbase | MIMAT0000161 | mmu-miR-151-3p | Clmn | 94040 | ENSMUSG00000021097 | Degradome sequencing | positive | | validated |
| tarbase | MIMAT0000161 | mmu-miR-151-3p | Aldh6a1 | 104776 | ENSMUSG00000021238 | Degradome sequencing//Degradome sequencing | positive | | validated |
| tarbase | MIMAT0000161 | mmu-miR-151-3p | Elovl2 | 54326 | ENSMUSG00000021364 | Degradome sequencing | positive | | validated |
| tarbase | MIMAT0000161 | mmu-miR-151-3p | Dek | 110052 | ENSMUSG00000021377 | Degradome sequencing//Degradome sequencing//Degradome sequencing | positive | | validated |
| tarbase | MIMAT0000161 | mmu-miR-151-3p | Nsd1 | 18193 | ENSMUSG00000021488 | Degradome sequencing | positive | | validated |
| tarbase | MIMAT0000161 | mmu-miR-151-3p | Mapk8 | 26419 | ENSMUSG00000021936 | Degradome sequencing | positive | | validated |
| tarbase | MIMAT0000161 | mmu-miR-151-3p | Retreg1 | 66270 | ENSMUSG00000022270 | Degradome sequencing | positive | | validated |
| tarbase | MIMAT0000161 | mmu-miR-151-3p | Pycrl | 66194 | ENSMUSG00000022571 | Degradome sequencing | positive | | validated |
| tarbase | MIMAT0000161 | mmu-miR-151-3p | Mylk | 107589 | ENSMUSG00000022836 | Degradome sequencing | positive | | validated |
| tarbase | MIMAT0000161 | mmu-miR-151-3p | St6gal1 | 20440 | ENSMUSG00000022885 | Degradome sequencing | positive | | validated |
| tarbase | MIMAT0000161 | mmu-miR-151-3p | Ivns1abp | 117198 | ENSMUSG00000023150 | Degradome sequencing | positive | | validated |

| | | | | | | | | | |
|---|---|---|---|---|---|---|---|---|---|
| tarbase | MIMAT0000161 | mmu-miR-151-3p | Tnfrsf21 | 94185 | ENSMUSG00000023915 | Degradome sequencing | positive | | validated |
| tarbase | MIMAT0000161 | mmu-miR-151-3p | Brd4 | 57261 | ENSMUSG00000024002 | Degradome sequencing | positive | | validated |
| tarbase | MIMAT0000161 | mmu-miR-151-3p | Etf1 | 225363 | ENSMUSG00000024360 | Degradome sequencing | positive | | validated |
| tarbase | MIMAT0000161 | mmu-miR-151-3p | Ttc39c | 72747 | ENSMUSG00000024424 | Degradome sequencing | positive | | validated |
| tarbase | MIMAT0000161 | mmu-miR-151-3p | 3110040N11Rik | 67290 | ENSMUSG00000025102 | Degradome sequencing | positive | | validated |
| tarbase | MIMAT0000161 | mmu-miR-151-3p | Idh1 | 15926 | ENSMUSG00000025950 | Degradome sequencing//Degradome sequencing | positive | | validated |
| tarbase | MIMAT0000161 | mmu-miR-151-3p | Wdr12 | 57750 | ENSMUSG00000026019 | Degradome sequencing | positive | | validated |
| tarbase | MIMAT0000161 | mmu-miR-151-3p | Cyp27a1 | 104086 | ENSMUSG00000026170 | Degradome sequencing//Degradome sequencing | positive | | validated |
| tarbase | MIMAT0000161 | mmu-miR-151-3p | Csrp1 | 13007 | ENSMUSG00000026421 | Degradome sequencing | positive | | validated |
| tarbase | MIMAT0000161 | mmu-miR-151-3p | Tor1aip1 | 208263 | ENSMUSG00000026466 | Degradome sequencing//Degradome sequencing | positive | | validated |
| tarbase | MIMAT0000161 | mmu-miR-151-3p | Eprs | 107508 | ENSMUSG00000026615 | Degradome sequencing | positive | | validated |
| tarbase | MIMAT0000161 | mmu-miR-151-3p | Zc3h15 | 69082 | ENSMUSG00000027091 | Degradome sequencing//Degradome sequencing | positive | | validated |
| tarbase | MIMAT0000161 | mmu-miR-151-3p | Hipk3 | 15259 | ENSMUSG00000027177 | Degradome sequencing | positive | | validated |
| tarbase | MIMAT0000161 | mmu-miR-151-3p | Mttp | 17777 | ENSMUSG00000028158 | Degradome sequencing//Degradome sequencing | positive | | validated |
| tarbase | MIMAT0000161 | mmu-miR-151-3p | Decr1 | 67460 | ENSMUSG00000028223 | Degradome sequencing | positive | | validated |
| tarbase | MIMAT0000161 | mmu-miR-151-3p | Aldob | 230163 | ENSMUSG00000028307 | Degradome sequencing//Degradome sequencing | positive | | validated |
| tarbase | MIMAT0000161 | mmu-miR-151-3p | Echdc2 | 52430 | ENSMUSG00000028601 | Degradome sequencing | positive | | validated |
| tarbase | MIMAT0000161 | mmu-miR-151-3p | Cyp4a14 | 13119 | ENSMUSG00000028715 | Degradome sequencing//Degradome sequencing//Degradome sequencing | positive | | validated |

| | | | | | | | | | |
|---|---|---|---|---|---|---|---|---|---|
| tarbase | MIMAT0000161 | mmu-miR-151-3p | Ripk2 | 192656 | ENSMUSG00000041135 | Degradome sequencing | positive | | validated |
| tarbase | MIMAT0000161 | mmu-miR-151-3p | Zfp451 | 98403 | ENSMUSG00000042197 | Degradome sequencing | positive | | validated |
| tarbase | MIMAT0000161 | mmu-miR-151-3p | Gnl3 | 30877 | ENSMUSG00000042354 | Degradome sequencing | positive | | validated |
| tarbase | MIMAT0000161 | mmu-miR-151-3p | Wdfy3 | 72145 | ENSMUSG00000043940 | Degradome sequencing//Degradome sequencing | positive | | validated |
| tarbase | MIMAT0000161 | mmu-miR-151-3p | Thrap3 | 230753 | ENSMUSG00000043962 | Degradome sequencing | positive | | validated |
| tarbase | MIMAT0000161 | mmu-miR-151-3p | Foxo1 | 56458 | ENSMUSG00000044167 | Degradome sequencing | positive | | validated |
| tarbase | MIMAT0000161 | mmu-miR-151-3p | Tst | 22117 | ENSMUSG00000044986 | Degradome sequencing//Degradome sequencing | positive | | validated |
| tarbase | MIMAT0000161 | mmu-miR-151-3p | Hs6st1 | 50785 | ENSMUSG00000045216 | Degradome sequencing | positive | | validated |
| tarbase | MIMAT0000161 | mmu-miR-151-3p | Ppp1r15b | 108954 | ENSMUSG00000046062 | Degradome sequencing | positive | | validated |
| tarbase | MIMAT0000161 | mmu-miR-151-3p | Slc25a23 | 66972 | ENSMUSG00000046329 | Degradome sequencing//Degradome sequencing | positive | | validated |
| tarbase | MIMAT0000161 | mmu-miR-151-3p | Gjb2 | 14619 | ENSMUSG00000046352 | Degradome sequencing | positive | | validated |
| tarbase | MIMAT0000161 | mmu-miR-151-3p | Rnf24 | 51902 | ENSMUSG00000048911 | Degradome sequencing | positive | | validated |
| tarbase | MIMAT0000161 | mmu-miR-151-3p | Lrrc3 | 237387 | ENSMUSG00000051652 | Degradome sequencing | positive | | validated |
| tarbase | MIMAT0000161 | mmu-miR-151-3p | Selenop | 20363 | ENSMUSG00000064373 | Degradome sequencing//Degradome sequencing//Degradome sequencing | positive | | validated |
| tarbase | MIMAT0000161 | mmu-miR-151-3p | Mcc | 328949 | ENSMUSG00000071856 | Degradome sequencing | positive | | validated |
| tarbase | MIMAT0000161 | mmu-miR-151-3p | Serpina1e | | ENSMUSG00000072849 | Degradome sequencing | positive | | validated |
| tarbase | MIMAT0000161 | mmu-miR-151-3p | Ugt1a7c | 394432 | ENSMUSG00000090124 | Degradome sequencing | positive | | validated |

#### Supplementary Data 2

**Unique target genes of miR-203-3p**
ZNF148
Cav1
Cp
Cpox
Itga4
Myd88
Prkcb
Uri1
Rs1
Stxbp4
Tlr4
Trp63
Ung
Mapk8
Commd10
Gtpbp3
Ces2g
Cyld
Fgf16
Pcdhb19
Snora62
Thsd7b
Ccdc110
Srek1
Zfp281
Katnal1

Frmd3
Bnc2
Zfp300
Alkbh5
Srsf12
H2bc21
Zmiz1
Rbm44
Ptchd4
Ndufa9
Kdelr1
Cdip1
Pgrmc1
Pdk1
Cdc42
Tgfbr1
Slc38a3
Steap4
Hspa8
Lamp2
Crk
Sparc
Acsl1
Psen1
Nedd1
Xpo1
Ppp2ca

Canx
Lpin1
Rffl
Susd6
Lgmn
Yy1
Nsd1
Samd8
Parg
Atad2
Slc38a4
Atp13a3
Son
Lpin2
Vapa
Tcf7l2
Hhex
Flnb
Prim1
Shisa5
Idh1
Slc40a1
Tsn
Slc25a25
Rbms1
Cat
Trpm7

Stx16
Kcnq5
Adh5
Ptbp3
Plpp3
Sepsecs
Paics
Foxp2
Aass
Pon3
Lrig1
Rps6ka3
Sorbs2
Dnaja2
Elovl5
Tomm6
Slc16a2
Pds5b
Ctnnd1
Arhgap5
Hykk
Rnf139
4932438A13Rik
Fam13a
Rdh7
Arhgap32
Insig1

Slc25a23
Aff4
Maml1
Pcbp1
Hbb-bs
Gckr
Nup98
Cyp4a12a
Irgm2
Rbm47
Ganab
Gm5878
Trim71

**Unique target genes of miR-150-5p**
Myb
Vegfa
Pdgfb
Cxcr4
Abl2
Akt1
Zfp36l1
Chic1
Btc
Btrc
Camk4
Cbl
Col4a5
Cycs
Egr2
Eif4g2
Elf1
Elk1
F2r
Gad2
Gpr12
Grm1
H2-T3
Hyal1
Ifngr2
Kif5b
Mc1r

Mpv17
Myd88
Nfic
Emc8
Notch4
Npy1r
Ogdh
Pik3c2a
Pkd2
Pla2g2d
Ppargc1a
Ppp2r3d
Mob4
Rhag
Slc6a2
Srf
Stxbp4
Cntn2
Map3k12
Ercc4
Fbxl3
Chek2
Tmem38b
Kcnk6
Tet1
Angel2
Diaph2
Cxcl11

Car5b
Mrpl19
Arhgap23
Ing5
Sorcs3
Magt1
Lrp2bp
Rgs8
Slc25a39
Tprkb
Prdm16
Pwwp2a
Grap
Mau2
Asxl2
Mpp7
Bicd2
Usp45
H2az2
Rab27b
Kcnn1
Rabif
Unc5b
Ssr1
Chrnb4
Acat1
Adarb1
Cds2

Zfp607b
Tmlhe
Ripk2
Gabpb2
Nav1
Ybey
Heatr6
Tns4
Cd300a
Nol10
Mlh3
Cdc14b
Rprd1a
Pbxip1
Sdad1
Glyctk
Zfp811
Tmem245
Zfp568
Dkc1
AY074887
Depdc5
Zfyve27
A830018L16Rik
Gm14325
Zfp866
Dixdc1
Zfp882

Zfp488
Mettl21e
Gm14430
Zfp970
Gm14326
Gm4631
Mef2d
Kmt2a
Stat6
Cnot11
Slc2a3
Mmd
Hdgf
Metap1
Apc
Pdk1
Mark3
Tgfbr1
Snrpb2
Cbx5
Ptprs
Rnf168
Anapc1
Capza2
Nfe2l2
Cd274
Zfp207
Top2b

Mybl2
Srsf1
Ikzf1
Ywhah
Zfp687
Utrn
Prep
Atp2b1
Ccar1
Vps54
Ahsa2
Hnrnpab
Patz1
Pum2
Prkar1a
Tlk2
Cluh
Blmh
Ywhae
Prpf8
Top2a
Stat5b
Zfyve21
Irf4
Nsd1
Kat6b
Arhgef3
Rnf19a

Twf1
Nde1
Tfrc
Iqcb1
Brwd1
Pim1
Lbh
Birc6
Etf1
Gabbr1
Dcp2
Tcerg1
Csnk1a1
Nars
Zfp91
Cdc37l1
Tnks2
Vldlr
Lcor
Add3
Scd2
Pikfyve
Cd28
Acbd3
Dcaf6
Il2ra
Hspa5
Hipk3

Fbxo3
Caprin1
Pdia3
Snx5
Gnas
Mbnl1
Tpm3
Hadh
Ccne2
Tmeff1
Zdhhc21
Rad23b
Macf1
Nasp
AU040320
Mtor
Rnf4
Gfi1
Ccng2
Abcb9
Sppl3
Exoc4
Mtpn
Mkrn1
Isy1
Foxp1
Bhlhe40
Adipor2

Arhgdib
Sec13
Pagr1a
Il21r
Ash2l
Tnpo2
Ccnb2
Anp32a
Pkm
Ulk3
Pdcd6ip
Topbp1
Cish
Cip2a
Atg9a
Szt2
Mif
Fem1c
Mapkbp1
Mga
Setd5
Smc4
Zfp395
Bmp2k
Tet3
Ap2b1
Rabgap1
Rsf1

Ggta1
Pip4p1
Ube2r2
Oxsr1
Map11
Slc12a9
Cd81
Nufip2
Stk35
Bri3bp
Rara
Ccdc50
Ttc39b
Mllt6
Jarid2
Rpl12
Baz2a
Gramd1b
Bach2
Phc1
Fam117b
Ppp4r3a
Kdm5b
Zmym4
Sem1
Smg7
Hilpda
Zbtb37

Swi5
Kmt5b
Zfp36l2
Atxn1
Cltc
Tnrc6b
Gli2
Mlec
Foxo3
Smcr8
Aff4
Tmsb4x
Fgf4
Prkca
Irf2bp2
Mafg
Tnrc6a
Cbx7
Parp4
Kdm2a
Nfil3
Npm1
Slc44a2
Cep170
Zfp329
Marf1
H2-K1
Cfl2

Zfp26
Ldha
Arih2
Arfgef1
Tcp1
Rnf121
Ptgr2
Csnk1g3
Sox2
Pdzd8
Wipf1
Selenot
Ddi2
Ube2d3
Chd2
Alkbh1
Ccr5
Hmgcs1
H1f0
Psmb9

**Unique target genes of let-7e-5p**
Tlr4
Atp2b2
Col3a1
Col1a1
Col1a2
Ifnar1
Il10
Il13
Itga4
Meis2
Nf2
Scd1
Thbs1
Apc2
Cts8
Col24a1
Lins1
Zfp444
Msi2
Eif3j1
Fbxl14
Itga1
Armc2
Nup214
Gnl3l
Tnfrsf26
Col27a1

Trim71
Mup11
Mup17
Xpo6
Ccnd2
Trim25
Mnt
Itgb2
Tmprss2
Acvr1b
Cd52
Usp32
Mxd1
Calm1
Agpat3
Sp1
Ell2
Ergic1
Gstt1
Gramd3
Chordc1
Kmt2a
Celf2
Def6
Angptl4
Med6
Baz1b
Flii

Nab1
Hbp1
Cdk12
Slc2a3
Dgcr2
Sh3gl1
Prkcsh
Dusp3
Prodh
Aven
Snrnp200
Nfat5
Stat3
Phb2
Lpcat3
Clcn3
Utp20
Coro1c
Ndrg2
Pkn2
Arid3b
Myo9b
Ulk2
Lbr
Pex5
Wdr1
Polr2a
Prlr

Txn2
Crbn
Celf1
Por
Insr
Eif4g2
Pcyt1a
Cs
Slc30a4
Metap1
Ncoa2
Odr4
Kmt2b
Gcat
Mark3
Tgfbr1
Zmiz1
Fgfrl1
Prpf31
Sertad1
Herpud2
Pou2f2
Slc16a12
Tnpo1
Cbx5
Pla2g12b
Slc38a3
Raver1

Kif11
Atp6v0d1
Tmem175
Dedd
Map3k4
Dnajb9
Slc25a13
Slamf1
Zdhhc12
Cacfd1
Arnt
Setdb1
Plekho1
Anp32e
Tab2
Chdh
Fli1
Hsd11b1
Cd274
Lamp2
Atxn10
Eif3d
H3f3b
Cmah
Plcg1
Taok1
Nlk
Aldoc

Arl5b
Suz12
Serinc3
Mmp9
Ctsa
Crk
Cyth3
Ro60
Il12rb2
Srsf1
Akap1
Appbp2
Ncor1
G3bp1
Tbx3
Pnpo
Dync1h1
Ywhah
Nars2
Slc35b4
H13
Gyg
Arid3a
Cd164
Utrn
Slc16a10
Prep
Tnfaip3

Rtn4ip1
Rhobtb1
Uhrf1bp1l
Dusp6
Psen1
Apaf1
Fgd6
Tmcc3
Rfx4
Igf1
Rufy2
Eif4ebp2
Lrig3
Tbk1
Egfr
Vps54
Cpm
Mdm2
Mknk2
Ncln
Hcfc2
Xpo1
Cpeb4
Sptbn1
Ube2b
Btg2
Ppp4r3b
Rnft1

Fam49a
Lpin1
Trib2
Apob
Itsn2
Bcap29
Dnmt3a
Itgb3
Rffl
Map3k3
Gga3
Ankfy1
Stat5b
Tmem101
Map3k14
Sel1l
Pnn
Hif1a
Pacs2
Wdr37
Esyt2
Pfkp
Entpd5
Aldh6a1
Traf3
Gpr132
Irf4
Elovl2

Nup153
Sema4d
Ptch1
Ccnh
Dapk1
Trip13
Clptm1l
Marveld2
Gfm2
Iqgap2
Kif2a
Slc4a7
Map3k1
Vcl
Ap3m1
Mettl6
Arhgef3
Ctsb
Xpo4
Rb1
Lrp10
Golph3
Dcaf11
Myc
Fbxo32
Asap1
Mief1
Myh9

Slc38a2
Txndc11
1810013L24Rik
Crebbp
Atp13a3
Hsf1
Plec
Cep97
Alcam
Cblb
Cpox
Senp5
Tfrc
Rabl3
Ehhadh
Kpna1
Brwd1
Il10rb
Lmbr1l
Akirin1
Igf2r
Chd1
Cenpq
Hsp90ab1
Vegfa
Ubr2
Glo1
Wiz

Lbh
Birc6
Strn
Man2a1
Pdpk1
Abca3
C3
Dusp1
Cul2
Svil
Epc1
Sos1
Map4k3
Abcg8
Mapre2
Wac
Mib1
Tapbp
Tmem173
C2
Map3k2
Bag6
Tnf
Riok3
Hdac3
Tcerg1
Pmaip1
Csnk1a1

Lmnb1
Fads3
Ms4a6b
Ddb1
Cemip2
Rtn3
Syvn1
Uhrf2
Slc25a45
Rps6kb2
Rab1b
Scyl1
Cep55
Prdx3
Ccnj
Msr1
Taf5
Cenpx
Slc16a3
Loxl4
Chuk
Scd2
Sema4g
Arih1
Huwe1
Pfkfb1
Dnajc14
Dgka

Mbd6
Tmem192
Tnrc6c
Mkln1
Bach1
Shisa5
Pard3
Nrp2
Ctla4
Cflar
Gls
Nabp1
Sema4c
Dst
Fam135a
Pnkd
Tuba4a
Trip12
Itm2c
Atg16l1
Lrrfip1
Ubxn4
Ptpn4
Ptprc
Nr5a2
Pfkfb2
Srgap2
Tor1aip1

Xpr1
Rgs16
Ahctf1
Sec16b
Abl2
Kctd3
Lpgat1
Plxna2
Uhmk1
Uap1
Pigc
Rabgap1l
Mllt10
Eng
Crat
Gapvd1
Hc
Brd3
Sec16a
Pmpca
Abca2
Rbms1
45358
Psd4
Abcb11
Clp1
Katnbl1
Nop10

Hipk3
Hao1
Snap23
Ubox5
4930402H24Rik
Rpusd2
Gpcpd1
Rasgrp1
Spred1
Stard7
Bcl2l11
Slc20a1
Snx5
Pag1
Gid8
Helz2
Rpn2
Tti1
Skil
Pik3ca
Ttc14
Ncoa3
Sec62
Ccna2
Kpna4
Tsc22d2
Nras
Notch2

Adar
Ash1l
Cd1d1
Lrba
Rnf115
Bcar3
Tmem56
Tspan5
Rpf1
Trp53inp1
Ccne2
Tgs1
Ndufaf4
Gbp2
Tnfsf8
Ugcg
Ptbp3
Ptprd
Zdhhc21
Rad23b
Elp1
Ubap2
Ubap1
Rgp1
Usp24
Plpp3
Prkaa2
Ak4

Jak1
Tnfrsf1b
Macf1
Eloa
Nsun4
Pqlc2
Ddost
Ak2
Psmb2
Gpatch3
Eya3
Per3
Ube4b
Errfi1
Slc2a5
Mtor
Kmt2e
Nelfa
0
Tmem165
Tgfbr3
Slc10a6
Abcb9
Scarb2
Mapkapk5
Kdm2b
Ncor2
Ints1

Ddx54
Cdk8
Polr1d
Hsph1
Wasl
Lrwd1
Gigyf1
Cald1
Gars
Smarcad1
Pcyox1
Cnbp
Copg1
Foxp1
Slc6a13
Plxnd1
Adipor2
Atf7ip
Golt1b
Etnk1
Cmas
Ergic2
Tnfrsf1a
Vasp
Nipa2
Psd3
Ctsc
Pak4

Cyp2r1
Ppp4c
Lipt2
Pold3
Slco2b1
Spns1
Dgat2
Itgal
Abcc6
Plk1
Abraxas2
Acsm5
Pgap2
Otud5
Abcd1
Piga
F10
Adrb3
Eif4ebp1
Col4a2
Dusp4
Kat6a
Gpat4
Plpp5
Klhl2
Casp3
Slc25a4
Sh3rf1

N4bp1
Sall1
Dnaja2
Gab1
Usp10
Hsbp1
Hsd17b2
Nfatc3
Pla2g15
Ankrd49
Kars
Aars
Abcb10
Urb2
Vps26b
Oaf
Sc5d
Crtam
Acat1
Rdx
Apoa5
Cd3e
Slc37a4
Tyk2
Smarca4
Ldlr
Glce
1700017B05Rik

Tmem30a
Ciao2a
Parp16
Pias1
Atr
Xrn1
Nt5e
Mlh1
Trib1
Wdr48
Vipr1
Dnajc13
Qars
Atxn2l
Trib3
Arap1
Zswim6
Cyb5r4
Mycbp2
Slc35d2
Atg9a
Phldb2
Rac2
Slc35a4
Ptprf
Ctdp1
Dnm2
Map2k4

Agl
Ctdspl2
Lpar6
Fam160b1
Fndc3a
Larp4b
Gtf2a2
Rbfox2
Pcgf3
Cep350
Ucp2
Asb13
Atp7a
Alg3
Lgals3bp
Slc16a2
Rpap1
Phka1
Hdlbp
Pomt2
Lrrc58
Chmp7
Usp54
Setd5
Nek9
Cbl
Smc4
Wdr5b

Trmt5
Eda2r
Slc41a2
Tut4
Pik3ip1
Zyg11b
Bmp2k
Xpot
Pbx2
Tmx4
Cnot6l
Map4k5
Dusp5
Tet3
Acot11
Rnf44
Dhx8
Eef2
Baz1a
Arhgap5
Ap2b1
Hectd1
Adrb1
Galm
Rnf38
Syt1
Hykk
Uba6

Fnip1
Ripor2
Smug1
Sigmar1
Slc17a3
Arl5a
Slf2
Vezt
Gxylt1
Naa30
Gramd1c
Hif1an
Cnot1
Plxnb2
Clcn7
Ago2
Oxsr1
Jmjd4
Ehmt1
C1qb
Aifm1
Rab8b
Sh3glb1
Slc35b2
Ralgapa2
E330009J07Rik
Spry1
Mospd3

Fgf2
Mxd4
Matr3
Clic4
Larp1
Paqr7
Letmd1
Kdm6a
Tbc1d2b
Ranbp10
Klf10
Ppfia1
Rfx7
Inf2
Ddhd1
Lzts3
Tmem33
Fam222b
Jmjd1c
Ccr7
Cramp1l
Socs1
Haus6
Kmt2c
Cdkn2aip
Trappc11
Ccdc50
Prdm1

Ttc39b
Slc43a2
Fbxo8
Tapbpl
Serpinf2
Usp38
Slc22a23
Ostm1
Snx25
Pdzk1
Sesn1
Mlxip
Ncoa6
Egr1
Cdc40
Parp12
Jarid2
Rgs4
Zfp280d
Ubn2
Pptc7
Dock10
Ric1
Herc1
Trps1
Mapkap1
Taf5l
Dsel

Golga4
Wdr26
Ptpn3
Gabpb2
Itpkb
Zfhx3
Irs2
Prc1
Pdk2
Cables2
Rreb1
Nrn1
Akna
Dap
Daglb
Gpatch2
Srrm2
Rftn1
Sec24c
Mtmr12
Mocos
Hnrnpu
Mrpl12
Lap3
Nploc4
Exo1
Alkbh4
Rhbdd2

Hip1
Saa4
Rab11fip2
Elavl1
Baz2a
Gramd1b
Prrc2c
Trappc5
Lrp1
Bach2
BC052040
Ddx58
Suco
Dcaf1
Trim41
Rc3h1
Os9
45360
Pvr
Tex2
Ppip5k2
Efhd2
Rad54l2
Nup88
Kremen2
Limd2
Rnf167
Psme4

Ino80d
S100pbp
Tet2
Chd7
Wapl
Dicer1
Pik3r1
Zfp281
Kif14
Rnf123
Ago1
Sox5
Kif21b
Pcca
Fcho2
Gpr155
Tpp2
Slc16a6
Mvk
Zbed3
Tlk1
Cox10
Fbxo38
Ccr8
S100a13
Snx18
Abcb4
Elmsan1

Asxl1
Mier2
Ube2q1
Sh2b3
Stk40
Arrdc4
Mapk6
Bmt2
Smg7
Klhl6
Onecut1
Edem3
Mob1a
Setx
Ptpn11
Taf10
S1pr2
Mgat2
Rsbn1
Foxo1
Crb3
Tent5c
Zfand3
2510039O18Rik
Zbtb39
Zfp36
Setd2
Sft2d3

Kmt5b
Rtn4rl1
Wdr81
Akr1e1
Dipk2a
Tmem60
Trrap
Penk
Adrb2
Slc16a5
Eif4g1
Onecut2
Pofut1
Ppp1r15b
Igfals
Patl1
Yod1
Cxxc5
Cldn12
Zfp740
Zfp654
Ythdf3
Gphn
Dynlrb1
Zc3hav1l
Gjb1
Zfp473
Kmt2d

Asb8
Pskh1
Zfp407
Nrip1
Lemd3
Rhno1
Ggnbp1
Phf3
Sephs2
Zfp644
Zbtb5
Zfp280b
Suox
Slc35c1
H2ax
Grem2
1600012H06Rik
Pigm
Hic2
Rictor
Tor1aip2
Maml1
Tmem37
Tmem123
Ankrd37
Gprc5c
Adnp
Eml5

Zfp524
Fam174a
Rnf7
Usp42
Plagl2
Mical3
B3gnt2
Trim32
Zfp217
Dock8
Zfp622
Ankrd44
Nrros
Ezr
Nav2
Tnrc6a
Swt1
Junb
Rnf26
Trmt1l
Kdm3a
Nrd1
Sh3pxd2a
Dusp7
Ndst1
Ercc6
Lifr
Mdm4

Spcs3
Mettl7a1
1600014C10Rik
Nsd3
Pcnx3
Zzef1
Ghr
Mia3
Dennd1b
Lig1
Chd9
Nbeal2
Bcl2
Cep170
Nsd2
E2f6
Sptan1
Abat
Arhgap35
Rrp1b
Dhcr7
0610030E20Rik
Egln2
Usp47
Tonsl
Gckr
Max
Kbtbd2

Arhgef12
Zfp266
Fnip2
Cxcl12
Hipk2
Dot1l
Armt1
Hsd3b3
Tlr12
Phactr2
Klhl24
Mtap
Kdr
Kif1b
Zfp26
Mapk1
Rnf217
Uqcrh
Crem
Nupl1
Med14
Rab3ip
Cpped1
Cyp4a10
Slc31a1
Slc31a2
Rtp3
Zfyve26

4931406P16Rik
Sptbn2
Afdn
Il2rb
Mgam
Cry2
Sort1
Tmem19
Scyl2
Arid1b
Ahnak
Rnf213
Rbm47
Rraga
Acnat1
Smim15
Ccnf
Smim10l1
Nfxl1
Nbeal1
Nlrc5
Zfp568
Csnk2a1
Atxn7l3b
Qser1
Heg1
Fign
Rc3h2

Zfp652
Klhdc7a
Ddi2
Slfn1
Ifi47
Kdelr2
Zfp664
Tmppe
Plcxd2
Galnt2
Shkbp1
Acad11
4930523C07Rik
Ccdc71l
Lrch4
Hmgcs1
Rnasek
Gm38394
Fignl2
Syne1
Cmtm4

**unique target genes of miR-328-3p**

Bace1
Dcc
Grk4
H2-Q4
Maoa
Map4
Rad51d
Bace2
Rint1
Ms4a6c
Dhx8
Tbc1d8b
Nkain3
Lig4
Aim2
Zfp970
Gm7609
Klf6
Comt
Itga5
Zfp512b
Cct3
Snd1
Ell2
Ugp2
Supt6

Supt5
Inmt
Calr
Stat3
Psap
Mcm5
Metap1
Rbm14
Hap1
Pole
2610507B11Rik
Faf1
Fads1
Yes1
C8g
Ctsz
Hnf4a
Med1
Stard3
Pip4k2b
Phf23
Sparc
Nudt4
Rfx4
Ncln
Mgat1
Canx

Rack1
Med7
Cnot8
Zfp36l1
Susd6
Entpd5
Nsd1
F12
Hmgcr
Lrp10
Dhrs4
Mief1
Myh9
Srebf2
Gpt
Cyc1
Maf1
Plec
Tymp
Serpind1
Mylk
Ets2
Rcan1
Gart
Gpd1
Nagpa
Cpn2

Tpi1
Cyp4f14
Slc39a7
Lmnb1
Hmgxb3
Tle4
Cdc42bpg
Habp2
P4hb
Actr1a
Hsd17b10
Itgb1
Glul
Coq8a
Dcaf8
F5
Atf2
Hsd17b12
Ivd
Pck1
Hmgcs2
Trp53inp1
Nbn
Stra6l
Nr4a3
Nol6
Ubap2

Tln1
Macf1
Capzb
Tinagl1
Sfpq
H6pd
Kmt2e
Ugdh
Pds5a
Scarb2
Aldh2
Dhx37
Aacs
Ints1
Arpc1a
Gigyf1
Itpr1
Edem1
Slc6a13
Gabarapl1
Aebp2
Furin
Arl6ip1
Mrpl17
Arfip2
Taf1
Slc25a15

Gpat4
Tnpo2
Cfdp1
Aars
Agt
Apoa4
Neil1
Atr
Gnai2
Slc16a1
Ptprf
Tomm6
Cct2
Lrrc58
Rnf43
Mtmr3
Gulo
Mon2
Colgalt1
Tet3
Rnf44
Eef2
Stab2
Zhx3
Hykk
Zfp57
Sigmar1

Slc17a2
Fgf1
Sun1
Lrrc20
Ctbp1
Ranbp10
Fam168b
Crp
Sesn1
Txlng
Foxq1
Mesd
Jarid2
Irs2
Lsm14b
Nrn1
Prrc2b
Ccdc32
Baz2a
Thbs1
Pklr
Pcf11
Lman1
Mvk
Dmgdh
Cmklr1
Pelo

Hsd3b7
Rbm7
Atxn2
Dhx9
Arf6
Gpr146
P2ry4
Tst
Wdr81
Ccdc149
Zfp36l2
Eif4g1
Olig1
E2f8
Zfp319
Gm5424
Rnf152
Apof
Wbp1l
Angptl8
Mlec
Foxo3
Agtr1a
Retreg2
Tsku
Armcx4
Adra1b

Gprc5c
Pcdh1
Irf2bp2
Pcbp1
Rab4b
Gon4l
Fgfr3
Nfic
Per2
Mia3
Ilrun
Sptan1
Rnf169
Vwa8
Hnrnpm
Hopx
Serpina6
Zcchc14
Rnf217
Bmpr2
Sptbn2
Gm5096
Ugt2b36
Prr16
Ttpa
Osgin1
Rpl13a

Zfp568
Plxnc1
Plcxd2
Bcl2l2
Rbm4

**Unique target genes of miR-151-3p**

Clns1a
Unc5b
Rmi2
Lrrc4c
Mup10
Mup11
Mup12
Mup14
Mup15
Mup17
Mup19
Ell2
Ergic1
Trim28
Hnrnph1
Tnpo1
Cbx5
Slc38a3
Efr3a
Tox4
Hnf4a
Reep3
Ccar1
Apob
Gna13
Ddx5

Spag9
Clmn
Aldh6a1
Elovl2
Dek
Nsd1
Mapk8
Retreg1
Pycrl
Mylk
St6gal1
Ivns1abp
Tnfrsf21
Brd4
Etf1
Ttc39c
3110040N11Rik
Idh1
Wdr12
Cyp27a1
Csrp1
Tor1aip1
Eprs
Zc3h15
Hipk3
Mttp
Decr1

Aldob
Echdc2
Cyp4a14
Tinagl1
Cyp3a25
Foxp1
Otc
Pdha1
Dusp4
Got2
Aqp9
Ctsh
Dis3l
Scap
Klf9
Tns2
Scd1
Dzank1
Crp
Trp53inp2
Ypel5
Wdr43
Ripk2
Zfp451
Gnl3
Wdfy3
Thrap3

Foxo1
Tst
Hs6st1
Ppp1r15b
Slc25a23
Gjb2
Rnf24
Lrrc3
Selenop
Mcc
Serpina1e
Ugt1a7c

#### Supplementary Data 3

| | |
|---|---|
| Analysis Type: | PANTHER Overrepresentation Test (Released 20240807) |
| Annotation Version and Release Date: | PANTHER version 19.0 Released 2024-06-20 |
| Analyzed List: | miR-203-3p_all targets.txt (Mus musculus) |
| Reference List: | Mus musculus (all genes in database) |
| Test Type: | FISHER |
| Correction: | FDR |

| PANTHER Pathways | Mus musculus - REFLIST (21836) | miR-203-3p_all targets.txt (119) | miR-203-3p_all targets.txt (expected) | miR-203-3p_all targets.txt (over/under) | miR-203-3p_all targets.txt (fold Enrichment) | miR-203-3p_all targets.txt (raw P-value) | miR-203-3p_all targets.txt (FDR) |
|---|---|---|---|---|---|---|---|
| **Ras Pathway (P04393)** | 72 | 4 | 0.39 | + | 10.19 | 6.48E-04 | 3.48E-02 |
| **CCKR signaling map (P06959)** | 164 | 6 | 0.89 | + | 6.71 | 2.82E-04 | 4.55E-02 |
| **Integrin signalling pathway (P00034)** | 189 | 6 | 1.03 | + | 5.83 | 6.00E-04 | 4.83E-02 |

| | |
|---|---|
| Analysis Type: | PANTHER Overrepresentation Test (Released 20240807) |
| Annotation Version and Release Date: | PANTHER version 19.0 Released 2024-06-20 |
| Analyzed List: | miR-150-5p.txt (Mus musculus) |
| Reference List: | Mus musculus (all genes in database) |
| Test Type: | FISHER |
| Correction: | FDR |

| PANTHER Pathways | Mus musculus - REFLIST (21836) | miR-150-5p.txt (327) | miR-150-5p.txt (expected) | miR-150-5p.txt (over/under) | miR-150-5p.txt (fold Enrichment) | miR-150-5p.txt (raw P-value) | miR-150-5p.txt (FDR) |
|---|---|---|---|---|---|---|---|
| **Hedgehog signaling pathway (P00025)** | 20 | 4 | 0.3 | + | 13.36 | 1.98E-04 | 0.01 |
| **Insulin/IGF pathway-protein kinase B signaling cascade (P00033)** | 38 | 4 | 0.57 | + | 7.03 | 2.44E-03 | 0.04 |
| **Interleukin signaling pathway (P00036)** | 94 | 9 | 1.41 | + | 6.39 | 1.18E-05 | 0.00 |
| **Parkinson disease (P00049)** | 92 | 7 | 1.38 | + | 5.08 | 4.67E-04 | 0.01 |
| **EGF receptor signaling pathway (P00018)** | 133 | 10 | 1.99 | + | 5.02 | 3.28E-05 | 0.00 |
| **PDGF signaling pathway (P00047)** | 140 | 10 | 2.1 | + | 4.77 | 5.09E-05 | 0.00 |
| **FGF signaling pathway (P00021)** | 122 | 7 | 1.83 | + | 3.83 | 2.44E-03 | 0.04 |
| **CCKR signaling map (P06959)** | 164 | 9 | 2.46 | + | 3.66 | 8.45E-04 | 0.02 |
| **Angiogenesis (P00005)** | 169 | 9 | 2.53 | + | 3.56 | 1.04E-03 | 0.02 |
| Unclassified (UNCLASSIFIED) | 19255 | 259 | 288.35 | - | 0.9 | 3.65E-06 | 0.00 |

| | |
|---|---|
| Analysis Type: | PANTHER Overrepresentation Test (Released 20240807) |
| Annotation Version and Release Date: | PANTHER version 19.0 Released 2024-06-20 |
| Analyzed List: | let-7e-5p_all targets.txt (Mus musculus) |
| Reference List: | Mus musculus (all genes in database) |
| Test Type: | FISHER |
| Correction: | FDR |

| PANTHER Pathways | Mus musculus - REFLIST (21836) | let-7e-5p_all targets.txt (1022) | let-7e-5p_all targets.txt (expected) | let-7e-5p_all targets.txt (over/under) | let-7e-5p_all targets.txt (fold Enrichment) | let-7e-5p_all targets.txt (raw P-value) | let-7e-5p_all targets.txt (FDR) |
|---|---|---|---|---|---|---|---|
| **JAK/STAT signaling pathway (P00038)** | 17 | 6 | 0.8 | + | 7.54 | 8.21E-05 | 1.02E-03 |
| **Hypoxia response via HIF activation (P00030)** | 29 | 8 | 1.36 | + | 5.89 | 3.99E-05 | 5.84E-04 |
| **Insulin/IGF pathway-protein kinase B signaling cascade (P00033)** | 38 | 10 | 1.78 | + | 5.62 | 6.89E-06 | 1.11E-04 |
| **Oxidative stress response (P00046)** | 54 | 12 | 2.53 | + | 4.75 | 5.75E-06 | 1.03E-04 |
| **Insulin/IGF pathway-mitogen activated protein kinase kinase/MAP kinase cascade (P00032)** | 36 | 8 | 1.68 | + | 4.75 | 2.10E-04 | 2.26E-03 |
| **p53 pathway feedback loops 2 (P04398)** | 49 | 10 | 2.29 | + | 4.36 | 7.50E-05 | 1.01E-03 |
| **Interleukin signaling pathway (P00036)** | 94 | 19 | 4.4 | + | 4.32 | 5.92E-08 | 3.18E-06 |
| **PDGF signaling pathway (P00047)** | 140 | 25 | 6.55 | + | 3.82 | 7.35E-09 | 5.92E-07 |
| **Axon guidance mediated by netrin (P00009)** | 35 | 6 | 1.64 | + | 3.66 | 5.24E-03 | 3.52E-02 |
| **p38 MAPK pathway (P05918)** | 41 | 7 | 1.92 | + | 3.65 | 2.69E-03 | 1.88E-02 |
| **PI3 kinase pathway (P00048)** | 53 | 9 | 2.48 | + | 3.63 | 7.20E-04 | 5.79E-03 |
| **Apoptosis signaling pathway (P00006)** | 116 | 19 | 5.43 | + | 3.5 | 1.80E-06 | 4.14E-05 |
| **EGF receptor signaling pathway (P00018)** | 133 | 21 | 6.22 | + | 3.37 | 9.74E-07 | 2.61E-05 |
| **B cell activation (P00010)** | 70 | 11 | 3.28 | + | 3.36 | 3.84E-04 | 3.25E-03 |
| **VEGF signaling pathway (P00056)** | 68 | 10 | 3.18 | + | 3.14 | 1.19E-03 | 9.11E-03 |
| **T cell activation (P00053)** | 89 | 13 | 4.17 | + | 3.12 | 2.48E-04 | 2.49E-03 |
| **CCKR signaling map (P06959)** | 164 | 23 | 7.68 | + | 3 | 2.52E-06 | 5.07E-05 |
| **Ras Pathway (P04393)** | 72 | 10 | 3.37 | + | 2.97 | 1.85E-03 | 1.36E-02 |
| **Integrin signalling pathway (P00034)** | 189 | 26 | 8.85 | + | 2.94 | 8.30E-07 | 2.67E-05 |
| **Gonadotropin-releasing hormone receptor pathway (P06664)** | 235 | 31 | 11 | + | 2.82 | 1.96E-07 | 7.90E-06 |
| **FGF signaling pathway (P00021)** | 122 | 16 | 5.71 | + | 2.8 | 1.84E-04 | 2.11E-03 |
| **p53 pathway (P00059)** | 85 | 10 | 3.98 | + | 2.51 | 6.28E-03 | 4.04E-02 |
| **Angiogenesis (P00005)** | 169 | 19 | 7.91 | + | 2.4 | 3.69E-04 | 3.30E-03 |

| | | | | | | | |
|---|---|---|---|---|---|---|---|
| **Inflammation mediated by chemokine and cytokine signaling pathway (P00031)** | 259 | 26 | 12.12 | + | 2.14 | 2.72E-04 | 2.57E-03 |
| Unclassified (UNCLASSIFIED) | 19255 | 817 | 901.2 | - | 0.91 | 1.16E-14 | 1.86E-12 |

| | |
|---|---|
| Analysis Type: | PANTHER Overrepresentation Test (Released 20240807) |
| Annotation Version and Release Date: | PANTHER version 19.0 Released 2024-06-20 |
| Analyzed List: | miR-328-3p_all targets.txt (Mus musculus) |
| Reference List: | Mus musculus (all genes in database) |
| Test Type: | FISHER |
| Correction: | FDR |

| PANTHER Pathways | Mus musculus - REFLIST (21836) | miR-328-3p_all targets.txt (251) | miR-328-3p_all targets.txt (expected) | miR-328-3p_all targets.txt (over/under) | miR-328-3p_all targets.txt (fold Enrichment) | miR-328-3p_all targets.txt (raw P-value) | miR-328-3p_all targets.txt (FDR) |
|---|---|---|---|---|---|---|---|
| Unclassified (UNCLASSIFIED) | 19255 | 195 | 221.33 | - | 0.88 | 2.61E-06 | 4.21E-04 |

| | |
|---|---|
| Analysis Type: | PANTHER Overrepresentation Test (Released 20240807) |
| Annotation Version and Release Date: | PANTHER version 19.0 Released 2024-06-20 |
| Analyzed List: | miR-151-3p_all targets.txt (Mus musculus) |
| Reference List: | Mus musculus (all genes in database) |
| Test Type: | FISHER |
| Correction: | FDR |

| PANTHER Pathways | Mus musculus - REFLIST (21836) | miR-151-3p_all targets.txt (89) | miR-151-3p_all targets.txt (expected) | miR-151-3p_all targets.txt (over/under) | miR-151-3p_all targets.txt (fold Enrichment) | miR-151-3p_all targets.txt (raw P-value) | miR-151-3p_all targets.txt (FDR) |
|---|---|---|---|---|---|---|---|

#### Supplementary Data 4

**miR-203-3p**

| PANTHER Pathways | No. of pathway genes in whole genome | No. of pathway genes regulated by miR-203-3p | Names of pathway genes regulated by miR-203-3p | miR-203-3p_all targets.txt (fold Enrichment) | miR-203-3p_all targets.txt (FDR) |
|---|---|---|---|---|---|
| **Ras Pathway (P04393)** | 72 | 4 | Rps6ka3, Pdk1, Mapk8, Cdc42 | 10.19 | 0.0348 |
| **CCKR signaling map (P06959)** | 164 | 6 | Crk, Rps6ka3, Pdk1, Mapk8, Cdc42, Prkcb | 6.71 | 0.0455 |
| **Integrin signalling pathway (P00034)** | 189 | 6 | Crk, Mapk8, Cav1, Cdc42, Flnb, Itga4 | 5.83 | 0.0483 |

miR-150-5p

| PANTHER Pathways | No. of pathway genes in whole genome | No. of pathway genes regulated by miR-150-5p | Names of pathway genes regulated by miR-150-5p | miR-150-5p_all targets.txt (fold Enrichment) | miR-150-5p_all targets.txt (FDR) |
|---|---|---|---|---|---|
| **Interleukin signaling pathway (P00036)** | 94 | 9 | Pdk1, Stat5b, Elk1, Stat6, Akt1, Foxo3, Mtor, Il2ra, Srf | 6.39 | 0.000951 |
| **EGF receptor signaling pathway (P00018)** | 133 | 10 | Stat5b, Grap, Stat6, Akt1, Pik3c2a, Prkca, Ywhae, Ywhah, Cbl, Btc | 5.02 | 0.00176 |
| **PDGF signaling pathway (P00047)** | 140 | 10 | Pdk1, Elf1, Stat5b, Elk1, Grap, Stat6, Prkca, Mtor, Srf, Pdgfb | 4.77 | 0.00205 |
| **Hedgehog signaling pathway (P00025)** | 20 | 4 | Btrc, Prkar1a, Csnk1a1, Gli2 | 13.36 | 0.00637 |
| **Parkinson disease (P00049)** | 92 | 7 | Csnk1a1, Elk1, Ywhae, Ywhah, Hspa5, Ccne2, Csnk1g3 | 5.08 | 0.0125 |
| **CCKR signaling map (P06959)** | 164 | 9 | Pdk1, Mef2d, Elk1, Akt1, Foxo3, Prkca, Camk4, Srf, Acat1 | 3.66 | 0.0194 |
| **Angiogenesis (P00005)** | 169 | 9 | Apc, Grap, Akt1, Pik3c2a, F2r, Prkca, Notch4, Pdgfb, Vegfa | 3.56 | 0.021 |
| **FGF signaling pathway (P00021)** | 122 | 7 | Fgf4, Grap, Akt1, Pik3c2a, Prkca, Ywhae, Ywhah | 3.83 | 0.0393 |
| **Insulin/IGF pathway-protein kinase B signaling cascade (P00033)** | 38 | 4 | Pdk1, Akt1, Pik3c2a, Foxo3 | 7.03 | 0.0436 |

let-7e-5p

| PANTHER Pathways | No. of pathway genes in whole genome | No. of pathway genes regulated by let-7e-5p | Names of pathway genes regulated by let-7e-5p | let-7e-5p_all targets.txt (fold Enrichment) | let-7e-5p_all targets.txt (FDR) |
|---|---|---|---|---|---|
| **PDGF signaling pathway (P00047)** | 140 | 25 | Map3k4, Elp1, Arhgap5, Stat3, Pkn2, Rps6kb2, Pdpk1, Mapk1, Pik3r1, Stat5b, Gab1, Srgap2, Plcg1, Nras, Mapk6, Srgap2, Sos1, Mknk2, Jak1, Map3k2, Fli1, Mtor, Chuk, Myc, Pik3ca | 3.82 | 0.000000592 |
| **Interleukin signaling pathway (P00036)** | 94 | 19 | Il10, Map3k4, Irs2, Stat3, Pdpk1, Mapk1, Stat5b, Nras, Mapk6, Sos1, Mknk2, Il2rb, Il13, Mtor, Il12rb2, Chuk, Myc, Pik3ca, Il10rb | 4.32 | 0.00000318 |
| **Gonadotropin-releasing hormone receptor pathway (P06664)** | 235 | 31 | Vcl, Map4k3, Map3k4, Irs2, Ncoa3, Nfatc3, Stat3, Map2k4, Mapk1, Pik3r1, Sp1, Map3k1, Nab1, Skil, Dusp1, Egfr, Adipor2, Tgfbr3, Map3k3, Junb, Prlr, Map4k5, Igf1, Sos1, Map3k2, Crebbp, Egr1, Itga1, Map3k14, Acvr1b, Insr | 2.82 | 0.0000079 |
| **EGF receptor signaling pathway (P00018)** | 133 | 21 | Map3k4, Cblb, Stat3, Map2k4, Mapk1, Spry1, Stat5b, Map3k1, Gab1, Egfr, Phldb2, Plcg1, Map3k3, Nras, Sos1, Map3k2, Ppp4c, Rac2, Ywhah, Cbl, Pik3ca | 3.37 | 0.0000261 |
| **Integrin signalling pathway (P00034)** | 189 | 26 | Crk, Vcl, Map3k4, Itgal, Vasp, Map2k4, Col1a1, Itgb3, Mapk1, Pik3r1, Map3k1, Col1a2, Map3k3, Nras, Mapk6, Sos1, Col4a2, Map3k2, Itga1, Rac2, Itga4, Col27a1, Asap1, Itgb2, Col3a1, Pik3ca | 2.94 | 0.0000267 |
| **Apoptosis signaling pathway (P00006)** | 116 | 19 | Map4k3, Map2k4, Mapk1, Map3k1, Bcl2, Cflar, Igf2r, Aifm1, Map4k5, Tnfrsf1b, Map3k14, Chuk, Tnfrsf1a, Bcl2l11, Crem, Tnf, Apaf1, Casp3, Pik3ca | 3.5 | 0.0000414 |

| | | | | | |
|---|---|---|---|---|---|
| **CCKR signaling map (P06959)** | 164 | 23 | Crk, Akap1, Stat3, Calm1, Pdpk1, Mapk1, Pik3r1, Sp1, Bcl2, Ptpn11, Plcg1, Foxo1, Mob1a, Sos1, Elavl1, Egr1, Map3k14, Mmp9, Eif4ebp1, Crem, Myc, Acat1, Casp3 | 3 | 0.0000507 |
| **Oxidative stress response (P00046)** | 54 | 12 | Map3k4, Dusp4, Map2k4, Bcl2, Dusp1, Mknk2, Dusp5, Max, Dusp7, Dusp6, Dusp3, Myc | 4.75 | 0.000103 |
| **Insulin/IGF pathway-protein kinase B signaling cascade (P00033)** | 38 | 10 | Irs2, Mdm4, Pdpk1, Pik3r1, Igf2r, Foxo1, Igf1, Mdm2, Insr, Pik3ca | 5.62 | 0.000111 |
| **Hypoxia response via HIF activation (P00030)** | 29 | 8 | Pik3r1, Arnt, Egln2, Txn2, Crebbp, Mtor, Hif1a, Pik3ca | 5.89 | 0.000584 |
| **p53 pathway feedback loops 2 (P04398)** | 49 | 10 | Rb1, Atr, Mdm4, Pdpk1, Pik3r1, Nras, Ccna2, Mdm2, Myc, Pik3ca | 4.36 | 0.00101 |
| **JAK/STAT signaling pathway (P00038)** | 17 | 6 | Stat3, Stat5b, Socs1, Jak1, Pias1, Ptprc | 7.54 | 0.00102 |
| **FGF signaling pathway (P00021)** | 122 | 16 | Map3k4, Map2k4, Mapk1, Spry1, Map3k1, Ptpn11, Plcg1, Map3k3, Nras, Sos1, Map3k2, Fgf2, Ppp4c, Rac2, Ywhah, Pik3ca | 2.8 | 0.00211 |
| **Insulin/IGF pathway-mitogen activated protein kinase kinase/MAP kinase cascade (P00032)** | 36 | 8 | Irs2, Map2k4, Rps6kb2, Mapk1, Igf2r, Igf1, Sos1, Insr | 4.75 | 0.00226 |
| **T cell activation (P00053)** | 89 | 13 | Cd3e, Nfatc3, Calm1, Mapk1, Pik3r1, Map3k1, Plcg1, Nras, Sos1, Rac2, Chuk, Ptprc, Pik3ca | 3.12 | 0.00249 |
| **Inflammation mediated by chemokine and cytokine signaling pathway (P00031)** | 259 | 26 | Tyk2, Map3k4, Ccr8, Itgal, Pak4, Nfatc3, Nfat5, Stat3, Pdpk1, Mapk1, Abcd1, Plcg1, Junb, Nras, Ccr7, Ifnar1, Sos1, Myh9, Rgs4, Rac2, Il13, Itga4, Chuk, Itgb2, Pik3ca, Il10rb | 2.14 | 0.00257 |

| | | | | | |
|---|---|---|---|---|---|
| **B cell activation (P00010)** | 70 | 11 | Nfatc3, Calm1, Mapk1, Map3k3, Nras, Sos1, Map3k2, Rac2, Chuk, Ptprc, Pik3ca | 3.36 | 0.00325 |
| **Angiogenesis (P00005)** | 169 | 19 | Crk, Apc2, Stat3, Map2k4, Mapk1, Pik3r1, Map3k1, Ptpn11, Plcg1, Nras, Mapk6, Notch2, Sos1, Jak1, Fgf2, Hif1a, Vegfa, Kdr, Pik3ca | 2.4 | 0.0033 |
| **PI3 kinase pathway (P00048)** | 53 | 9 | Rps6kb2, Ccnd2, Pdpk1, Pik3r1, Nras, Foxo1, Sos1, Insr, Pik3ca | 3.63 | 0.00579 |
| **VEGF signaling pathway (P00056)** | 68 | 10 | Mapk1, Pik3r1, Plcg1, Nras, Mapk6, Rac2, Hif1a, Vegfa, Kdr, Pik3ca | 3.14 | 0.00911 |
| **Ras Pathway (P04393)** | 72 | 10 | Map3k4, Stat3, Map2k4, Pdpk1, Mapk1, Map3k1, Nras, Sos1, Rac2, Pik3ca | 2.97 | 0.0136 |
| **p38 MAPK pathway (P05918)** | 41 | 7 | Map3k4, Map2k4, Dusp1, Tab2, Mknk2, Rac2, Eif4ebp1 | 3.65 | 0.0188 |
| **Axon guidance mediated by netrin (P00009)** | 35 | 6 | Nfatc3, Vasp, Pik3r1, Plcg1, Rac2, Pik3ca | 3.66 | 0.0352 |
| **p53 pathway (P00059)** | 85 | 10 | Atr, Mdm4, Pmaip1, Pdpk1, Pik3r1, Mdm2, Crebbp, Thbs1, Apaf1, Pik3ca | 2.51 | 0.0404 |
